# *E2F3* amplification primes bladder cancer cells for premature mitosis

**DOI:** 10.1101/2025.09.16.676590

**Authors:** Kathryn A. Wierenga, Isabel Nieland, Qingwu Liu, Pepijn Rakers, Anita van den Heuvel, Mara Pateli, Richard Wubbolts, Frank Riemers, Saskia van Essen-Dorresteijn, Elsbeth van Liere, Bart Westendorp

**Author notes:** Correspondence to: Bart Westendorp. **Competing interest statements:** This work was partly sponsored by Repare Therapeutics.

## Abstract

The E2F–RB pathway controls the G1/S checkpoint, and tumors often bypass it through *RB1* or *CDKN2A* loss. Unlike these lesions, *E2F3* amplification drives persistent excessive E2F-dependent transcription through S and G2 phases. This oncogene is frequently amplified in for example bladder cancer, but its impact on the cancer cell cycle remains unclear. Using isogenic bladder cancer models and patient data, we show that *E2F3* amplification hyperactivates a mitotic gene expression program, including cyclin B1. This predisposes cells to unscheduled mitosis when the G2/M checkpoint is inhibited using the PKMYT1 inhibitor lunresertib, alone or in combination with low dosages of the WEE1 inhibitor zederosertib. *E2F3*-amplified cells acquired resistance to lunresertib by permanently reducing cyclin B1 expression, thereby preventing premature mitotic entry. Importantly, this resistance was reversed by co-treatment with a low dose of WEE1 inhibitor. These findings identify PKMYT1-dependent CDK1 inhibition as a critical safeguard against premature mitosis in *E2F3*-amplified bladder cancer. Thus, we uncover an opportunity for precision medicine strategies aimed at G2/M checkpoint inhibition to promote catastrophic mitosis in bladder cancer patients with *E2F3* amplification and excessive cyclin B1 expression.

## Introduction

In mammalian cells, the CDK-RB-E2F signaling pathway is the main driver of cell cycle entry and DNA replication. During G1 phase, phosphorylation of the retinoblastoma protein (RB) by CDK4/6-cyclin D complexes releases E2F activators, E2F1-3, allowing them to bind to the promoter of target genes and upregulate their expression ^1^. Many E2F target genes are critical for the G1/S transition, and the synthesis, replication, and repair of DNA. In addition, E2Fs activate cyclin E expression, which binds and activates CDK2. CDK2 also phosphorylates RB, thus engaging in a positive feedback loop to assure S-phase entry and progression. Hence, the CDK-RB-E2F axis is strictly regulated in normal cells to avoid unscheduled cell cycle entry and the formation of cancers. Indeed, *RB1* (which encodes the E2F inhibitor RB) and *CDKN2A* (which encodes the CDK inhibitor P16^INK4A^) are tumor suppressor genes that are commonly mutated or lost in cancer. Furthermore, genes encoding cyclin D or cyclin E proteins are amplified in many cancers, and act as oncogenes. Though amplification of genes encoding activating E2Fs is less common, *E2F3* locus amplification is still observed in many cancers. It is particularly frequent in patients with muscle-invasive bladder cancer (Figure S1A), but high incidences have also been reported in retinoblastoma and melanoma ^2-5^.

Previous work showed that knockdown of *E2F3* expression in bladder cancers with *E2F3* gene amplification inhibits DNA replication and proliferation ^6^. This indicates that ectopic E2F3 protein levels provide a growth advantage to tumor cells and thereby contribute to bladder cancer progression. Similarly, *E2F3* overexpression and *E2F3* amplification are associated with enhanced proliferation, increased malignant behavior and worse prognosis in various cancers, including bladder cancer ^5, 7-10^. Despite this large body of evidence, carefully controlled mechanistic studies using isogenic cell lines with acute *E2F3* overexpression systems are lacking. Hence, potential therapeutic vulnerabilities conferred by *E2F3* amplification may be missed. We recently showed that inducible overexpression of E2F3 in non-transformed epithelial cells caused a defect in cell cycle exit during G2 after the induction of DNA damage ^11^. This indicates that the consequences of *E2F3* amplification on proliferating cancer cells could extend beyond simply facilitating S-phase entry.Oncogene-induced accelerated S-phase entry and unconstrained E2F-dependent transcription could for example causes replication stress ^12^. DNA replication stress can be defined as events that cause stalling and/or collapse of DNA replication forks. This causes DNA damage and activation of cell cycle checkpoints. In response to endogenous or exogenous DNA damage, cycling cells have several options to delay or halt cell cycle progression. The intra-S-phase checkpoint can delay DNA replication. This checkpoint is enforced by the ATR and CHK1 kinases, which can inhibit CDK1 and CDK2 activity via phosphorylation and subsequent degradation of CDC25 phosphatases and activation of the G2 checkpoint kinase WEE1. Damaged cells can also undergo a p53-dependent cell cycle exit from G2 ^13^. Finally, the G2/M checkpoint can block unscheduled mitotic entry through inhibition of CDK1-cyclin B complexes. CDK1 activation is essential for mitotic entry and its activation can be blocked via the aforementioned WEE1 kinase as well as the related kinase PKMYT1. These kinases can thus prevent premature activation of CDK1 to avoid mitotic catastrophe ^14^.

Here, we reveal that *E2F3* amplification primes cells for unscheduled mitosis by upregulating the mitotic gene expression program, including cyclin B. This causes a heavy reliance on the PKMYT1 kinase, and we found that E2F3 amplification is synthetic lethal with the PKMYT1 inhibitor lunresertib (also known as RP-6306 and Debio-2513). We found that bladder cancer cell lines with E2F3 amplification can develop resistance to lunresertib by reducing the expression of cyclin B1, while keeping E2F3 protein levels elevated. Even though *E2F3* amplification does not sensitize cells to WEE1 inhibition, this lunresertib resistance could be overcome by combining this drug with a low dose of WEE1 inhibitor. These results suggest that PKMYT1 plays a crucial role in preventing CDK1 hyperactivation and mitotic catastrophe in bladder cancer cells with elevated expression of cyclin B1. CDK1 hyperactivation is thus a promising strategy for targeted therapy in cancers with *E2F3* amplification and/or overexpression of cyclin B1.

## Results

### E2F3 overexpression alters cell cycle progression

To study the impact of *E2F3* amplification on cancer cell proliferation we decided to use isogenic cell culture models by introducing an doxycycline-inducible overexpression system into T24 and UMUC3 bladder cancer cell lines. We also introduced this system into the non-transformed RPE1-hTert cell line, and combined this system with *TP53*^*ko*^ deletion, hereafter referred to as RPE-*TP53*^*ko*^. This enabled us to dissect the genetic interactions between *E2F3* amplification and TP53 mutation. Doxycycline treatment caused overexpression levels comparable with bladder cancer cell lines that carry endogenous *E2F3* amplifications (Figure 1A, S1A). Moreover, the *TP53* cells did not express P53 and its canonical target gene P21, even after treatment with the MDM2 inhibitor nutlin-3a (Figure S1A). We verified that the mTurq-E2F3 fusion protein localized to cell nuclei, and that its expression caused upregulation of canonical E2F target genes (Figure S1B-C).

**Figure 1.**
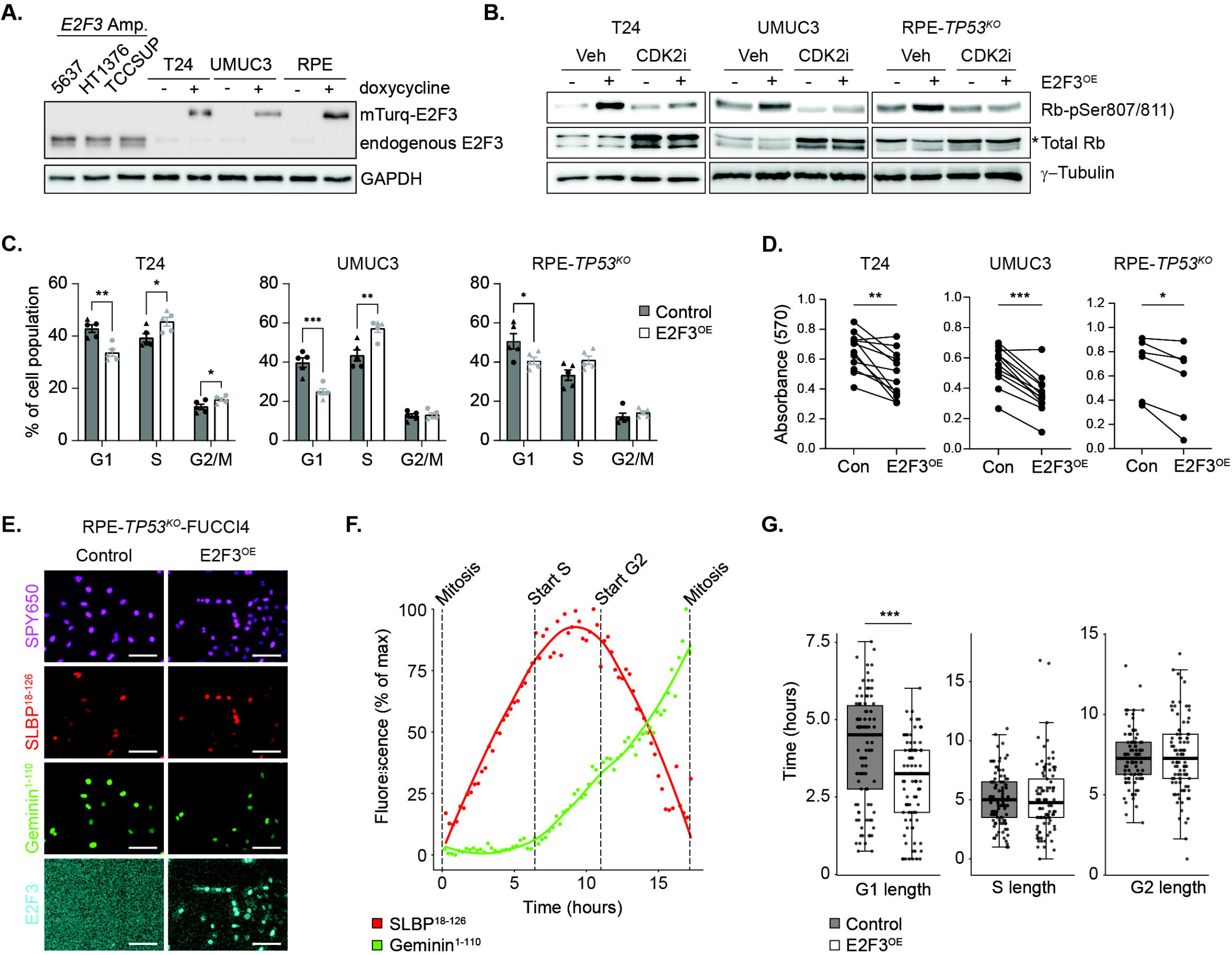
Acute E2F3 overexpression alters cell cycle progression. **A)** Immunoblot showing E2F3 expression in bladder cancer cell lines. To avoid cell cycle bias, cells were synchronized with 2mM hydroxyurea for 16 hours prior to harvesting. Doxycycline was added to T24, UMUC3, and *TP53*-proficient RPE cells 24 hours prior to hydroxyurea to induce overexpression of E2F3. Representative example of n=2 independent replicates. **B)** Immunoblot of control and E2F3^OE^ cells treated for 24 hours with doxycycline, and the last 4 hours with 1 μM of the CDK2 inhibitor INX-315; representative example of *n*=2 biological replicates. **C)** Quantification of flow cytometric cell cycle distribution analysis of cells in G1, S, and G2/M phase in control and E2F3^OE^ conditions. Each point represents an independent experiment. E2F3 overexpressing cells where treated with doxycycline for 48 hours (circles) or 72 hours (triangles). Differences between control and E2F3^OE^ cells within each cell cycle phase were analyzed using unpaired t-tests. **D)** MTT absorbance of control and E2F3^OE^ cells. Data shows samples collected from several experiments spanning 2-8 days. Each point represents an independent experiment and lines connect samples analyzed in the same experiment, meaning identical time between plating the cells and measuring MTT absorbance for the control and E2F3^OE^ groups. Differences between control and E2F3^OE^ cells was analyzed using a 2-tailed paired T-test. \**p*<0.05, \*\**p*<0.01, \*\*\*\**p*<0.0001. **E)** Fluorescence microscopy showing nuclear expression of FUCCI4 constructs and mTurqoise-tagged E2F3. E2F3-inducible UMUC3 and RPE-*TP53*^*ko*^ cells expressing the FUCCI4 markers SLBP-mKO and Geminin-mClover. SPY650 was used to track nuclei. mTurqoise-tagged E2F3 is only present in E2F3^OE^ cells. Scalebar: 50 µm. **F)** Example of mean fluorescence intensity of FUCCI4 over time in one RPE-*TP53*^KO^-FUCCI4 cell. Start of S-phase was defined as appearance of Geminin, start of G2 was defined the first 3 frames after peak fluorescence showing a continuous decrease. G1 starts the first frame after telophase (DNA decondensation), G2 ends the last frame before nuclear envelope breakdown. **G)** Quantification of RPE-TP53KO-FUCCI4 cells with and without E2F3 induction (*n*=98 and *n*=87, respectively). \*\*\**p*<0.00005.

E2F3 induction caused a marked increase in RB phosphorylation, which could be completely blocked after 4 hours of treatment with the CDK2 inhibitor INX-315 (Figure 1B). In contrast, treating the E2F3-overexpressing cell lines with CDK4/6 inhibitor palbociclib did not cause a noticeable change in RB phosphorylation (Figure S1D). This suggests that *E2F3* amplification can override its negative regulator RB via hyperactivation of CDK2. We then analyzed the impact of E2F3 expression on cell cycle progression. Flow cytometry showed that E2F3 induction caused consistent reductions of G1 populations, indicating G1-phase shortening (Figure 1C, S1E). Remarkably, this did not result in increased proliferation rates. In fact, E2F3 overexpression reduced proliferation rates in all three cell lines (Figure 1D). To study this in more detail, we inserted the FUCCI4 cell cycle reporter system ^15^ in the RPE-*TP53*^*ko*^ -E2F3 overexpression cell lines and subjected them to live cell imaging to determine duration of the different cell cycle phases (Figure 1E, F). Cell cycle tracking showed that E2F3 overexpression shortened G1 phase (Figure 1G). We did not have bladder cancer cell lines with these reporters available, but in a complementary experiment we released T24 bladder cancer cell lines from a G1 block and found that E2F3^OE^ significantly increased the percentage EdU-positive cells within 5 hours after release (Figure S1F). The same experiment in UMUC3 did not yield a significant increase, but a similar trend.

Collectively, these experiments show that E2F3 overexpression alters cell cycle progression, reduces proliferation rates, and causes hyperactivation of CDK2.

### E2F3 overexpression triggers context-dependent replication stress signaling in bladder cancer cells

A potential reason for the reduced proliferation rates in cells with E2F3 overexpression could be endogenous replication stress. In line with this hypothesis, we observed that 48 hours of E2F3 induction caused modest, but consistent activation of the replication stress markers RPA-pSer4/8, CHK1-pSer345, and KAP1-pSer824 (Figure 2A). DNA fiber analysis showed reduced replication fork speeds in UMUC3 cells and RPE-*TP53*^*ko*^ cells (Figure 2B, S2A). Strikingly, E2F3 overexpression caused increased fork speeds in T24 cells (Figure 2B). Increased fork speeds can also be linked to replication stress ^12, 16^, but these conflicting findings in the three different cell lines suggest a genetic context-dependent replication stress phenotype in cells with E2F3 overexpression. Replication stress may result in accumulation of double-stranded breaks. In line with this, we found that UMUC3 cells showed a modest but significant increase in the double-stranded break marker γH2AX (Figure 2C-D). However, the RPE and T24 cell lines did not show an increase after 48 hours of doxycycline treatment, illustrating that the replication stress caused by E2F3 overexpression is mild, and context-dependent.

**Figure 2.**
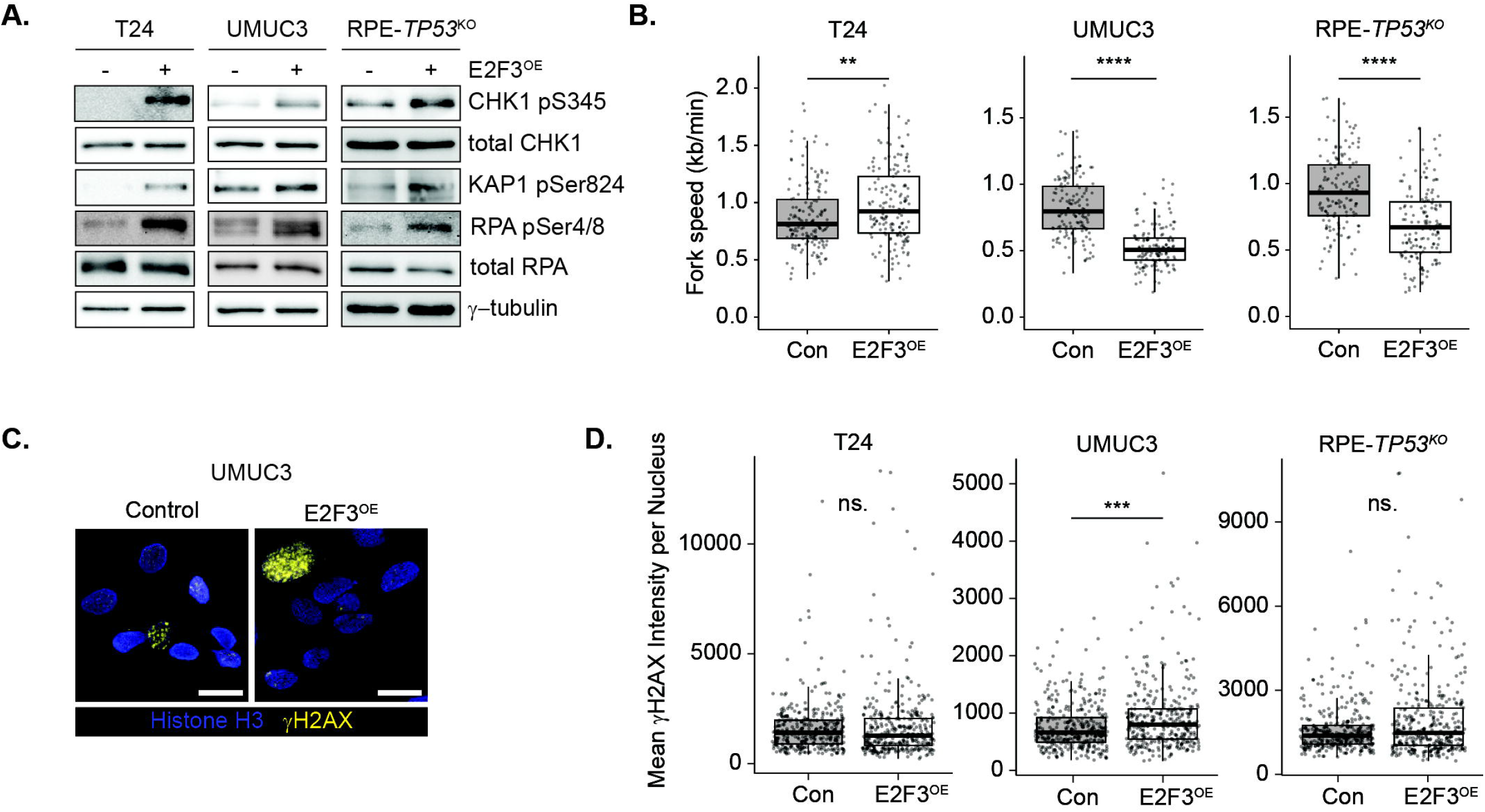
E2F3 overexpression in bladder cancer cell lines induces context-dependent replication stress signaling. **A)** Representative immunoblot of replication stress markers in control and E2F3^OE^ cells. Representative of 3 independent experiments per cell line. **B)** Boxplots depicting replication fork speeds (kb/min) of individual DNA fibers after 72 hours of E2F3 overexpression. Differences between groups determined by two-sided Wilcoxon Rank Sum test. **p<0.01, \*\*\*\**p*<0.0001 (Wilcoxon Rank Sum test). Average number of fibers quantified per condition was 140-170. Representative of 2 independent experiments per cell line. **C)** Immunofluorescence staining of γH2AX on UMUC3 cells with and without 72 hours of doxycycline-induced E2F3 overexpression. **D)** Boxplots quantifying mean γH2AX intensity per nucleus after 72 hours of E2F3 overexpression. Differences between groups determined by two-sided Wilcoxon Rank Sum test. \*\*\**p*<0.001 (Mann-Whitney Rank Sum test). Representative of 2-3 independent experiments per cell line. In all panels of this figure doxycycline treatments were maintained for 72 hours.

To complement these *in vitro* results, we investigated the impact of *E2F3* amplification on transcriptomes of bladder cancer patients using the public available TCGA dataset. First, we confirmed that patients with modest and severe copy number gains (GISTIC scores 2-7 and >7, respectively) showed an increase in *E2F3* transcript counts (Figure 3A). These groups were comparable, in that they did not show differences in tumor stage or tumor subtype (Figure 3B). We then performed differential gene expression between patients with and without *E2F3* copy number gains. To balance the numbers of patients in this analysis, we pooled patients with modest and severe copy E2F3 amplifications. Pathway analysis on genes upregulated in patients with E2F3 amplification showed that ‘E2F targets’, ‘G2-M checkpoint’ and ‘Mitotic spindle’ were among the most strongly enriched pathways (Figure 3C, Table S1). Accordingly, FOXM1 and E2F target genes topped the ranking of transcription factor target enrichment analysis (Figure 3C and Table S2). Among the downregulated genes in tumors with *E2F3* amplification we observed pathways related to metabolism and inflammation (Figure S2B and Table S2).

**Figure 3.**
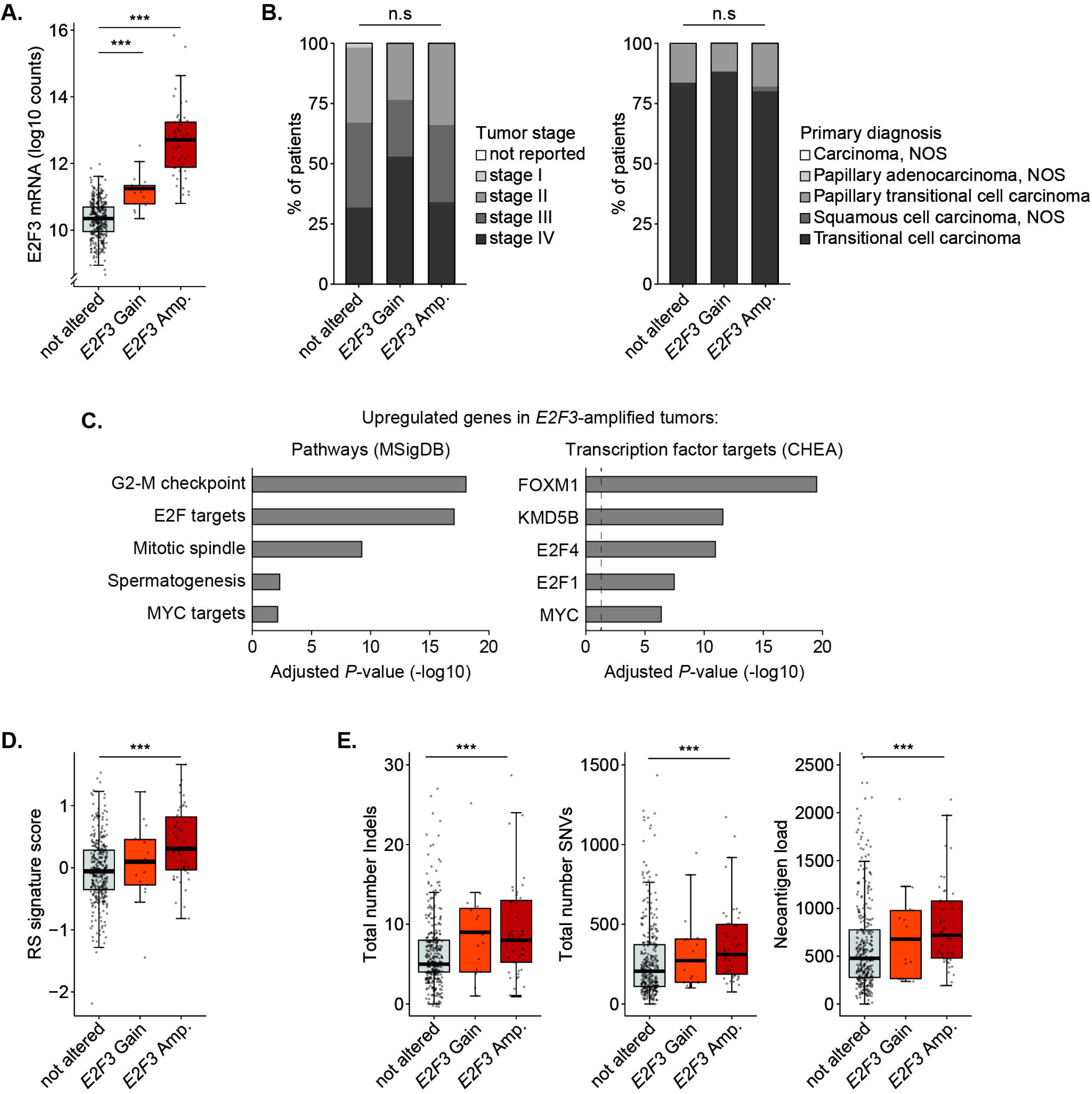
Replication stress signature expression and deregulated expression of mitotic genes in transcriptomes of bladder tumors with *E2F3* amplification. **A)** Normalized E2F3 transcript counts from RNA-sequencing data of the TCGA bladder cancer cohort. **B)** Comparison of clinical features in the TCGA bladder cancer cohort between patients with ad without *E2F3* copy number gains. **C)** Corrected *p* values of pathway enrichment analysis (left panel) and transcription factor target enrichment analysis (right panel) in genes significantly upregulated in bladder patients with *E2F3* amplification. For this statistical analysis, patients with moderate and strong *E2F3* amplifications were pooled to balance the group sizes as good as possible (*n*=61 altered versus *n*=341 unaltered patients). **D)** Replication stress signature scores in TCGA bladder cancer patients with and without *E2F3* copy number alterations. **E)** Indicators of mutation load and genetic instability in patients with and without *E2F3* copy number gains. Gain and Amp. represent groups with respectively moderate (GISTIC scores 2-7) and strong (GISTIC scores >7) copy number gains under A, B, D, and E. \*\*\**p*<0.001 as indicated.

To test whether bladder cancer patients with *E2F3* amplification show evidence of replication stress, we then analyzed expression of a published gene signature for oncogene-induced replication stress consisting of the genes *C8ORF33, DDX27, MOCS3, NAT10*, and *MPP6* ^17^. Quantitative PCR revealed that two of the 3 bladder cancer cell lines with endogenous E2F3 amplification showed significantly increased expression of these signature genes (Figure S2C). Moreover, replication stress signature scores were also increased after doxycycline-induced E2F3 overexpression (Figure S2D). Signature score analysis in TCGA bladder cancer patients indicated that replication stress was significantly increased in patients with *E2F3* copy number gains (Figure 3D). We verified that the 6 genes all contributed to the signature scores in the cancer samples (Figure S2E). This elevated replication stress may lead to enhanced genomic instability. And indeed, bladder tumors with *E2F3* amplification also showed increases in indicators of increased mutation burden, such as the numbers of indels, single nucleotide variations, and neoantigen load (Figure 3E). Altogether, these results indicate that *E2F3* amplification is associated with modest, context-dependent replication stress signaling and genomic instability in bladder cancer cells.

### E2F3 amplification sensitizes bladder cancer cells to PKMYT1 inhibition

Although the replication stress-signaling phenotypes caused by E2F3 overexpression were mild and context-dependent, we reasoned that it may render cells vulnerable to pharmacological inhibition of checkpoint kinases involved in the replication stress response. Using our inducible cell lines, we tested if E2F3 overexpression sensitizes cells to inhibition of the intra-S-phase checkpoint (ATR, CHK1) or the G2/M checkpoint (WEE1, or PKMYT1). Remarkably, the PKMYT1 inhibitor lunresertib was the only drug that was significantly more efficient in E2F3-overexpressing cells compared to cells without E2F3 induction, although all drugs inhibited their respective target kinases (Figure 4A, S3A-C). We found that E2F3 induction caused a marked increase in phosphorylation of Thr14 in CDK1 – the main target of PKMYT1-in all inducible E2F3 overexpression cell lines (Figure 4B, S3C). This elevated Thr14 phosphorylation appears to be independent from DNA damage signaling because T24 and RPE-*TP53*^*ko*^ cells did not show elevated DNA breaks (Figure 2C). We also inactivated PKMYT1 with siRNA in the E2F3 overexpression cell lines, and observed reduced colony formation of E2F3^OE^ cells compared with control cells (Figure S3D-E). Immunofluorescence staining revealed that PKMYT1 inhibition with lunresertib resulted in aberrant mitotic figures, as well as lobulated and pan-γH2AX nuclei in E2F3-overexpressing cells, indicative of mitotic catastrophe (Figure S3F-G).

**Figure 4.**
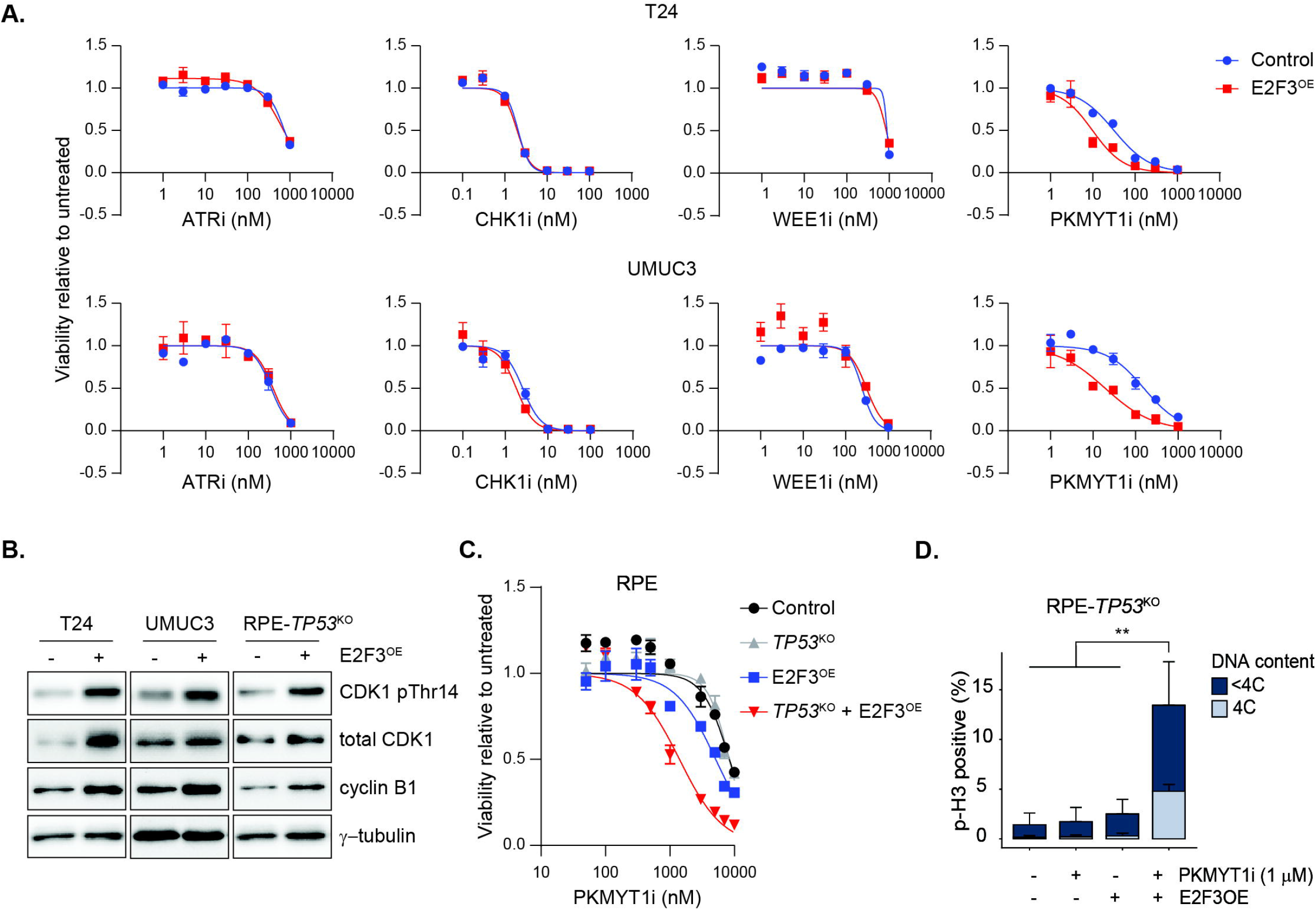
Induction of E2F3 overexpression sensitizes cells to PKMYT1 inhibition. **A)** Dose response curves (MTT assays) of DNA damage checkpoint inhibitors in control and E2F3^OE^ T24 and UMUC3 cells. E2F3 was induced 24 hours prior to the start of drug treatments, and cells were treated with the indicated drugs for 7 days. Individual points represent mean ± SEM of the percent of viable cells relative to untreated cells. Best fit curves were generated using a 4-parameter logistic curve constrained between 0 and 1. Representative examples of 3 independent experiments. The following drugs were used: ATR inhibitor ceralasertib, CHK1 inhibitor prexasertib, WEE1 inhibitor Debio 0123, PKMYT1 inhibitor lunresertib. **B)** Immunoblots showing the effect of 48 hours of doxycylin-induced E2F3 overexpression on CDK1 phosphorylation and cyclin B1 expression. Representative examples of *n*=3. **C)** Dose response curves (MTT assays) of control and E2F3^OE^ cells in WT RPE and TP53^KO^ RPE cells. Cells were treated 48 hr with lunresertib at the indicated doses. Doxycycline was added 24 hours prior to lunresertib treatment. Individual points represent mean ± SEM of the percent of viable cells relative to untreated cells. Best fit curves were generated using a 4-parameter logistic curve constrained between 0 and 1. Representative of 3 independent experiments. **D)** Quantification of flow cytometry analysis of phospho-H3 showing increased percentages of mitotic RPE-*TP53*^KO^ cells with 48 E2F3 overexpression in combination with 1 µM of the PKMYT1 inhibitor lunresertib. Bars represent mean ± SEM of 3 biological replicates. \*\**p*<0.005.

lunresertib was initially developed because PKMYT1 inactivation was found to be synthetic lethal with amplification of *CCNE1*, the gene encoding cyclin E1, in ovarian cancer ^18^. As *CCNE1* is an E2F target gene, we wondered whether *CCNE1* knockdown would rescue the increased sensitivity of E2F3-overexpressing cells to lunresertib. However, this was not the case, suggesting that E2F3 overexpression causes PKMYT1i sensitization independent from cyclin E1 (Figure S3H-I).

Our transcriptome analysis of bladder tumors with *E2F3* amplification suggested a consistent upregulation of FOXM1 target genes. This is consistent with the notion that *FOXM1* is a transcriptional target of E2F transcription factors ^19^. Accordingly, quantitative PCR showed that E2F3 overexpression consistently increased transcription of *FOXM1* and a panel of its target genes including *CCNB1* (Figure S4A-B). Moreover, protein levels of cyclin B1 were consistently increased by E2F3 (Figure 4B). This could be mechanistically relevant, because E2F3 overexpression could thus lead to elevated expression of the mitotic gene expression program, and thus make cells rely heavily on PKMYT1-dependent phosphorylation of CDK1-Thr14 to prevent premature mitosis.

One likely premise for this reliance on PKMYT1 could be the absence of functional P53 protein ^18^. Our bladder cancer cell lines all carry *TP53* mutations that impair P53 transcription factor activity, seen as inability to induce the CDK-inhibiting protein P21 in response to DNA damage (Table S3, Figure S4C). Therefore, we used our isogenic *TP53*^*KO*^ and *TP53*-proficient RPE cells. Cell viability assays unequivocally demonstrated that E2F3 overexpression only sensitizes RPE cells to PKMYT1 inhibition only when *TP53* was mutated (Figure 4C). Flow cytometry showed that PKMYT1 inhibition forced these *TP53*^*KO*^ E2F3-overexpressing cells into premature mitosis (Figure 4D, S4D).

Collectively, these data suggest that combined *TP53* mutation and *E2F3* amplification sensitizes cells to PKMYT1 inhibition. Importantly, these are highly co-occurring mutations in muscle-invasive bladder cancer ^3^.

### Resistance to PKMYT1 inhibition is associated with reduced cyclin B1 expression and negative regulation of CDK1

We then treated TCCSUP and 5637 bladder cells, which carry endogenous *E2F3* amplifications, with lunresertib. MTT assays after 7 days of treatment showed that both cell lines have IC50 below 100 nM (Figure 5A). To establish that the mechanism of action of lunresertib treatment is unscheduled mitotic entry, we then synchronized TCCSUP and 5637 cells using hydroxyurea and released them into culture medium containing lunresertib (Figure S5A). After 8 hours, untreated cells were nearly all in G2, while lunresertib caused a massive increase in mitotic cells (Figure S5B-C). This effect was rescued by co-treating the cells with the CDK1 inhibitor RO-3306. These results demonstrate that bladder cancer cells with *E2F3* amplification are susceptible to undergo premature mitosis via CDK1 hyperactivation.

**Figure 5.**
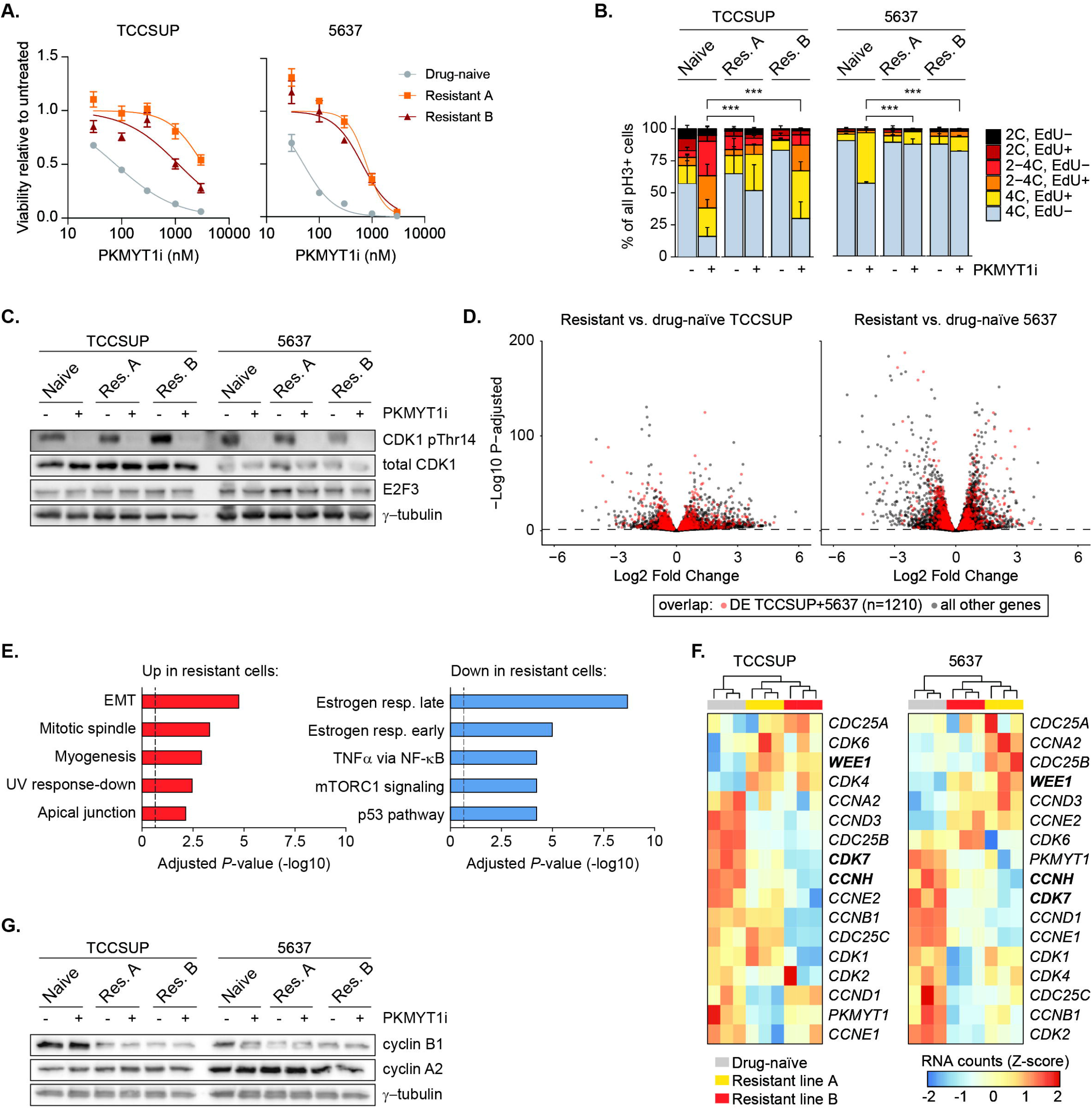
Bladder cancer cells with *E2F3* amplification become resistant to PKMYT1 inhibition via repression of cyclin B1. **A)** Dose response curves (MTT assays) of drug-naïve E2F3-amplified bladder cancer cells and cell lines made resistant to lunresertib. Resistance was achieved by gradually increasing the percent of lunresertib consistently present in the cell culture media. Individual points represent mean ± SEM of the percent of viable cells relative to untreated cells. Best fit curves were generated using a 4-parameter logistic curve constrained between 0 and 1. Representative example of 3 independent experiments. **B)** Quantification of distribution of phospho-H3 positive cells per cell cycle phase after 24 hours of PKMYT1 inhibition with 1 µM lunresertib. Drug-naïve parental cell lines are compared with lunresertib-resistant cell lines. For representative flow cytometry plots, see Figure S5B. Significance was determined with two-way ANOVA and Tukey post-hoc testing. Bars represent mean ± SEM of *n*=3 replicates. **C)** Immunoblots showing comparable levels of E2F3 protein between drug-naïve and resistant TCCSUP and 5637 bladder cancer cell lines. The PKMYT1 substrate CDK1-Thr14 was also effectively inhibited by treatment with the PKMYT1 inhibitor lunresertib (500 nM) for 4 hours. Representative of *n*=3 independent experiments **D)** Volcano plots of differentially expressed genes in bulk RNA-sequencing of asynchronously growing PKMYT1 inhibitor-resistant bladder cancer cell lines. **E)** MSigDB Pathway enrichment scores showing the 5 most significantly enriched pathways among the differentially expressed genes described in D). **F)** Heatmaps showing normalized transcript counts of a selected panel of key cell cycle regulators (cyclins, CDKs, phosphatases, and G2/M kinases). Genes in bold font were consistently and significantly up-or down regulated in all 4 resistant cell lines. G) Immunoblots showing expression of cyclin B1 and cyclin A2 in drug-naïve versus resistant cell lines. Cells were released from a hydroxyurea block and collected during G2 phase, 8 hours after release. The PKMYT1 inhibitor lunresertib (500 nM) was added 4 hours prior to cell collection to rule out cell cycle progression bias. Representative example of *n*=2 independent experiments.

We then investigated if and how resistance to lunresertib can emerge in bladder cancer cells. To this end, we made TCCSUP and 5637 cell lines resistant to lunresertib by exposing them to gradually increasing dosages of the drug in the cell culture medium. For each cell line we made two resistant polyclonal lines (hereafter referred to as A and B). All cell lines could tolerate much higher dosages of lunresertib than the parental drug-naive cell lines (Figure 5A). This resistance was retained for several (up to 20) passages after drug withdrawal. Flow cytometry analysis of cell cycle profiles showed that under normal growth conditions the resistant cell lines were nearly undistinguishable from the parental cell lines (Figure S6A-B). However, when treated with 500 nM lunresertib, the drug-naïve TCCSUP cells showed an increase in mitotic cells with <4C DNA content, which was rescued in resistant cell lines Figure S6B-C). The drug naïve 5637 cells responded less strongly to 24 hours of this dosage of lunresertib. However, all resistant cell lines consistently showed a significant rescue of the fractions of EdU-positive mitotic cells during PKMYT1i treatment (Figure 5B).

We then explored the mechanism behind the lunresertib resistance. One simple explanation could be that E2F3 expression is silenced, but this was clearly not the case in any of the cell lines (Figure 5C). Another mechanism could be enhanced drug efflux, but this was also not the case, as lunresertib still inhibited Thr14 phosphorylation of CDK1 in all 4 resistant cell lines, indicating it to be pharmacologically active in the resistant cells (Figure 5C).

Next, we performed RNA-sequencing on untreated resistant versus naïve cell lines to study how gene expression profiles are altered. We found 1207 genes that were consistently up- and downregulated in all four resistant cell lines (Figure 5D). Pathway analysis showed that genes related to the terms “epithelial-to-mesenchymal transition” and “mitotic spindle” were upregulated (Figure 5E, Table S4). Protection of the mitotic spindle could be a mechanism to protect cells from PKMYT1i-induced mitotic catastrophe, but inspection of the upregulated genes in this pathway did not yield genes that could be related to such a protective function (Table S5). We then took a more supervised approach and analyzed the expression of the main cell cycle cyclins, CDKs and CDK1 regulators in our RNA-seq dataset (Figure 5F). In both TCCSUP and 5637 cell lines, the drug-resistant cell lines clustered separately from the parental cell lines, mainly due to downregulation of different cyclins or CDKs. *CDK7* and *CCNH* were significantly decreased in all resistant cell lines. Together, these genes form the CDK-activating complex (CAK), which facilitates the activation of cyclin B1-CDK1 complexes via phosphorylation of Thr161 on CDK1 ^20^. This could potentially offer a mechanism to counteract CDK1 hyperactivation. However, phosphorylation of Thr161 on CDK1 was not reduced in the untreated resistant cell lines (Figure S6C). Interestingly, PKMYT1 inhibition reduced Thr161 phosphorylation in the resistant 5637 cells but not it resistant TCCSUP cells (Figure S6C).

The RNA-sequencing also revealed modest but consistent increases of *WEE1* transcripts in the resistant cell lines (Figure 5F). WEE1 can phosphorylate CDK1-Tyr15 to inhibit CDK1 activity. However, immunoblotting revealed that levels of total WEE1, active WEE1, and phosphorylation of CDK1 on Tyrosine 15 were not consistently elevated in the lunresertib-resistant cell lines compared to the drug-naïve parental cell lines (Figure S6C).

Another way how resistant cells could prevent mitotic CDK1 hyperactivity is by reducing cycling B levels. RNA-sequencing showed that *CCNB1* transcripts were reduced in 3 of the 4 lunresertib-resistant cell lines (Figure 5F). We then analyzed cyclin B1 protein levels in resistant cell lines. To avoid potential cell cycle bias, we synchronized the cells with hydroxyurea, and analyzed proteins 8 hours after release. We observed consistently reduced cyclin B1 levels in all resistant cell lines (Figure 5G). This reduction cannot be explained by enhanced degradation via the anaphase promoting complex/cyclosome (APC/C) E3 ligase complex, as cyclin A2 levels were not altered. Cyclin A2 is also a canonical APC/C substrate.

We then hypothesized that acute depletion of cyclin B1 would desensitize cells towards PKMYT1 inhibition. To test this, we transfected T24- and UMUC3-E2F3OE cells with siRNA against CCNB1. The siRNA caused highly effective knockdown, resulting in a reduction in cell proliferation under otherwise unperturbed conditions (Figure 6A-B). Nonetheless, PKMYT1 inhibitor treatment had far less impact on cell viability when CCNB1 was knocked down (Figure 6B).

**Figure 6.**
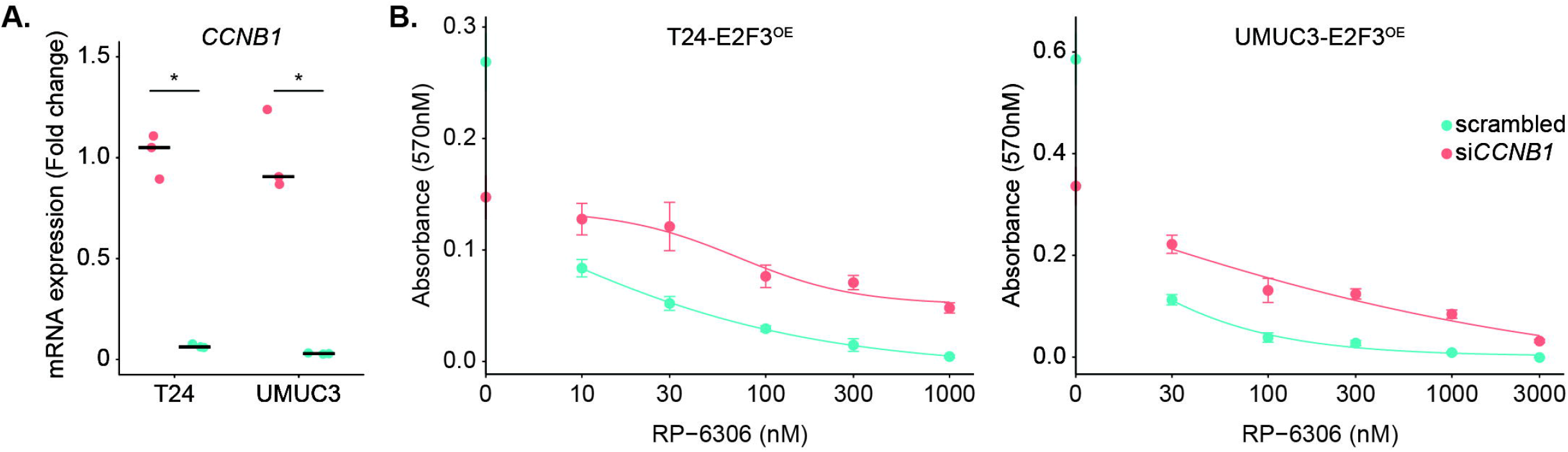
Reduced sensitivity of E2F3-overexpressing bladder cancer cell lines to PKMYT1 inhibition after cyclin B1 knockdown. **A)** Quantitative PCR of *CCNB1* expression 24 hours after siRNA transfection. \**p*<0.05 **B)** Dose-response curves to PKMYT1 inhibition with lunresertib in cells 96 hours after transfection with *CCNB1* or scrambled non-targeting siRNA. lunresertib was added during the last 72 hours. Doxycycline to induce E2F3 was added to the cells 24 hours prior to siRNA transfection. Dots represent mean ± SEM of n=3 replicates. Best fit curves were generated using a 4-parameter logistic curve constrained between 0 and 1.

**Figure 7.**
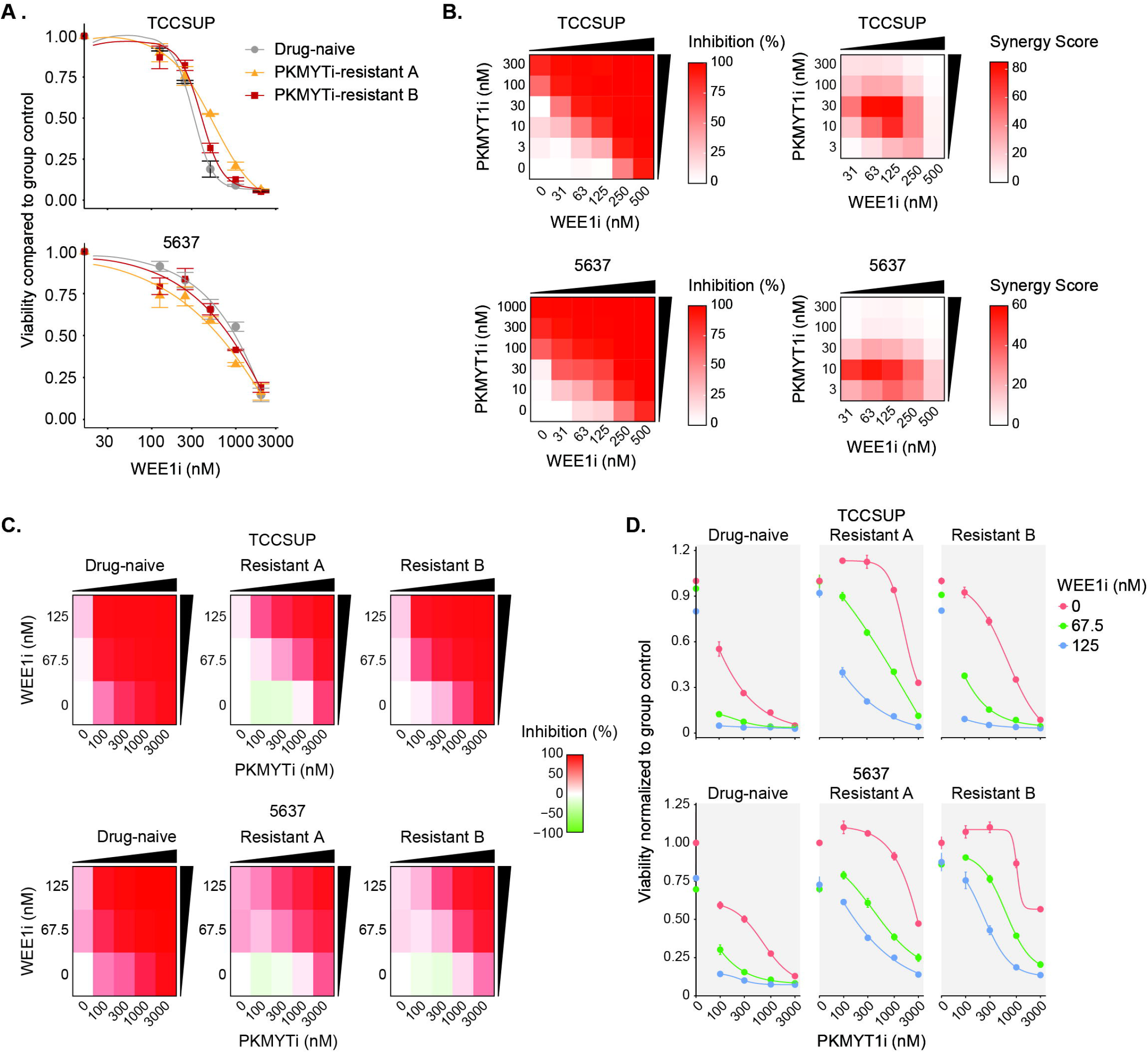
Additional treatment with a WEE1 inhibitor overcomes resistance to PKMYT1 inhibition. **A)** MTT viability assay for monotherapy with the WEE1 inhibitor zedoresertib in RP6306-resistant *E2F3*-amplified cell lines. Data are mean ± SD (*n*=3). **B)** Viability and synergy matrices showing synergistic effects of lunresertib and zedoresertib at low nano-molar doses in E2F3-amplified bladder cancer cell lines. Synergy is represented by the ZIP synergy score. Each value represents the average of *n*=3 technical replicates. **C)** MTT viability matrices for lunresertib and zedoresertib combination treatment showing mean percentage inhibition per combination. Each value represents the average of *n*=3 technical replicates. **D)** MTT viability assay dose response curves for combination treatment of lunresertib with the WEE1 inhibitor zedoresertib in lunresertib-resistant *E2F3*-amplified cell lines. Data are mean ± SD (*n*=3). Best fit curves were generated using a 4-parameter logistic curve constrained between 0 and 1.

Together, these results suggest that bladder cancer cells with *E2F3* amplification can become resistant to PKMYT1 inhibitors by reducing cyclin B1 expression to prevent premature CDK1 activation and mitosis entry.

### WEE1 inhibition overcomes PKMYT1 inhibitor resistance in E2F3-amplified bladder cancer cells

Although WEE1 activity was not elevated in lunresertib-resistant cell lines, this kinase can be expected to compensate for inactivation of PKMYT1, as both kinases catalyze inhibitory phosphorylations on CDK1. WEE1 and PKMYT1 inhibitors were previously found to act synergistically in ovarian cancers with cyclin E overexpression ^21^. Therefore, we asked whether WEE1 inhibition could overcome lunresertib resistance in bladder cancer cells with *E2F3* amplification.

Our WEE1 inhibitor of choice was the novel, highly selective compound zedoresertib (also known as Debio-0123) ^22, 23^. Zedoresertib is currently being evaluated in combination with lunresertib in a clinical trial (NCT04855656). Dose-response curves for zedoresertib monotreatment were identical in resistant and naïve cell lines, indicating that sensitivity of the lunresertib-resistant cell lines to this WEE1 inhibitor was not altered (Figure 6A). Next, we found that drug-naive TCCSUP and 5637 cells were synergistically killed at very low dosages of lunresertib and zedoresertib (Figure 6B). Importantly, all our lunresertib-resistant cell lines were also highly sensitive to a low dosage of 125 nM zedoresertib in combination with lunresertib (Figure 6C-D, S6E). This low dose of WEE1i did not cause significant cell death by itself. Together, these data show that a combination of WEE1 inhibition and PKMYT1 inhibition is very toxic to E2F3-overexpressing bladder cancer cells, even when these cells are initially made resistant to PKMYT1 inhibitor monotreatment. Thus, additional WEE1 inhibition may overcome resistance to PKMYT1 inhibition in bladder cancer cells with *E2F3* amplification.

## Discussion

The oncogenes *CCNE1* and *E2F3* engage in positive feedback, allowing cells to enter S-phase. However, this does not mean that the physiological consequences of amplification of these genes in cancer cells is interchangeable. Whereas cyclin E1 is a strong inducer of replication stress ^24-26^, our work now shows that E2F3 overexpression causes only mild and context-dependent replication stress signaling, which is insufficient to sensitize cells to ATR or CHK1 inhibition. An explanation for this difference could be that excessive cyclin E1 primarily causes hyperactivation of CDK2 and origin firing, while overexpression of the transcription factor E2F3 causes a broader hyperactivation of genes encoding DNA replication and metabolism proteins. Yet, despite absence of severe replication stress and DNA damage, E2F3 overexpression strongly increases PKMYT1-dependent inhibitory phosphorylation of CDK1 on Thr14. Our findings thus challenge the existing idea that replication stress is a prerequisite for a cell killing effect of lunresertib ^27^. It remains to be uncovered how elevated E2F3 can activate *PKMYT1*, but this could be achieved at least in part through direct transcriptional activation, as the proximal promoter of *PKMYT1* contains E2F binding sites, and chromatin-immunoprecipitation data from the ENCODE consortium show binding of E2F1 and E2F4 to this promoter ^28^.

Although we cannot not exclude contributions from a broader panel of FOXM1-associated mitosis regulators, our results strongly suggest that high levels of the FOXM1 target cyclin B1 are a prerequisite to drive premature mitosis in absence of PKMYT1. Conversely cyclin B1 downregulation and potentially rewiring of other factors controlling CDK1 activity was the most prominent putative resistance mechanism om PKMYT1i-resistant bladder cancer cell lines. The regulation of cyclin B1 protein activity during G2 phase is complex, involving for example phosphorylation and nuclear shuttling ^29, 30^. Nevertheless, recent work showed that E2Fs and CDK2 activate FOXM1 to promote cyclin B1 expression and facilitate the G2/M transition ^31^. This supports the notion that the mere abundance of cyclin B1 is important for cell fate decisions during G2 phase. It will be interesting to evaluate if cyclin B1 overexpression by itself is enough to prime for premature mitosis as E2F3 does, and if so, whether it would be a biomarker for sensitivity to PKMYT1 inhibition therapies.

PKMYT1 is not essential for survival of unperturbed cells ^32^. This makes it an attractive candidate for personalized cancer treatment. Our work shows that cells with combined *TP53* mutation and *E2F3* amplification are strongly sensitized to PKMYT1 inhibition compared with cells lacking both or either of these two mutations. There is substantial co-occurrence of these two genetic alterations; it is for example seen in ∼10-12% of patients with muscle-invasive bladder cancer ^3^. Currently, the PKMYT1 inhibitor lunresertib is being evaluated in clinical trials ^27^, and it will be of interest to explore its efficacy in bladder cancer patients with *E2F3* amplification. Mounting evidence shows that lunresertib acts synergistically with gemcitabine and cisplatin ^18, 33^, which are standard-of-care drug for patients with muscle-invasive bladder cancer ^34^.

The combination of PKMYT1 inhibition and WEE1 inhibitors holds particular promise to cause a strong synergistic cell killing of bladder cancer cells with low toxicity on non-tumor cells. Interestingly, PKMYT1 localizes to the cytosol, and WEE1 to the nucleus ^35, 36^. As CDK1 shuttles rapidly between cytosolic and nucleus ^37^, the dual inhibition of PKMYT1 and WEE1 fits with a conceptual model in which CDK1 pools are hyperactivated regardless of its subcellular localization, and thus higher propensity to undergo mitotic catastrophe. Strong synergy was also reported in ovarian cancer cells ^21^. Furthermore PKMYT1 upregulation was found to confer resistance to WEE1 inhibitors ^38^. We now show the opposite, namely that WEE1 inhibition can overcome resistance to PKMYT1 inhibitor treatment. Surprisingly, WEE1 activity was not overtly increased in our lunresertib-resistant cell lines, but his may be explained by the notion that the experiments were done in non-DNA damaging growth conditions.

Finally, combinations of PKMYT1 inhibition with immunotherapy will be worth exploring, as mitotic defects caused by PKMYT1 inhibition could potentially exacerbate genomic instability and increase neo-epitope formation in bladder cancer cells that survive lunresertib treatment. This could in turn trigger improved immune responses in patients treated with immune checkpoint inhibitors, although the predictive value of immunotherapy response is nuanced and context-specific ^39^. Nevertheless, exploring this strategy in bladder cancer with *E2F3* amplification is of particular relevance. A subset of bladder cancer patients can be successfully treated with immunotherapy ^40^ and the *E2F3* oncogene by itself already causes enhanced genomic instability and neo-epitope burden. Concluding, our work shows that *E2F3* gene amplification primes bladder cancer cells for premature mitosis, which can be leveraged to design precision medicine strategies aimed at inducing mitotic catastrophe in cancer patients carrying this genetic alteration.

## Methods

### Cell culture and cell line engineering

HEK293T, TCCSUP, HT1376, 5637, HT1197, T24, UMUC3, SW780, RT4, and hTERT RPE1 cell lines were purchased from ATCC. HEK293T and hTERT RPE1 cells were cultured in DMEM (41966052, Thermo); TCCSUP, HT1376, HT1197 and UMUC3 cells were cultured in MEM (10370088, Thermo) supplemented with L-glutamine and sodium pyruvate; 5637 cells were cultured in RPMI-1640 (21875-034, Gibco); SW780 cells were cultured in Leibovitz’s L-15 medium (30-2008, ATCC); and RT4 and T24 cells were cultured in McCoy’s 5A medium (36600088, Thermo). All the forementioned media contain 10% fetal bovine serum (10500064, Life Technologies) and 1% penicillin-streptomycin (Gibco, 15140-122). All cells were cultures at 37°C, 5% CO2. Cell lines were regularly tested for mycoplasma.

T24, UMUC3, and RPE1-hTert cell lines containing the Tet Repressor, E2F3, and the FUCCI4 system were created using lentiviral transduction with the third-generation lentiviral packaging system as previously described ^41^. Briefly, HEK293T cells were transfected with 9 μg lentiviral packaging plasmids and 9 μg constructs of interest using 90 μg PEI (Polyethylenimine, 23966). Additionally, *TP53* was knocked out in RPE1-hTERT cells with FUCCI4 and inducible E2F3 using CRISPR/Cas9 genome editing. Briefly, cells harboring the FUCCI4 system were transduced with lentiviral plasmids containing a Cas9-gRNA construct for the knock-out (sgTP53_F1: CACCGCAGAATGCAAGAAGCCCAGA, sgTP53_R1: AAACTCTGGGCTTCTTGCATTCTG). Following transfection, p53 knockout cells were selected using Nutlin. Single cell clones were generated and indels in the expected genomic location were confirmed by Sanger sequencing three different clones were selected for further analysis. Clones were validated based on p53 protein levels and sanger sequencing. For all cell lines, E2F3 overexpression was induced by adding 200 ng/ml doxycycline (D9891, Sigma Aldrich).

The *E2F3*-amplified bladder cancer cell lines TCCSUP and 5637 were made resistant to RP6306 by gradually increasing the concentration of the inhibitor in the cell culture media until cells continued to replicate in the presence of 1000 nM RP6306 (a lethal dose in the parental cell line). For both cell lines, clone A was initially treated with 500 nM, which killed 80-90% of cells. The remaining cells grew into large colonies even in the presence of the drug. After ∼2 weeks of growing in the presence of 500 nM RP6306, the dose of the drug was increased to 750 nM. When cells grew stably at 750 nM (1-2 weeks), the dose was increased a final time to 1000 nM. Clone B was generated in a similar manner, but the starting concentration was lower (50 nM). When cells grew stably at 1000 nM, RP6306 was removed and cells cultured in drug-free media unless otherwise noted.

### RNAi transfections

For siRNA experiments, cells were transfected with a final concentration of 10Cnm siRNA targeting the gene of interest or a scrambled control using Lipofectamine RNAiMAX according to manufacturers’ instructions (Life Technologies, 13778030). *CCNE1, PKMYT1*, and *CCNB1* knockdown experiments were done with smartpools of 4 siRNAs targeting each respective gene. Knockdown of the gene of interest was confirmed by qPCR. All siRNAs were purchased from Dharmacon. Transfection medium was replaced after 16-24 hours.

### Live cell imaging

RPE-*TP53*^*ko*^ -E2F3 cells expressing the FUCCI4 system were plated at 500 cells/well in 18-chambered coverglass plates (CellVis, C18SB-1.5H). Doxycycline (200 ng/mL) was added 6 hours prior to the start of imaging and SPY650 (Spirochrome, 1:1000) was added 2 hour prior to imaging to label all nuclei for cell tracking. A gas-permeable film (IbidiSeal 25×52, 10873) was used to seal the plate to prevent evaporation of media in the imaging chamber. Images were acquired using a 20x objective (UPLXAPO, NA 0.8) on an Evident SoRa W1 Confocal Spinning disk microscope (SR10, Evident, Leidendorp, NL) equipped with a Orca Fusion sCMOS camera (Hamamatsu) that was exposed for 250 ms for the channels in the live cell recording. Focus was maintained by the hardware focus ZDC system. Fluorescence emission was collected by sequential illumination with 640, 561 and 488 nm lasers set to 20% through a quad band dichroic (D405/488/561/640) and emission filters B685/40, B607/37 and B525/50 for the channels for SP650, mKO-SLBP_18-128_ and mClover-Geminin_1-110_ respectively. After time series recording, an end stage recording was performed to detect E2F3-CFP (with excitation of 455nm at 38% laserpower, dichroic D445/514/640 and emission filter 482/35 nm) with an exposure of 500 ms. Intervals were 15 minutes for 48 hours. Cell tracking and calculation of fluorescent intensity was performed in an automated manner using the FIJI plugin Trackmate ^42^ after detecting nuclei instances using Stardist. Cell cycle durations were calculated and plotted in Rstudio using the packages dplyr, ggplot2, and Gridextra. The start of G1 was defined as first frame after DNA decondensation. S-phase entry was defined as the 2^nd^ of the three consecutive frames in which Geminin signal was higher than 5% of maximum intensity for the first time after the start of G1. G2 phase entry was defined as the 2^nd^ of 3 frames showing consecutive drops in SLBP expression after the signal had reached its maximum.

### Immunoblotting

Cells were washed with ice-cold PBS and lysed in ice-cold RIPA-buffer (50⍰nM Tris-HCl pH 7.5, 1 ⍰mM EDTA, 150 ⍰mM NaCl, 0.25% deoxycholate, 1% NP-40) supplemented with NaF (1⍰mM), NaV_3_O_4_ (1⍰mM) and protease inhibitor cocktail (11873580001, Sigma Aldrich). A BCA assay was performed to measure protein concentration. Then, each protein sample was subjected to SDS-PAGE on 7.5%, 10%, or 12% polyacrylamide gels depending on the protein being evaluated. Proteins were transferred into PVDF membranes using a semi-dry Trans-Blot Turbo system according to the manufacturers protocol (Bio-Rad). The membranes were blocked in 5% BSA (for antibodies against phospho-proteins) or 5% milk (for antibodies against all other proteins) in TBS-T for an hour. Blots were then incubated with primary antibody overnight at 4°C (antibodies and dilutions are listed in Table S6). After being washed 3×10mins in TBS-Tween, blots were incubated with HRP-linked secondary antibody (diluted 1:3000) for 1 hour at room temperature. Blots were then washed 3×10mins in TBS-Tween and imaged using an ECL detection system with SuperSignal West Pico PLUS Chemiluminescent Substrate or SuperSignal West Femto Chemiluminescent Substrate (Thermo Fisher Scientific).

### Flow cytometry

Cells were lifted by trypsinization and the supernatant was carefully removed by inverting the tubes, leaving only the cell pellet. The cell pellet was resuspended in 180 μL of 0.1% PBS-BSA. Samples were then transferred to round bottom plates for fixing, permeabilization, and staining. Cells were fixed with 4% paraformaldehyde (PFA) for 30 minutes at room temperature. Cells were then washed with 0.1% PBS-BSA. For intracellular staining, cells were permeabilized in 0.2% Triton-X in PBS for 30 minutes, washed 1x 0.1% PBS-BSA, and incubated for 1-2 hour at room temperature with primary antibody diluted in 0.1% PBS-BSA. Cells were then washed 1x 0.1% PBS-BSA and incubated 1 hour with a fluorescent-labeled secondary antibody. (Antibodies and dilutions are listed in Table S6). For nuclear staining, DAPI (2μg/1E6 cells) was added and samples were incubated for 30 minutes.

For EdU incorporation assays, cells were pulse-labeled with 10 μM EdU for 90 minutes prior to collection. EdU staining was carried out according to the manufacturer’s instructions in the Click-iT Plus EdU Alexa Fluor 647 assay kit (C10634, Thermo Fisher).

For PI staining, cells were fixed with 1x PBS containing 70% Ethanol in 4° overnight. Cells were washed twice with ice cold 1x TBS and resuspend with propidium iodide (PI, P4170, Sigma Aldrich) staining buffer (20 μg/ml PI, 250 μg/ml RNase A and 0.1% BSA). PI-stained cells were not stained with additional antibodies.

For all experiments, data were collected using a Beckman Coulter CytoFlex LX flow cytometer, and analyses were performed using FlowJo v10 software (BD Life Sciences).

### Immunofluorescence staining

Cells were seeded on 13mm glass coverslips (Boom, LLG9161065) disinfected with 70% ethanol and washed 2x PBS prior to transfer in 6-well adhesion plates. Cells were immediately treated with doxycycline. After 72 hours coverslips were fixed using 4% paraformaldehyde (PFA in 0.1 M PBS) for 20 minutes, washed twice with PBS, and permeabilized with 0.1% Triton-PBS for 10 minutes.

Coverslips were transferred to 12-well plates for immunofluorescent staining. Coverslips were blocked with 5% goat serum in 0.1% PBS-Tween20 for 30 minutes at room temperature to prevent nonspecific binding. Following blocking, cells were incubated with primary antibodies (dilutions listed in Table S6) for 2 hours at room temperature. UMUC3 cells were additionally stained with an antibody against histone H3 to label nuclei. After incubation, coverslips were washed twice with PBS and incubated with secondary antibodies (dilutions listed in Table S6) for 1 hour at room temperature. Coverslips were then washed 2x with PBS. Samples were then stained with DAPI (at 0.3 ug/ml) for 10 minutes, unless H3 staining was used instead of DAPI to label nuclei. Coverslips were mounted on slides using 10^μ^L of FluorSave Reagent (Millipore, 3237994). Slides were analyzed on a Leica SP8 X confocal microscope using a 40X objective. Intensity of the yH2AX per nuclei was determined with a custom-made Fiji script. For each condition at least 100 cells were quantified.

### Fiber assays

Cells were pulsed with CIdU (50 μM, Sigma) for 20 minutes, washed 3x with PBS, and subsequently pulsed with IdU (250 μM, Sigma) for 20 minutes. DNA fibers were prepared as previously described ^43^. Briefly, cells were collected and lysed with spreading buffer (0.5% SDS, 200 mM Tris-HCl pH 7.4, 50 mM EDTA pH 8). The lysate was spread onto tilted slides (approximately 15°) to allow the DNA suspension to run down the slide. The slides were then fixed in methanol: acetic acid (3:1), treated with 2.5 M HCl for 75 minutes and blocked for 1 hour in blocking solution (1% BSA, 0.1% Tween-20 in PBS). Primary antibodies were applied to the slides, which were incubated for 1 hour at room temperature. After incubation, slides were fixed in 4% paraformaldehyde (PFA) and then incubated with secondary antibodies for 90 minutes at room temperature in the dark (antibodies and dilutions in Table S6). Images were obtained using a Leica TCS SP8 X confocal microscope using 63X oil objective. For each condition, approximately 140-170 fibers were quantified. The length and quantification of DNA tracks were manually analyzed with ImageJ software. The track length was calculated using the conversion factor 1 μm = 2.59 kb.

### Quantitative PCR

RNA isolation, cDNA synthesis and quantitative PCR were performed as previously described (Westendorp B, et al., 2012). Gene mRNA levels were determined using ΔΔC_t_ method for multiple-reference gene correction (*GAPDH, ACTB, RSP18* were used). Primer sequences are provided in Table S7.

### Viability and Synergy assays

Cells were plated in 96-well plate and treated as described. Treatment media was replaced with with full medium containing 5 mg/ml MTT (3-(4,5-Dimethylthiazol-2-yl, M6494, Invitrogen) for 4 hours at 37°C, 5% CO_2_. Medium containing MTT was discarded and 100 μl DMSO was added to dissolve formazan crystals. Samples were measured at the wavelength of 570 nm with a Clariostar Plus Microplate Reader. Absorbance was plotted relative to the untreated cells in each condition to determine viability. Synergy between lunresertib and zedoresertib was analyzed using the ZIP model in the R package SynergyFinder Plus version 3.4.5 ^44, 45^.

### Colony formation assays

UMUC3-E2F3 and T24-E2F3 cells were seeded in 24-well plates and immediately treated with doxycycline. 24 hours after plating, cells were transfected with 10 nM scrambled or PKMYT1 siRNA using Lipofectamine RNAiMAX (Life Technologies, 13778030), according to the manufacturer’s protocol. After 24 hours, the transfection medium was removed and replaced with medium containing doxycycline. After another 24 hours, cells were trypsinized and replated in 6-well plates at 1000 cells per well. The medium was refreshed every 2–3 days until colonies containing >50 cells were visible. For the staining procedure, the medium was removed, and cells were washed once with PBS before fixation for 5–10 minutes in 7:1 (v/v) methanol:acetic acid. After removal of the fixative and washing with PBS, cells were stained with 0.5% (w/v) crystal violet in a 1:4 methanol:Milli-Q solution for 1 hour. The crystal violet solution was then removed, and plates were rinsed under running tap water until the water ran clear. Plates were dried overnight and imaged using the Coomassie brilliant blue setting on the Bio-Rad ChemiDoc Imaging System.

### RNA sequencing

RNA was isolated from drug-naïve and lunresertib-resistant TCCSUP and 5637 cells using the QIAGEN RNAeasy kit according to manufacturers’ recommendations. Next, samples were subjected to sequencing. Quality control on the single-end reads was done with FastQC (version 0.11.9) TrimGalore (version 0.6.7) as used to trim reads based on quality and adapter presence after which FastQC was again used to check the resulting quality. rRNA reads were filtered out using SortMeRNA (version 4.3.6) after which the resulting reads were aligned to the reference genome (hg38) using STAR (version 2.7.10b) aligner. Follow-up quality control on the mapped (.bam) files was done using Sambamba (version 0.8.2), RSeQC (version 5.0.1) and PreSeq (v3.2.0). Read counts were then generated using the Subread FeatureCounts module (version 2.0.3) with the Homo_sapiens.GRCh38.106.ncbi.gtf file as annotation, after which normalization was done using the R-package edgeR (version 3.40). Raw read counts were further analysed with DEseq2 (version 1.28.0) using default analysis parameters and the differential gene expression between groups was assessed as shrunken log2 fold changes (LFC). Pathway analysis was performed on genes that were significantly changed in all drug-resistant clones compared to their drug-naïve counterparts (Table S3-4) with with Enrichr ^46^.

### Analysis of TCGA data

RNA sequencing data from the TCGA bladder cancer cohort were downloaded using the TCGAbiolinks package (version 2.26.0). Raw counts were compiled with metadata into a SummarizedExperiment object, and normalized using variance stabilizing transformation in DEseq2 version 1.38.3 for data visualization. Copy number data (GISTIC scores) were downloaded from the cBioportal website, and added as metadata to the data object. Differential expression analysis was done on raw read counts.

### Statistical analysis

Details of statistical tests and number of independent experiments are described in the figure legends. All analysis was performed using either GraphPad Prism v10 or R Studio (R version 4.2.0).

## Data accessibility

Raw and processed RNA-sequencing data from the PKMYT1 inhibitor-resistant bladder cancer cell lines are available on Gene Expression Omnibus under accession number GSE302069.

## Acknowledgement

We thank Reinier van der Linden and Stefan van der Elst (Hubrecht Institute-KNAW, NL) for assistance with FACS sorting experiments. We thank David Gallo and Gary Marshall (Repare Therapeutics) for valuable advice. Images were acquired in the Center for Cellular Imaging (CCI) at the Faculty of Veterinary Medicine Utrecht. This research was funded by grants from the Dutch Cancer Foundation (projects 11941/2018-2 and 15812/2024-1), ZonMW (91116011), NWO (OCENW.XS21.3.066) to B.W. and from the China Scholarship Counsil (201706140153) to Q.L.). We also received RP-6306 (lunresertib) and funding from Repare Therapeutics. We thank the members of the Division of Cell Biology, Metabolism and Cancer at Utrecht University for helpful suggestions and critical discussions.

## Supplemental Figure legends

**Supplemental figure S1, related to Figure 1**.

**A)** Immunoblots revealing absence of P53 protein and P21 expression in our *TP53*^KO^ RPE cell line; under control conditions or after 24 hours treatment with 1 μM of the MDM2 inhibitor nutlin-3a. **B)** E2F target gene expression after 24 hours of E2F3 overexpression, measured with qPCR. **C)** Nuclear localization of mTurq-E2F3 overexpression construct, shown by Immunofluorsence staining in T24 cells treated with doxycycline for 24 hours. **D)** Immunoblots of control and E2F3^OE^ cells treated for 24 hours with doxycycline, and the last 4 hours with 1 μM of the CDK2 inhibitor palbociclib; representative example of *n*=2 biological replicates. **E)** Representative DAPI histograms of control and E2F3^OE^ cells. E2F3 was overexpressed by 72 hr doxycycline treatment. Blue, orange, and green shading represent cells in G1, S, and G2 phase respectively, as calculated using the built-in cell cycle analysis in the FlowJo software. **F)** Quantification of EdU labeling in cells released from a G1 arrest with 1 mM palbociclib. The colors represent three different biological replicates. \**p*<0.05 as indicated.

**Supplemental figure S2, related to Figure 3**.

**A)** Boxplots depicting replication fork speeds (kb/min) of individual DNA fibers after 72 hours of E2F3 overexpression. Differences between groups determined by two-sided Wilcoxon Rank Sum test. \*\*\*\**p*<0.0001 (Wilcoxon Rank Sum test). Average number of fibers quantified per condition was 140-170. Replication experiment of Figure 2A. **B)** Corrected *p* values of pathway enrichment analysis (upper panel) and transcription factor target enrichment analysis (lower panel) in genes significantly upregulated in bladder patients with *E2F3* amplification. For this statistical analysis, patients with moderate and strong *E2F3* amplifications were pooled to balance the group sizes as good as possible (*n*=61 altered versus *n*=341 unaltered patients). **B)** Heatmap showing normalized expression of RS signature genes, determined by qPCR. Each observation represent a mean of two replicates **C)** Quantification of RS signature scores after 48 hours of doxycycline-induced E2F3 overexpression. Scores were calculated by converting normalized fold changes to z-scores, and averaging the z-scores for all 6 signature genes. \**p*<0.05, Student’s t-test **D)** Heatmap showing the normalized mRNA counts of all 6 replication stress signature genes in the TCGA bladder cancer cohort. Samples were ordered by total signature scores. For each sample, the *E2F3* and *TP53* mutation status are indicated on above the heatmap.

**Supplemental figure S3, related to Figure 4**

**A)** Immunoblots showing effective inhibition of the ATR substrate site CHK1-Ser345 by treatment with the ATR inhibitor ceralasertib (1 µM) in T24 bladder cancer cells with inducible E2F3 overexpression. Doxycyline was added for 48 hours prior to the addition of ceralasertib, which was added for 16 hours. **B)** Immunoblots showing effective inhibition of the CHK1 autophosporylation site Ser296 by treatment with the CHK1 inhibitor prexasertib (10 nM) in T24 bladder cancer cells with inducible E2F3 overexpression. Doxycyline was added for 48 hours prior to the addition of prexasertib, which was added for 16 hr. **C)** Immunoblots showing effective inhibition of the PKMYT1 substrate site CDK1-Thr14, and the WEE1 substrate site CDK1-Tyr15 by treatment with respectively the PKMYT1 inhibitor lunresertib and the WEE1 inhibitor zedoresertib in T24 bladder cancer cells with inducible E2F3 overexpression. Doxycyline was added for 48 hours. All immunoblots are representative examples of n=2 experiments in T24 cells; moreover *n*=2 parallel experiments in UMUC3 cells with inducible E2F3 expression showed identical results. **D)** Quantitative PCR of *PKMYT1* expression 24 hours after siRNA transcfection. * *P*<0.05 **E)** Colony formation assays of E2F3-overexpressing bladder cancer cell lines after siRNA-mediated knockdown of *PKMYT1*. Representative examples of n=3 replicates. **F)** Immunofluorescence staining showing abberant nuclear shapes and mitotic figures in UMUC3 cells after 48 hours E2F3 overexpression in presence of 300 nM lunresertib. **G)** Immunofluorescence staining showing abberant nuclear shapes and pan-γH2AX-positive cells in UMUC3 cells after 48 hours E2F3 overexpression in presence of 300 nM lunresertib. **H)** Dose response curves (MTT assays) of control and E2F3^OE^ cells where *CCNE1* was knocked down. Cells were treated 4 days with lunresertib at the indicated doses. Individual points represent mean ± SEM of the percent of viable cells relative to untreated cells. Best fit curves were generated using a 4-parameter logistic curve constrained between 0 and 1. Representative of *n*=3 independent experiments. **I)** Quantitative PCR showing efficient knockdown of *CCNE1* transcripts after siRNA transfection. Bars represent mean ± SEM of *n*=2 replicates.

**Supplemental figure S4, related to Figure 4**

**A)** Heatmaps showing normalized expressions of *FOXM1* and a panel of its target genes, obtained with quantitative PCR analysis. E2F3 overexpression was induced with doxycycline for 48 hours. **B)** Quantification of FOXM1 target gene signature scores. Scores were calculated by taking the mean z-scores of all genes shown in (A), for each replicate separately. \**p*<0.05, Student’s t-tests. **C)** Immunoblots of P35 and P21 in indicated cell lines, 24 hours after treatment with 100 nM gemcitabine or vehicle (DMSO). Representative example of n=2 separate experiments. **D)** Flow cytometry analysis of phospho-H3 showing increased percentages of mitotic RPE-*TP53*^KO^ cells with 48 E2F3 overexpression in combination with 1 µM of the PKMYT1 inhibitor lunresertib. Representative examples of *n*=3 replicates; quantification in Figure 4D.

**Supplemental figure S5, related to Figure 5**.

**A)** Schematic overview of experiment to study premature mitosis in synchronized bladder cancer cells. Hydroxurea (HU, 2 mM) was added to arrest cells at the beginning of S-phase, and after 16 hours the cells were released into fresh medium containing the PKMYT1 inhibitor lunresertib (1 µM), the CDK1 inhibitor RO-3306 (3 µM), or a combination thereof. **B)** Representative flow cytometry plots of phospho-H3 staining showing an increase in premature mitosis in synchronized lunresertib treated cells, which was rescued by addition of the CDK1 inhibitor RO-3306. **C)** Quantifications of B). Stacked bars represent mean percentages ± SEM. \**p*<0.05 \*\*\**p*<0.0001 determined by one-way ANOVA with Tukey’s post-hoc tests.

**Supplemental figure S6, related to Figure 5 and 6**.

**A)** Cell cycle distributions of untreated naïve and lunresertib-resistant bladder cancer cell lines with *E2F3* amplification. Stacked bars represent mean ± SD or *n*=4 replicates. Significance was determined with Pearson’s Chi-squared test. **B)** Flow cytometry analysis of EdU labeling in unsynchronized TCCSUP and 5637 bladder cancer cell lines, in presence or absence of 1 µM lunresertib. **C)** Quantification of percentage of phospho-H3 positive (i.e. mitotic) cells after 24 hours of PKMYT1 inhibition with 1 µM lunresertib. Drug-naïve parental cell lines are compared with lunresertib-resistant cell lines. For representative flow cytometry plots, see Figure S5A. Significance was determined with two-way ANOVA and Tukey post-hoc testing. Bars represent mean ± SEM of *n*=3 replicates. *** *p*<0.005 **D)** Immunoblots of asynchronously growing lunresertib-resistant TCCSUP and 5637 bladder cancer cells and the drug-naïve parental cell lines, untreated or treated for 4 hours with the PKMYT1 inhibitor lunresertib (500nM). Representative examples of *n*=2 independent replicate experiments are shown. **E)** Quantification of Figure 6D; area under the dose-response-curve plots for lunresertib with a combination of different Debio0123 concentrations. Significance was determined with one-way ANOVA and Tukey post-hoc testing. \**p*<0.05, **=*p*<0.01, ***=*p*<0.001)

## Notes

https://www.ncbi.nlm.nih.gov/geo/query/acc.cgi?acc=GSE302069

### Competing Interest Statement

This study was partly funded by Repare Therapeutics.

### Summary of Updates

We addressed a list of comments and concerns from the 3 reviewers. Major changes involve experiments with siRNA against cyclin B1 to demonstrate that low levels of cyclin B1 confer resistance to PKMYT1 inhibition. Furthermore we characterized our E2F3 overexpressing cell better, showing nuclear protein localization and activation of E2F target genes including CCNE1 and CCNE2.We also show that TP53 is transciptionally inactive in the used bladder cancer cell lines. Finally, we show changed the naming of the PKMYT1 inhibitor from RP-6306 to its more widely used name lunrasertib.

