## Supplementary material for "*E2F3* amplification primes bladder cancer cells for premature mitosis": Table S1

**Table S1** Differentially expressed genes in TCGA-BLCA with *E2F3* amplified tumors versus *E2F3* -intact tumors

| ensembl ID | gene symbol | baseMean | log2FoldChange | padj |
|---|---|---|---|---|
| ENSG00000198033 | TUBA3C | 13.201 | 4.023 | 7.23991E-07 |
| ENSG00000223564 | CYP4F32P | 35.319 | 3.715 | 5.10959E-09 |
| ENSG00000140505 | CYP1A2 | 100.533 | 3.697 | 5.16339E-09 |
| ENSG00000253898 | LINC01419 | 88.593 | 3.436 | 1.31755E-05 |
| ENSG00000228496 | | 30.418 | 3.403 | 6.31988E-07 |
| ENSG00000101180 | HRH3 | 52.000 | 3.379 | 2.45978E-12 |
| ENSG00000143194 | MAEL | 31.749 | 3.265 | 1.08915E-18 |
| ENSG00000164434 | FABP7 | 22.980 | 3.159 | 1.56842E-07 |
| ENSG00000178233 | TMEM151B | 15.309 | 3.124 | 2.10342E-32 |
| ENSG00000007350 | TKTL1 | 19.068 | 3.023 | 5.60525E-10 |
| ENSG00000155495 | MAGEC1 | 82.425 | 2.929 | 0.001347674 |
| ENSG00000198597 | ZNF536 | 33.178 | 2.915 | 2.36867E-14 |
| ENSG00000084628 | NKAIN1 | 78.720 | 2.907 | 1.89426E-28 |
| ENSG00000158164 | TMSB15A | 134.769 | 2.895 | 9.48401E-27 |
| ENSG00000253138 | LINC00967 | 225.350 | 2.884 | 1.00272E-05 |
| ENSG00000260019 | LINC01992 | 10.706 | 2.833 | 2.24707E-05 |
| ENSG00000157851 | DPYSL5 | 29.933 | 2.782 | 7.33386E-07 |
| ENSG00000166426 | CRABP1 | 41.069 | 2.756 | 1.77003E-09 |
| ENSG00000175229 | GAL3ST3 | 8.170 | 2.749 | 3.38865E-08 |
| ENSG00000260455 | NBAT1 | 8.223 | 2.729 | 1.56851E-15 |
| ENSG00000109424 | UCP1 | 19.407 | 2.700 | 4.76634E-09 |
| ENSG00000197408 | CYP2B6 | 13.714 | 2.611 | 3.8712E-06 |
| ENSG00000198914 | POU3F3 | 11.585 | 2.552 | 9.96513E-05 |
| ENSG00000140557 | ST8SIA2 | 40.829 | 2.522 | 8.69586E-11 |
| ENSG00000168269 | FOXI1 | 15.882 | 2.513 | 4.13192E-06 |
| ENSG00000279305 | | 7.750 | 2.513 | 0.000614734 |
| ENSG00000164900 | GBX1 | 13.699 | 2.513 | 4.37058E-07 |
| ENSG00000272168 | CASC15 | 297.459 | 2.505 | 1.96106E-40 |
| ENSG00000154162 | CDH12 | 110.521 | 2.411 | 0.000366917 |
| ENSG00000253998 | IGKV2-29 | 33.179 | 2.409 | 6.27404E-05 |
| ENSG00000112242 | E2F3 | 2388.387 | 2.404 | 1.151E-124 |
| ENSG00000198822 | GRM3 | 43.342 | 2.374 | 1.51218E-09 |
| ENSG00000234068 | PAGE2 | 44.682 | 2.366 | 0.017635538 |
| ENSG00000104833 | TUBB4A | 77.341 | 2.347 | 3.14878E-10 |
| ENSG00000250337 | PURPL | 24.612 | 2.341 | 0.000126028 |
| ENSG00000244094 | SPRR2F | 18.116 | 2.331 | 2.57693E-05 |
| ENSG00000226508 | LINC01918 | 5.138 | 2.324 | 3.45492E-11 |
| ENSG00000007314 | SCN4A | 23.582 | 2.288 | 2.03976E-15 |
| ENSG00000186487 | MYT1L | 6.506 | 2.279 | 2.81521E-05 |
| ENSG00000269779 | | 95.224 | 2.277 | 0.000198288 |
| ENSG00000120088 | CRHR1 | 6.890 | 2.273 | 7.07128E-07 |
| ENSG00000145996 | CDKAL1 | 1402.152 | 2.270 | 1.12005E-97 |
| ENSG00000273335 | | 414.615 | 2.258 | 1.35495E-07 |
| ENSG00000130294 | KIF1A | 370.535 | 2.249 | 6.81374E-06 |
| ENSG00000161609 | KASH5 | 10.452 | 2.223 | 3.17644E-07 |
| ENSG00000259417 | CTXND1 | 234.622 | 2.207 | 4.43686E-07 |
| ENSG00000171551 | ECEL1 | 108.865 | 2.206 | 1.87279E-10 |
| ENSG00000213420 | GPC2 | 314.735 | 2.202 | 2.27819E-22 |
| ENSG00000215475 | SIAH3 | 9.523 | 2.200 | 1.753E-09 |
| ENSG00000180660 | MAB21L1 | 38.505 | 2.197 | 4.2597E-11 |
| ENSG00000184956 | MUC6 | 24.741 | 2.194 | 3.16621E-10 |
| ENSG00000163202 | LCE3D | 14.147 | 2.173 | 0.014693027 |
| ENSG00000227932 | SELENOOLP | 13.393 | 2.170 | 0.011229221 |
| ENSG00000280171 | | 74.130 | 2.153 | 0.000569796 |
| ENSG00000258702 | | 9.554 | 2.150 | 1.20346E-07 |
| ENSG00000224927 | NDUFA5P10 | 5.907 | 2.139 | 0.012428028 |
| ENSG00000227195 | MIR663AHG | 4.232 | 2.136 | 3.04744E-05 |

| | | | | |
|---|---|---|---|---|
| ENSG00000197376 | | 6.101 | 2.132 | 2.54301E-05 |
| ENSG00000241720 | | 67.616 | 2.121 | 0.000514606 |
| ENSG00000126752 | SSX1 | 67.715 | 2.110 | 0.038334312 |
| ENSG00000145248 | SLC10A4 | 49.103 | 2.109 | 9.50424E-11 |
| ENSG00000169059 | VCX3A | 38.973 | 2.101 | 0.000349981 |
| ENSG00000156269 | NAA11 | 63.153 | 2.098 | 0.010207078 |
| ENSG00000139209 | SLC38A4 | 837.296 | 2.097 | 7.34287E-07 |
| ENSG00000260777 | | 37.720 | 2.081 | 1.59984E-06 |
| ENSG00000225792 | | 22.696 | 2.075 | 2.48021E-12 |
| ENSG00000227234 | SPANXB1 | 9.474 | 2.073 | 0.047702629 |
| ENSG00000185686 | PRAME | 732.227 | 2.056 | 0.000532451 |
| ENSG00000259342 | | 66.570 | 2.051 | 2.75786E-06 |
| ENSG00000260518 | BMS1P8 | 16.019 | 2.050 | 0.00043893 |
| ENSG00000135220 | UGT2A3 | 37.457 | 2.047 | 0.023435029 |
| ENSG00000083067 | TRPM3 | 9.318 | 2.040 | 7.1424E-11 |
| ENSG00000242766 | IGKV1D-17 | 11.435 | 2.039 | 7.71593E-05 |
| ENSG00000172201 | ID4 | 1368.124 | 2.037 | 4.61657E-15 |
| ENSG00000233723 | LINC01122 | 14.059 | 2.034 | 1.4382E-05 |
| ENSG00000233316 | DSCR10 | 3.353 | 2.030 | 0.004782983 |
| ENSG00000182674 | KCNB2 | 6.197 | 2.018 | 0.000287187 |
| ENSG00000215612 | HMX1 | 14.474 | 2.009 | 0.038996097 |
| ENSG00000228412 | LNC-LBCS | 31.800 | 1.999 | 4.64864E-09 |
| ENSG00000265060 | PPY2P | 12.675 | 1.996 | 0.00619122 |
| ENSG00000188404 | SELL | 2009.302 | 1.992 | 1.73527E-09 |
| ENSG00000166800 | LDHAL6A | 29.697 | 1.988 | 1.55478E-05 |
| ENSG00000156009 | MAGEA8 | 51.201 | 1.977 | 0.000180352 |
| ENSG00000226057 | PHF2P2 | 48.704 | 1.957 | 0.039878608 |
| ENSG00000229700 | | 5.006 | 1.947 | 4.38692E-17 |
| ENSG00000248498 | ASNSP1 | 20.032 | 1.941 | 0.007187669 |
| ENSG00000218048 | | 8.989 | 1.929 | 0.007302532 |
| ENSG00000278626 | | 11.875 | 1.920 | 8.85181E-05 |
| ENSG00000172197 | MBOAT1 | 525.982 | 1.918 | 8.92143E-23 |
| ENSG00000188322 | SBK1 | 1184.500 | 1.915 | 5.72639E-12 |
| ENSG00000279516 | FAM230C | 9.513 | 1.911 | 0.031647787 |
| ENSG00000231431 | FAR2P4 | 55.585 | 1.908 | 0.003350779 |
| ENSG00000246465 | | 52.894 | 1.907 | 8.10025E-10 |
| ENSG00000161682 | FAM171A2 | 305.109 | 1.895 | 3.75974E-20 |
| ENSG00000187621 | TCL6 | 19.708 | 1.895 | 4.57168E-06 |
| ENSG00000205231 | TTLL10-AS1 | 5.782 | 1.887 | 9.33686E-09 |
| ENSG00000174948 | GPR149 | 8.582 | 1.885 | 0.015077756 |
| ENSG00000236516 | KLF2P4 | 10.484 | 1.883 | 0.006231359 |
| ENSG00000241697 | TMEFF1 | 4.966 | 1.881 | 5.03803E-07 |
| ENSG00000132911 | NMUR2 | 6.706 | 1.870 | 2.10567E-05 |
| ENSG00000182256 | GABRG3 | 16.918 | 1.860 | 0.000416249 |
| ENSG00000143494 | VASH2 | 316.575 | 1.853 | 1.24282E-17 |
| ENSG00000253661 | ZFHX4-AS1 | 23.737 | 1.830 | 8.15387E-05 |
| ENSG00000165828 | PRAP1 | 307.802 | 1.826 | 3.27172E-05 |
| ENSG00000249948 | GBA3 | 26.593 | 1.825 | 0.003816666 |
| ENSG00000184029 | DSCR4 | 19.827 | 1.825 | 0.031636816 |
| ENSG00000105251 | SHD | 37.733 | 1.825 | 7.39748E-06 |
| ENSG00000198054 | DSCR8 | 127.955 | 1.819 | 0.030591042 |
| ENSG00000253363 | | 8.864 | 1.809 | 0.023787428 |
| ENSG00000046774 | MAGEC2 | 210.403 | 1.808 | 0.039691411 |
| ENSG00000105255 | FSD1 | 249.872 | 1.808 | 1.49993E-08 |
| ENSG00000230306 | BANF1P2 | 9.853 | 1.807 | 3.02599E-08 |
| ENSG00000104901 | DKKL1 | 16.363 | 1.807 | 2.28132E-09 |
| ENSG00000176490 | DIRAS1 | 137.448 | 1.801 | 2.68695E-07 |
| ENSG00000169862 | CTNND2 | 71.454 | 1.798 | 6.56361E-05 |
| ENSG00000205325 | | 8.609 | 1.796 | 0.028335004 |

| | | | | |
|---|---|---|---|---|
| ENSG00000251226 | LINC02714 | 14.740 | 1.789 | 0.003063537 |
| ENSG00000184330 | S100A7A | 355.474 | 1.787 | 0.023714687 |
| ENSG00000173406 | DAB1 | 377.188 | 1.785 | 7.71593E-05 |
| ENSG00000160963 | COL26A1 | 92.949 | 1.775 | 7.87259E-06 |
| ENSG00000279579 | LINC01666 | 6.523 | 1.767 | 0.000670535 |
| ENSG00000226699 | | 4.493 | 1.766 | 7.67014E-06 |
| ENSG00000196090 | PTPRT | 42.856 | 1.764 | 8.08543E-05 |
| ENSG00000238284 | LINC01448 | 3.655 | 1.756 | 0.013209266 |
| ENSG00000105642 | KCNN1 | 36.616 | 1.756 | 7.46921E-10 |
| ENSG00000174672 | BRSK2 | 95.911 | 1.755 | 1.73203E-07 |
| ENSG00000254726 | MEX3A | 2164.728 | 1.754 | 9.87467E-18 |
| ENSG00000250420 | AACSP1 | 27.468 | 1.753 | 0.000718557 |
| ENSG00000278931 | | 6.044 | 1.748 | 1.52654E-06 |
| ENSG00000125851 | PCSK2 | 13.870 | 1.746 | 0.001137275 |
| ENSG00000151640 | DPYSL4 | 212.142 | 1.744 | 2.07703E-06 |
| ENSG00000149970 | CNKSR2 | 30.047 | 1.743 | 1.873E-06 |
| ENSG00000156414 | TDRD9 | 89.370 | 1.743 | 7.94028E-05 |
| ENSG00000130876 | SLC7A10 | 21.705 | 1.737 | 3.38048E-05 |
| ENSG00000186335 | SLC36A2 | 12.770 | 1.734 | 8.91487E-05 |
| ENSG00000197410 | DCHS2 | 21.958 | 1.733 | 3.80923E-07 |
| ENSG00000135097 | MSI1 | 258.378 | 1.727 | 1.3428E-07 |
| ENSG00000224607 | IGKV1D-27 | 5.303 | 1.724 | 0.002566891 |
| ENSG00000056487 | PHF21B | 9.404 | 1.724 | 1.91387E-05 |
| ENSG00000266976 | | 23.842 | 1.719 | 0.020386414 |
| ENSG00000259129 | LINC00648 | 81.538 | 1.714 | 0.010228544 |
| ENSG00000258754 | LINC01579 | 35.478 | 1.713 | 2.07354E-05 |
| ENSG00000124557 | BTN1A1 | 6.988 | 1.712 | 1.56839E-06 |
| ENSG00000165061 | ZMAT4 | 24.430 | 1.707 | 0.00193498 |
| ENSG00000242173 | ARHGDIG | 19.675 | 1.705 | 0.000328711 |
| ENSG00000159516 | SPRR2G | 55.268 | 1.703 | 0.041356882 |
| ENSG00000272108 | | 14.461 | 1.698 | 4.39096E-09 |
| ENSG00000228420 | LINC01768 | 8.683 | 1.697 | 0.024112746 |
| ENSG00000165309 | ARMC3 | 4.640 | 1.696 | 0.000460317 |
| ENSG00000102230 | PCYT1B | 47.393 | 1.694 | 1.52654E-06 |
| ENSG00000240280 | TCAM1P | 92.257 | 1.686 | 0.000144354 |
| ENSG00000231532 | LINC01249 | 21.102 | 1.678 | 0.047912401 |
| ENSG00000129951 | PLPPR3 | 13.053 | 1.675 | 9.47934E-05 |
| ENSG00000167768 | KRT1 | 1107.528 | 1.669 | 0.004661921 |
| ENSG00000260644 | HERC2P5 | 3.365 | 1.668 | 0.001136712 |
| ENSG00000122133 | PAEP | 23.222 | 1.668 | 0.001146966 |
| ENSG00000182272 | B4GALNT4 | 910.066 | 1.668 | 2.8099E-07 |
| ENSG00000205293 | LINC01602 | 9.906 | 1.666 | 0.019603036 |
| ENSG00000171243 | SOSTDC1 | 37.781 | 1.655 | 0.000345821 |
| ENSG00000279208 | | 5.189 | 1.654 | 1.24458E-06 |
| ENSG00000226762 | LINC02668 | 7.568 | 1.653 | 0.006091081 |
| ENSG00000107018 | RLN1 | 15.513 | 1.651 | 1.88395E-08 |
| ENSG00000130226 | DPP6 | 8.623 | 1.648 | 1.41925E-05 |
| ENSG00000233358 | | 4.194 | 1.647 | 0.000317181 |
| ENSG00000242808 | SOX2-OT | 36.118 | 1.641 | 5.60525E-10 |
| ENSG00000180537 | RNF182 | 119.520 | 1.641 | 6.17803E-06 |
| ENSG00000261168 | | 10.813 | 1.639 | 2.47909E-14 |
| ENSG00000220771 | BOLA2P3 | 5.761 | 1.638 | 2.12941E-08 |
| ENSG00000233098 | CCDC144NL-AS1 | 220.913 | 1.635 | 2.37361E-06 |
| ENSG00000250519 | | 14.803 | 1.631 | 8.91487E-05 |
| ENSG00000260088 | DDX59-AS1 | 10.284 | 1.627 | 1.38232E-08 |
| ENSG00000161798 | AQP5 | 44.330 | 1.622 | 0.00114845 |
| ENSG00000100167 | SEPTIN3 | 424.112 | 1.618 | 1.4803E-08 |
| ENSG00000134760 | DSG1 | 185.311 | 1.618 | 0.000645203 |
| ENSG00000118276 | B4GALT6 | 347.721 | 1.614 | 9.48575E-14 |

| | | | | |
|---|---|---|---|---|
| ENSG00000250376 | | 6.075 | 1.612 | 0.037064237 |
| ENSG00000139200 | PIANP | 66.433 | 1.606 | 2.60228E-09 |
| ENSG00000183496 | MEX3B | 193.353 | 1.602 | 1.56851E-15 |
| ENSG00000253125 | | 12.180 | 1.601 | 0.000799739 |
| ENSG00000175497 | DPP10 | 32.229 | 1.598 | 0.028640359 |
| ENSG00000187527 | ATP13A5 | 5.226 | 1.597 | 0.004633575 |
| ENSG00000251056 | ANKRD20A17P | 2.830 | 1.596 | 0.006855881 |
| ENSG00000203585 | LINC02408 | 8.108 | 1.592 | 3.97716E-08 |
| ENSG00000106689 | LHX2 | 46.804 | 1.592 | 6.86309E-05 |
| ENSG00000154143 | PANX3 | 3.589 | 1.591 | 0.021665187 |
| ENSG00000181378 | CFAP65 | 4.291 | 1.591 | 0.000190456 |
| ENSG00000224559 | LINC01087 | 12.371 | 1.586 | 0.00397074 |
| ENSG00000090932 | DLL3 | 55.822 | 1.581 | 2.81521E-05 |
| ENSG00000077063 | CTTNBP2 | 281.896 | 1.578 | 5.33003E-06 |
| ENSG00000204969 | PCDHA2 | 7.854 | 1.576 | 0.000378018 |
| ENSG00000105278 | ZFR2 | 9.633 | 1.568 | 1.24157E-05 |
| ENSG00000266830 | | 53.807 | 1.562 | 0.025704916 |
| ENSG00000235961 | PNMA6A | 36.464 | 1.557 | 8.72265E-06 |
| ENSG00000233585 | | 9.869 | 1.553 | 4.10424E-11 |
| ENSG00000012504 | NR1H4 | 123.040 | 1.553 | 0.006916472 |
| ENSG00000069188 | SDK2 | 237.617 | 1.552 | 6.18735E-07 |
| ENSG00000125462 | MIR9-1HG | 38.731 | 1.550 | 1.71958E-05 |
| ENSG00000084710 | EFR3B | 90.553 | 1.546 | 3.31264E-13 |
| ENSG00000170523 | KRT83 | 45.830 | 1.546 | 0.000465953 |
| ENSG00000137285 | TUBB2B | 582.795 | 1.544 | 0.000140511 |
| ENSG00000027644 | INSRR | 7.310 | 1.535 | 4.82563E-05 |
| ENSG00000146166 | LGSN | 29.988 | 1.531 | 0.016900686 |
| ENSG00000137571 | SLCO5A1 | 57.246 | 1.524 | 1.39508E-06 |
| ENSG00000170558 | CDH2 | 313.717 | 1.520 | 1.40712E-05 |
| ENSG00000244142 | ATP6V0CP2 | 10.825 | 1.519 | 0.004013059 |
| ENSG00000258590 | NBEAP1 | 29.340 | 1.508 | 0.000812666 |
| ENSG00000100604 | CHGA | 34.135 | 1.508 | 0.001171176 |
| ENSG00000126950 | TMEM35A | 118.617 | 1.503 | 7.92839E-05 |
| ENSG00000227121 | LINC02672 | 71.935 | 1.502 | 0.040620344 |
| ENSG00000185818 | NAT8L | 120.216 | 1.502 | 4.5297E-05 |
| ENSG00000276966 | H4C5 | 38.042 | 1.499 | 3.02684E-08 |
| ENSG00000260230 | FRRS1L | 29.514 | 1.496 | 0.000264715 |
| ENSG00000166105 | GLB1L3 | 35.076 | 1.495 | 0.001039452 |
| ENSG00000075340 | ADD2 | 269.331 | 1.495 | 6.6039E-05 |
| ENSG00000165349 | SLC7A3 | 6.137 | 1.492 | 0.002675023 |
| ENSG00000182798 | MAGEB17 | 17.279 | 1.491 | 0.002668768 |
| ENSG00000237515 | SHISA9 | 81.876 | 1.490 | 0.016175855 |
| ENSG00000259495 | | 44.875 | 1.488 | 7.89112E-05 |
| ENSG00000164638 | SLC29A4 | 388.534 | 1.486 | 8.97701E-10 |
| ENSG00000255201 | | 7.086 | 1.484 | 1.15446E-05 |
| ENSG00000167434 | CA4 | 190.631 | 1.471 | 0.003768097 |
| ENSG00000183242 | WT1-AS | 18.755 | 1.471 | 0.004356096 |
| ENSG00000101098 | RIMS4 | 58.926 | 1.470 | 0.004412094 |
| ENSG00000169548 | ZNF280A | 8.812 | 1.468 | 0.038197356 |
| ENSG00000101489 | CELF4 | 34.498 | 1.458 | 8.22922E-07 |
| ENSG00000131409 | LRRC4B | 76.361 | 1.456 | 6.53838E-12 |
| ENSG00000064218 | DMRT3 | 14.350 | 1.455 | 0.001382159 |
| ENSG00000224885 | EIPR1-IT1 | 5.609 | 1.453 | 2.92856E-08 |
| ENSG00000271369 | | 17.224 | 1.452 | 0.004963605 |
| ENSG00000237510 | GPAT2P1 | 13.680 | 1.445 | 0.024990521 |
| ENSG00000105675 | ATP4A | 16.774 | 1.445 | 0.003117218 |
| ENSG00000250753 | | 4.940 | 1.444 | 0.035190787 |
| ENSG00000267919 | | 5.628 | 1.439 | 0.000175104 |
| ENSG00000138741 | TRPC3 | 32.008 | 1.437 | 4.76037E-08 |

| | | | | |
|---|---|---|---|---|
| ENSG00000130643 | CALY | 6.916 | 1.436 | 0.000179459 |
| ENSG00000248360 | LINC00504 | 61.046 | 1.436 | 0.003075266 |
| ENSG00000261594 | TPBGL | 86.447 | 1.433 | 7.33386E-07 |
| ENSG00000134216 | CHIA | 8.756 | 1.431 | 0.038265997 |
| ENSG00000183960 | KCNH8 | 73.617 | 1.431 | 0.000212941 |
| ENSG00000184937 | WT1 | 47.658 | 1.429 | 0.003250914 |
| ENSG00000227012 | LINC02527 | 5.367 | 1.427 | 0.003314014 |
| ENSG00000124766 | SOX4 | 11467.901 | 1.412 | 8.64972E-26 |
| ENSG00000258986 | TMEM179 | 21.851 | 1.412 | 0.007313197 |
| ENSG00000180178 | FAR2P1 | 243.731 | 1.410 | 0.010407187 |
| ENSG00000154493 | C10orf90 | 16.242 | 1.407 | 0.00475405 |
| ENSG00000185666 | SYN3 | 15.293 | 1.405 | 1.81389E-06 |
| ENSG00000141668 | CBLN2 | 18.236 | 1.400 | 0.0083842 |
| ENSG00000176826 | FKBP9P1 | 107.600 | 1.400 | 7.6118E-09 |
| ENSG00000226097 | | 15.216 | 1.399 | 0.002068673 |
| ENSG00000268460 | | 140.060 | 1.399 | 0.000568739 |
| ENSG00000231702 | | 7.216 | 1.397 | 0.000381012 |
| ENSG00000179111 | HES7 | 12.812 | 1.396 | 3.84388E-06 |
| ENSG00000267413 | LINC01901 | 8.477 | 1.396 | 0.020187611 |
| ENSG00000146648 | EGFR | 10531.890 | 1.395 | 2.0988E-08 |
| ENSG00000249790 | | 21.309 | 1.394 | 0.002272737 |
| ENSG00000126262 | FFAR2 | 89.014 | 1.391 | 4.02603E-05 |
| ENSG00000105088 | OLFM2 | 646.353 | 1.388 | 1.08691E-08 |
| ENSG00000147488 | ST18 | 12.639 | 1.387 | 3.11078E-07 |
| ENSG00000197813 | | 16.437 | 1.385 | 8.39295E-08 |
| ENSG00000099617 | EFNA2 | 27.399 | 1.384 | 0.002750999 |
| ENSG00000124194 | GDAP1L1 | 4.825 | 1.383 | 7.42139E-06 |
| ENSG00000154118 | JPH3 | 73.739 | 1.382 | 0.000807424 |
| ENSG00000185615 | PDIA2 | 53.754 | 1.382 | 6.89025E-05 |
| ENSG00000279479 | | 9.131 | 1.381 | 0.009781321 |
| ENSG00000221946 | FXYD7 | 12.847 | 1.378 | 3.57948E-06 |
| ENSG00000133665 | DYDC2 | 7.537 | 1.378 | 0.00339979 |
| ENSG00000152932 | RAB3C | 32.162 | 1.374 | 0.000152583 |
| ENSG00000142549 | IGLON5 | 29.773 | 1.373 | 1.10936E-06 |
| ENSG00000253239 | IGLVI-70 | 5.293 | 1.371 | 0.020149201 |
| ENSG00000234948 | LINC01524 | 6.277 | 1.369 | 0.01069268 |
| ENSG00000281566 | | 12.173 | 1.369 | 0.041984331 |
| ENSG00000197421 | GGT3P | 13.556 | 1.367 | 0.010207078 |
| ENSG00000272461 | | 3.455 | 1.365 | 0.004232104 |
| ENSG00000164175 | SLC45A2 | 18.645 | 1.362 | 0.000247755 |
| ENSG00000148948 | LRRC4C | 25.685 | 1.361 | 2.88639E-05 |
| ENSG00000113205 | PCDHB3 | 54.669 | 1.361 | 9.69094E-07 |
| ENSG00000162490 | DRAXIN | 82.317 | 1.359 | 1.97201E-07 |
| ENSG00000085552 | IGSF9 | 2849.537 | 1.359 | 9.08943E-12 |
| ENSG00000181291 | TMEM132E | 25.120 | 1.355 | 1.3075E-07 |
| ENSG00000132204 | LINC00470 | 72.881 | 1.350 | 0.017714386 |
| ENSG00000268686 | | 70.331 | 1.350 | 0.000887652 |
| ENSG00000264016 | | 8.189 | 1.349 | 2.84164E-05 |
| ENSG00000135374 | ELF5 | 725.108 | 1.348 | 8.97911E-05 |
| ENSG00000275383 | | 6.182 | 1.348 | 1.73527E-09 |
| ENSG00000265702 | | 11.350 | 1.347 | 1.79458E-05 |
| ENSG00000158683 | PKD1L1 | 225.013 | 1.347 | 2.07529E-05 |
| ENSG00000149599 | DUSP15 | 84.733 | 1.344 | 3.26876E-06 |
| ENSG00000100276 | RASL10A | 112.348 | 1.344 | 1.73203E-07 |
| ENSG00000081842 | PCDHA6 | 15.034 | 1.342 | 0.00545018 |
| ENSG00000237693 | IRGM | 4.666 | 1.341 | 2.37361E-06 |
| ENSG00000240975 | | 6.532 | 1.339 | 0.000405065 |
| ENSG00000250590 | LINC02492 | 6.276 | 1.335 | 0.034205796 |
| ENSG00000258081 | | 3.322 | 1.332 | 0.02332761 |

| | | | | |
|---|---|---|---|---|
| ENSG00000183798 | EMILIN3 | 36.123 | 1.332 | 4.79016E-06 |
| ENSG00000166707 | ZCCHC18 | 46.661 | 1.331 | 3.04436E-10 |
| ENSG00000175868 | CALCB | 12.135 | 1.328 | 0.001567891 |
| ENSG00000169618 | PROKR1 | 9.016 | 1.327 | 0.002248212 |
| ENSG00000135298 | ADGRB3 | 35.150 | 1.327 | 0.000797536 |
| ENSG00000063127 | SLC6A16 | 57.185 | 1.325 | 3.80923E-07 |
| ENSG00000184486 | POU3F2 | 5.105 | 1.324 | 6.77019E-05 |
| ENSG00000174460 | ZCCHC12 | 27.013 | 1.323 | 9.7035E-05 |
| ENSG00000166863 | TAC3 | 806.259 | 1.323 | 0.028879851 |
| ENSG00000082482 | KCNK2 | 29.261 | 1.319 | 0.001033577 |
| ENSG00000163673 | DCLK3 | 61.578 | 1.317 | 2.42727E-06 |
| ENSG00000234537 | | 6.956 | 1.317 | 4.39339E-05 |
| ENSG00000117598 | PLPPR5 | 7.559 | 1.314 | 0.002176091 |
| ENSG00000116983 | HPCAL4 | 22.178 | 1.312 | 0.000245125 |
| ENSG00000102935 | ZNF423 | 299.220 | 1.310 | 1.07427E-09 |
| ENSG00000205456 | TP53TG3D | 3.208 | 1.310 | 0.01374468 |
| ENSG00000125869 | LAMP5 | 177.067 | 1.307 | 0.000147046 |
| ENSG00000272438 | | 10.737 | 1.304 | 0.000815893 |
| ENSG00000173894 | CBX2 | 1286.325 | 1.304 | 1.49307E-09 |
| ENSG00000196109 | ZNF676 | 40.502 | 1.302 | 0.023454808 |
| ENSG00000131126 | TEX101 | 6.220 | 1.298 | 0.007709472 |
| ENSG00000265933 | LINC00668 | 129.060 | 1.298 | 0.017148707 |
| ENSG00000173404 | INSM1 | 21.409 | 1.296 | 0.014298017 |
| ENSG00000226913 | BSN-DT | 4.371 | 1.294 | 5.23622E-05 |
| ENSG00000177519 | RPRM | 100.796 | 1.291 | 0.000476444 |
| ENSG00000112273 | HDGFL1 | 7.354 | 1.290 | 0.005757963 |
| ENSG00000263551 | | 6.998 | 1.289 | 0.012358086 |
| ENSG00000227857 | | 15.588 | 1.287 | 2.06028E-06 |
| ENSG00000205426 | KRT81 | 399.016 | 1.284 | 0.006621968 |
| ENSG00000168702 | LRP1B | 50.800 | 1.280 | 0.004614475 |
| ENSG00000228956 | SATB1-AS1 | 188.329 | 1.279 | 9.94139E-07 |
| ENSG00000277775 | H3C7 | 7.434 | 1.277 | 4.76426E-06 |
| ENSG00000179915 | NRXN1 | 19.843 | 1.276 | 0.009255761 |
| ENSG00000101812 | H2BW2 | 7.380 | 1.272 | 0.021853213 |
| ENSG00000162891 | IL20 | 19.695 | 1.265 | 0.010047179 |
| ENSG00000176749 | CDK5R1 | 435.128 | 1.265 | 6.2786E-10 |
| ENSG00000251574 | | 3.236 | 1.263 | 0.028209944 |
| ENSG00000183307 | TMEM121B | 44.345 | 1.260 | 1.15688E-08 |
| ENSG00000112852 | PCDHB2 | 284.880 | 1.255 | 2.73918E-05 |
| ENSG00000167355 | OR51B5 | 9.838 | 1.255 | 0.024832432 |
| ENSG00000139988 | RDH12 | 96.166 | 1.253 | 0.000279441 |
| ENSG00000086506 | HBQ1 | 6.233 | 1.253 | 0.007418942 |
| ENSG00000229676 | ZNF492 | 42.789 | 1.253 | 0.002339543 |
| ENSG00000183908 | LRRC55 | 16.440 | 1.252 | 2.97145E-06 |
| ENSG00000270972 | | 3.916 | 1.245 | 0.002511113 |
| ENSG00000258947 | TUBB3 | 332.020 | 1.244 | 1.25112E-06 |
| ENSG00000231240 | KLF2P1 | 25.883 | 1.242 | 0.034059583 |
| ENSG00000196277 | GRM7 | 10.950 | 1.238 | 0.002567876 |
| ENSG00000277224 | H2BC7 | 28.565 | 1.232 | 4.84364E-05 |
| ENSG00000204420 | MPIG6B | 21.423 | 1.232 | 1.52913E-06 |
| ENSG00000152092 | ASTN1 | 17.167 | 1.229 | 0.008815723 |
| ENSG00000231327 | LINC01816 | 29.412 | 1.228 | 2.85282E-10 |
| ENSG00000246350 | | 4.154 | 1.226 | 0.000106614 |
| ENSG00000185640 | KRT79 | 116.209 | 1.223 | 0.011613955 |
| ENSG00000234323 | LINC01505 | 18.602 | 1.221 | 0.008059354 |
| ENSG00000261754 | | 16.552 | 1.221 | 3.79882E-06 |
| ENSG00000268949 | MRPS17P1 | 6.786 | 1.221 | 0.000104745 |
| ENSG00000138829 | FBN2 | 1768.726 | 1.218 | 0.001334022 |
| ENSG00000162975 | KCNF1 | 136.191 | 1.214 | 0.004569821 |

| | | | | |
|---|---|---|---|---|
| ENSG00000120057 | SFRP5 | 11.562 | 1.213 | 0.028599657 |
| ENSG00000040731 | CDH10 | 5.370 | 1.213 | 0.038559863 |
| ENSG00000197705 | KLHL14 | 17.699 | 1.211 | 0.000451281 |
| ENSG00000122733 | PHF24 | 19.894 | 1.211 | 0.000403833 |
| ENSG00000266578 | | 7.544 | 1.210 | 0.000428498 |
| ENSG00000165973 | NELL1 | 21.130 | 1.210 | 0.016480898 |
| ENSG00000230102 | LINC02028 | 13.898 | 1.208 | 0.00028544 |
| ENSG00000113209 | PCDHB5 | 198.702 | 1.207 | 0.000618654 |
| ENSG00000267767 | LINC01801 | 6.639 | 1.206 | 0.000162285 |
| ENSG00000163735 | CXCL5 | 519.406 | 1.206 | 0.014359267 |
| ENSG00000170074 | FAM153A | 14.628 | 1.206 | 0.00163025 |
| ENSG00000132639 | SNAP25 | 69.324 | 1.205 | 0.000412682 |
| ENSG00000070886 | EPHA8 | 9.726 | 1.204 | 0.0135408 |
| ENSG00000187242 | KRT12 | 37.317 | 1.203 | 0.003298687 |
| ENSG00000265158 | LRRC37A7P | 16.984 | 1.202 | 0.000830816 |
| ENSG00000064763 | FAR2 | 936.304 | 1.200 | 2.11323E-07 |
| ENSG00000101349 | PAK5 | 10.505 | 1.199 | 0.030717997 |
| ENSG00000134323 | MYCN | 470.034 | 1.194 | 0.003185315 |
| ENSG00000250979 | | 5.223 | 1.194 | 0.001188396 |
| ENSG00000157927 | RADIL | 66.894 | 1.194 | 7.20369E-06 |
| ENSG00000258525 | | 6.389 | 1.193 | 0.000293774 |
| ENSG00000254024 | | 181.588 | 1.192 | 0.000504626 |
| ENSG00000228261 | ITPRIP-AS1 | 8.066 | 1.189 | 1.60086E-05 |
| ENSG00000261586 | | 39.030 | 1.187 | 9.00266E-05 |
| ENSG00000167749 | KLK4 | 24.781 | 1.184 | 0.015879782 |
| ENSG00000065609 | SNAP91 | 123.567 | 1.184 | 0.009759467 |
| ENSG00000275329 | | 6.633 | 1.182 | 1.01078E-05 |
| ENSG00000260209 | | 13.565 | 1.181 | 0.024497121 |
| ENSG00000233381 | AK4P3 | 4.362 | 1.179 | 0.003411971 |
| ENSG00000105889 | STEAP1B | 90.346 | 1.178 | 0.005098621 |
| ENSG00000104059 | FAM189A1 | 213.759 | 1.173 | 0.007642179 |
| ENSG00000118402 | ELOVL4 | 201.774 | 1.172 | 5.54096E-05 |
| ENSG00000074219 | TEAD2 | 1728.579 | 1.172 | 1.81293E-07 |
| ENSG00000188859 | FAM78B | 374.083 | 1.168 | 0.000851709 |
| ENSG00000237742 | | 5.277 | 1.167 | 0.003465073 |
| ENSG00000225096 | | 5.417 | 1.166 | 0.00254431 |
| ENSG00000237390 | | 5.484 | 1.165 | 0.00143382 |
| ENSG00000100027 | YPEL1 | 155.106 | 1.165 | 5.18719E-08 |
| ENSG00000278952 | | 24.877 | 1.164 | 5.05495E-05 |
| ENSG00000266278 | LINC01910 | 8.187 | 1.164 | 0.022231094 |
| ENSG00000250056 | LINC01018 | 23.469 | 1.162 | 0.003786577 |
| ENSG00000183248 | PRR36 | 723.870 | 1.160 | 0.000332808 |
| ENSG00000072182 | ASIC4 | 6.929 | 1.158 | 2.49514E-05 |
| ENSG00000135477 | KRT87P | 140.340 | 1.156 | 1.67269E-05 |
| ENSG00000205502 | C2CD4B | 99.666 | 1.156 | 9.67748E-05 |
| ENSG00000254349 | MIR2052HG | 17.697 | 1.155 | 0.003570015 |
| ENSG00000175161 | CADM2 | 34.163 | 1.153 | 0.020299228 |
| ENSG00000109265 | CRACD | 202.295 | 1.152 | 2.07192E-05 |
| ENSG00000237940 | LINC01238 | 27.127 | 1.147 | 9.91916E-05 |
| ENSG00000248383 | PCDHAC1 | 12.158 | 1.147 | 0.012099514 |
| ENSG00000145808 | ADAMTS19 | 37.820 | 1.147 | 0.014003055 |
| ENSG00000176204 | LRRTM4 | 10.086 | 1.145 | 0.019535191 |
| ENSG00000114349 | GNAT1 | 5.601 | 1.143 | 0.002823393 |
| ENSG00000133640 | LRRIQ1 | 39.845 | 1.143 | 0.001281383 |
| ENSG00000230067 | HSPD1P6 | 69.999 | 1.141 | 0.002583794 |
| ENSG00000148408 | CACNA1B | 27.385 | 1.140 | 0.019898475 |
| ENSG00000273252 | OR7E39P | 9.011 | 1.135 | 0.010096639 |
| ENSG00000189127 | ANKRD34B | 31.066 | 1.133 | 0.020638593 |
| ENSG00000164796 | CSMD3 | 18.957 | 1.132 | 0.022770995 |

| | | | | |
|---|---|---|---|---|
| ENSG00000169760 | NLGN1 | 55.935 | 1.131 | 0.000852441 |
| ENSG00000254602 | | 15.237 | 1.131 | 5.6917E-06 |
| ENSG00000214146 | LINC02026 | 10.171 | 1.131 | 0.000127447 |
| ENSG00000180251 | SLC9A4 | 439.973 | 1.130 | 0.014567797 |
| ENSG00000107014 | RLN2 | 29.943 | 1.130 | 8.26059E-05 |
| ENSG00000198846 | TOX | 176.456 | 1.130 | 0.000169912 |
| ENSG00000113758 | DBN1 | 4347.165 | 1.129 | 2.77745E-11 |
| ENSG00000257818 | C1GALT1P1 | 16.735 | 1.125 | 0.001596151 |
| ENSG00000198720 | ANKRD13B | 811.618 | 1.122 | 3.97515E-13 |
| ENSG00000188517 | COL25A1 | 7.984 | 1.120 | 0.001253303 |
| ENSG00000163661 | PTX3 | 219.566 | 1.120 | 0.00300784 |
| ENSG00000214796 | TUBA5P | 549.417 | 1.120 | 1.89637E-06 |
| ENSG00000125538 | IL1B | 1336.577 | 1.119 | 0.00096976 |
| ENSG00000013297 | CLDN11 | 404.339 | 1.119 | 0.000291765 |
| ENSG00000182583 | VCX | 43.872 | 1.118 | 0.034859854 |
| ENSG00000196427 | NBPF4 | 16.237 | 1.117 | 0.018530599 |
| ENSG00000105610 | KLF1 | 5.165 | 1.117 | 1.81795E-05 |
| ENSG00000187486 | KCNJ11 | 66.420 | 1.116 | 2.83234E-05 |
| ENSG00000160469 | BRSK1 | 200.848 | 1.116 | 7.33386E-07 |
| ENSG00000144285 | SCN1A | 3.779 | 1.115 | 0.044732764 |
| ENSG00000273321 | | 6.176 | 1.113 | 0.003104055 |
| ENSG00000172367 | PDZD3 | 103.011 | 1.112 | 0.019222144 |
| ENSG00000230583 | GTF2IRD1P1 | 4.840 | 1.112 | 8.87342E-07 |
| ENSG00000225072 | OR7E116P | 3.737 | 1.112 | 0.028597662 |
| ENSG00000033122 | LRRC7 | 31.571 | 1.112 | 4.90457E-11 |
| ENSG00000134343 | ANO3 | 40.209 | 1.110 | 0.003490926 |
| ENSG00000167912 | | 40.047 | 1.109 | 0.000169833 |
| ENSG00000026559 | KCNG1 | 2250.460 | 1.108 | 0.001523372 |
| ENSG00000177182 | CLVS1 | 25.689 | 1.106 | 2.48883E-05 |
| ENSG00000175175 | PPM1E | 36.395 | 1.106 | 7.26663E-06 |
| ENSG00000143434 | SEMA6C | 609.261 | 1.105 | 2.90105E-11 |
| ENSG00000118156 | ZNF541 | 67.528 | 1.104 | 0.001049597 |
| ENSG00000145908 | ZNF300 | 524.745 | 1.104 | 2.73918E-05 |
| ENSG00000185046 | ANKS1B | 28.019 | 1.102 | 0.000727332 |
| ENSG00000163449 | TMEM169 | 39.682 | 1.100 | 2.24544E-05 |
| ENSG00000233718 | MYCNOS | 34.713 | 1.098 | 0.030965698 |
| ENSG00000273443 | | 24.161 | 1.097 | 0.000382421 |
| ENSG00000188176 | SMTNL2 | 111.179 | 1.097 | 0.000760064 |
| ENSG00000155816 | FMN2 | 13.592 | 1.097 | 0.023233255 |
| ENSG00000185130 | H2BC13 | 9.060 | 1.097 | 1.5028E-06 |
| ENSG00000272002 | | 4.603 | 1.096 | 6.55371E-06 |
| ENSG00000138606 | SHF | 307.492 | 1.096 | 8.74603E-09 |
| ENSG00000248925 | | 5.218 | 1.087 | 1.99993E-06 |
| ENSG00000175928 | LRRN1 | 470.044 | 1.085 | 0.00146751 |
| ENSG00000140600 | SH3GL3 | 50.222 | 1.084 | 0.044132357 |
| ENSG00000077264 | PAK3 | 51.169 | 1.082 | 0.005130459 |
| ENSG00000244485 | RPL18P13 | 6.774 | 1.082 | 0.00687453 |
| ENSG00000170423 | KRT78 | 99.638 | 1.081 | 0.005502933 |
| ENSG00000230699 | | 36.099 | 1.080 | 0.010692114 |
| ENSG00000087250 | MT3 | 13.205 | 1.078 | 0.009796055 |
| ENSG00000179774 | ATOH7 | 10.282 | 1.078 | 0.000106906 |
| ENSG00000160145 | KALRN | 763.145 | 1.076 | 1.14198E-05 |
| ENSG00000233926 | | 4.705 | 1.076 | 0.000489376 |
| ENSG00000077420 | APBB1IP | 750.511 | 1.072 | 0.000667024 |
| ENSG00000148942 | SLC5A12 | 110.599 | 1.070 | 0.000595469 |
| ENSG00000129195 | PIMREG | 672.676 | 1.070 | 2.04478E-11 |
| ENSG00000119927 | GPAM | 795.834 | 1.069 | 2.18342E-15 |
| ENSG00000145681 | HAPLN1 | 174.147 | 1.069 | 0.001438315 |
| ENSG00000179455 | MKRN3 | 101.390 | 1.067 | 0.034611744 |

| | | | | |
|---|---|---|---|---|
| ENSG00000134201 | GSTM5 | 251.943 | 1.066 | 0.002059617 |
| ENSG00000055813 | CCDC85A | 15.053 | 1.066 | 0.001259394 |
| ENSG00000184702 | SEPTIN5 | 946.220 | 1.065 | 1.59984E-06 |
| ENSG00000178531 | CTXN1 | 591.006 | 1.065 | 7.66976E-06 |
| ENSG00000162374 | ELAVL4 | 13.506 | 1.065 | 0.001965266 |
| ENSG00000092850 | TEKT2 | 39.878 | 1.064 | 0.000550196 |
| ENSG00000131773 | KHDRBS3 | 364.747 | 1.062 | 8.38015E-05 |
| ENSG00000253686 | LINC01484 | 3.643 | 1.062 | 0.003237632 |
| ENSG00000270035 | | 4.805 | 1.061 | 0.012582669 |
| ENSG00000122778 | KIAA1549 | 645.891 | 1.061 | 1.92288E-09 |
| ENSG00000124657 | OR2B6 | 23.802 | 1.058 | 0.000495819 |
| ENSG00000242078 | | 5.306 | 1.055 | 7.51633E-05 |
| ENSG00000211979 | IGHV7-81 | 13.382 | 1.054 | 0.00647999 |
| ENSG00000223561 | | 6.709 | 1.053 | 0.003483495 |
| ENSG00000184949 | FAM227A | 170.263 | 1.053 | 1.64774E-08 |
| ENSG00000139352 | ASCL1 | 10.224 | 1.052 | 0.040225312 |
| ENSG00000079101 | CLUL1 | 25.279 | 1.048 | 0.00032547 |
| ENSG00000105767 | CADM4 | 1692.264 | 1.046 | 3.16621E-10 |
| ENSG00000164398 | ACSL6 | 20.873 | 1.044 | 0.000201742 |
| ENSG00000182912 | TSPEAR-AS2 | 93.923 | 1.043 | 0.034151998 |
| ENSG00000272056 | | 7.237 | 1.042 | 2.51273E-08 |
| ENSG00000144057 | ST6GAL2 | 144.732 | 1.042 | 0.003176359 |
| ENSG00000164236 | ANKRD33B | 251.501 | 1.042 | 0.000801116 |
| ENSG00000177359 | | 85.609 | 1.041 | 0.000719861 |
| ENSG00000167889 | MGAT5B | 58.367 | 1.040 | 0.002921401 |
| ENSG00000224265 | LINC02633 | 5.810 | 1.038 | 0.029075742 |
| ENSG00000188175 | HEPACAM2 | 15.853 | 1.038 | 0.034691886 |
| ENSG00000067840 | PDZD4 | 130.999 | 1.037 | 0.000188214 |
| ENSG00000165799 | RNASE7 | 232.781 | 1.037 | 0.010093137 |
| ENSG00000176049 | JAKMIP2 | 104.905 | 1.037 | 0.001523372 |
| ENSG00000250709 | CCDC169-SOHLH | 3.045 | 1.037 | 0.016425039 |
| ENSG00000133519 | ZDHHC8P1 | 77.096 | 1.034 | 0.009931972 |
| ENSG00000267432 | DNAH17-AS1 | 15.540 | 1.033 | 0.007907286 |
| ENSG00000034053 | APBA2 | 535.897 | 1.033 | 0.000163397 |
| ENSG00000176387 | HSD11B2 | 630.368 | 1.030 | 2.03513E-05 |
| ENSG00000273802 | H2BC8 | 163.792 | 1.028 | 0.000544127 |
| ENSG00000260132 | | 4.733 | 1.027 | 0.000716547 |
| ENSG00000128045 | RASL11B | 387.067 | 1.027 | 0.000166568 |
| ENSG00000258405 | ZNF578 | 26.044 | 1.026 | 0.00067629 |
| ENSG00000146038 | DCDC2 | 138.075 | 1.023 | 0.021207591 |
| ENSG00000114757 | PEX5L | 12.944 | 1.023 | 0.004083282 |
| ENSG00000104332 | SFRP1 | 721.432 | 1.022 | 0.014705216 |
| ENSG00000240040 | | 9.153 | 1.021 | 0.018001921 |
| ENSG00000170681 | CAVIN4 | 41.266 | 1.021 | 1.48887E-09 |
| ENSG00000153404 | PLEKHG4B | 516.615 | 1.021 | 0.017714386 |
| ENSG00000161031 | PGLYRP2 | 26.885 | 1.019 | 0.012438412 |
| ENSG00000141744 | PNMT | 305.612 | 1.018 | 0.010420465 |
| ENSG00000159409 | CELF3 | 5.050 | 1.018 | 0.002925245 |
| ENSG00000266602 | | 48.599 | 1.017 | 0.049737507 |
| ENSG00000232938 | RPL23AP87 | 17.552 | 1.014 | 0.001500314 |
| ENSG00000108309 | RUNDC3A | 154.596 | 1.013 | 0.000798679 |
| ENSG00000274276 | CBSL | 23.254 | 1.011 | 0.025260562 |
| ENSG00000242797 | GLYCTK-AS1 | 7.066 | 1.010 | 0.00158229 |
| ENSG00000127585 | FBXL16 | 158.576 | 1.010 | 0.000996092 |
| ENSG00000124391 | IL17C | 21.195 | 1.009 | 0.034537566 |
| ENSG00000274376 | ADAMTS7P1 | 10.228 | 1.006 | 2.35886E-05 |
| ENSG00000152503 | TRIM36 | 71.647 | 1.006 | 0.000390558 |
| ENSG00000272180 | | 5.302 | 1.005 | 0.001830652 |
| ENSG00000186231 | KLHL32 | 16.056 | 1.005 | 0.000690805 |

| | | | | |
|---|---|---|---|---|
| ENSG00000259755 | | 7.061 | 1.003 | 0.008518741 |
| ENSG00000234511 | C5orf58 | 26.378 | 1.002 | 0.010033199 |
| ENSG00000223823 | LINC01342 | 8.276 | 1.000 | 0.021195447 |
| ENSG00000160781 | PAQR6 | 324.239 | 0.999 | 1.37547E-06 |
| ENSG00000263809 | | 6.844 | 0.999 | 8.86146E-05 |
| ENSG00000117707 | PROX1 | 141.154 | 0.998 | 0.00397074 |
| ENSG00000136378 | ADAMTS7 | 1086.850 | 0.997 | 4.00536E-08 |
| ENSG00000279791 | | 4.605 | 0.995 | 0.001644789 |
| ENSG00000188523 | CFAP77 | 20.444 | 0.995 | 0.024856283 |
| ENSG00000235436 | DPY19L2P4 | 5.456 | 0.994 | 0.022791146 |
| ENSG00000163879 | DNALI1 | 324.633 | 0.994 | 0.001523372 |
| ENSG00000255202 | | 22.916 | 0.993 | 0.005821514 |
| ENSG00000273032 | DGCR5 | 22.556 | 0.992 | 0.00425828 |
| ENSG00000158427 | TMSB15B | 25.560 | 0.991 | 4.72394E-08 |
| ENSG00000183833 | CFAP91 | 108.962 | 0.990 | 0.003794784 |
| ENSG00000259985 | | 42.040 | 0.988 | 2.97145E-06 |
| ENSG00000092295 | TGM1 | 1141.059 | 0.986 | 0.024424542 |
| ENSG00000116183 | PAPPA2 | 56.298 | 0.985 | 0.016664218 |
| ENSG00000236708 | SDK1-AS1 | 5.804 | 0.983 | 0.008794328 |
| ENSG00000137819 | PAQR5 | 410.239 | 0.983 | 0.000624096 |
| ENSG00000173825 | TIGD3 | 68.938 | 0.982 | 1.01794E-06 |
| ENSG00000162188 | GNG3 | 15.868 | 0.979 | 5.24725E-08 |
| ENSG00000237517 | DGCR5 | 57.234 | 0.977 | 0.007534797 |
| ENSG00000138622 | HCN4 | 12.978 | 0.976 | 0.048577914 |
| ENSG00000278635 | | 15.933 | 0.976 | 5.29386E-05 |
| ENSG00000126010 | GRPR | 5.107 | 0.974 | 0.001908202 |
| ENSG00000105880 | DLX5 | 420.658 | 0.974 | 0.001877796 |
| ENSG00000251994 | RNU2-27P | 27.481 | 0.972 | 5.97344E-07 |
| ENSG00000075461 | CACNG4 | 667.615 | 0.971 | 0.031643876 |
| ENSG00000162373 | BEND5 | 257.139 | 0.971 | 8.90681E-05 |
| ENSG00000172379 | ARNT2 | 560.610 | 0.971 | 0.000102349 |
| ENSG00000109758 | HGFAC | 17.452 | 0.970 | 0.001060696 |
| ENSG00000217275 | | 60.592 | 0.970 | 0.001310274 |
| ENSG00000183850 | ZNF730 | 37.580 | 0.970 | 0.009401007 |
| ENSG00000215146 | | 134.651 | 0.968 | 0.000830816 |
| ENSG00000162174 | ASRGL1 | 277.683 | 0.965 | 0.000239196 |
| ENSG00000236924 | | 13.737 | 0.963 | 0.038258344 |
| ENSG00000269994 | LINC02893 | 195.320 | 0.963 | 0.015511213 |
| ENSG00000163734 | CXCL3 | 159.345 | 0.960 | 0.008281352 |
| ENSG00000168779 | SHOX2 | 114.081 | 0.959 | 0.004726708 |
| ENSG00000184368 | MAP7D2 | 108.169 | 0.956 | 0.01402105 |
| ENSG00000152192 | POU4F1 | 37.519 | 0.955 | 0.041534504 |
| ENSG00000146966 | DENND2A | 386.727 | 0.954 | 2.26095E-06 |
| ENSG00000073670 | ADAM11 | 112.053 | 0.953 | 0.000121714 |
| ENSG00000112276 | BVES | 256.057 | 0.953 | 0.000727332 |
| ENSG00000144583 | MARCHF4 | 74.420 | 0.952 | 0.036349236 |
| ENSG00000279693 | | 4.331 | 0.951 | 0.017971158 |
| ENSG00000233922 | LINC01694 | 72.006 | 0.949 | 0.007534797 |
| ENSG00000108576 | SLC6A4 | 81.530 | 0.949 | 0.010206306 |
| ENSG00000165810 | BTNL9 | 301.461 | 0.948 | 0.003075266 |
| ENSG00000154917 | RAB6B | 374.918 | 0.948 | 2.05837E-05 |
| ENSG00000149968 | MMP3 | 586.720 | 0.946 | 0.022127552 |
| ENSG00000198723 | TEX45 | 133.390 | 0.946 | 0.007113768 |
| ENSG00000266805 | | 14.209 | 0.946 | 0.002761034 |
| ENSG00000047617 | ANO2 | 39.867 | 0.944 | 6.63342E-06 |
| ENSG00000184194 | GPR173 | 220.933 | 0.943 | 8.26384E-05 |
| ENSG00000169474 | SPRR1A | 1627.277 | 0.943 | 0.032353838 |
| ENSG00000255129 | | 5.430 | 0.942 | 0.00254431 |
| ENSG00000203808 | BVES-AS1 | 13.492 | 0.939 | 0.017748446 |

| | | | | |
|---|---|---|---|---|
| ENSG00000254872 | LINC02688 | 8.772 | 0.937 | 0.019928782 |
| ENSG00000141622 | RNF165 | 144.024 | 0.936 | 0.002364528 |
| ENSG00000254777 | | 5.275 | 0.936 | 0.003465073 |
| ENSG00000225868 | WDR87BP | 24.836 | 0.935 | 0.005556638 |
| ENSG00000112394 | SLC16A10 | 212.020 | 0.934 | 0.000703753 |
| ENSG00000255753 | | 4.166 | 0.934 | 0.000902417 |
| ENSG00000174226 | SNX31 | 3456.226 | 0.933 | 0.049599902 |
| ENSG00000258815 | LINC02820 | 41.007 | 0.933 | 0.009425299 |
| ENSG00000162669 | HFM1 | 13.310 | 0.932 | 0.003781847 |
| ENSG00000152578 | GRIA4 | 3.958 | 0.932 | 0.004465829 |
| ENSG00000105894 | PTN | 3598.584 | 0.931 | 0.027670415 |
| ENSG00000243230 | | 5.676 | 0.930 | 0.009067265 |
| ENSG00000146005 | PSD2 | 59.689 | 0.929 | 0.024742098 |
| ENSG00000105929 | ATP6V0A4 | 184.745 | 0.927 | 0.044878046 |
| ENSG00000131771 | PPP1R1B | 1169.391 | 0.927 | 0.049742785 |
| ENSG00000214870 | | 33.096 | 0.926 | 7.56294E-05 |
| ENSG00000238178 | | 7.531 | 0.926 | 0.017152757 |
| ENSG00000267886 | | 20.024 | 0.925 | 0.033567426 |
| ENSG00000233384 | CNIH3-AS2 | 18.014 | 0.924 | 0.03816747 |
| ENSG00000184005 | ST6GALNAC3 | 74.233 | 0.924 | 4.37632E-06 |
| ENSG00000170959 | DCDC1 | 18.643 | 0.924 | 0.00248395 |
| ENSG00000269226 | TMSB15B | 5.228 | 0.923 | 3.52704E-05 |
| ENSG00000255843 | | 12.034 | 0.922 | 0.002487983 |
| ENSG00000249661 | TNRC18P1 | 13.494 | 0.921 | 0.00294612 |
| ENSG00000272398 | CD24 | 32218.355 | 0.919 | 0.000314552 |
| ENSG00000196747 | H2AC13 | 32.625 | 0.919 | 0.00128602 |
| ENSG00000234261 | | 10.842 | 0.918 | 0.02467023 |
| ENSG00000109084 | TMEM97 | 7814.255 | 0.917 | 0.000659171 |
| ENSG00000278291 | | 173.476 | 0.917 | 0.000353863 |
| ENSG00000272138 | LINC01607 | 19.981 | 0.916 | 0.000486228 |
| ENSG00000197153 | H3C12 | 6.395 | 0.915 | 0.001911113 |
| ENSG00000267640 | | 28.593 | 0.914 | 0.01492346 |
| ENSG00000273687 | | 7.351 | 0.914 | 0.000200172 |
| ENSG00000136866 | ZFP37 | 108.777 | 0.912 | 0.000109544 |
| ENSG00000254585 | MAGEL2 | 32.352 | 0.912 | 0.0072632 |
| ENSG00000099256 | PRTFDC1 | 373.954 | 0.910 | 4.78272E-05 |
| ENSG00000165097 | KDM1B | 1185.109 | 0.909 | 3.65358E-17 |
| ENSG00000269978 | | 8.498 | 0.909 | 0.024193495 |
| ENSG00000231892 | | 6.650 | 0.905 | 0.015092306 |
| ENSG00000077935 | SMC1B | 145.845 | 0.904 | 0.004306538 |
| ENSG00000134940 | ACRV1 | 23.442 | 0.902 | 1.21099E-07 |
| ENSG00000147255 | IGSF1 | 91.532 | 0.902 | 0.018705751 |
| ENSG00000261087 | ZNNT1 | 165.736 | 0.902 | 2.1439E-06 |
| ENSG00000107105 | ELAVL2 | 114.451 | 0.901 | 0.042694754 |
| ENSG00000115616 | SLC9A2 | 831.405 | 0.900 | 0.014479214 |
| ENSG00000267156 | TPMTP1 | 3.898 | 0.899 | 8.83966E-05 |
| ENSG00000106078 | COBL | 215.029 | 0.898 | 0.035327125 |
| ENSG00000182263 | FIGN | 297.129 | 0.898 | 0.00071448 |
| ENSG00000265480 | KRT18P55 | 13.349 | 0.898 | 0.003147365 |
| ENSG00000265975 | | 8.085 | 0.898 | 0.002165518 |
| ENSG00000167552 | TUBA1A | 6295.419 | 0.897 | 5.99267E-05 |
| ENSG00000240567 | LINC02067 | 5.895 | 0.897 | 0.009407285 |
| ENSG00000089169 | RPH3A | 4.164 | 0.895 | 0.009551665 |
| ENSG00000215246 | | 14.792 | 0.894 | 0.002750226 |
| ENSG00000115290 | GRB14 | 84.094 | 0.894 | 0.001579919 |
| ENSG00000149926 | TLCD3B | 16.907 | 0.893 | 0.002161406 |
| ENSG00000174871 | CNIH2 | 153.882 | 0.893 | 0.000317181 |
| ENSG00000217801 | | 438.483 | 0.893 | 0.000265031 |
| ENSG00000183831 | ANKRD45 | 23.329 | 0.892 | 0.005715467 |

| | | | | |
|---|---|---|---|---|
| ENSG00000225173 | | 7.451 | 0.892 | 0.008197884 |
| ENSG00000204434 | POTEKP | 16.271 | 0.891 | 0.026755191 |
| ENSG00000159753 | CARMIL2 | 335.297 | 0.891 | 0.001319023 |
| ENSG00000274641 | H2BC17 | 22.761 | 0.891 | 0.002721135 |
| ENSG00000225742 | LINC02036 | 18.001 | 0.890 | 0.002627399 |
| ENSG00000180458 | | 8.041 | 0.890 | 0.023413057 |
| ENSG00000092054 | MYH7 | 8.500 | 0.889 | 0.046865616 |
| ENSG00000153132 | CLGN | 113.347 | 0.888 | 0.007112721 |
| ENSG00000166582 | CENPV | 720.659 | 0.887 | 7.33188E-05 |
| ENSG00000167525 | PROCA1 | 222.953 | 0.887 | 1.28173E-06 |
| ENSG00000130413 | STK33 | 88.260 | 0.886 | 0.007275124 |
| ENSG00000187068 | C3orf70 | 283.361 | 0.885 | 0.000260707 |
| ENSG00000129437 | KLK14 | 92.005 | 0.883 | 0.041420703 |
| ENSG00000186019 | | 15.315 | 0.883 | 0.000684391 |
| ENSG00000261308 | FIGNL2 | 11.345 | 0.882 | 0.029019774 |
| ENSG00000179750 | APOBEC3B | 972.131 | 0.881 | 0.000364748 |
| ENSG00000157087 | ATP2B2 | 9.400 | 0.878 | 0.007275124 |
| ENSG00000076864 | RAP1GAP | 1673.991 | 0.878 | 0.000957572 |
| ENSG00000164061 | BSN | 81.395 | 0.877 | 0.000306905 |
| ENSG00000184774 | MGAT4EP | 7.920 | 0.876 | 0.002550495 |
| ENSG00000142611 | PRDM16 | 88.715 | 0.876 | 0.014794776 |
| ENSG00000163958 | ZDHHC19 | 19.735 | 0.875 | 0.005447367 |
| ENSG00000115423 | DNAH6 | 46.625 | 0.874 | 0.002597817 |
| ENSG00000148935 | GAS2 | 87.750 | 0.873 | 0.024770055 |
| ENSG00000176244 | ACBD7 | 44.177 | 0.872 | 0.00118519 |
| ENSG00000197889 | MEIG1 | 15.317 | 0.871 | 3.1081E-05 |
| ENSG00000151322 | NPAS3 | 44.938 | 0.871 | 0.007997346 |
| ENSG00000149488 | TMC2 | 5.353 | 0.870 | 0.001826135 |
| ENSG00000105048 | TNNT1 | 602.187 | 0.870 | 0.029495381 |
| ENSG00000127252 | PLAAT1 | 37.231 | 0.868 | 0.032992198 |
| ENSG00000226091 | LINC00937 | 43.775 | 0.868 | 0.001218417 |
| ENSG00000198216 | CACNA1E | 12.469 | 0.868 | 0.014286791 |
| ENSG00000137142 | IGFBPL1 | 18.423 | 0.867 | 0.031280042 |
| ENSG00000196876 | SCN8A | 157.668 | 0.867 | 0.004614475 |
| ENSG00000214999 | | 16.541 | 0.866 | 0.012654852 |
| ENSG00000197083 | ZNF300P1 | 33.269 | 0.865 | 0.005484771 |
| ENSG00000144847 | IGSF11 | 141.902 | 0.864 | 0.005655432 |
| ENSG00000243069 | ARHGEF26-AS1 | 130.354 | 0.864 | 0.019368776 |
| ENSG00000128266 | GNAZ | 160.819 | 0.862 | 0.004338069 |
| ENSG00000137261 | KIAA0319 | 62.620 | 0.862 | 0.017002862 |
| ENSG00000232000 | CLCN3P1 | 12.425 | 0.862 | 0.022990056 |
| ENSG00000250451 | HOXC-AS1 | 25.936 | 0.862 | 0.047709691 |
| ENSG00000139219 | COL2A1 | 27.121 | 0.860 | 0.029032111 |
| ENSG00000149403 | GRIK4 | 45.912 | 0.860 | 0.009012894 |
| ENSG00000108947 | EFNB3 | 261.615 | 0.859 | 0.004412094 |
| ENSG00000275713 | H2BC9 | 123.538 | 0.859 | 0.004933013 |
| ENSG00000115008 | IL1A | 1706.662 | 0.859 | 0.028664234 |
| ENSG00000242600 | MBL1P | 24.826 | 0.858 | 0.000286307 |
| ENSG00000255240 | | 5.122 | 0.858 | 0.043288959 |
| ENSG00000276445 | | 19.222 | 0.857 | 7.21857E-05 |
| ENSG00000112182 | BACH2 | 139.563 | 0.853 | 0.007671762 |
| ENSG00000121236 | TRIM6 | 399.474 | 0.853 | 0.000199601 |
| ENSG00000172733 | PURG | 6.920 | 0.852 | 0.010547683 |
| ENSG00000104177 | MYEF2 | 312.556 | 0.852 | 0.008112524 |
| ENSG00000185652 | NTF3 | 35.276 | 0.851 | 0.004684555 |
| ENSG00000261159 | | 40.552 | 0.849 | 0.000902417 |
| ENSG00000005961 | ITGA2B | 66.357 | 0.848 | 0.001711231 |
| ENSG00000204556 | | 15.388 | 0.847 | 0.000427891 |
| ENSG00000160200 | CBS | 63.336 | 0.847 | 0.017936776 |

| | | | | |
|---|---|---|---|---|
| ENSG00000222014 | RAB6C | 6.683 | 0.846 | 0.005843511 |
| ENSG00000134986 | NREP | 1746.246 | 0.845 | 1.76041E-05 |
| ENSG00000100473 | COCH | 314.153 | 0.845 | 0.008659813 |
| ENSG00000251026 | LINC02163 | 22.345 | 0.843 | 0.0279269 |
| ENSG00000196812 | ZSCAN16 | 372.233 | 0.842 | 5.10959E-09 |
| ENSG00000269416 | LINC01224 | 147.039 | 0.842 | 0.037586386 |
| ENSG00000152672 | CLEC4F | 16.388 | 0.842 | 0.010522886 |
| ENSG00000149571 | KIRREL3 | 26.431 | 0.840 | 0.000378686 |
| ENSG00000247151 | CSTF3-DT | 4.215 | 0.839 | 0.000865725 |
| ENSG00000133169 | BEX1 | 12.440 | 0.838 | 0.032245677 |
| ENSG00000175697 | GPR156 | 60.232 | 0.838 | 0.007187669 |
| ENSG00000112290 | WASF1 | 642.121 | 0.837 | 5.06351E-08 |
| ENSG00000105219 | CCNP | 58.606 | 0.837 | 0.00744183 |
| ENSG00000167646 | DNAAF3 | 56.805 | 0.836 | 0.002884727 |
| ENSG00000227051 | C14orf132 | 406.515 | 0.836 | 0.002274941 |
| ENSG00000167771 | RCOR2 | 428.600 | 0.834 | 0.001850381 |
| ENSG00000146938 | NLGN4X | 297.741 | 0.833 | 0.025800793 |
| ENSG00000122824 | NUDT10 | 24.657 | 0.832 | 0.036082859 |
| ENSG00000006377 | DLX6 | 163.288 | 0.831 | 0.015099696 |
| ENSG00000164082 | GRM2 | 40.181 | 0.831 | 8.15387E-05 |
| ENSG00000237732 | CT75 | 27.248 | 0.831 | 0.003786928 |
| ENSG00000017797 | RALBP1 | 11055.328 | 0.829 | 5.75285E-06 |
| ENSG00000272347 | | 19.782 | 0.828 | 0.017786834 |
| ENSG00000242715 | CCDC169 | 162.391 | 0.828 | 0.002005333 |
| ENSG00000225857 | LINC02816 | 3.445 | 0.828 | 0.013449747 |
| ENSG00000101542 | CDH20 | 11.025 | 0.828 | 0.007363733 |
| ENSG00000128602 | SMO | 1004.434 | 0.828 | 0.000639159 |
| ENSG00000206344 | HCG27 | 52.934 | 0.827 | 0.000146642 |
| ENSG00000153157 | SYCP2L | 13.592 | 0.826 | 0.012881254 |
| ENSG00000234409 | CCDC188 | 34.915 | 0.826 | 0.003096471 |
| ENSG00000215022 | | 31.730 | 0.826 | 2.18273E-06 |
| ENSG00000258290 | BRWD1P2 | 3.477 | 0.826 | 0.048332072 |
| ENSG00000138587 | MNS1 | 230.472 | 0.825 | 9.30155E-06 |
| ENSG00000135144 | DTX1 | 165.564 | 0.824 | 0.002049444 |
| ENSG00000230091 | TMEM254-AS1 | 192.961 | 0.823 | 0.001235284 |
| ENSG00000176428 | VPS37D | 161.331 | 0.823 | 0.000214949 |
| ENSG00000273983 | H3C8 | 107.766 | 0.822 | 0.008658662 |
| ENSG00000251013 | GAPDHP62 | 8.784 | 0.820 | 0.002946131 |
| ENSG00000196381 | ZNF781 | 78.709 | 0.818 | 0.001492698 |
| ENSG00000197472 | ZNF695 | 97.243 | 0.818 | 8.76445E-07 |
| ENSG00000114923 | SLC4A3 | 600.319 | 0.818 | 0.000188327 |
| ENSG00000106236 | NPTX2 | 242.380 | 0.817 | 0.001192227 |
| ENSG00000274525 | | 3.670 | 0.817 | 0.032301925 |
| ENSG00000003137 | CYP26B1 | 334.049 | 0.816 | 0.015466871 |
| ENSG00000273216 | | 41.203 | 0.816 | 0.007534797 |
| ENSG00000230023 | LINC02800 | 7.973 | 0.815 | 0.038884198 |
| ENSG00000196476 | C20orf96 | 569.146 | 0.815 | 4.81197E-09 |
| ENSG00000144730 | IL17RD | 408.283 | 0.815 | 5.17296E-05 |
| ENSG00000233251 | | 18.924 | 0.814 | 0.005637163 |
| ENSG00000266389 | PIK3R5-DT | 4.115 | 0.814 | 0.019076928 |
| ENSG00000235033 | DAAM2-AS1 | 55.549 | 0.813 | 0.004602264 |
| ENSG00000124795 | DEK | 8171.842 | 0.812 | 1.42218E-10 |
| ENSG00000279135 | | 4.693 | 0.812 | 0.006650412 |
| ENSG00000169258 | GPRIN1 | 496.878 | 0.811 | 0.000185629 |
| ENSG00000278200 | LINC01971 | 3.625 | 0.810 | 0.041762168 |
| ENSG00000279878 | | 8.584 | 0.808 | 0.046136109 |
| ENSG00000154188 | ANGPT1 | 219.394 | 0.808 | 0.008362818 |
| ENSG00000236824 | BCYRN1 | 60.204 | 0.806 | 0.001310422 |
| ENSG00000223511 | | 11.695 | 0.806 | 0.027266908 |

| | | | | |
|---|---|---|---|---|
| ENSG00000255330 | | 7.872 | 0.806 | 0.002412375 |
| ENSG00000258957 | | 7.773 | 0.804 | 0.000165047 |
| ENSG00000135747 | ZNF670-ZNF695 | 17.922 | 0.803 | 8.95754E-07 |
| ENSG00000182963 | GJC1 | 821.821 | 0.802 | 4.63725E-05 |
| ENSG00000101004 | NINL | 678.731 | 0.798 | 0.000551206 |
| ENSG00000137434 | C6orf52 | 55.575 | 0.798 | 0.000476865 |
| ENSG00000236780 | LINC01829 | 4.084 | 0.797 | 0.0119868 |
| ENSG00000112293 | GPLD1 | 141.779 | 0.797 | 0.006562245 |
| ENSG00000231057 | | 5.391 | 0.797 | 0.00248395 |
| ENSG00000124532 | MRS2 | 1871.304 | 0.796 | 1.35562E-19 |
| ENSG00000117632 | STMN1 | 15195.819 | 0.796 | 8.69714E-09 |
| ENSG00000263164 | | 4.959 | 0.795 | 0.000869578 |
| ENSG00000279970 | | 44.480 | 0.795 | 0.004593125 |
| ENSG00000115457 | IGFBP2 | 4035.618 | 0.794 | 0.012853798 |
| ENSG00000278090 | LUNAR1 | 9.309 | 0.793 | 0.008115606 |
| ENSG00000166546 | BEAN1 | 151.688 | 0.793 | 0.010858048 |
| ENSG00000278231 | | 6.634 | 0.793 | 0.007145216 |
| ENSG00000248027 | ARHGAP42-AS1 | 13.480 | 0.793 | 0.001461594 |
| ENSG00000256037 | MRPL40P1 | 11.047 | 0.792 | 0.016144508 |
| ENSG00000169031 | COL4A3 | 82.757 | 0.792 | 0.034495023 |
| ENSG00000180815 | MAP3K15 | 8.433 | 0.789 | 0.003980974 |
| ENSG00000278828 | H3C10 | 347.612 | 0.788 | 0.006213269 |
| ENSG00000213190 | MLLT11 | 1270.918 | 0.788 | 0.000101907 |
| ENSG00000167077 | 37012 | 162.144 | 0.786 | 0.007364683 |
| ENSG00000246985 | SOCS2-AS1 | 120.719 | 0.784 | 0.004485456 |
| ENSG00000166450 | PRTG | 107.990 | 0.784 | 0.014708491 |
| ENSG00000178187 | ZNF454 | 19.676 | 0.784 | 0.00859398 |
| ENSG00000074276 | CDHR2 | 21.975 | 0.783 | 0.001069289 |
| ENSG00000104899 | AMH | 188.398 | 0.783 | 0.024622559 |
| ENSG00000143147 | GPR161 | 823.195 | 0.782 | 1.11524E-05 |
| ENSG00000167914 | GSDMA | 129.126 | 0.781 | 0.009767915 |
| ENSG00000180806 | HOXC9 | 169.222 | 0.781 | 0.043431653 |
| ENSG00000260920 | | 182.444 | 0.780 | 7.04598E-08 |
| ENSG00000125409 | TEKT3 | 14.103 | 0.779 | 0.004657447 |
| ENSG00000127328 | RAB3IP | 1929.091 | 0.779 | 3.59337E-05 |
| ENSG00000256673 | | 40.848 | 0.778 | 0.000203925 |
| ENSG00000116990 | MYCL | 6661.827 | 0.778 | 0.007654967 |
| ENSG00000255867 | DENND5B-AS1 | 8.186 | 0.777 | 0.040335522 |
| ENSG00000149927 | DOC2A | 81.204 | 0.776 | 0.037197331 |
| ENSG00000114631 | PODXL2 | 2581.018 | 0.776 | 0.001584116 |
| ENSG00000262904 | TMPOP2 | 8.858 | 0.775 | 5.79061E-05 |
| ENSG00000224057 | EGFR-AS1 | 10.446 | 0.774 | 0.023638415 |
| ENSG00000166035 | LIPC | 270.311 | 0.773 | 0.034633147 |
| ENSG00000273274 | ZBTB8B | 139.357 | 0.773 | 0.024768721 |
| ENSG00000161860 | SYCE2 | 49.406 | 0.773 | 1.58112E-05 |
| ENSG00000219891 | ZSCAN12P1 | 72.991 | 0.773 | 7.14276E-06 |
| ENSG00000138061 | CYP1B1 | 1017.238 | 0.772 | 0.034590007 |
| ENSG00000140623 | SEPTIN12 | 4.196 | 0.770 | 0.027843487 |
| ENSG00000115155 | OTOF | 169.528 | 0.770 | 0.043687978 |
| ENSG00000164683 | HEY1 | 705.836 | 0.769 | 0.002260535 |
| ENSG00000213462 | ERV3-1 | 1322.268 | 0.768 | 0.00375655 |
| ENSG00000216331 | H1-12P | 28.051 | 0.768 | 0.010212424 |
| ENSG00000115526 | CHST10 | 386.382 | 0.768 | 0.000712541 |
| ENSG00000258592 | | 10.975 | 0.766 | 0.014230056 |
| ENSG00000213722 | DDAH2 | 2835.171 | 0.766 | 9.52264E-07 |
| ENSG00000265683 | SYPL1P2 | 12.024 | 0.766 | 6.33155E-05 |
| ENSG00000243795 | LINC02044 | 4.032 | 0.765 | 0.001039452 |
| ENSG00000242220 | TCP10L | 14.720 | 0.764 | 0.003445395 |
| ENSG00000021300 | PLEKHB1 | 417.386 | 0.764 | 0.005655432 |

| | | | | |
|---|---|---|---|---|
| ENSG00000154127 | UBASH3B | 1224.369 | 0.764 | 0.000196039 |
| ENSG00000197061 | H4C3 | 6.137 | 0.763 | 0.00123897 |
| ENSG00000108830 | RND2 | 143.622 | 0.762 | 0.004730388 |
| ENSG00000100336 | APOL4 | 2944.087 | 0.761 | 0.003251456 |
| ENSG00000231305 | | 36.762 | 0.761 | 0.000190618 |
| ENSG00000187122 | SLIT1 | 40.792 | 0.760 | 0.006877035 |
| ENSG00000008083 | JARID2 | 2138.652 | 0.760 | 1.8047E-08 |
| ENSG00000154548 | SRSF12 | 49.482 | 0.759 | 0.013148678 |
| ENSG00000265749 | | 56.300 | 0.758 | 0.001087715 |
| ENSG00000165716 | DIPK1B | 235.978 | 0.756 | 0.000732666 |
| ENSG00000162641 | AKNAD1 | 13.929 | 0.755 | 0.028600424 |
| ENSG00000168913 | ENHO | 57.376 | 0.753 | 0.035327125 |
| ENSG00000102098 | SCML2 | 97.671 | 0.753 | 0.020057041 |
| ENSG00000154134 | ROBO3 | 316.673 | 0.753 | 0.001422928 |
| ENSG00000173838 | MARCHF10 | 32.894 | 0.753 | 0.004180806 |
| ENSG00000135502 | SLC26A10 | 19.417 | 0.752 | 0.003794784 |
| ENSG00000275936 | | 6.249 | 0.752 | 0.008815723 |
| ENSG00000110427 | KIAA1549L | 627.138 | 0.750 | 0.010605589 |
| ENSG00000007372 | PAX6 | 219.103 | 0.750 | 0.001759953 |
| ENSG00000166123 | GPT2 | 2475.777 | 0.750 | 2.26095E-06 |
| ENSG00000172687 | ZNF738 | 599.887 | 0.749 | 8.03358E-06 |
| ENSG00000143786 | CNIH3 | 271.504 | 0.749 | 0.006689532 |
| ENSG00000278932 | | 10.873 | 0.748 | 0.000904643 |
| ENSG00000158055 | GRHL3 | 5184.203 | 0.748 | 0.02416512 |
| ENSG00000275491 | LINC01730 | 9.303 | 0.747 | 0.01284303 |
| ENSG00000229757 | | 8.022 | 0.746 | 0.032114822 |
| ENSG00000169783 | LINGO1 | 253.895 | 0.746 | 0.001382152 |
| ENSG00000183914 | DNAH2 | 160.319 | 0.746 | 0.033382722 |
| ENSG00000233822 | H2BC15 | 92.882 | 0.745 | 0.00072571 |
| ENSG00000264569 | DCXR-DT | 11.293 | 0.745 | 0.014724678 |
| ENSG00000155761 | SPAG17 | 364.248 | 0.745 | 0.027652242 |
| ENSG00000198221 | AFDN-DT | 28.229 | 0.744 | 0.000222325 |
| ENSG00000228415 | PTMAP1 | 6.201 | 0.743 | 0.001224344 |
| ENSG00000273305 | | 22.542 | 0.742 | 0.002213298 |
| ENSG00000196581 | AJAP1 | 50.522 | 0.742 | 0.019178383 |
| ENSG00000179241 | LDLRAD3 | 541.663 | 0.742 | 0.000298531 |
| ENSG00000092621 | PHGDH | 6362.314 | 0.740 | 0.000328711 |
| ENSG00000258734 | | 4.683 | 0.739 | 0.00269561 |
| ENSG00000122574 | WIPF3 | 189.120 | 0.738 | 0.007738209 |
| ENSG00000279667 | | 7.069 | 0.738 | 0.004710052 |
| ENSG00000178665 | ZNF713 | 136.960 | 0.738 | 2.7302E-08 |
| ENSG00000106701 | FSD1L | 148.678 | 0.737 | 0.001224499 |
| ENSG00000253953 | PCDHGB4 | 43.223 | 0.736 | 0.028207869 |
| ENSG00000108515 | ENO3 | 208.434 | 0.736 | 2.00634E-05 |
| ENSG00000213876 | RPL7AP64 | 7.580 | 0.736 | 0.011719226 |
| ENSG00000250091 | DNAH10OS | 52.565 | 0.734 | 0.002351702 |
| ENSG00000144681 | STAC | 93.621 | 0.733 | 0.032717637 |
| ENSG00000175130 | MARCKSL1 | 12863.628 | 0.733 | 1.45173E-06 |
| ENSG00000262772 | LINC01977 | 41.926 | 0.732 | 0.010452084 |
| ENSG00000111341 | MGP | 8475.569 | 0.732 | 0.031685277 |
| ENSG00000214121 | PRDX1P1 | 8.185 | 0.731 | 0.015617803 |
| ENSG00000067445 | TRO | 328.984 | 0.730 | 0.006364661 |
| ENSG00000089101 | CFAP61 | 13.988 | 0.730 | 0.00387662 |
| ENSG00000132432 | SEC61G | 3837.659 | 0.729 | 1.72786E-08 |
| ENSG00000141576 | RNF157 | 444.286 | 0.729 | 0.017680396 |
| ENSG00000228203 | GRASLND | 80.602 | 0.728 | 0.007907286 |
| ENSG00000090924 | PLEKHG2 | 1576.656 | 0.727 | 8.2392E-08 |
| ENSG00000274286 | ADRA2B | 66.261 | 0.727 | 0.019222657 |
| ENSG00000120645 | IQSEC3 | 137.848 | 0.727 | 0.019843894 |

| | | | | |
|---|---|---|---|---|
| ENSG00000141371 | C17orf64 | 4.398 | 0.727 | 0.041726916 |
| ENSG00000179388 | EGR3 | 437.567 | 0.727 | 0.014035422 |
| ENSG00000273107 | | 11.660 | 0.727 | 0.009165546 |
| ENSG00000111664 | GNB3 | 69.730 | 0.726 | 0.00118519 |
| ENSG00000111644 | ACRBP | 139.803 | 0.726 | 0.000243704 |
| ENSG00000269425 | | 4.122 | 0.725 | 0.000848976 |
| ENSG00000179899 | PHC1P1 | 24.252 | 0.725 | 0.000131777 |
| ENSG00000163006 | CCDC138 | 333.920 | 0.724 | 2.76661E-11 |
| ENSG00000141431 | ASXL3 | 41.223 | 0.722 | 0.03697738 |
| ENSG00000272049 | | 5.157 | 0.721 | 0.011762645 |
| ENSG00000152926 | ZNF117 | 2317.707 | 0.720 | 0.005216025 |
| ENSG00000260941 | LINC00622 | 31.554 | 0.720 | 0.001764446 |
| ENSG00000280018 | | 3.704 | 0.719 | 0.013449747 |
| ENSG00000281344 | HELLPAR | 74.466 | 0.718 | 0.000124973 |
| ENSG00000233966 | UBE2SP1 | 46.104 | 0.718 | 0.000971038 |
| ENSG00000179363 | TMEM31 | 6.558 | 0.718 | 0.023394232 |
| ENSG00000099889 | ARVCF | 1183.218 | 0.715 | 3.43144E-05 |
| ENSG00000257696 | | 8.164 | 0.714 | 0.0111017 |
| ENSG00000169122 | FAM110B | 167.218 | 0.714 | 0.009668798 |
| ENSG00000168280 | KIF5C | 700.458 | 0.713 | 0.023638415 |
| ENSG00000236021 | | 6.778 | 0.712 | 0.000989315 |
| ENSG00000262484 | CCER2 | 23.802 | 0.711 | 0.011467941 |
| ENSG00000261126 | RBFADN | 45.754 | 0.711 | 0.000315093 |
| ENSG00000196503 | ARL9 | 72.999 | 0.710 | 0.012843068 |
| ENSG00000250254 | PTTG2 | 6.394 | 0.708 | 0.028879851 |
| ENSG00000172458 | IL17D | 95.139 | 0.707 | 0.000974875 |
| ENSG00000188383 | GPAT2P2 | 4.550 | 0.706 | 0.02572401 |
| ENSG00000187607 | ZNF286A | 273.716 | 0.705 | 2.2494E-11 |
| ENSG00000213085 | CFAP45 | 225.031 | 0.705 | 0.008408708 |
| ENSG00000269486 | ERVK9-11 | 13.465 | 0.704 | 0.002272737 |
| ENSG00000234345 | ELF2P1 | 4.367 | 0.704 | 0.01069268 |
| ENSG00000006118 | TMEM132A | 4386.951 | 0.704 | 1.19047E-05 |
| ENSG00000101265 | RASSF2 | 1407.364 | 0.703 | 0.001983014 |
| ENSG00000184270 | H2AC21 | 4.714 | 0.703 | 0.010090966 |
| ENSG00000162738 | VANGL2 | 2075.040 | 0.702 | 0.01326457 |
| ENSG00000112320 | SOBP | 191.672 | 0.701 | 0.003824482 |
| ENSG00000118503 | TNFAIP3 | 3106.114 | 0.701 | 0.002518274 |
| ENSG00000102104 | RS1 | 9.593 | 0.701 | 0.042144777 |
| ENSG00000167554 | ZNF610 | 194.195 | 0.701 | 0.001619995 |
| ENSG00000261204 | | 5.190 | 0.701 | 0.003149238 |
| ENSG00000137558 | PI15 | 238.400 | 0.701 | 0.033406203 |
| ENSG00000184619 | KRBA2 | 25.506 | 0.700 | 0.002721135 |
| ENSG00000220378 | KRT8P42 | 9.812 | 0.700 | 0.02373258 |
| ENSG00000183943 | PRKX | 1278.942 | 0.700 | 2.32366E-07 |
| ENSG00000138336 | TET1 | 107.486 | 0.698 | 0.000760899 |
| ENSG00000164627 | KIF6 | 20.192 | 0.697 | 0.019843894 |
| ENSG00000145506 | NKD2 | 666.414 | 0.697 | 0.035438689 |
| ENSG00000131242 | RAB11FIP4 | 3025.566 | 0.696 | 0.000291212 |
| ENSG00000163462 | TRIM46 | 146.484 | 0.694 | 0.000316626 |
| ENSG00000137338 | PGBD1 | 249.046 | 0.694 | 9.72653E-06 |
| ENSG00000105737 | GRIK5 | 87.587 | 0.693 | 0.016271117 |
| ENSG00000003249 | DBNDD1 | 1264.049 | 0.693 | 0.001329679 |
| ENSG00000178409 | BEND3 | 408.121 | 0.692 | 4.33381E-09 |
| ENSG00000167315 | ACAA2 | 2141.379 | 0.692 | 0.000725512 |
| ENSG00000144395 | CCDC150 | 169.875 | 0.691 | 1.00272E-05 |
| ENSG00000270948 | MTDHP1 | 15.718 | 0.691 | 0.01064431 |
| ENSG00000280020 | | 8.088 | 0.690 | 0.000112495 |
| ENSG00000164221 | CCDC112 | 318.874 | 0.690 | 7.70571E-09 |
| ENSG00000143184 | XCL1 | 129.276 | 0.690 | 0.023868362 |

| | | | | |
|---|---|---|---|---|
| ENSG00000232022 | FAAHP1 | 29.284 | 0.690 | 0.006015577 |
| ENSG00000240668 | KRT8P36 | 30.253 | 0.690 | 0.001313418 |
| ENSG00000226856 | THORLNC | 23.621 | 0.690 | 0.005688973 |
| ENSG00000233560 | KRT8P39 | 12.424 | 0.687 | 0.000746567 |
| ENSG00000007402 | CACNA2D2 | 117.436 | 0.687 | 0.001138935 |
| ENSG00000063176 | SPHK2 | 1072.776 | 0.686 | 1.47136E-06 |
| ENSG00000160716 | CHRNB2 | 10.522 | 0.685 | 0.012027351 |
| ENSG00000253395 | | 9.805 | 0.684 | 0.023714687 |
| ENSG00000269102 | | 5.936 | 0.684 | 0.003980974 |
| ENSG00000271936 | | 91.331 | 0.684 | 1.74379E-07 |
| ENSG00000176371 | ZSCAN2 | 384.531 | 0.684 | 1.64332E-09 |
| ENSG00000239415 | | 96.461 | 0.683 | 1.65626E-06 |
| ENSG00000206140 | TMEM191C | 49.277 | 0.682 | 0.021115255 |
| ENSG00000175352 | NRIP3 | 265.516 | 0.682 | 0.000989077 |
| ENSG00000162873 | KLHDC8A | 30.367 | 0.682 | 0.006879741 |
| ENSG00000162571 | TTLL10 | 24.290 | 0.680 | 0.024627424 |
| ENSG00000135414 | GDF11 | 950.616 | 0.680 | 2.53834E-06 |
| ENSG00000076382 | SPAG5 | 2571.350 | 0.680 | 4.72394E-08 |
| ENSG00000229891 | LINC01315 | 40.577 | 0.679 | 0.015467775 |
| ENSG00000144485 | HES6 | 417.902 | 0.679 | 0.002067733 |
| ENSG00000151692 | RNF144A | 1106.045 | 0.678 | 1.85712E-05 |
| ENSG00000167840 | ZNF232 | 410.616 | 0.677 | 9.87449E-10 |
| ENSG00000277938 | | 27.716 | 0.676 | 0.001345225 |
| ENSG00000203814 | H2BC18 | 169.441 | 0.676 | 0.018157405 |
| ENSG00000125571 | IL37 | 9.358 | 0.675 | 0.047069292 |
| ENSG00000224126 | UBE2SP2 | 5.984 | 0.675 | 0.000931439 |
| ENSG00000173991 | TCAP | 82.156 | 0.674 | 0.000767156 |
| ENSG00000230630 | DNM3OS | 128.586 | 0.674 | 0.017575505 |
| ENSG00000214465 | SMARCE1P6 | 8.140 | 0.673 | 0.024213378 |
| ENSG00000269155 | | 12.928 | 0.672 | 0.036884139 |
| ENSG00000239713 | APOBEC3G | 1778.792 | 0.671 | 0.001268658 |
| ENSG00000153395 | LPCAT1 | 6988.046 | 0.669 | 0.000447048 |
| ENSG00000234776 | C11orf94 | 5.520 | 0.668 | 0.001622544 |
| ENSG00000249459 | ZNF286B | 70.377 | 0.668 | 1.31958E-06 |
| ENSG00000227725 | GCOM2 | 4.532 | 0.668 | 0.032090116 |
| ENSG00000121690 | DEPDC7 | 202.352 | 0.667 | 0.035730444 |
| ENSG00000235072 | ARNILA | 16.528 | 0.667 | 0.004180806 |
| ENSG00000205704 | LINC00634 | 16.607 | 0.666 | 0.004501578 |
| ENSG00000218073 | | 9.549 | 0.665 | 0.036617736 |
| ENSG00000267041 | ZNF850 | 213.369 | 0.665 | 0.000204359 |
| ENSG00000237594 | | 84.687 | 0.664 | 0.004265887 |
| ENSG00000152284 | TCF7L1 | 825.212 | 0.663 | 0.006322796 |
| ENSG00000085662 | AKR1B1 | 10453.445 | 0.662 | 0.006705868 |
| ENSG00000088305 | DNMT3B | 932.549 | 0.662 | 0.000858384 |
| ENSG00000240288 | GHRLOS | 60.616 | 0.661 | 0.001460998 |
| ENSG00000152582 | SPEF2 | 135.104 | 0.661 | 0.030460545 |
| ENSG00000129646 | QRICH2 | 200.460 | 0.659 | 0.000489502 |
| ENSG00000206432 | TMEM200C | 55.126 | 0.659 | 0.045844753 |
| ENSG00000196793 | ZNF239 | 244.524 | 0.659 | 0.000336961 |
| ENSG00000270638 | | 8.705 | 0.659 | 0.001897864 |
| ENSG00000139174 | PRICKLE1 | 273.772 | 0.659 | 0.019843894 |
| ENSG00000275720 | | 12.317 | 0.658 | 0.021160333 |
| ENSG00000225439 | BOLA3-AS1 | 106.039 | 0.657 | 0.002621294 |
| ENSG00000114279 | FGF12 | 207.187 | 0.656 | 0.025199256 |
| ENSG00000101280 | ANGPT4 | 27.205 | 0.656 | 0.03816747 |
| ENSG00000279692 | | 69.075 | 0.656 | 0.017285542 |
| ENSG00000135119 | RNFT2 | 278.958 | 0.656 | 0.001568565 |
| ENSG00000080709 | KCNN2 | 49.404 | 0.655 | 0.021450748 |
| ENSG00000157827 | FMNL2 | 1490.066 | 0.655 | 0.000521222 |

| | | | | |
|---|---|---|---|---|
| ENSG00000250510 | GPR162 | 109.758 | 0.655 | 0.004678812 |
| ENSG00000134253 | TRIM45 | 625.234 | 0.655 | 9.6398E-06 |
| ENSG00000267199 | | 13.443 | 0.655 | 0.014746118 |
| ENSG00000231806 | PCAT7 | 89.744 | 0.654 | 0.010420465 |
| ENSG00000234719 | NPIPB2 | 38.195 | 0.654 | 0.032091312 |
| ENSG00000266916 | ZNF793-AS1 | 107.065 | 0.654 | 0.004014223 |
| ENSG00000116852 | KIF21B | 487.273 | 0.653 | 9.91236E-05 |
| ENSG00000129810 | SGO1 | 481.140 | 0.652 | 6.71828E-06 |
| ENSG00000230042 | AK3P3 | 6.038 | 0.652 | 0.001841634 |
| ENSG00000175305 | CCNE2 | 300.021 | 0.652 | 0.000460319 |
| ENSG00000132434 | LANCL2 | 1256.994 | 0.652 | 4.38898E-07 |
| ENSG00000213160 | KLHL23 | 679.405 | 0.651 | 5.87454E-05 |
| ENSG00000271947 | | 7.643 | 0.651 | 0.028550225 |
| ENSG00000264968 | | 7.177 | 0.651 | 0.021860765 |
| ENSG00000167874 | TMEM88 | 156.033 | 0.651 | 0.000265686 |
| ENSG00000175322 | ZNF519 | 225.691 | 0.650 | 9.40561E-06 |
| ENSG00000187987 | ZSCAN23 | 32.842 | 0.650 | 0.035478364 |
| ENSG00000229689 | | 109.379 | 0.649 | 0.004348082 |
| ENSG00000224420 | ADM5 | 145.895 | 0.649 | 0.001235284 |
| ENSG00000127561 | SYNGR3 | 125.467 | 0.649 | 0.044462529 |
| ENSG00000111879 | FAM184A | 113.347 | 0.649 | 0.008489274 |
| ENSG00000040933 | INPP4A | 2258.318 | 0.649 | 4.13192E-06 |
| ENSG00000254477 | | 16.489 | 0.648 | 0.0111017 |
| ENSG00000214077 | GNAQP1 | 4.468 | 0.647 | 0.019925999 |
| ENSG00000147180 | ZNF711 | 452.024 | 0.647 | 0.005502933 |
| ENSG00000213096 | ZNF254 | 897.447 | 0.646 | 0.000127693 |
| ENSG00000180336 | MEIOC | 24.670 | 0.646 | 0.001194947 |
| ENSG00000168772 | CXXC4 | 120.380 | 0.645 | 0.042028123 |
| ENSG00000169607 | CKAP2L | 885.261 | 0.645 | 5.72661E-05 |
| ENSG00000261215 | | 7.706 | 0.645 | 0.007421838 |
| ENSG00000271387 | C1orf21-DT | 14.056 | 0.645 | 0.005369132 |
| ENSG00000171208 | NETO2 | 1157.437 | 0.644 | 0.004014223 |
| ENSG00000135643 | KCNMB4 | 548.622 | 0.644 | 0.018139669 |
| ENSG00000106069 | CHN2 | 448.850 | 0.644 | 0.00508104 |
| ENSG00000276529 | | 129.002 | 0.643 | 2.94155E-05 |
| ENSG00000147536 | GINS4 | 392.693 | 0.643 | 4.43592E-05 |
| ENSG00000160360 | GPSM1 | 758.369 | 0.643 | 0.001476721 |
| ENSG00000175832 | ETV4 | 1939.387 | 0.641 | 0.007268369 |
| ENSG00000159164 | SV2A | 505.287 | 0.641 | 0.018106846 |
| ENSG00000231359 | | 17.909 | 0.641 | 0.020931603 |
| ENSG00000259917 | HNRNPLP2 | 35.145 | 0.640 | 0.000279441 |
| ENSG00000241717 | VWFP1 | 7.508 | 0.639 | 0.042878413 |
| ENSG00000169679 | BUB1 | 1810.632 | 0.639 | 8.64516E-07 |
| ENSG00000115507 | OTX1 | 251.986 | 0.638 | 0.000823046 |
| ENSG00000182264 | IZUMO1 | 17.774 | 0.638 | 0.022632447 |
| ENSG00000128596 | CCDC136 | 149.458 | 0.637 | 0.00480316 |
| ENSG00000149557 | FEZ1 | 934.016 | 0.637 | 0.011435054 |
| ENSG00000137343 | ATAT1 | 1048.940 | 0.636 | 1.24458E-06 |
| ENSG00000082684 | SEMA5B | 195.419 | 0.635 | 0.023540375 |
| ENSG00000161509 | GRIN2C | 41.219 | 0.634 | 0.014916655 |
| ENSG00000168298 | H1-4 | 51.307 | 0.634 | 0.027176046 |
| ENSG00000213260 | YWHAZP5 | 68.222 | 0.634 | 0.002438976 |
| ENSG00000253978 | CTB-178M22.2 | 7.739 | 0.633 | 0.032486473 |
| ENSG00000143882 | ATP6V1C2 | 203.955 | 0.633 | 0.0024848 |
| ENSG00000271888 | JARID2-DT | 32.445 | 0.633 | 0.00016036 |
| ENSG00000121764 | HCRTR1 | 8.048 | 0.633 | 0.016890123 |
| ENSG00000042781 | USH2A | 10.992 | 0.633 | 0.010500023 |
| ENSG00000166900 | STX3 | 2839.096 | 0.633 | 8.41755E-07 |
| ENSG00000006042 | TMEM98 | 1980.905 | 0.632 | 0.002286871 |

| | | | | |
|---|---|---|---|---|
| ENSG00000273230 | | 103.994 | 0.632 | 0.000651425 |
| ENSG00000184530 | C6orf58 | 6.177 | 0.631 | 0.004444873 |
| ENSG00000184635 | ZNF93 | 480.224 | 0.630 | 0.000628604 |
| ENSG00000120756 | PLS1 | 1638.327 | 0.630 | 0.005484771 |
| ENSG00000166532 | RIMKLB | 1047.548 | 0.629 | 0.000631475 |
| ENSG00000259845 | HERC2P10 | 5.028 | 0.628 | 0.033380651 |
| ENSG00000272275 | | 27.828 | 0.627 | 0.003465073 |
| ENSG00000188610 | FAM72B | 112.920 | 0.627 | 0.00019003 |
| ENSG00000130347 | RTN4IP1 | 446.235 | 0.626 | 1.52408E-06 |
| ENSG00000258376 | | 72.909 | 0.626 | 0.023928057 |
| ENSG00000227740 | LINC02803 | 19.300 | 0.625 | 0.0105216 |
| ENSG00000197238 | H4C11 | 7.925 | 0.625 | 0.027990922 |
| ENSG00000165730 | STOX1 | 158.252 | 0.625 | 0.009468191 |
| ENSG00000133874 | RNF122 | 466.704 | 0.624 | 9.92006E-06 |
| ENSG00000272540 | | 20.436 | 0.624 | 1.76505E-05 |
| ENSG00000259531 | | 5.977 | 0.624 | 0.026274767 |
| ENSG00000248774 | | 6.343 | 0.624 | 0.012027351 |
| ENSG00000115112 | TFCP2L1 | 908.202 | 0.623 | 0.030565976 |
| ENSG00000151023 | ENKUR | 26.418 | 0.623 | 0.027132809 |
| ENSG00000112137 | PHACTR1 | 226.778 | 0.623 | 0.00785892 |
| ENSG00000228794 | LINC01128 | 576.207 | 0.623 | 5.90099E-06 |
| ENSG00000141519 | CCDC40 | 215.994 | 0.623 | 0.000105321 |
| ENSG00000113212 | PCDHB7 | 125.535 | 0.623 | 0.024339786 |
| ENSG00000166793 | YPEL4 | 60.065 | 0.622 | 0.000977349 |
| ENSG00000144712 | CAND2 | 228.160 | 0.622 | 0.024942814 |
| ENSG00000124374 | PAIP2B | 541.882 | 0.622 | 0.000201977 |
| ENSG00000251050 | RBX1P2 | 6.772 | 0.622 | 0.028112937 |
| ENSG00000165821 | SALL2 | 423.871 | 0.622 | 0.010093137 |
| ENSG00000163235 | TGFA | 2141.506 | 0.622 | 0.00509627 |
| ENSG00000271109 | | 39.267 | 0.621 | 0.00534063 |
| ENSG00000279641 | | 13.940 | 0.620 | 0.000114851 |
| ENSG00000106633 | GCK | 16.361 | 0.619 | 0.046814921 |
| ENSG00000185252 | ZNF74 | 964.710 | 0.619 | 9.49609E-08 |
| ENSG00000229932 | YWHAZP3 | 20.324 | 0.619 | 0.002133395 |
| ENSG00000228010 | | 10.152 | 0.619 | 0.006015577 |
| ENSG00000096654 | ZNF184 | 504.997 | 0.619 | 4.04947E-11 |
| ENSG00000258101 | | 16.827 | 0.617 | 0.000603104 |
| ENSG00000250067 | YJEFN3 | 310.023 | 0.617 | 0.009723142 |
| ENSG00000188599 | NPIPP1 | 201.189 | 0.616 | 0.00016036 |
| ENSG00000176788 | BASP1 | 5238.005 | 0.616 | 0.025702226 |
| ENSG00000167535 | CACNB3 | 2409.984 | 0.616 | 6.69322E-06 |
| ENSG00000108852 | MPP2 | 287.008 | 0.616 | 0.007669961 |
| ENSG00000172350 | ABCG4 | 19.917 | 0.615 | 0.014853865 |
| ENSG00000132016 | BRME1 | 255.057 | 0.615 | 0.002203441 |
| ENSG00000272128 | | 10.860 | 0.615 | 0.018185479 |
| ENSG00000274997 | H2AC12 | 3.924 | 0.615 | 0.029168991 |
| ENSG00000272462 | | 126.090 | 0.614 | 0.000189933 |
| ENSG00000160191 | PDE9A | 904.396 | 0.614 | 0.030921208 |
| ENSG00000263235 | | 25.118 | 0.614 | 0.002433679 |
| ENSG00000105173 | CCNE1 | 1370.769 | 0.614 | 0.001280606 |
| ENSG00000258317 | | 8.770 | 0.614 | 0.008110955 |
| ENSG00000147872 | PLIN2 | 4503.261 | 0.613 | 0.002457302 |
| ENSG00000175820 | CCDC168 | 5.892 | 0.613 | 0.016052872 |
| ENSG00000228126 | FALEC | 4.766 | 0.612 | 0.030707777 |
| ENSG00000260167 | | 8.583 | 0.611 | 0.002179856 |
| ENSG00000084731 | KIF3C | 1577.736 | 0.611 | 2.10567E-05 |
| ENSG00000271524 | BNIP3P17 | 24.133 | 0.611 | 0.035201594 |
| ENSG00000005187 | ACSM3 | 448.549 | 0.611 | 0.046996908 |
| ENSG00000273209 | | 15.430 | 0.610 | 0.022535 |

| | | | | |
|---|---|---|---|---|
| ENSG00000253392 | | 18.633 | 0.610 | 0.0084105 |
| ENSG00000166503 | HDGFL3 | 2016.076 | 0.609 | 6.19677E-05 |
| ENSG00000272931 | LRRC8D-DT | 20.122 | 0.609 | 0.004598212 |
| ENSG00000257950 | P2RX5-TAX1BP3 | 37.788 | 0.609 | 7.34287E-07 |
| ENSG00000272402 | | 21.031 | 0.608 | 0.000267793 |
| ENSG00000261587 | TMEM249 | 12.873 | 0.607 | 0.008257545 |
| ENSG00000229739 | PDC-AS1 | 6.984 | 0.606 | 0.007163031 |
| ENSG00000196917 | HCAR1 | 920.698 | 0.606 | 0.043952792 |
| ENSG00000227201 | CNN2P1 | 13.151 | 0.605 | 0.014531285 |
| ENSG00000230606 | | 11.621 | 0.605 | 0.040335522 |
| ENSG00000169885 | CALML6 | 31.730 | 0.605 | 0.035225515 |
| ENSG00000088325 | TPX2 | 5606.445 | 0.604 | 0.000129244 |
| ENSG00000166226 | CCT2 | 12708.342 | 0.604 | 0.00152339 |
| ENSG00000109099 | PMP22 | 2664.836 | 0.604 | 0.016008085 |
| ENSG00000124789 | NUP153 | 3005.917 | 0.604 | 4.11822E-08 |
| ENSG00000267565 | | 38.989 | 0.603 | 0.003148989 |
| ENSG00000126217 | MCF2L | 1763.278 | 0.603 | 0.007669961 |
| ENSG00000274750 | H3C6 | 35.561 | 0.603 | 0.015703493 |
| ENSG00000265415 | | 132.876 | 0.602 | 0.00120993 |
| ENSG00000232369 | PPIAP35 | 4.984 | 0.601 | 0.009908836 |
| ENSG00000119471 | HSDL2 | 3179.711 | 0.601 | 5.26672E-05 |
| ENSG00000153982 | GDPD1 | 234.948 | 0.601 | 0.000208337 |
| ENSG00000241155 | ARHGAP31-AS1 | 11.990 | 0.600 | 0.014497696 |
| ENSG00000000460 | C1orf112 | 711.289 | 0.599 | 5.55099E-05 |
| ENSG00000124140 | SLC12A5 | 51.918 | 0.599 | 0.027110625 |
| ENSG00000196081 | ZNF724 | 74.688 | 0.599 | 0.002272737 |
| ENSG00000111752 | PHC1 | 432.486 | 0.598 | 1.34734E-06 |
| ENSG00000117477 | CCDC181 | 65.414 | 0.598 | 0.040870034 |
| ENSG00000267980 | | 41.509 | 0.597 | 0.000115192 |
| ENSG00000204815 | ODAD4 | 73.253 | 0.597 | 0.002448829 |
| ENSG00000023330 | ALAS1 | 5951.926 | 0.596 | 0.000447013 |
| ENSG00000176358 | TAC4 | 13.071 | 0.596 | 0.047787397 |
| ENSG00000273786 | | 4.679 | 0.594 | 0.029950456 |
| ENSG00000112294 | ALDH5A1 | 2401.989 | 0.593 | 0.002810843 |
| ENSG00000227542 | | 7.548 | 0.593 | 0.005400645 |
| ENSG00000260727 | SLC7A5P1 | 6.713 | 0.593 | 0.008786183 |
| ENSG00000154839 | SKA1 | 691.039 | 0.593 | 3.72085E-05 |
| ENSG00000250251 | PKD1P6 | 182.478 | 0.592 | 0.000408727 |
| ENSG00000213131 | YWHAZP4 | 61.653 | 0.592 | 0.000906664 |
| ENSG00000165185 | KIAA1958 | 439.210 | 0.592 | 1.50389E-05 |
| ENSG00000187801 | ZFP69B | 122.436 | 0.592 | 3.54873E-05 |
| ENSG00000143228 | NUF2 | 1150.412 | 0.591 | 0.000304642 |
| ENSG00000217624 | YWHAZP10 | 40.757 | 0.591 | 0.004092005 |
| ENSG00000255397 | NASPP1 | 4.992 | 0.591 | 0.00061471 |
| ENSG00000186648 | CARMIL3 | 85.084 | 0.590 | 0.022072356 |
| ENSG00000146555 | SDK1 | 1529.664 | 0.590 | 0.020149201 |
| ENSG00000270996 | | 4.606 | 0.589 | 0.036617736 |
| ENSG00000118298 | CA14 | 37.985 | 0.589 | 0.000614863 |
| ENSG00000254285 | KRT8P3 | 178.264 | 0.589 | 0.030662342 |
| ENSG00000108106 | UBE2S | 2648.588 | 0.589 | 4.73268E-05 |
| ENSG00000272417 | | 8.585 | 0.588 | 0.032787321 |
| ENSG00000214578 | HMGN2P15 | 16.214 | 0.588 | 0.020077498 |
| ENSG00000141505 | ASGR1 | 101.099 | 0.587 | 0.005984602 |
| ENSG00000270885 | RASL10B | 121.597 | 0.586 | 0.043837637 |
| ENSG00000159023 | EPB41 | 2747.796 | 0.585 | 0.000111163 |
| ENSG00000197099 | | 5.817 | 0.585 | 0.019496104 |
| ENSG00000239523 | MYLK-AS1 | 48.398 | 0.584 | 0.000279441 |
| ENSG00000167992 | VWCE | 95.347 | 0.583 | 0.013975102 |
| ENSG00000157833 | GAREM2 | 515.001 | 0.583 | 0.001619872 |

| | | | | |
|---|---|---|---|---|
| ENSG00000088756 | ARHGAP28 | 259.705 | 0.583 | 0.036414394 |
| ENSG00000184678 | H2BC21 | 1864.645 | 0.581 | 0.009468191 |
| ENSG00000167524 | RSKR | 217.488 | 0.581 | 0.002099161 |
| ENSG00000101695 | RNF125 | 229.662 | 0.581 | 0.003799228 |
| ENSG00000197748 | CFAP43 | 41.432 | 0.580 | 0.004898627 |
| ENSG00000197299 | BLM | 650.366 | 0.580 | 1.94959E-07 |
| ENSG00000254536 | | 4.701 | 0.580 | 0.014912615 |
| ENSG00000197093 | GAL3ST4 | 569.850 | 0.579 | 0.002810843 |
| ENSG00000128408 | RIBC2 | 170.825 | 0.579 | 0.002009499 |
| ENSG00000147862 | NFIB | 1797.339 | 0.579 | 0.001850381 |
| ENSG00000205622 | | 31.780 | 0.579 | 0.044275973 |
| ENSG00000162367 | TAL1 | 89.703 | 0.579 | 0.01069268 |
| ENSG00000166856 | GPR182 | 8.846 | 0.578 | 0.008558202 |
| ENSG00000271914 | | 11.424 | 0.578 | 0.002605743 |
| ENSG00000196208 | GREB1 | 145.426 | 0.578 | 0.032102779 |
| ENSG00000116128 | BCL9 | 1668.616 | 0.578 | 3.71009E-07 |
| ENSG00000273080 | | 25.837 | 0.578 | 0.000380599 |
| ENSG00000099338 | CATSPERG | 132.088 | 0.577 | 0.009468191 |
| ENSG00000196418 | ZNF124 | 271.595 | 0.577 | 0.000272707 |
| ENSG00000105613 | MAST1 | 158.370 | 0.577 | 0.028600424 |
| ENSG00000186185 | KIF18B | 1300.074 | 0.576 | 9.97149E-05 |
| ENSG00000215283 | HMGB3P24 | 12.545 | 0.573 | 0.001264473 |
| ENSG00000071051 | NCK2 | 2854.378 | 0.572 | 1.19272E-11 |
| ENSG00000120647 | CCDC77 | 552.339 | 0.572 | 1.91664E-08 |
| ENSG00000184792 | OSBP2 | 987.499 | 0.572 | 0.033113603 |
| ENSG00000156509 | FBXO43 | 57.141 | 0.571 | 0.0119868 |
| ENSG00000147874 | HAUS6 | 1127.167 | 0.571 | 1.63782E-06 |
| ENSG00000259172 | | 30.759 | 0.570 | 0.007788237 |
| ENSG00000160298 | C21orf58 | 611.104 | 0.570 | 0.000378268 |
| ENSG00000227470 | RPL39P16 | 3.681 | 0.570 | 0.013400315 |
| ENSG00000180574 | EIF2S3B | 17.037 | 0.569 | 0.015466871 |
| ENSG00000237976 | RFX5-AS1 | 21.900 | 0.569 | 0.006762577 |
| ENSG00000154479 | CCDC173 | 14.085 | 0.569 | 0.010624959 |
| ENSG00000225125 | RANP4 | 10.106 | 0.569 | 0.020796243 |
| ENSG00000250326 | | 7.731 | 0.568 | 0.006365077 |
| ENSG00000173261 | PLAC8L1 | 15.112 | 0.568 | 0.020070425 |
| ENSG00000269834 | ZNF528-AS1 | 425.352 | 0.568 | 0.010047183 |
| ENSG00000121152 | NCAPH | 1143.773 | 0.567 | 8.35926E-05 |
| ENSG00000050393 | MCUR1 | 1941.214 | 0.567 | 1.51218E-09 |
| ENSG00000198075 | SULT1C4 | 26.674 | 0.567 | 0.024059603 |
| ENSG00000135476 | ESPL1 | 1418.796 | 0.566 | 0.000199601 |
| ENSG00000080298 | RFX3 | 415.331 | 0.566 | 7.56294E-05 |
| ENSG00000109881 | CCDC34 | 859.798 | 0.565 | 4.06442E-06 |
| ENSG00000223393 | | 10.314 | 0.565 | 0.007863784 |
| ENSG00000260368 | | 27.425 | 0.564 | 0.005401527 |
| ENSG00000246859 | STARD4-AS1 | 51.740 | 0.564 | 0.024622559 |
| ENSG00000176076 | KCNE5 | 11.715 | 0.563 | 0.021656392 |
| ENSG00000128011 | LRFN1 | 173.335 | 0.562 | 0.025199256 |
| ENSG00000186907 | RTN4RL2 | 67.788 | 0.562 | 0.019496104 |
| ENSG00000159556 | ISL2 | 99.940 | 0.562 | 0.006213269 |
| ENSG00000146267 | FAXC | 205.645 | 0.562 | 0.033382722 |
| ENSG00000107249 | GLIS3 | 440.630 | 0.562 | 0.026184813 |
| ENSG00000092853 | CLSPN | 460.007 | 0.561 | 0.004882371 |
| ENSG00000178999 | AURKB | 1712.483 | 0.561 | 0.000289934 |
| ENSG00000233757 | | 21.256 | 0.560 | 0.001613795 |
| ENSG00000144671 | SLC22A14 | 13.759 | 0.560 | 0.023319465 |
| ENSG00000157045 | NTAN1 | 1864.402 | 0.560 | 5.98089E-06 |
| ENSG00000101057 | MYBL2 | 5153.004 | 0.559 | 0.003726507 |
| ENSG00000233246 | PHC2-AS1 | 5.421 | 0.559 | 0.035617632 |

| | | | | |
|---|---|---|---|---|
| ENSG00000170264 | FAM161A | 217.952 | 0.559 | 0.000152513 |
| ENSG00000197457 | STMN3 | 1144.992 | 0.558 | 0.046296692 |
| ENSG00000166352 | IFTAP | 374.126 | 0.558 | 6.48025E-06 |
| ENSG00000186193 | SAPCD2 | 1576.290 | 0.557 | 0.003661524 |
| ENSG00000197291 | RAMP2-AS1 | 44.429 | 0.557 | 0.039335515 |
| ENSG00000187186 | | 29.312 | 0.557 | 0.006671094 |
| ENSG00000232934 | | 41.621 | 0.557 | 0.013779325 |
| ENSG00000270580 | PKD1P6-NPIPP1 | 72.172 | 0.556 | 7.7086E-05 |
| ENSG00000230658 | KLHL7-DT | 34.449 | 0.556 | 0.020518289 |
| ENSG00000230002 | ALMS1-IT1 | 45.279 | 0.556 | 0.002610707 |
| ENSG00000273045 | C2orf15 | 200.857 | 0.556 | 0.004633575 |
| ENSG00000180998 | GPR137C | 108.640 | 0.555 | 0.021011475 |
| ENSG00000236519 | LINC01424 | 12.233 | 0.555 | 0.004712992 |
| ENSG00000139354 | GAS2L3 | 410.808 | 0.555 | 0.00043316 |
| ENSG00000166289 | PLEKHF1 | 1968.516 | 0.555 | 0.027249848 |
| ENSG00000244184 | | 13.077 | 0.555 | 0.00694236 |
| ENSG00000121957 | GPSM2 | 1305.944 | 0.554 | 0.000111743 |
| ENSG00000224728 | IMPDH1P8 | 11.873 | 0.554 | 0.010514269 |
| ENSG00000258830 | | 6.195 | 0.554 | 0.031808359 |
| ENSG00000168993 | CPLX1 | 104.552 | 0.554 | 0.012358086 |
| ENSG00000233155 | HMGA1P8 | 7.759 | 0.553 | 0.002457302 |
| ENSG00000180694 | TMEM64 | 2139.227 | 0.552 | 0.017786834 |
| ENSG00000137185 | ZSCAN9 | 756.014 | 0.551 | 1.4666E-06 |
| ENSG00000082397 | EPB41L3 | 818.776 | 0.550 | 0.047867242 |
| ENSG00000272263 | | 8.191 | 0.550 | 0.016664218 |
| ENSG00000162814 | SPATA17 | 162.242 | 0.550 | 0.022111561 |
| ENSG00000234996 | ACTG1P25 | 30.174 | 0.548 | 0.010946144 |
| ENSG00000108666 | C17orf75 | 1078.789 | 0.548 | 4.11822E-08 |
| ENSG00000227372 | TP73-AS1 | 389.116 | 0.548 | 0.018408594 |
| ENSG00000270571 | | 21.782 | 0.548 | 0.022025812 |
| ENSG00000279266 | | 5.843 | 0.548 | 0.042201594 |
| ENSG00000101306 | MYLK2 | 28.746 | 0.548 | 0.005323567 |
| ENSG00000265800 | | 23.112 | 0.547 | 0.00156076 |
| ENSG00000163808 | KIF15 | 790.645 | 0.547 | 0.00020988 |
| ENSG00000065328 | MCM10 | 793.488 | 0.546 | 0.001019191 |
| ENSG00000133065 | SLC41A1 | 2213.599 | 0.546 | 6.0544E-05 |
| ENSG00000068489 | PRR11 | 1350.613 | 0.545 | 0.000452517 |
| ENSG00000267707 | | 11.009 | 0.545 | 0.036617736 |
| ENSG00000005448 | WDR54 | 1132.995 | 0.545 | 0.000159362 |
| ENSG00000011332 | DPF1 | 72.923 | 0.545 | 0.041283268 |
| ENSG00000273473 | | 10.757 | 0.544 | 0.013123801 |
| ENSG00000162929 | KIAA1841 | 818.238 | 0.544 | 3.16138E-07 |
| ENSG00000272562 | H3-3A-DT | 5.755 | 0.544 | 0.012529782 |
| ENSG00000170522 | ELOVL6 | 2194.690 | 0.544 | 0.003490926 |
| ENSG00000278627 | | 9.328 | 0.543 | 0.021023226 |
| ENSG00000213760 | ATP6V1G2 | 26.176 | 0.543 | 0.009931972 |
| ENSG00000001460 | STPG1 | 965.838 | 0.543 | 0.000204132 |
| ENSG00000106976 | DNM1 | 849.867 | 0.543 | 0.022576053 |
| ENSG00000170921 | TANC2 | 3756.154 | 0.542 | 0.000279441 |
| ENSG00000100312 | ACR | 4.451 | 0.542 | 0.043648657 |
| ENSG00000101224 | CDC25B | 4682.249 | 0.542 | 0.007602949 |
| ENSG00000131080 | EDA2R | 124.772 | 0.542 | 0.030567724 |
| ENSG00000230185 | C9orf147 | 13.231 | 0.542 | 0.000200759 |
| ENSG00000204366 | ZBTB12 | 467.525 | 0.541 | 6.56361E-05 |
| ENSG00000139890 | REM2 | 38.283 | 0.541 | 0.002133395 |
| ENSG00000227210 | | 9.638 | 0.541 | 0.01069268 |
| ENSG00000132510 | KDM6B | 3068.021 | 0.541 | 3.08817E-06 |
| ENSG00000261335 | | 10.287 | 0.540 | 0.005688973 |
| ENSG00000237094 | | 25.645 | 0.540 | 0.024986517 |

| | | | | |
|---|---|---|---|---|
| ENSG00000166596 | CFAP52 | 17.473 | 0.540 | 0.014724678 |
| ENSG00000213694 | S1PR3 | 1290.329 | 0.540 | 0.041998935 |
| ENSG00000152270 | PDE3B | 589.462 | 0.539 | 0.029036172 |
| ENSG00000010017 | RANBP9 | 2788.164 | 0.539 | 1.99503E-09 |
| ENSG00000263513 | FAM72C | 52.764 | 0.539 | 0.004254273 |
| ENSG00000187605 | TET3 | 2505.019 | 0.539 | 4.61554E-05 |
| ENSG00000134594 | RAB33A | 53.219 | 0.538 | 0.013562181 |
| ENSG00000158402 | CDC25C | 381.160 | 0.538 | 0.000167905 |
| ENSG00000138092 | CENPO | 778.495 | 0.538 | 6.332E-08 |
| ENSG00000214776 | | 64.004 | 0.538 | 0.006730571 |
| ENSG00000106070 | GRB10 | 1453.877 | 0.537 | 0.005655432 |
| ENSG00000102221 | JADE3 | 844.394 | 0.537 | 4.39087E-06 |
| ENSG00000130783 | CCDC62 | 12.672 | 0.536 | 0.005326481 |
| ENSG00000236901 | MIR600HG | 179.748 | 0.535 | 0.004993427 |
| ENSG00000269737 | | 43.521 | 0.535 | 0.033971922 |
| ENSG00000103056 | SMPD3 | 244.617 | 0.535 | 0.017125074 |
| ENSG00000112304 | ACOT13 | 1358.935 | 0.535 | 2.04235E-07 |
| ENSG00000179935 | LINC00652 | 11.441 | 0.534 | 0.049419352 |
| ENSG00000156299 | TIAM1 | 2012.605 | 0.534 | 0.003185946 |
| ENSG00000213713 | PIGCP1 | 60.604 | 0.533 | 0.007579917 |
| ENSG00000008226 | DLEC1 | 78.384 | 0.533 | 0.038266238 |
| ENSG00000120833 | SOCS2 | 1014.605 | 0.532 | 0.031015636 |
| ENSG00000129749 | CHRNA10 | 42.916 | 0.532 | 0.003502682 |
| ENSG00000136449 | MYCBPAP | 42.309 | 0.531 | 0.020446905 |
| ENSG00000146757 | ZNF92 | 520.783 | 0.530 | 1.55478E-05 |
| ENSG00000198513 | ATL1 | 267.869 | 0.529 | 0.00163025 |
| ENSG00000119772 | DNMT3A | 2303.666 | 0.529 | 7.07128E-07 |
| ENSG00000131370 | SH3BP5 | 441.346 | 0.529 | 0.008337088 |
| ENSG00000127946 | HIP1 | 1843.972 | 0.528 | 5.73393E-05 |
| ENSG00000261437 | LINC02894 | 163.722 | 0.527 | 0.018860521 |
| ENSG00000128394 | APOBEC3F | 489.351 | 0.527 | 0.000196701 |
| ENSG00000127589 | TUBBP1 | 56.138 | 0.527 | 1.9027E-05 |
| ENSG00000229525 | | 4.999 | 0.527 | 0.031891635 |
| ENSG00000205853 | RFPL3S | 16.801 | 0.526 | 0.023454808 |
| ENSG00000180385 | EMC3-AS1 | 230.395 | 0.526 | 0.002522424 |
| ENSG00000231084 | RPL22P24 | 20.182 | 0.525 | 5.08248E-05 |
| ENSG00000197279 | ZNF165 | 441.308 | 0.525 | 0.002005179 |
| ENSG00000213236 | YWHAZP2 | 15.982 | 0.524 | 0.005222747 |
| ENSG00000117724 | CENPF | 4161.229 | 0.524 | 0.000513067 |
| ENSG00000108671 | PSMD11 | 7320.712 | 0.524 | 5.75663E-08 |
| ENSG00000065600 | PACC1 | 752.795 | 0.523 | 8.2392E-08 |
| ENSG00000196591 | HDAC2 | 6168.908 | 0.523 | 5.46221E-11 |
| ENSG00000185055 | EFCAB10 | 29.673 | 0.522 | 0.001039452 |
| ENSG00000177602 | HASPIN | 200.248 | 0.522 | 0.001974571 |
| ENSG00000197951 | ZNF71 | 538.363 | 0.522 | 0.001093557 |
| ENSG00000137563 | GGH | 2329.332 | 0.522 | 0.005552243 |
| ENSG00000144031 | ANKRD53 | 25.465 | 0.521 | 0.016915399 |
| ENSG00000159086 | PAXBP1 | 1799.722 | 0.520 | 9.34279E-06 |
| ENSG00000106665 | CLIP2 | 1546.329 | 0.520 | 0.001850381 |
| ENSG00000267278 | MAP3K14-AS1 | 137.664 | 0.520 | 0.002033711 |
| ENSG00000154099 | DNAAF1 | 30.384 | 0.520 | 0.021600326 |
| ENSG00000133740 | E2F5 | 508.911 | 0.520 | 0.000510749 |
| ENSG00000234327 | ZNF232-AS1 | 94.852 | 0.518 | 0.005400645 |
| ENSG00000196781 | TLE1 | 1864.492 | 0.518 | 0.001684444 |
| ENSG00000186106 | ANKRD46 | 1139.835 | 0.517 | 0.001610314 |
| ENSG00000126858 | RHOT1 | 1656.373 | 0.517 | 6.00016E-09 |
| ENSG00000198521 | ZNF43 | 1009.252 | 0.517 | 0.012079357 |
| ENSG00000267834 | | 18.792 | 0.517 | 0.029150168 |
| ENSG00000172731 | LRRC20 | 727.432 | 0.517 | 0.000497746 |

| | | | | |
|---|---|---|---|---|
| ENSG00000215417 | MIR17HG | 57.525 | 0.517 | 0.018255115 |
| ENSG00000256897 | | 6.050 | 0.516 | 0.042861159 |
| ENSG00000234882 | EIF3EP1 | 50.763 | 0.516 | 0.005777655 |
| ENSG00000230454 | | 40.416 | 0.515 | 0.028887097 |
| ENSG00000180884 | ZNF792 | 290.565 | 0.514 | 0.000449195 |
| ENSG00000254806 | SYS1-DBNDD2 | 8.936 | 0.512 | 0.003887309 |
| ENSG00000077616 | NAALAD2 | 52.061 | 0.512 | 0.006565438 |
| ENSG00000151490 | PTPRO | 136.345 | 0.512 | 0.042861159 |
| ENSG00000271789 | | 20.105 | 0.511 | 0.006873614 |
| ENSG00000121621 | KIF18A | 447.880 | 0.511 | 0.002044279 |
| ENSG00000273064 | | 24.941 | 0.511 | 0.003348371 |
| ENSG00000265806 | MIR4292 | 15.477 | 0.511 | 0.001094328 |
| ENSG00000232640 | | 70.284 | 0.510 | 0.000200323 |
| ENSG00000108984 | MAP2K6 | 299.290 | 0.510 | 0.004926895 |
| ENSG00000225921 | NOL7 | 2928.925 | 0.509 | 8.51096E-08 |
| ENSG00000176208 | ATAD5 | 522.212 | 0.509 | 8.01469E-05 |
| ENSG00000243024 | | 10.571 | 0.508 | 0.039121481 |
| ENSG00000105204 | DYRK1B | 1648.466 | 0.507 | 0.000698413 |
| ENSG00000075975 | MKRN2 | 1766.130 | 0.507 | 2.73113E-06 |
| ENSG00000254485 | | 18.545 | 0.507 | 0.036404761 |
| ENSG00000148308 | GTF3C5 | 4210.778 | 0.507 | 9.20432E-07 |
| ENSG00000154743 | TSEN2 | 849.012 | 0.505 | 0.000847297 |
| ENSG00000140534 | TICRR | 621.047 | 0.505 | 0.00051798 |
| ENSG00000268362 | | 128.622 | 0.504 | 0.035342844 |
| ENSG00000116670 | MAD2L2 | 2093.926 | 0.504 | 0.000242578 |
| ENSG00000121053 | EPX | 8.915 | 0.504 | 0.030357905 |
| ENSG00000278023 | RDM1 | 131.766 | 0.503 | 0.031235606 |
| ENSG00000259349 | | 11.189 | 0.503 | 0.016860929 |
| ENSG00000198088 | NUP62CL | 124.769 | 0.503 | 0.023928057 |
| ENSG00000114739 | ACVR2B | 934.813 | 0.502 | 0.003889097 |
| ENSG00000184117 | NIPSNAP1 | 4395.434 | 0.502 | 1.97355E-05 |
| ENSG00000140451 | PIF1 | 485.150 | 0.502 | 0.000920138 |
| ENSG00000260735 | | 4.052 | 0.502 | 0.030194653 |
| ENSG00000132359 | RAP1GAP2 | 1088.187 | 0.501 | 0.019754322 |
| ENSG00000215784 | FAM72D | 71.896 | 0.501 | 0.004730388 |
| ENSG00000204248 | COL11A2 | 97.601 | 0.501 | 0.041045779 |
| ENSG00000162062 | TEDC2 | 507.885 | 0.501 | 0.000456342 |
| ENSG00000165490 | DDIAS | 551.178 | 0.500 | 0.000511903 |
| ENSG00000188070 | C11orf95 | 684.918 | 0.500 | 4.06908E-05 |
| ENSG00000141391 | PRELID3A | 119.050 | 0.500 | 0.044416784 |
| ENSG00000228727 | SAPCD1 | 34.784 | 0.500 | 0.021070356 |
| ENSG00000246174 | KCTD21-AS1 | 107.498 | 0.500 | 0.005940064 |
| ENSG00000112742 | TTK | 1011.802 | 0.499 | 0.000616416 |
| ENSG00000272994 | C2orf49-DT | 109.004 | 0.499 | 6.69965E-05 |
| ENSG00000136982 | DSCC1 | 513.178 | 0.498 | 0.000906664 |
| ENSG00000153574 | RPIA | 1796.768 | 0.498 | 5.60281E-06 |
| ENSG00000106462 | EZH2 | 1538.231 | 0.498 | 0.000396085 |
| ENSG00000126970 | ZC4H2 | 454.825 | 0.498 | 0.042174007 |
| ENSG00000267508 | ZNF285 | 169.330 | 0.498 | 0.028016899 |
| ENSG00000113248 | PCDHB15 | 149.299 | 0.497 | 0.049798651 |
| ENSG00000112149 | CD83 | 886.494 | 0.496 | 0.017748446 |
| ENSG00000224738 | | 35.088 | 0.496 | 0.000567707 |
| ENSG00000234492 | RPL34-DT | 5.530 | 0.496 | 0.025638599 |
| ENSG00000187566 | NHLRC1 | 109.348 | 0.495 | 0.007803237 |
| ENSG00000224799 | | 4.514 | 0.494 | 0.01115206 |
| ENSG00000115163 | CENPA | 574.305 | 0.494 | 0.003295543 |
| ENSG00000147036 | LANCL3 | 145.241 | 0.493 | 0.042626005 |
| ENSG00000255458 | | 21.057 | 0.493 | 0.002991571 |
| ENSG00000165895 | ARHGAP42 | 399.201 | 0.493 | 0.010826073 |

| | | | | |
|---|---|---|---|---|
| ENSG00000240445 | FOXO3B | 32.593 | 0.492 | 0.002000854 |
| ENSG00000165028 | NIPSNAP3B | 54.310 | 0.492 | 0.001577561 |
| ENSG00000225411 | | 18.957 | 0.492 | 0.032992198 |
| ENSG00000230330 | HMGN2P3 | 28.550 | 0.492 | 0.014209324 |
| ENSG00000046889 | PREX2 | 210.184 | 0.491 | 0.040140912 |
| ENSG00000104081 | BMF | 1316.966 | 0.491 | 0.004952109 |
| ENSG00000228624 | HDAC2-AS2 | 55.280 | 0.490 | 0.006389919 |
| ENSG00000159239 | | 390.141 | 0.489 | 0.008089779 |
| ENSG00000112312 | GMNN | 1986.413 | 0.489 | 0.000244876 |
| ENSG00000143578 | CREB3L4 | 1090.225 | 0.489 | 0.000937292 |
| ENSG00000091651 | ORC6 | 785.882 | 0.488 | 0.000550196 |
| ENSG00000137942 | FNBP1L | 3079.648 | 0.488 | 0.000797536 |
| ENSG00000148143 | ZNF462 | 776.140 | 0.488 | 0.007531244 |
| ENSG00000166845 | C18orf54 | 387.989 | 0.488 | 0.000212941 |
| ENSG00000233396 | LINC01719 | 31.513 | 0.487 | 0.015077005 |
| ENSG00000185250 | PPIL6 | 78.452 | 0.487 | 0.007677712 |
| ENSG00000186638 | KIF24 | 332.683 | 0.486 | 0.001036344 |
| ENSG00000137364 | TPMT | 1629.724 | 0.486 | 1.89666E-06 |
| ENSG00000254004 | ZNF260 | 1258.111 | 0.486 | 3.13244E-06 |
| ENSG00000138346 | DNA2 | 464.105 | 0.486 | 7.67747E-05 |
| ENSG00000226245 | ZNF32-AS1 | 12.870 | 0.486 | 0.019417546 |
| ENSG00000155085 | AK9 | 276.376 | 0.486 | 0.000718557 |
| ENSG00000276571 | | 20.905 | 0.486 | 0.001455311 |
| ENSG00000231519 | | 4.981 | 0.486 | 0.019810947 |
| ENSG00000145979 | TBC1D7 | 1047.961 | 0.486 | 2.50797E-06 |
| ENSG00000245017 | LINC02453 | 17.899 | 0.486 | 0.010994952 |
| ENSG00000121653 | MAPK8IP1 | 350.127 | 0.485 | 0.029425111 |
| ENSG00000134690 | CDCA8 | 1603.644 | 0.485 | 0.000752409 |
| ENSG00000187840 | EIF4EBP1 | 3835.832 | 0.483 | 0.010776848 |
| ENSG00000147650 | LRP12 | 700.110 | 0.482 | 0.017257108 |
| ENSG00000260852 | FBXL19-AS1 | 133.039 | 0.481 | 0.005211553 |
| ENSG00000147535 | PLPP5 | 1992.352 | 0.481 | 0.001138935 |
| ENSG00000108389 | MTMR4 | 2773.429 | 0.481 | 9.14206E-06 |
| ENSG00000164985 | PSIP1 | 2726.783 | 0.481 | 0.002825892 |
| ENSG00000181588 | MEX3D | 1752.675 | 0.480 | 6.14151E-06 |
| ENSG00000258472 | | 138.900 | 0.480 | 0.009832054 |
| ENSG00000134222 | PSRC1 | 663.962 | 0.480 | 0.00072571 |
| ENSG00000169750 | RAC3 | 758.006 | 0.479 | 0.027904144 |
| ENSG00000132155 | RAF1 | 6575.570 | 0.479 | 5.60281E-06 |
| ENSG00000005801 | ZNF195 | 1226.223 | 0.479 | 3.43144E-05 |
| ENSG00000071073 | MGAT4A | 1908.756 | 0.479 | 0.017985433 |
| ENSG00000158966 | CACHD1 | 421.386 | 0.479 | 0.041346535 |
| ENSG00000165304 | MELK | 1498.114 | 0.479 | 0.006650041 |
| ENSG00000099284 | MACROH2A2 | 2291.913 | 0.479 | 0.005691556 |
| ENSG00000133026 | MYH10 | 3353.994 | 0.478 | 0.001911788 |
| ENSG00000071564 | TCF3 | 5574.488 | 0.478 | 1.59329E-08 |
| ENSG00000125046 | SSUH2 | 21.133 | 0.477 | 0.04103711 |
| ENSG00000147166 | ITGB1BP2 | 68.167 | 0.477 | 0.014724678 |
| ENSG00000233834 | | 54.154 | 0.477 | 0.048879164 |
| ENSG00000104147 | OIP5 | 216.693 | 0.477 | 0.00132772 |
| ENSG00000198844 | ARHGEF15 | 415.832 | 0.477 | 0.010067738 |
| ENSG00000196172 | ZNF681 | 422.589 | 0.477 | 0.027892327 |
| ENSG00000234241 | | 3.770 | 0.476 | 0.033979311 |
| ENSG00000112984 | KIF20A | 1925.816 | 0.476 | 0.001924121 |
| ENSG00000156876 | SASS6 | 496.628 | 0.476 | 9.01764E-06 |
| ENSG00000174705 | SH3PXD2B | 2545.821 | 0.476 | 0.00174025 |
| ENSG00000108468 | CBX1 | 4001.756 | 0.476 | 1.51103E-08 |
| ENSG00000071054 | MAP4K4 | 6509.811 | 0.476 | 0.001001166 |
| ENSG00000033327 | GAB2 | 1017.108 | 0.476 | 0.007654967 |

| | | | | |
|---|---|---|---|---|
| ENSG00000120963 | ZNF706 | 5171.671 | 0.476 | 0.001899557 |
| ENSG00000175063 | UBE2C | 4188.382 | 0.475 | 0.001880911 |
| ENSG00000119888 | EPCAM | 7304.963 | 0.474 | 0.035582554 |
| ENSG00000146670 | CDCA5 | 2082.252 | 0.474 | 0.003237632 |
| ENSG00000077097 | TOP2B | 12830.143 | 0.474 | 0.002011832 |
| ENSG00000049759 | NEDD4L | 3844.716 | 0.473 | 0.004956352 |
| ENSG00000112186 | CAP2 | 426.509 | 0.473 | 0.043204227 |
| ENSG00000141068 | KSR1 | 1095.553 | 0.473 | 0.010660429 |
| ENSG00000132967 | HMGB1P5 | 686.726 | 0.473 | 3.17031E-05 |
| ENSG00000112308 | C6orf62 | 9459.564 | 0.472 | 1.94992E-08 |
| ENSG00000180953 | ST20 | 137.004 | 0.472 | 0.010227588 |
| ENSG00000170456 | DENND5B | 311.580 | 0.471 | 0.046479925 |
| ENSG00000124181 | PLCG1 | 4063.473 | 0.471 | 2.31474E-07 |
| ENSG00000189339 | SLC35E2B | 2014.776 | 0.471 | 0.001377057 |
| ENSG00000182134 | TDRKH | 880.302 | 0.471 | 0.006669946 |
| ENSG00000146733 | PSPH | 873.333 | 0.471 | 0.001850381 |
| ENSG00000158321 | AUTS2 | 1641.050 | 0.471 | 0.042028123 |
| ENSG00000158373 | H2BC5 | 1038.272 | 0.470 | 0.034253759 |
| ENSG00000213080 | | 46.583 | 0.469 | 0.005541141 |
| ENSG00000241015 | TPM3P9 | 488.540 | 0.469 | 0.010229998 |
| ENSG00000261716 | H2BC20P | 608.639 | 0.469 | 0.011658528 |
| ENSG00000230793 | SMARCE1P5 | 4.174 | 0.469 | 0.00318753 |
| ENSG00000183889 | | 20.859 | 0.468 | 0.012805495 |
| ENSG00000185158 | LRRC37B | 369.508 | 0.468 | 7.01397E-06 |
| ENSG00000279443 | | 42.428 | 0.467 | 0.046282432 |
| ENSG00000196296 | ATP2A1 | 106.716 | 0.467 | 0.013924712 |
| ENSG00000164104 | HMGB2 | 7644.073 | 0.466 | 0.00167745 |
| ENSG00000167807 | | 17.974 | 0.465 | 0.009566359 |
| ENSG00000268635 | | 4.051 | 0.465 | 0.045844753 |
| ENSG00000173401 | GLIPR1L1 | 11.105 | 0.464 | 0.01633784 |
| ENSG00000129197 | RPAIN | 1284.624 | 0.464 | 2.54674E-06 |
| ENSG00000144554 | FANCD2 | 1515.104 | 0.463 | 0.000110205 |
| ENSG00000253047 | | 9.290 | 0.463 | 0.044275973 |
| ENSG00000232303 | DFFBP1 | 5.766 | 0.463 | 0.024946564 |
| ENSG00000160336 | ZNF761 | 1305.873 | 0.463 | 0.016196776 |
| ENSG00000076604 | TRAF4 | 8603.157 | 0.462 | 0.000974875 |
| ENSG00000227598 | | 6.197 | 0.462 | 0.023503675 |
| ENSG00000035141 | FAM136A | 2813.322 | 0.462 | 1.15872E-07 |
| ENSG00000088826 | SMOX | 822.275 | 0.462 | 0.031893217 |
| ENSG00000278916 | CEP83-DT | 25.354 | 0.462 | 0.0044935 |
| ENSG00000149554 | CHEK1 | 1174.914 | 0.462 | 0.000180231 |
| ENSG00000233225 | SSBL3P1 | 21.605 | 0.462 | 0.002070778 |
| ENSG00000128000 | ZNF780B | 621.417 | 0.462 | 0.000730614 |
| ENSG00000261019 | | 6.824 | 0.461 | 0.034611744 |
| ENSG00000260949 | | 9.301 | 0.461 | 0.03398285 |
| ENSG00000272345 | | 73.445 | 0.461 | 0.000707039 |
| ENSG00000142632 | ARHGEF19 | 2317.930 | 0.461 | 0.0083842 |
| ENSG00000261762 | | 85.799 | 0.460 | 0.006228643 |
| ENSG00000259736 | CRTC3-AS1 | 14.183 | 0.460 | 0.007174418 |
| ENSG00000229431 | MED8-AS1 | 28.630 | 0.460 | 0.006730571 |
| ENSG00000250731 | TPM3P6 | 41.633 | 0.459 | 0.047565853 |
| ENSG00000161405 | IKZF3 | 1465.548 | 0.458 | 0.038334312 |
| ENSG00000122779 | TRIM24 | 3892.497 | 0.458 | 0.01465285 |
| ENSG00000174721 | FGFBP3 | 127.952 | 0.458 | 0.019989659 |
| ENSG00000165209 | STRBP | 1379.717 | 0.458 | 1.84341E-05 |
| ENSG00000227973 | PIN4P1 | 6.307 | 0.457 | 0.013778794 |
| ENSG00000164542 | KIAA0895 | 313.279 | 0.457 | 0.035950757 |
| ENSG00000123485 | HJURP | 1190.901 | 0.457 | 0.002011832 |
| ENSG00000047662 | FAM184B | 17.656 | 0.457 | 0.008370735 |

| | | | | |
|---|---|---|---|---|
| ENSG00000264324 | | 4.362 | 0.457 | 0.033774796 |
| ENSG00000182481 | KPNA2 | 8294.514 | 0.457 | 0.000554345 |
| ENSG00000177868 | SVBP | 881.282 | 0.457 | 1.55478E-05 |
| ENSG00000156970 | BUB1B | 1184.230 | 0.457 | 0.002053125 |
| ENSG00000116127 | ALMS1 | 1368.936 | 0.456 | 1.34124E-05 |
| ENSG00000154920 | EME1 | 298.626 | 0.456 | 0.002991571 |
| ENSG00000188227 | ZNF793 | 451.172 | 0.455 | 0.033582875 |
| ENSG00000152127 | MGAT5 | 2889.194 | 0.455 | 0.000330554 |
| ENSG00000263786 | | 16.901 | 0.455 | 0.024768721 |
| ENSG00000151846 | PABPC3 | 57.176 | 0.455 | 0.039454308 |
| ENSG00000262304 | | 7.706 | 0.455 | 0.046496336 |
| ENSG00000229852 | | 98.239 | 0.454 | 0.025704916 |
| ENSG00000272356 | | 72.063 | 0.453 | 0.017953414 |
| ENSG00000131152 | | 4.420 | 0.453 | 0.041199065 |
| ENSG00000280385 | | 137.775 | 0.453 | 0.001675797 |
| ENSG00000140009 | ESR2 | 67.011 | 0.453 | 0.032473188 |
| ENSG00000235413 | KRT18P63 | 6.426 | 0.452 | 0.033322085 |
| ENSG00000271855 | | 33.693 | 0.452 | 0.006885493 |
| ENSG00000232485 | RPL37A-DT | 13.803 | 0.452 | 0.016894477 |
| ENSG00000071539 | TRIP13 | 1583.009 | 0.452 | 0.01466898 |
| ENSG00000070756 | PABPC1 | 155534.767 | 0.452 | 0.022055554 |
| ENSG00000025156 | HSF2 | 618.941 | 0.451 | 8.11461E-07 |
| ENSG00000128791 | TWSG1 | 4185.859 | 0.451 | 0.004397835 |
| ENSG00000115318 | LOXL3 | 286.433 | 0.450 | 0.01376926 |
| ENSG00000255517 | | 178.008 | 0.450 | 0.004398929 |
| ENSG00000272205 | POU2F1-DT | 26.322 | 0.449 | 0.047468694 |
| ENSG00000267248 | | 53.038 | 0.449 | 0.012704779 |
| ENSG00000125247 | TMTC4 | 1326.750 | 0.448 | 0.001111629 |
| ENSG00000134324 | LPIN1 | 1013.085 | 0.448 | 0.006325 |
| ENSG00000110925 | CSRNP2 | 1767.738 | 0.447 | 9.90445E-07 |
| ENSG00000136490 | LIMD2 | 1948.955 | 0.447 | 0.024798997 |
| ENSG00000170085 | SIMC1 | 368.665 | 0.447 | 0.048459918 |
| ENSG00000176927 | EFCAB5 | 19.238 | 0.447 | 0.043952792 |
| ENSG00000131747 | TOP2A | 8607.168 | 0.447 | 0.006066924 |
| ENSG00000113368 | LMNB1 | 3447.218 | 0.447 | 0.001401622 |
| ENSG00000182568 | SATB1 | 1963.657 | 0.447 | 0.024504048 |
| ENSG00000232742 | RHOQP2 | 7.172 | 0.447 | 0.013533816 |
| ENSG00000255435 | | 16.412 | 0.446 | 0.025224566 |
| ENSG00000229348 | HYI-AS1 | 24.539 | 0.446 | 0.023404858 |
| ENSG00000146410 | MTFR2 | 343.318 | 0.446 | 0.001523372 |
| ENSG00000165113 | GKAP1 | 177.113 | 0.445 | 0.008178519 |
| ENSG00000079462 | PAFAH1B3 | 3592.559 | 0.445 | 0.003570015 |
| ENSG00000165914 | TTC7B | 1458.405 | 0.445 | 0.002067733 |
| ENSG00000095203 | EPB41L4B | 1012.262 | 0.444 | 0.048400591 |
| ENSG00000155330 | C16orf87 | 652.116 | 0.444 | 0.000536985 |
| ENSG00000118965 | WDR35 | 656.699 | 0.444 | 2.00756E-05 |
| ENSG00000166263 | STXBP4 | 421.882 | 0.444 | 8.42697E-05 |
| ENSG00000132819 | RBM38 | 2134.625 | 0.443 | 0.002761034 |
| ENSG00000267500 | ZNF887P | 17.149 | 0.443 | 0.014004651 |
| ENSG00000255568 | BRWD1-AS2 | 35.803 | 0.442 | 0.02369537 |
| ENSG00000089685 | BIRC5 | 3391.763 | 0.442 | 0.005598288 |
| ENSG00000273987 | | 16.632 | 0.442 | 0.035893849 |
| ENSG00000169989 | TIGD4 | 16.706 | 0.442 | 0.02262168 |
| ENSG00000128944 | KNSTRN | 1087.372 | 0.442 | 9.7035E-05 |
| ENSG00000124383 | MPHOSPH10 | 2707.447 | 0.441 | 4.15347E-08 |
| ENSG00000072571 | HMMR | 994.147 | 0.441 | 0.007215818 |
| ENSG00000109576 | AADAT | 495.183 | 0.441 | 0.030642667 |
| ENSG00000163159 | VPS72 | 3085.538 | 0.441 | 1.58125E-06 |
| ENSG00000268575 | | 115.362 | 0.441 | 0.008987478 |

| | | | | |
|---|---|---|---|---|
| ENSG00000196912 | ANKRD36B | 49.967 | 0.441 | 0.042144777 |
| ENSG00000126787 | DLGAP5 | 1294.789 | 0.439 | 0.009191976 |
| ENSG00000090889 | KIF4A | 1550.673 | 0.439 | 0.004602264 |
| ENSG00000249786 | EAF1-AS1 | 7.816 | 0.439 | 0.035894638 |
| ENSG00000065882 | TBC1D1 | 4510.427 | 0.438 | 0.002007772 |
| ENSG00000255389 | | 34.064 | 0.438 | 0.026000765 |
| ENSG00000123975 | CKS2 | 2181.716 | 0.438 | 0.002005333 |
| ENSG00000181104 | F2R | 1412.691 | 0.437 | 0.043255739 |
| ENSG00000226833 | | 19.111 | 0.437 | 0.022346319 |
| ENSG00000070669 | ASNS | 1898.318 | 0.437 | 0.006873614 |
| ENSG00000108960 | MMD | 997.163 | 0.437 | 0.0396563 |
| ENSG00000223891 | OSER1-DT | 535.038 | 0.437 | 0.013449747 |
| ENSG00000148950 | IMMP1L | 456.908 | 0.436 | 7.91271E-07 |
| ENSG00000161692 | DBF4B | 633.983 | 0.436 | 3.09955E-05 |
| ENSG00000244607 | CCDC13 | 38.496 | 0.436 | 0.037458318 |
| ENSG00000180035 | ZNF48 | 770.891 | 0.435 | 0.000430978 |
| ENSG00000122483 | CCDC18 | 373.289 | 0.435 | 0.000486584 |
| ENSG00000256061 | DNAAF4 | 46.603 | 0.434 | 0.004656809 |
| ENSG00000235554 | SRSF6P2 | 6.444 | 0.434 | 0.013003483 |
| ENSG00000185774 | KCNIP4 | 53.025 | 0.434 | 0.002286871 |
| ENSG00000171365 | CLCN5 | 621.283 | 0.434 | 0.002059617 |
| ENSG00000272501 | | 70.017 | 0.433 | 0.009276268 |
| ENSG00000102172 | SMS | 5061.466 | 0.433 | 0.000939153 |
| ENSG00000272602 | ZNF595 | 289.632 | 0.432 | 0.008489274 |
| ENSG00000117155 | SSX2IP | 1035.057 | 0.431 | 0.000229671 |
| ENSG00000243819 | RN7SL832P | 28.153 | 0.431 | 0.013975102 |
| ENSG00000137337 | MDC1 | 1869.016 | 0.431 | 1.65131E-05 |
| ENSG00000231066 | NPM1P9 | 9.611 | 0.431 | 0.017125074 |
| ENSG00000197332 | | 27.023 | 0.431 | 0.019907235 |
| ENSG00000128536 | CDHR3 | 64.732 | 0.431 | 0.017117782 |
| ENSG00000187838 | PLSCR3 | 68.436 | 0.431 | 0.001955126 |
| ENSG00000230565 | ZNF32-AS2 | 33.849 | 0.431 | 0.016204497 |
| ENSG00000276384 | | 8.993 | 0.430 | 0.049466607 |
| ENSG00000141510 | TP53 | 3836.074 | 0.430 | 0.005363556 |
| ENSG00000122550 | KLHL7 | 1395.427 | 0.430 | 1.27529E-09 |
| ENSG00000137812 | KNL1 | 933.593 | 0.430 | 0.00537622 |
| ENSG00000269743 | SLC25A53 | 125.466 | 0.429 | 2.05058E-06 |
| ENSG00000013275 | PSMC4 | 6279.902 | 0.429 | 7.39748E-06 |
| ENSG00000065911 | MTHFD2 | 3335.989 | 0.429 | 0.008658662 |
| ENSG00000111665 | CDCA3 | 1319.112 | 0.429 | 0.003648814 |
| ENSG00000149646 | CNBD2 | 13.379 | 0.429 | 0.027539316 |
| ENSG00000164494 | PDSS2 | 857.864 | 0.428 | 6.3548E-05 |
| ENSG00000232611 | | 78.536 | 0.428 | 0.024882953 |
| ENSG00000243943 | ZNF512 | 1158.345 | 0.427 | 6.95087E-05 |
| ENSG00000204147 | ASAH2B | 172.556 | 0.426 | 5.72661E-05 |
| ENSG00000136237 | RAPGEF5 | 659.256 | 0.426 | 0.036667505 |
| ENSG00000264575 | LINC00526 | 70.529 | 0.426 | 0.036082859 |
| ENSG00000117399 | CDC20 | 4160.660 | 0.426 | 0.0159609 |
| ENSG00000260077 | | 126.474 | 0.425 | 0.01537969 |
| ENSG00000264859 | DSG2-AS1 | 22.891 | 0.425 | 0.049088973 |
| ENSG00000100749 | VRK1 | 1078.846 | 0.425 | 7.44241E-05 |
| ENSG00000232677 | LINC00665 | 1667.425 | 0.424 | 0.040677038 |
| ENSG00000280968 | | 10.228 | 0.424 | 0.038371061 |
| ENSG00000280798 | LINC00294 | 500.653 | 0.424 | 0.005688973 |
| ENSG00000029993 | HMGB3 | 5074.044 | 0.423 | 0.001695625 |
| ENSG00000174327 | SLC16A13 | 412.422 | 0.423 | 0.004835609 |
| ENSG00000008086 | CDKL5 | 1005.501 | 0.423 | 0.010955008 |
| ENSG00000279048 | | 34.223 | 0.422 | 0.025082436 |
| ENSG00000093009 | CDC45 | 1076.203 | 0.422 | 0.006138921 |

| | | | | |
|---|---|---|---|---|
| ENSG00000151575 | TEX9 | 174.852 | 0.422 | 0.008319533 |
| ENSG00000111206 | FOXM1 | 2998.758 | 0.422 | 0.019570841 |
| ENSG00000137804 | NUSAP1 | 2614.208 | 0.422 | 0.00547118 |
| ENSG00000005156 | LIG3 | 1761.627 | 0.422 | 2.81683E-05 |
| ENSG00000214826 | DDX12P | 230.125 | 0.421 | 0.008740609 |
| ENSG00000206573 | THUMPD3-AS1 | 842.863 | 0.421 | 0.003980974 |
| ENSG00000149136 | SSRP1 | 11248.709 | 0.420 | 6.11323E-07 |
| ENSG00000161533 | ACOX1 | 8585.378 | 0.420 | 0.035122996 |
| ENSG00000251161 | | 48.016 | 0.420 | 0.024190893 |
| ENSG00000163026 | WDCP | 584.243 | 0.419 | 6.93142E-07 |
| ENSG00000141570 | CBX8 | 634.549 | 0.418 | 0.000571433 |
| ENSG00000205208 | C4orf46 | 591.888 | 0.418 | 6.66146E-05 |
| ENSG00000178295 | GEN1 | 1010.233 | 0.418 | 0.000147164 |
| ENSG00000130348 | QRSL1 | 892.416 | 0.417 | 0.000195155 |
| ENSG00000171316 | CHD7 | 1855.584 | 0.417 | 0.000777981 |
| ENSG00000135387 | CAPRIN1 | 10097.482 | 0.417 | 1.58001E-05 |
| ENSG00000165480 | SKA3 | 734.516 | 0.417 | 0.002495641 |
| ENSG00000256771 | ZNF253 | 446.562 | 0.417 | 0.013343686 |
| ENSG00000107443 | CCNJ | 499.753 | 0.417 | 0.01069268 |
| ENSG00000095321 | CRAT | 3970.745 | 0.417 | 0.034907576 |
| ENSG00000112029 | FBXO5 | 510.508 | 0.417 | 0.002184484 |
| ENSG00000187678 | SPRY4 | 1640.114 | 0.416 | 0.033967385 |
| ENSG00000157657 | ZNF618 | 863.325 | 0.416 | 0.025645137 |
| ENSG00000142731 | PLK4 | 703.364 | 0.416 | 0.001001166 |
| ENSG00000276259 | | 48.008 | 0.415 | 0.010027575 |
| ENSG00000102984 | ZNF821 | 169.599 | 0.415 | 0.000213167 |
| ENSG00000188352 | FOCAD | 1693.338 | 0.415 | 0.013890257 |
| ENSG00000214485 | RPL7P1 | 80.919 | 0.414 | 0.034611744 |
| ENSG00000266910 | | 40.970 | 0.414 | 0.023992693 |
| ENSG00000281189 | GHET1 | 34.826 | 0.413 | 0.007174418 |
| ENSG00000133247 | KMT5C | 809.417 | 0.413 | 0.001119334 |
| ENSG00000111554 | MDM1 | 537.908 | 0.413 | 0.001986389 |
| ENSG00000164924 | YWHAZ | 59556.603 | 0.413 | 0.00096976 |
| ENSG00000196230 | TUBB | 49506.850 | 0.413 | 3.51836E-05 |
| ENSG00000085999 | RAD54L | 602.074 | 0.412 | 0.005688973 |
| ENSG00000270419 | CAHM | 27.810 | 0.412 | 0.027176046 |
| ENSG00000135451 | TROAP | 1470.960 | 0.412 | 0.008469788 |
| ENSG00000182327 | GLTPD2 | 28.370 | 0.412 | 0.028600424 |
| ENSG00000196584 | XRCC2 | 413.795 | 0.410 | 0.005438097 |
| ENSG00000196526 | AFAP1 | 1566.075 | 0.408 | 0.014001403 |
| ENSG00000148019 | CEP78 | 1068.123 | 0.408 | 3.27172E-05 |
| ENSG00000176714 | CCDC121 | 121.668 | 0.408 | 0.001720636 |
| ENSG00000176890 | TYMS | 2276.403 | 0.408 | 0.022088127 |
| ENSG00000248559 | | 10.571 | 0.408 | 0.00687453 |
| ENSG00000111247 | RAD51AP1 | 660.748 | 0.408 | 0.015227286 |
| ENSG00000156313 | RPGR | 376.808 | 0.407 | 6.69537E-05 |
| ENSG00000085185 | BCORL1 | 906.189 | 0.407 | 0.000386964 |
| ENSG00000198908 | BHLHB9 | 345.601 | 0.406 | 0.001213942 |
| ENSG00000198346 | ZNF813 | 562.862 | 0.406 | 0.028977515 |
| ENSG00000251503 | CENPS-CORT | 38.962 | 0.406 | 0.003117218 |
| ENSG00000163126 | ANKRD23 | 65.831 | 0.405 | 0.017406504 |
| ENSG00000260400 | | 55.214 | 0.405 | 0.03991284 |
| ENSG00000184481 | FOXO4 | 1247.368 | 0.404 | 0.01032959 |
| ENSG00000070950 | RAD18 | 929.775 | 0.404 | 0.000151488 |
| ENSG00000178691 | SUZ12 | 2619.582 | 0.403 | 5.4166E-05 |
| ENSG00000156802 | ATAD2 | 2881.741 | 0.403 | 0.004963605 |
| ENSG00000172167 | MTBP | 394.249 | 0.402 | 0.002630363 |
| ENSG00000087095 | NLK | 1454.235 | 0.402 | 2.81683E-05 |
| ENSG00000213089 | PDCL3P5 | 10.597 | 0.401 | 0.020030616 |

| | | | | |
|---|---|---|---|---|
| ENSG00000127334 | DYRK2 | 2707.569 | 0.401 | 0.003934368 |
| ENSG00000111802 | TDP2 | 2285.268 | 0.401 | 3.86855E-05 |
| ENSG00000077684 | JADE1 | 1201.499 | 0.401 | 0.00813815 |
| ENSG00000110987 | BCL7A | 1407.569 | 0.401 | 0.000529141 |
| ENSG00000236144 | TMEM147-AS1 | 568.930 | 0.401 | 0.01465285 |
| ENSG00000147140 | NONO | 18294.761 | 0.401 | 1.8398E-08 |
| ENSG00000100105 | PATZ1 | 3080.239 | 0.401 | 0.010594751 |
| ENSG00000165115 | KIF27 | 140.095 | 0.400 | 0.001247577 |
| ENSG00000234072 | | 366.396 | 0.400 | 0.001036344 |
| ENSG00000097046 | CDC7 | 727.298 | 0.400 | 0.003871917 |
| ENSG00000203668 | CHML | 843.441 | 0.399 | 0.014618107 |
| ENSG00000115484 | CCT4 | 9143.211 | 0.399 | 0.000162566 |
| ENSG00000166860 | ZBTB39 | 496.918 | 0.399 | 0.00072571 |
| ENSG00000231822 | SMC3P1 | 8.791 | 0.398 | 0.034611744 |
| ENSG00000175455 | CCDC14 | 2414.928 | 0.398 | 0.007052261 |
| ENSG00000141499 | WRAP53 | 833.501 | 0.397 | 0.000281139 |
| ENSG00000181192 | DHTKD1 | 2301.071 | 0.397 | 0.001885495 |
| ENSG00000120802 | TMPO | 6004.214 | 0.397 | 0.000638853 |
| ENSG00000160957 | RECQL4 | 2408.248 | 0.396 | 0.00913827 |
| ENSG00000108651 | UTP6 | 2614.858 | 0.396 | 2.88179E-06 |
| ENSG00000110048 | OSBP | 4076.745 | 0.395 | 1.94274E-06 |
| ENSG00000268798 | | 14.300 | 0.395 | 0.04831862 |
| ENSG00000181544 | FANCB | 116.920 | 0.395 | 0.009367775 |
| ENSG00000111788 | | 277.407 | 0.395 | 0.017125074 |
| ENSG00000197147 | LRRC8B | 1074.532 | 0.395 | 0.001492344 |
| ENSG00000139146 | SINHCAF | 2905.571 | 0.395 | 0.001467745 |
| ENSG00000272008 | | 17.525 | 0.394 | 0.017211126 |
| ENSG00000206567 | | 197.081 | 0.394 | 0.014912615 |
| ENSG00000197119 | SLC25A29 | 1898.116 | 0.394 | 0.013527107 |
| ENSG00000149503 | INCENP | 1571.974 | 0.393 | 0.00211452 |
| ENSG00000106086 | PLEKHA8 | 790.634 | 0.393 | 3.38221E-05 |
| ENSG00000152990 | ADGRA3 | 1426.095 | 0.393 | 0.020530799 |
| ENSG00000159055 | MIS18A | 1060.333 | 0.393 | 0.000317181 |
| ENSG00000260805 | | 87.403 | 0.392 | 0.032381639 |
| ENSG00000074855 | ANO8 | 890.714 | 0.392 | 0.02477578 |
| ENSG00000234945 | GTF3C2-AS1 | 16.615 | 0.392 | 0.041071879 |
| ENSG00000164754 | RAD21 | 10596.832 | 0.391 | 0.000184345 |
| ENSG00000117650 | NEK2 | 1025.697 | 0.391 | 0.011845383 |
| ENSG00000006625 | GGCT | 3743.953 | 0.391 | 0.001240472 |
| ENSG00000141504 | SAT2 | 2001.948 | 0.391 | 0.00215118 |
| ENSG00000136824 | SMC2 | 1621.850 | 0.391 | 0.006308813 |
| ENSG00000137414 | FAM8A1 | 1408.374 | 0.390 | 8.691E-05 |
| ENSG00000144034 | TPRKB | 1222.793 | 0.390 | 1.0325E-06 |
| ENSG00000261526 | | 154.610 | 0.390 | 0.009229572 |
| ENSG00000094975 | SUCO | 2916.002 | 0.390 | 0.001334226 |
| ENSG00000225151 | GOLGA2P7 | 45.906 | 0.389 | 0.028115137 |
| ENSG00000264538 | SUZ12P1 | 547.291 | 0.389 | 0.001048961 |
| ENSG00000163535 | SGO2 | 575.599 | 0.389 | 0.004509267 |
| ENSG00000112787 | FBRSL1 | 3112.131 | 0.388 | 3.69099E-05 |
| ENSG00000168268 | NT5DC2 | 5545.648 | 0.388 | 0.018860521 |
| ENSG00000135596 | MICAL1 | 2683.316 | 0.388 | 0.010876174 |
| ENSG00000143324 | XPR1 | 3524.398 | 0.388 | 0.005115259 |
| ENSG00000117528 | ABCD3 | 4289.425 | 0.387 | 0.021157364 |
| ENSG00000170364 | SETMAR | 939.930 | 0.386 | 0.003038874 |
| ENSG00000165632 | TAF3 | 894.123 | 0.386 | 0.000104745 |
| ENSG00000070501 | POLB | 1626.966 | 0.386 | 0.007187669 |
| ENSG00000175216 | CKAP5 | 5820.631 | 0.386 | 0.000105437 |
| ENSG00000105171 | POP4 | 2074.868 | 0.386 | 0.002030705 |
| ENSG00000198040 | ZNF84 | 1421.466 | 0.385 | 0.001675208 |

| | | | | |
|---|---|---|---|---|
| ENSG00000250539 | KRT8P33 | 40.527 | 0.385 | 0.027500677 |
| ENSG00000131470 | PSMC3IP | 317.857 | 0.385 | 0.000972247 |
| ENSG00000197933 | ZNF823 | 590.505 | 0.384 | 0.048671716 |
| ENSG00000132436 | FIGNL1 | 784.410 | 0.384 | 0.002721203 |
| ENSG00000104290 | FZD3 | 757.187 | 0.384 | 0.03284533 |
| ENSG00000015133 | CCDC88C | 1863.260 | 0.383 | 0.00534063 |
| ENSG00000108828 | VAT1 | 11373.069 | 0.383 | 0.000163397 |
| ENSG00000147050 | KDM6A | 1455.543 | 0.383 | 0.011658528 |
| ENSG00000278053 | DDX52 | 2249.235 | 0.383 | 3.44172E-06 |
| ENSG00000120533 | ENY2 | 2920.733 | 0.382 | 7.17839E-06 |
| ENSG00000262712 | | 32.909 | 0.382 | 0.026879287 |
| ENSG00000132846 | ZBED3 | 403.956 | 0.381 | 0.034498602 |
| ENSG00000141577 | CEP131 | 1861.208 | 0.381 | 0.003958871 |
| ENSG00000024526 | DEPDC1 | 819.667 | 0.381 | 0.034495023 |
| ENSG00000108559 | NUP88 | 1939.562 | 0.381 | 1.00272E-05 |
| ENSG00000100296 | THOC5 | 1745.624 | 0.380 | 0.000202285 |
| ENSG00000227256 | MIS18A-AS1 | 9.938 | 0.380 | 0.046624908 |
| ENSG00000154832 | CXXC1 | 3701.138 | 0.380 | 0.003724501 |
| ENSG00000227486 | | 52.024 | 0.380 | 0.010151555 |
| ENSG00000129245 | FXR2 | 2124.234 | 0.379 | 6.93142E-07 |
| ENSG00000259781 | HMGB1P6 | 565.510 | 0.379 | 0.001024227 |
| ENSG00000141562 | NARF | 3890.754 | 0.379 | 0.001219831 |
| ENSG00000182518 | FAM104B | 810.166 | 0.378 | 0.000973485 |
| ENSG00000272663 | PPP1R21-DT | 15.241 | 0.377 | 0.044655994 |
| ENSG00000278867 | | 31.296 | 0.377 | 0.018113895 |
| ENSG00000274943 | | 21.107 | 0.376 | 0.049296408 |
| ENSG00000162063 | CCNF | 1639.508 | 0.375 | 0.001008655 |
| ENSG00000171827 | ZNF570 | 334.383 | 0.375 | 0.023866739 |
| ENSG00000102384 | CENPI | 363.543 | 0.375 | 0.018198431 |
| ENSG00000105245 | NUMBL | 1483.812 | 0.374 | 0.010117016 |
| ENSG00000179532 | DNHD1 | 382.915 | 0.374 | 0.035034091 |
| ENSG00000152767 | FARP1 | 3746.549 | 0.374 | 0.024209264 |
| ENSG00000136295 | TTYH3 | 6946.986 | 0.374 | 0.00174441 |
| ENSG00000134461 | ANKRD16 | 473.891 | 0.374 | 0.000640372 |
| ENSG00000109089 | CDR2L | 2643.990 | 0.373 | 0.030180388 |
| ENSG00000111530 | CAND1 | 7024.862 | 0.373 | 0.006667099 |
| ENSG00000111653 | ING4 | 1527.833 | 0.373 | 0.000174452 |
| ENSG00000147799 | ARHGAP39 | 835.699 | 0.372 | 0.000948183 |
| ENSG00000115825 | PRKD3 | 1909.098 | 0.372 | 0.000654689 |
| ENSG00000160325 | CACFD1 | 1956.889 | 0.372 | 0.009462891 |
| ENSG00000273893 | | 49.357 | 0.372 | 0.025702226 |
| ENSG00000108262 | GIT1 | 5459.524 | 0.372 | 5.54096E-05 |
| ENSG00000120158 | RCL1 | 727.325 | 0.372 | 0.001718704 |
| ENSG00000130881 | LRP3 | 1569.962 | 0.372 | 0.039335515 |
| ENSG00000270587 | | 17.271 | 0.370 | 0.009767915 |
| ENSG00000133265 | HSPBP1 | 3474.396 | 0.370 | 0.001389013 |
| ENSG00000166889 | PATL1 | 3496.571 | 0.370 | 0.000182978 |
| ENSG00000230487 | PSMG3-AS1 | 185.715 | 0.370 | 0.033901189 |
| ENSG00000135953 | MFSD9 | 958.441 | 0.370 | 0.002922847 |
| ENSG00000146731 | CCT6A | 11799.079 | 0.369 | 0.000709062 |
| ENSG00000123374 | CDK2 | 3024.544 | 0.368 | 0.002213183 |
| ENSG00000115687 | PASK | 897.696 | 0.368 | 0.003051889 |
| ENSG00000086475 | SEPHS1 | 3131.325 | 0.368 | 2.88639E-05 |
| ENSG00000166508 | MCM7 | 9385.884 | 0.368 | 0.000869578 |
| ENSG00000134369 | NAV1 | 2352.024 | 0.368 | 0.049289464 |
| ENSG00000160392 | C19orf47 | 1231.008 | 0.368 | 0.000629216 |
| ENSG00000163029 | SMC6 | 1913.715 | 0.368 | 0.000215965 |
| ENSG00000184897 | H1-10 | 7099.148 | 0.367 | 0.000714308 |
| ENSG00000153487 | ING1 | 1117.676 | 0.367 | 1.92795E-05 |

| | | | | |
|---|---|---|---|---|
| ENSG00000235078 | | 9.624 | 0.367 | 0.039908366 |
| ENSG00000236565 | HNRNPA3P5 | 12.489 | 0.367 | 0.01050836 |
| ENSG00000259959 | | 171.964 | 0.367 | 0.005098621 |
| ENSG00000236199 | | 76.795 | 0.367 | 0.012843068 |
| ENSG00000239912 | RPL39P36 | 11.845 | 0.366 | 0.009468191 |
| ENSG00000142065 | ZFP14 | 492.812 | 0.366 | 0.007484437 |
| ENSG00000262115 | | 7.884 | 0.366 | 0.047446928 |
| ENSG00000185414 | MRPL30 | 1570.011 | 0.366 | 1.52781E-05 |
| ENSG00000177732 | SOX12 | 1382.045 | 0.365 | 0.026117569 |
| ENSG00000261740 | BOLA2-SMG1P6 | 209.576 | 0.365 | 0.008658662 |
| ENSG00000273329 | | 137.672 | 0.364 | 0.017847113 |
| ENSG00000232995 | RGS5 | 78.576 | 0.364 | 0.035810732 |
| ENSG00000104341 | LAPTM4B | 8031.622 | 0.364 | 0.027460788 |
| ENSG00000101624 | CEP76 | 392.384 | 0.363 | 0.000337054 |
| ENSG00000131116 | ZNF428 | 1618.005 | 0.363 | 0.004712992 |
| ENSG00000143367 | TUFT1 | 3207.018 | 0.363 | 0.027169363 |
| ENSG00000116001 | TIA1 | 4508.520 | 0.363 | 0.001880554 |
| ENSG00000143315 | PIGM | 1128.486 | 0.362 | 0.008268711 |
| ENSG00000127586 | CHTF18 | 1718.076 | 0.362 | 0.010798607 |
| ENSG00000229692 | SOS1-IT1 | 75.881 | 0.361 | 0.012060959 |
| ENSG00000163467 | TSACC | 34.477 | 0.361 | 0.034450394 |
| ENSG00000011258 | MBTD1 | 1264.708 | 0.361 | 0.005744819 |
| ENSG00000215158 | | 102.596 | 0.361 | 0.030357905 |
| ENSG00000105176 | URI1 | 4458.637 | 0.360 | 0.002513718 |
| ENSG00000270039 | | 47.360 | 0.360 | 0.015696763 |
| ENSG00000163918 | RFC4 | 1504.455 | 0.359 | 0.005176067 |
| ENSG00000235363 | SNRPGP10 | 15.538 | 0.359 | 0.019222657 |
| ENSG00000140525 | FANCI | 2621.470 | 0.359 | 0.002012691 |
| ENSG00000124357 | NAGK | 5128.082 | 0.359 | 0.000732286 |
| ENSG00000130559 | CAMSAP1 | 2102.404 | 0.359 | 0.000489502 |
| ENSG00000080608 | PUM3 | 1970.133 | 0.359 | 0.002107113 |
| ENSG00000179833 | SERTAD2 | 2527.038 | 0.359 | 0.002068673 |
| ENSG00000180626 | ZNF594 | 238.262 | 0.359 | 0.011989059 |
| ENSG00000279227 | | 37.289 | 0.359 | 0.015615255 |
| ENSG00000167291 | TBC1D16 | 2070.282 | 0.359 | 0.005851257 |
| ENSG00000241360 | PDXP | 29.890 | 0.358 | 0.047895945 |
| ENSG00000196150 | ZNF250 | 542.466 | 0.358 | 5.05333E-05 |
| ENSG00000148459 | PDSS1 | 379.925 | 0.358 | 0.000779025 |
| ENSG00000165169 | DYNLT3 | 2171.525 | 0.358 | 0.016480898 |
| ENSG00000157456 | CCNB2 | 2057.710 | 0.358 | 0.019236821 |
| ENSG00000251247 | ZNF345 | 230.942 | 0.357 | 0.020015263 |
| ENSG00000186871 | ERCC6L | 379.641 | 0.357 | 0.042317216 |
| ENSG00000162086 | ZNF75A | 693.896 | 0.357 | 0.008529464 |
| ENSG00000142945 | KIF2C | 2334.237 | 0.357 | 0.017552018 |
| ENSG00000047230 | CTPS2 | 1334.197 | 0.356 | 7.11799E-05 |
| ENSG00000133422 | MORC2 | 3098.551 | 0.356 | 2.81683E-05 |
| ENSG00000134453 | RBM17 | 5337.557 | 0.356 | 0.000100408 |
| ENSG00000257390 | | 15.723 | 0.355 | 0.003251456 |
| ENSG00000228315 | GUSBP11 | 69.480 | 0.355 | 0.046582654 |
| ENSG00000234444 | ZNF736 | 561.199 | 0.355 | 0.049880037 |
| ENSG00000247137 | | 73.350 | 0.355 | 0.03284533 |
| ENSG00000237649 | KIFC1 | 2391.764 | 0.355 | 0.023454808 |
| ENSG00000106443 | PHF14 | 2300.339 | 0.355 | 3.96263E-05 |
| ENSG00000189298 | ZKSCAN3 | 317.464 | 0.354 | 0.003017219 |
| ENSG00000105677 | TMEM147 | 5963.840 | 0.354 | 0.00475405 |
| ENSG00000084676 | NCOA1 | 2628.493 | 0.353 | 0.000850926 |
| ENSG00000168496 | FEN1 | 3014.930 | 0.353 | 0.016403345 |
| ENSG00000114346 | ECT2 | 2318.139 | 0.352 | 0.024942814 |
| ENSG00000166851 | PLK1 | 2577.590 | 0.352 | 0.036304758 |

| | | | | |
|---|---|---|---|---|
| ENSG00000183814 | LIN9 | 426.933 | 0.352 | 0.000977735 |
| ENSG00000139880 | CDH24 | 832.325 | 0.352 | 0.03883497 |
| ENSG00000114796 | KLHL24 | 2657.432 | 0.352 | 0.009221114 |
| ENSG00000080986 | NDC80 | 1241.213 | 0.352 | 0.031456188 |
| ENSG00000095002 | MSH2 | 2157.859 | 0.351 | 0.002184484 |
| ENSG00000188493 | C19orf54 | 923.433 | 0.351 | 0.002089549 |
| ENSG00000249115 | HAUS5 | 1544.459 | 0.351 | 0.001225539 |
| ENSG00000168005 | SPINDOC | 1629.945 | 0.350 | 0.000150377 |
| ENSG00000101811 | CSTF2 | 1247.746 | 0.350 | 0.000203126 |
| ENSG00000198134 | PTMAP9 | 24.845 | 0.350 | 0.005682641 |
| ENSG00000105738 | SIPA1L3 | 3169.342 | 0.350 | 0.010413205 |
| ENSG00000168234 | TTC39C | 993.784 | 0.350 | 0.008366701 |
| ENSG00000138439 | FAM117B | 979.601 | 0.350 | 0.004927235 |
| ENSG00000204822 | MRPL53 | 204.241 | 0.350 | 0.000839481 |
| ENSG00000168887 | C2orf68 | 2104.001 | 0.349 | 0.000115112 |
| ENSG00000183765 | CHEK2 | 927.636 | 0.349 | 0.001458554 |
| ENSG00000111875 | ASF1A | 949.642 | 0.349 | 0.00096976 |
| ENSG00000104517 | UBR5 | 5704.714 | 0.348 | 0.002554549 |
| ENSG00000144597 | EAF1 | 1414.384 | 0.348 | 0.000349981 |
| ENSG00000035499 | DEPDC1B | 584.972 | 0.348 | 0.042899903 |
| ENSG00000173638 | SLC19A1 | 1589.343 | 0.348 | 0.019843894 |
| ENSG00000105866 | SP4 | 363.190 | 0.347 | 0.003896236 |
| ENSG00000178921 | PFAS | 1259.756 | 0.347 | 0.001591141 |
| ENSG00000135372 | NAT10 | 3552.019 | 0.347 | 0.000899841 |
| ENSG00000230844 | ZNF674-AS1 | 160.726 | 0.347 | 0.002000854 |
| ENSG00000123213 | NLN | 1345.440 | 0.347 | 0.017699907 |
| ENSG00000255139 | | 30.071 | 0.347 | 0.00561624 |
| ENSG00000184271 | POU6F1 | 249.169 | 0.347 | 0.041978869 |
| ENSG00000157216 | SSBP3 | 2753.676 | 0.347 | 0.005046957 |
| ENSG00000218891 | ZNF579 | 872.333 | 0.347 | 0.033248755 |
| ENSG00000005189 | REXO5 | 322.919 | 0.347 | 0.018483901 |
| ENSG00000104221 | BRF2 | 380.721 | 0.346 | 0.044660439 |
| ENSG00000177855 | CACYBPP2 | 9.007 | 0.346 | 0.035403977 |
| ENSG00000112877 | CEP72 | 375.604 | 0.346 | 0.024742098 |
| ENSG00000156983 | BRPF1 | 1656.281 | 0.346 | 4.42613E-05 |
| ENSG00000161800 | RACGAP1 | 2425.553 | 0.346 | 0.016573531 |
| ENSG00000077152 | UBE2T | 1124.877 | 0.345 | 0.009012894 |
| ENSG00000159873 | CCDC117 | 93.805 | 0.344 | 0.042254001 |
| ENSG00000266490 | | 43.924 | 0.343 | 0.018918692 |
| ENSG00000110104 | CCDC86 | 2289.664 | 0.342 | 0.006755146 |
| ENSG00000146574 | CCZ1B | 428.215 | 0.342 | 0.000974875 |
| ENSG00000077232 | DNAJC10 | 4470.468 | 0.342 | 0.001371755 |
| ENSG00000135622 | SEMA4F | 699.353 | 0.341 | 0.021820186 |
| ENSG00000030066 | NUP160 | 2315.320 | 0.341 | 0.000624896 |
| ENSG00000160208 | RRP1B | 2769.824 | 0.341 | 0.000233913 |
| ENSG00000204371 | EHMT2 | 4758.735 | 0.341 | 7.87116E-06 |
| ENSG00000213551 | DNAJC9 | 1174.711 | 0.340 | 0.000852441 |
| ENSG00000015171 | ZMYND11 | 3714.972 | 0.340 | 0.000629818 |
| ENSG00000203362 | POLH-AS1 | 37.096 | 0.340 | 0.022687627 |
| ENSG00000179195 | ZNF664 | 5904.067 | 0.340 | 0.001001166 |
| ENSG00000228889 | UBAC2-AS1 | 81.527 | 0.340 | 0.045160691 |
| ENSG00000151657 | KIN | 829.652 | 0.340 | 0.000359928 |
| ENSG00000079134 | THOC1 | 1625.275 | 0.339 | 0.00072571 |
| ENSG00000141965 | FEM1A | 158.870 | 0.339 | 0.011648913 |
| ENSG00000188295 | ZNF669 | 197.337 | 0.339 | 0.001321151 |
| ENSG00000110107 | PRPF19 | 8148.024 | 0.339 | 1.31755E-05 |
| ENSG00000033867 | SLC4A7 | 1198.117 | 0.339 | 0.030934306 |
| ENSG00000186017 | ZNF566 | 350.846 | 0.339 | 0.002011832 |
| ENSG00000267152 | | 93.232 | 0.338 | 0.033979311 |

| | | | | |
|---|---|---|---|---|
| ENSG00000180198 | RCC1 | 4589.495 | 0.338 | 0.001920222 |
| ENSG00000170946 | DNAJC24 | 298.200 | 0.338 | 0.001743801 |
| ENSG00000173473 | SMARCC1 | 7880.327 | 0.338 | 0.001266945 |
| ENSG00000244627 | TPTEP2 | 114.964 | 0.337 | 0.029329848 |
| ENSG00000143977 | SNRPG | 2890.968 | 0.337 | 0.000176486 |
| ENSG00000279332 | | 77.750 | 0.337 | 0.049014956 |
| ENSG00000004975 | DVL2 | 1814.840 | 0.337 | 0.000172515 |
| ENSG00000158169 | FANCC | 670.707 | 0.337 | 0.010362091 |
| ENSG00000079616 | KIF22 | 4541.715 | 0.336 | 0.005655432 |
| ENSG00000168137 | SETD5 | 5731.316 | 0.336 | 0.000851709 |
| ENSG00000137038 | DMAC1 | 1884.602 | 0.336 | 0.006896338 |
| ENSG00000109189 | USP46 | 1329.845 | 0.335 | 0.010977935 |
| ENSG00000115317 | HTRA2 | 1260.810 | 0.335 | 7.06586E-05 |
| ENSG00000259994 | | 94.442 | 0.335 | 0.014308309 |
| ENSG00000198464 | ZNF480 | 1171.718 | 0.335 | 0.00509627 |
| ENSG00000214223 | HNRNPA1P10 | 38.388 | 0.335 | 0.037517349 |
| ENSG00000094804 | CDC6 | 1892.338 | 0.335 | 0.025905408 |
| ENSG00000108406 | DHX40 | 2543.356 | 0.334 | 0.000513065 |
| ENSG00000243364 | EFNA4 | 987.297 | 0.334 | 0.009583872 |
| ENSG00000117569 | PTBP2 | 730.402 | 0.334 | 0.002922847 |
| ENSG00000220875 | H3C9P | 8.142 | 0.334 | 0.034574941 |
| ENSG00000143079 | CTTNBP2NL | 2066.962 | 0.334 | 0.00100685 |
| ENSG00000167645 | YIF1B | 2494.627 | 0.334 | 0.0083842 |
| ENSG00000215790 | SLC35E2A | 88.142 | 0.333 | 0.037598669 |
| ENSG00000085840 | ORC1 | 540.033 | 0.333 | 0.048135216 |
| ENSG00000165138 | ANKS6 | 1403.957 | 0.333 | 0.045489837 |
| ENSG00000235655 | H3P6 | 169.422 | 0.332 | 0.043933307 |
| ENSG00000047579 | DTNBP1 | 1088.540 | 0.332 | 0.001521382 |
| ENSG00000156603 | MED19 | 782.107 | 0.332 | 0.000180352 |
| ENSG00000234028 | EIF2AK3-DT | 113.175 | 0.332 | 0.040469046 |
| ENSG00000167720 | SRR | 257.053 | 0.332 | 0.004139214 |
| ENSG00000172301 | COPRS | 1922.636 | 0.331 | 0.01327472 |
| ENSG00000187741 | FANCA | 1109.154 | 0.331 | 0.023343782 |
| ENSG00000109805 | NCAPG | 1196.962 | 0.331 | 0.049253411 |
| ENSG00000100523 | DDHD1 | 933.357 | 0.330 | 0.031929735 |
| ENSG00000198901 | PRC1 | 2810.880 | 0.330 | 0.031592975 |
| ENSG00000171224 | FAM241B | 608.291 | 0.330 | 0.037467211 |
| ENSG00000277462 | ZNF670 | 181.269 | 0.330 | 0.006407033 |
| ENSG00000116161 | CACYBP | 4356.792 | 0.329 | 0.000759653 |
| ENSG00000196670 | ZFP62 | 1330.553 | 0.329 | 0.006333382 |
| ENSG00000163602 | RYBP | 2844.114 | 0.329 | 0.003070023 |
| ENSG00000170836 | PPM1D | 752.592 | 0.329 | 0.001955659 |
| ENSG00000102302 | FGD1 | 864.586 | 0.329 | 0.043869926 |
| ENSG00000164975 | SNAPC3 | 1258.762 | 0.329 | 0.001197836 |
| ENSG00000136122 | BORA | 396.725 | 0.328 | 0.016184023 |
| ENSG00000100629 | CEP128 | 293.913 | 0.327 | 0.008815723 |
| ENSG00000255717 | SNHG1 | 2778.698 | 0.327 | 0.004837657 |
| ENSG00000266472 | MRPS21 | 4152.551 | 0.327 | 0.029638243 |
| ENSG00000102606 | ARHGEF7 | 3046.205 | 0.326 | 0.00072571 |
| ENSG00000245149 | RNF139-AS1 | 121.296 | 0.326 | 0.020447833 |
| ENSG00000182287 | AP1S2 | 665.978 | 0.326 | 0.041633519 |
| ENSG00000060749 | QSER1 | 2034.493 | 0.326 | 0.012968711 |
| ENSG00000004478 | FKBP4 | 10972.511 | 0.326 | 0.004654603 |
| ENSG00000143379 | SETDB1 | 2168.641 | 0.325 | 0.000526754 |
| ENSG00000152253 | SPC25 | 437.495 | 0.325 | 0.045884903 |
| ENSG00000140743 | CDR2 | 1871.912 | 0.325 | 0.022717502 |
| ENSG00000138035 | PNPT1 | 1928.300 | 0.325 | 0.000335545 |
| ENSG00000214719 | | 73.154 | 0.325 | 0.006696908 |
| ENSG00000049449 | RCN1 | 9684.574 | 0.324 | 0.025738938 |

| | | | | |
|---|---|---|---|---|
| ENSG00000164934 | DCAF13 | 3374.398 | 0.324 | 0.004583561 |
| ENSG00000149639 | SOGA1 | 2783.739 | 0.323 | 0.009931111 |
| ENSG00000148737 | TCF7L2 | 1335.491 | 0.323 | 0.014972697 |
| ENSG00000119729 | RHOQ | 2678.428 | 0.323 | 0.005655432 |
| ENSG00000183161 | FANCF | 859.279 | 0.323 | 0.009668798 |
| ENSG00000123473 | STIL | 825.328 | 0.323 | 0.031891572 |
| ENSG00000108395 | TRIM37 | 1492.124 | 0.323 | 0.001442381 |
| ENSG00000261188 | TFIP11-DT | 177.837 | 0.322 | 0.025905408 |
| ENSG00000261879 | ZNF594-DT | 47.220 | 0.322 | 0.013846123 |
| ENSG00000075218 | GTSE1 | 952.868 | 0.322 | 0.037598669 |
| ENSG00000134297 | PLEKHA8P1 | 209.142 | 0.322 | 0.002495641 |
| ENSG00000143621 | ILF2 | 11368.984 | 0.322 | 0.000341451 |
| ENSG00000134313 | KIDINS220 | 3342.166 | 0.322 | 0.003826484 |
| ENSG00000267249 | | 57.908 | 0.322 | 0.010212424 |
| ENSG00000271851 | | 57.719 | 0.321 | 0.010369294 |
| ENSG00000186812 | ZNF397 | 1225.909 | 0.321 | 0.005984602 |
| ENSG00000145014 | TMEM44 | 1048.464 | 0.321 | 0.017757408 |
| ENSG00000089220 | PEBP1 | 20789.066 | 0.321 | 0.041030424 |
| ENSG00000057608 | GDI2 | 14747.307 | 0.320 | 0.001171384 |
| ENSG00000082458 | DLG3 | 2607.293 | 0.320 | 0.0117034 |
| ENSG00000167523 | SPATA33 | 376.187 | 0.319 | 0.006303775 |
| ENSG00000183530 | PRR14L | 2369.204 | 0.319 | 0.004449272 |
| ENSG00000078246 | TULP3 | 1173.778 | 0.319 | 0.001458435 |
| ENSG00000166881 | NEMP1 | 1415.214 | 0.319 | 0.013363445 |
| ENSG00000171103 | TRMT61B | 511.943 | 0.318 | 0.000250074 |
| ENSG00000225507 | | 85.325 | 0.318 | 0.035201594 |
| ENSG00000263766 | KPNB1-DT | 20.484 | 0.318 | 0.039562862 |
| ENSG00000143198 | MGST3 | 4162.826 | 0.318 | 0.02130886 |
| ENSG00000125319 | HROB | 500.961 | 0.317 | 0.022338048 |
| ENSG00000149089 | APIP | 730.467 | 0.317 | 0.004437993 |
| ENSG00000116830 | TTF2 | 1421.096 | 0.317 | 0.004083282 |
| ENSG00000156787 | TBC1D31 | 515.561 | 0.316 | 0.002044279 |
| ENSG00000173207 | CKS1B | 1884.666 | 0.316 | 0.026995467 |
| ENSG00000127616 | SMARCA4 | 10097.367 | 0.316 | 0.000203126 |
| ENSG00000215041 | NEURL4 | 1073.166 | 0.316 | 0.000326248 |
| ENSG00000279413 | | 99.914 | 0.316 | 0.010027575 |
| ENSG00000160949 | TONSL | 1951.992 | 0.315 | 0.013974928 |
| ENSG00000160299 | PCNT | 2219.631 | 0.315 | 0.000571588 |
| ENSG00000021574 | SPAST | 1301.627 | 0.315 | 6.50933E-05 |
| ENSG00000198826 | ARHGAP11A | 1295.887 | 0.315 | 0.036964939 |
| ENSG00000264350 | SNRPGP2 | 139.205 | 0.315 | 0.004302458 |
| ENSG00000176236 | RPP38-DT | 14.025 | 0.314 | 0.039464078 |
| ENSG00000109083 | IFT20 | 1476.487 | 0.313 | 0.005032782 |
| ENSG00000198563 | DDX39B | 4321.335 | 0.313 | 0.014779237 |
| ENSG00000120278 | PLEKHG1 | 1210.739 | 0.313 | 0.047692546 |
| ENSG00000147669 | POLR2K | 3012.770 | 0.313 | 0.016671088 |
| ENSG00000085788 | DDHD2 | 1616.829 | 0.313 | 0.032749089 |
| ENSG00000198551 | ZNF627 | 692.003 | 0.312 | 0.004567065 |
| ENSG00000183763 | TRAIP | 540.499 | 0.312 | 0.017552018 |
| ENSG00000221829 | FANCG | 1471.284 | 0.312 | 0.009693918 |
| ENSG00000161036 | LRWD1 | 1385.728 | 0.312 | 0.000339097 |
| ENSG00000120539 | MASTL | 1100.699 | 0.311 | 0.004933013 |
| ENSG00000122218 | COPA | 13174.139 | 0.311 | 0.00526375 |
| ENSG00000143190 | POU2F1 | 1667.234 | 0.310 | 0.001764446 |
| ENSG00000143256 | PFDN2 | 4398.710 | 0.310 | 0.043679975 |
| ENSG00000178966 | RMI1 | 711.599 | 0.310 | 0.02525939 |
| ENSG00000104889 | RNASEH2A | 2197.069 | 0.310 | 0.011036461 |
| ENSG00000134086 | VHL | 2727.903 | 0.309 | 0.011002777 |
| ENSG00000143442 | POGZ | 4069.177 | 0.307 | 0.008592302 |

| | | | | |
|---|---|---|---|---|
| ENSG00000123908 | AGO2 | 2765.731 | 0.307 | 0.010515911 |
| ENSG00000065183 | WDR3 | 2105.170 | 0.307 | 0.004139403 |
| ENSG00000136261 | BZW2 | 5697.315 | 0.307 | 0.006218918 |
| ENSG00000103494 | RPGRIP1L | 424.778 | 0.307 | 0.00561624 |
| ENSG00000166526 | ZNF3 | 1664.446 | 0.307 | 0.001224499 |
| ENSG00000198554 | WDHD1 | 919.826 | 0.307 | 0.033948006 |
| ENSG00000121289 | CEP89 | 1941.654 | 0.307 | 0.00064489 |
| ENSG00000162607 | USP1 | 2488.960 | 0.306 | 0.022535 |
| ENSG00000137807 | KIF23 | 1624.535 | 0.305 | 0.048044278 |
| ENSG00000250462 | LRRC37BP1 | 347.343 | 0.305 | 0.002576696 |
| ENSG00000105223 | PLD3 | 13784.799 | 0.305 | 0.026795194 |
| ENSG00000120526 | NUDCD1 | 1384.842 | 0.305 | 0.007918241 |
| ENSG00000115514 | TXNDC9 | 1415.608 | 0.304 | 0.002192765 |
| ENSG00000135637 | CCDC142 | 372.137 | 0.304 | 0.003581774 |
| ENSG00000158850 | B4GALT3 | 4287.977 | 0.303 | 0.031863209 |
| ENSG00000232748 | | 29.182 | 0.303 | 0.010216647 |
| ENSG00000151092 | NGLY1 | 1801.708 | 0.303 | 0.005115259 |
| ENSG00000137073 | UBAP2 | 2280.062 | 0.303 | 0.001029677 |
| ENSG00000087338 | GMCL1 | 1551.649 | 0.301 | 0.000746567 |
| ENSG00000146535 | GNA12 | 3742.705 | 0.301 | 0.044835125 |
| ENSG00000101868 | POLA1 | 1146.652 | 0.301 | 0.016078075 |
| ENSG00000152455 | SUV39H2 | 636.655 | 0.300 | 0.003252558 |
| ENSG00000138160 | KIF11 | 2017.599 | 0.300 | 0.04986361 |
| ENSG00000102241 | HTATSF1 | 4906.600 | 0.300 | 0.00049048 |
| ENSG00000166788 | SAAL1 | 697.299 | 0.300 | 0.002825892 |
| ENSG00000168439 | STIP1 | 10118.822 | 0.300 | 0.000764086 |
| ENSG00000152240 | HAUS1 | 1063.475 | 0.300 | 0.014982239 |
| ENSG00000162639 | HENMT1 | 969.787 | 0.299 | 0.035459993 |
| ENSG00000143401 | ANP32E | 4352.085 | 0.299 | 0.020638463 |
| ENSG00000127884 | ECHS1 | 9274.114 | 0.299 | 0.018198431 |
| ENSG00000109685 | NSD2 | 4794.850 | 0.299 | 0.01617163 |
| ENSG00000167378 | IRGQ | 1603.931 | 0.299 | 0.001775953 |
| ENSG00000132300 | PTCD3 | 3294.991 | 0.299 | 0.000279441 |
| ENSG00000109079 | TNFAIP1 | 4484.358 | 0.299 | 0.000278821 |
| ENSG00000165506 | DNAAF2 | 689.429 | 0.298 | 0.000314552 |
| ENSG00000107937 | GTPBP4 | 3952.050 | 0.298 | 0.001492344 |
| ENSG00000043514 | TRIT1 | 2148.707 | 0.298 | 0.044531912 |
| ENSG00000197024 | ZNF398 | 1239.449 | 0.298 | 0.001438315 |
| ENSG00000130669 | PAK4 | 5285.671 | 0.298 | 0.009668798 |
| ENSG00000134452 | FBH1 | 3238.544 | 0.297 | 0.000475021 |
| ENSG00000272009 | | 76.715 | 0.297 | 0.024213378 |
| ENSG00000134779 | TPGS2 | 3830.919 | 0.297 | 0.001843459 |
| ENSG00000236810 | ELOA-AS1 | 66.594 | 0.297 | 0.017876481 |
| ENSG00000137274 | BPHL | 829.517 | 0.297 | 0.023503675 |
| ENSG00000119787 | ATL2 | 2799.602 | 0.297 | 0.019570841 |
| ENSG00000172175 | MALT1 | 1680.908 | 0.297 | 0.043600672 |
| ENSG00000106723 | SPIN1 | 4036.545 | 0.297 | 0.001453916 |
| ENSG00000133619 | KRBA1 | 839.526 | 0.297 | 0.01406253 |
| ENSG00000172239 | PAIP1 | 2817.031 | 0.297 | 0.001138935 |
| ENSG00000231889 | TRAF3IP2-AS1 | 88.431 | 0.296 | 0.022605204 |
| ENSG00000125482 | TTF1 | 820.555 | 0.296 | 9.47934E-05 |
| ENSG00000228343 | | 149.438 | 0.295 | 0.028054767 |
| ENSG00000023287 | RB1CC1 | 2937.556 | 0.295 | 0.0014635 |
| ENSG00000196741 | LINC01560 | 125.812 | 0.295 | 0.011843009 |
| ENSG00000241127 | YAE1 | 478.129 | 0.295 | 0.000202285 |
| ENSG00000173065 | FAM222B | 2065.415 | 0.294 | 0.005067383 |
| ENSG00000131944 | FAAP24 | 354.220 | 0.294 | 0.008149836 |
| ENSG00000042429 | MED17 | 1347.502 | 0.294 | 0.000284083 |
| ENSG00000144028 | SNRNP200 | 12139.989 | 0.294 | 5.62377E-05 |

| | | | | |
|---|---|---|---|---|
| ENSG00000115421 | PAPOLG | 907.298 | 0.292 | 0.000182548 |
| ENSG00000174799 | CEP135 | 528.288 | 0.292 | 0.011050554 |
| ENSG00000170340 | B3GNT2 | 1453.626 | 0.292 | 0.00985943 |
| ENSG00000115042 | FAHD2A | 746.656 | 0.292 | 0.00172882 |
| ENSG00000052723 | SIKE1 | 1763.982 | 0.291 | 8.08613E-05 |
| ENSG00000115207 | GTF3C2 | 3497.466 | 0.290 | 4.66083E-05 |
| ENSG00000122970 | IFT81 | 634.592 | 0.289 | 0.007192999 |
| ENSG00000122565 | CBX3 | 10599.400 | 0.289 | 4.75335E-05 |
| ENSG00000135632 | SMYD5 | 2126.550 | 0.289 | 0.000263975 |
| ENSG00000167842 | MIS12 | 799.782 | 0.287 | 0.001860313 |
| ENSG00000111605 | CPSF6 | 4499.005 | 0.287 | 0.009020741 |
| ENSG00000085382 | HACE1 | 518.368 | 0.287 | 0.039285358 |
| ENSG00000126249 | PDCD2L | 458.566 | 0.286 | 0.014990353 |
| ENSG00000133703 | KRAS | 2507.132 | 0.286 | 0.009179519 |
| ENSG00000146386 | ABRACL | 2660.652 | 0.286 | 0.026847664 |
| ENSG00000185480 | PARPBP | 479.414 | 0.286 | 0.043091157 |
| ENSG00000107890 | ANKRD26 | 484.390 | 0.286 | 0.014518492 |
| ENSG00000164941 | INTS8 | 2283.231 | 0.286 | 0.001275299 |
| ENSG00000172071 | EIF2AK3 | 1410.790 | 0.286 | 0.025575761 |
| ENSG00000179943 | FIZ1 | 1085.540 | 0.285 | 0.0024295 |
| ENSG00000141456 | PELP1 | 4498.861 | 0.285 | 0.005286587 |
| ENSG00000136518 | ACTL6A | 3469.213 | 0.284 | 0.001576413 |
| ENSG00000105443 | CYTH2 | 5368.279 | 0.282 | 0.003814369 |
| ENSG00000163872 | YEATS2 | 2690.983 | 0.282 | 0.003465073 |
| ENSG00000213281 | NRAS | 3460.530 | 0.282 | 0.010309544 |
| ENSG00000179134 | SAMD4B | 6252.543 | 0.281 | 0.000282878 |
| ENSG00000143033 | MTF2 | 1384.788 | 0.281 | 0.001126803 |
| ENSG00000268471 | MIR4453HG | 154.422 | 0.281 | 0.029425111 |
| ENSG00000134809 | TIMM10 | 1468.841 | 0.281 | 0.013309652 |
| ENSG00000244045 | TMEM199 | 924.005 | 0.281 | 0.000156465 |
| ENSG00000184675 | AMER1 | 415.193 | 0.281 | 0.041199065 |
| ENSG00000196417 | ZNF765 | 588.191 | 0.280 | 0.015656119 |
| ENSG00000152193 | OBI1 | 616.861 | 0.280 | 0.002236562 |
| ENSG00000090565 | RAB11FIP3 | 2099.321 | 0.280 | 0.006324412 |
| ENSG00000169925 | BRD3 | 1452.529 | 0.280 | 0.013248409 |
| ENSG00000131591 | C1orf159 | 1292.555 | 0.280 | 0.040565717 |
| ENSG00000132591 | ERAL1 | 3646.201 | 0.280 | 0.002561263 |
| ENSG00000149091 | DGKZ | 4699.952 | 0.279 | 0.016119944 |
| ENSG00000119397 | CNTRL | 1158.444 | 0.279 | 0.014567797 |
| ENSG00000179051 | RCC2 | 13033.781 | 0.279 | 0.000508037 |
| ENSG00000160214 | RRP1 | 3387.706 | 0.279 | 0.024213378 |
| ENSG00000258890 | CEP95 | 1612.551 | 0.278 | 0.004934205 |
| ENSG00000136197 | C7orf25 | 38.303 | 0.277 | 0.002884727 |
| ENSG00000132478 | UNK | 1675.390 | 0.277 | 0.001581084 |
| ENSG00000241973 | PI4KA | 3563.281 | 0.277 | 0.018224094 |
| ENSG00000231503 | PTMAP4 | 32.216 | 0.276 | 0.01684442 |
| ENSG00000077312 | SNRPA | 4499.675 | 0.275 | 0.003136037 |
| ENSG00000157796 | WDR19 | 756.731 | 0.275 | 0.0119868 |
| ENSG00000280088 | | 254.529 | 0.275 | 0.008172941 |
| ENSG00000103160 | HSDL1 | 1211.994 | 0.275 | 0.002091798 |
| ENSG00000197008 | ZNF138 | 545.147 | 0.275 | 0.027539316 |
| ENSG00000005436 | GCFC2 | 847.757 | 0.275 | 0.000247514 |
| ENSG00000198315 | ZKSCAN8 | 1220.713 | 0.274 | 0.015356468 |
| ENSG00000166902 | MRPL16 | 2331.553 | 0.274 | 0.015966835 |
| ENSG00000184402 | SS18L1 | 1270.679 | 0.274 | 0.025194137 |
| ENSG00000005100 | DHX33 | 1331.653 | 0.273 | 0.006671094 |
| ENSG00000108384 | RAD51C | 863.970 | 0.273 | 0.007602949 |
| ENSG00000131373 | HACL1 | 886.195 | 0.272 | 0.013902301 |
| ENSG00000183309 | ZNF623 | 1602.513 | 0.272 | 0.004730388 |

| | | | | |
|---|---|---|---|---|
| ENSG00000155592 | ZKSCAN2 | 316.994 | 0.272 | 0.031878425 |
| ENSG00000110429 | FBXO3 | 1624.216 | 0.272 | 0.012088176 |
| ENSG00000152102 | FAM168B | 4141.857 | 0.272 | 0.008577592 |
| ENSG00000105202 | FBL | 10267.883 | 0.271 | 0.038051593 |
| ENSG00000204394 | VARS1 | 6618.351 | 0.271 | 0.00651696 |
| ENSG00000167635 | ZNF146 | 5122.183 | 0.271 | 0.005285268 |
| ENSG00000104885 | DOT1L | 2130.312 | 0.271 | 0.009693918 |
| ENSG00000177302 | TOP3A | 1297.139 | 0.271 | 0.0083842 |
| ENSG00000237190 | CDKN2AIPNL | 1041.650 | 0.271 | 0.006916186 |
| ENSG00000120334 | CENPL | 433.481 | 0.271 | 0.027952534 |
| ENSG00000102309 | PIN4 | 1063.572 | 0.271 | 0.011078932 |
| ENSG00000179611 | DGKZP1 | 46.377 | 0.270 | 0.028680813 |
| ENSG00000116668 | SWT1 | 319.638 | 0.270 | 0.005562571 |
| ENSG00000101974 | ATP11C | 1225.184 | 0.270 | 0.019895059 |
| ENSG00000171612 | SLC25A33 | 533.831 | 0.270 | 0.025492726 |
| ENSG00000144231 | POLR2D | 1889.727 | 0.270 | 0.000349981 |
| ENSG00000143951 | WDPCP | 169.535 | 0.269 | 0.006916472 |
| ENSG00000227345 | PARG | 1000.535 | 0.269 | 0.003394402 |
| ENSG00000172409 | CLP1 | 646.235 | 0.269 | 0.003971331 |
| ENSG00000278129 | ZNF8 | 540.873 | 0.269 | 0.015713178 |
| ENSG00000170037 | CNTROB | 2086.023 | 0.269 | 0.001396095 |
| ENSG00000115364 | MRPL19 | 1956.297 | 0.269 | 0.000589941 |
| ENSG00000115977 | AAK1 | 2576.610 | 0.268 | 0.01163418 |
| ENSG00000150753 | CCT5 | 13696.399 | 0.268 | 0.036045873 |
| ENSG00000141298 | SSH2 | 1477.049 | 0.268 | 0.003786577 |
| ENSG00000130758 | MAP3K10 | 1022.854 | 0.268 | 0.032989699 |
| ENSG00000167987 | VPS37C | 1934.907 | 0.267 | 0.000303635 |
| ENSG00000170004 | CHD3 | 10249.791 | 0.267 | 0.024912509 |
| ENSG00000103550 | KNOP1 | 1132.562 | 0.267 | 0.0054746 |
| ENSG00000084693 | AGBL5 | 1943.488 | 0.267 | 0.004109609 |
| ENSG00000088356 | PDRG1 | 1589.902 | 0.267 | 0.018012035 |
| ENSG00000131473 | ACLY | 8891.820 | 0.267 | 0.005518503 |
| ENSG00000068654 | POLR1A | 2988.290 | 0.267 | 0.003768134 |
| ENSG00000115307 | AUP1 | 9785.911 | 0.266 | 0.002560666 |
| ENSG00000122515 | ZMIZ2 | 6396.379 | 0.265 | 0.005615053 |
| ENSG00000105197 | TIMM50 | 3580.340 | 0.265 | 0.009073693 |
| ENSG00000213780 | GTF2H4 | 153.631 | 0.265 | 0.007531244 |
| ENSG00000119640 | ACYP1 | 495.308 | 0.265 | 0.028600424 |
| ENSG00000165684 | SNAPC4 | 1209.431 | 0.264 | 0.007671762 |
| ENSG00000148498 | PARD3 | 3373.379 | 0.264 | 0.024671052 |
| ENSG00000064313 | TAF2 | 1718.221 | 0.263 | 0.045087961 |
| ENSG00000204463 | BAG6 | 12786.293 | 0.263 | 5.99267E-05 |
| ENSG00000189042 | ZNF567 | 324.121 | 0.263 | 0.034413906 |
| ENSG00000251192 | ZNF674 | 140.190 | 0.263 | 0.002763181 |
| ENSG00000173611 | SCAI | 454.126 | 0.263 | 0.012520844 |
| ENSG00000158417 | EIF5B | 8090.457 | 0.262 | 0.020613551 |
| ENSG00000164715 | LMTK2 | 2159.607 | 0.262 | 0.039044211 |
| ENSG00000138081 | FBXO11 | 1224.924 | 0.262 | 0.027394773 |
| ENSG00000176624 | MEX3C | 1742.566 | 0.261 | 0.011496358 |
| ENSG00000151461 | UPF2 | 2369.001 | 0.260 | 0.005425595 |
| ENSG00000134308 | YWHAQ | 14077.411 | 0.260 | 0.030517873 |
| ENSG00000243667 | WDR92 | 98.550 | 0.260 | 0.019178383 |
| ENSG00000100426 | ZBED4 | 1499.437 | 0.260 | 0.008547561 |
| ENSG00000143436 | MRPL9 | 2569.126 | 0.260 | 0.009496367 |
| ENSG00000054965 | FAM168A | 2237.109 | 0.260 | 0.009583872 |
| ENSG00000116679 | IVNS1ABP | 7739.090 | 0.260 | 0.039632219 |
| ENSG00000188010 | MORN2 | 682.954 | 0.260 | 0.032587372 |
| ENSG00000143514 | TP53BP2 | 2185.651 | 0.259 | 0.006058599 |
| ENSG00000075292 | ZNF638 | 4861.965 | 0.259 | 4.74575E-05 |

| | | | | |
|---|---|---|---|---|
| ENSG00000164610 | RP9 | 459.472 | 0.258 | 0.009855743 |
| ENSG00000187189 | TSPYL4 | 1181.725 | 0.258 | 0.029008733 |
| ENSG00000117543 | DPH5 | 855.132 | 0.258 | 0.002608207 |
| ENSG00000117602 | RCAN3 | 1733.715 | 0.257 | 0.046949903 |
| ENSG00000152117 | SMPD4BP | 1163.244 | 0.256 | 0.039846082 |
| ENSG00000162521 | RBBP4 | 8390.493 | 0.256 | 0.007153722 |
| ENSG00000143393 | PI4KB | 4995.065 | 0.255 | 0.008830455 |
| ENSG00000135624 | CCT7 | 14453.531 | 0.255 | 0.005196235 |
| ENSG00000111602 | TIMELESS | 3034.121 | 0.255 | 0.032521731 |
| ENSG00000196151 | WDSUB1 | 433.654 | 0.255 | 0.005447367 |
| ENSG00000073536 | NLE1 | 1271.429 | 0.254 | 0.022539232 |
| ENSG00000166326 | TRIM44 | 3578.546 | 0.254 | 0.01131244 |
| ENSG00000204576 | PRR3 | 804.007 | 0.254 | 0.023151875 |
| ENSG00000144134 | RABL2A | 250.868 | 0.253 | 0.037352304 |
| ENSG00000128581 | IFT22 | 963.661 | 0.253 | 0.010939096 |
| ENSG00000174013 | FBXO45 | 1672.506 | 0.252 | 0.0277419 |
| ENSG00000132780 | NASP | 7570.329 | 0.252 | 0.012784219 |
| ENSG00000144635 | DYNC1LI1 | 1781.320 | 0.251 | 0.001594504 |
| ENSG00000116691 | MIIP | 2086.042 | 0.251 | 0.048411363 |
| ENSG00000099956 | SMARCB1 | 4695.320 | 0.251 | 0.006325 |
| ENSG00000140265 | ZSCAN29 | 1038.606 | 0.251 | 0.011270819 |
| ENSG00000115392 | FANCL | 1057.534 | 0.251 | 0.021111441 |
| ENSG00000125834 | STK35 | 3065.310 | 0.251 | 0.015349349 |
| ENSG00000213923 | CSNK1E | 7508.895 | 0.250 | 0.014955536 |
| ENSG00000162694 | EXTL2 | 773.894 | 0.250 | 0.025704916 |
| ENSG00000110442 | COMMD9 | 1748.241 | 0.250 | 0.017406504 |
| ENSG00000165512 | ZNF22 | 989.095 | 0.250 | 0.016682088 |
| ENSG00000115548 | KDM3A | 2923.876 | 0.250 | 0.031636816 |
| ENSG00000253719 | ATXN7L3B | 5443.145 | 0.249 | 0.015368622 |
| ENSG00000233230 | | 40.759 | 0.249 | 0.035327125 |
| ENSG00000280789 | PAGR1 | 452.491 | 0.249 | 0.027220695 |
| ENSG00000115946 | PNO1 | 1349.198 | 0.249 | 0.002934036 |
| ENSG00000169021 | UQCRFS1 | 3353.582 | 0.249 | 0.041635702 |
| ENSG00000102316 | MAGED2 | 7902.508 | 0.248 | 0.036012315 |
| ENSG00000135521 | LTV1 | 1189.713 | 0.248 | 0.003683578 |
| ENSG00000081721 | DUSP12 | 1009.844 | 0.247 | 0.035438501 |
| ENSG00000147687 | TATDN1 | 961.511 | 0.247 | 0.01137452 |
| ENSG00000169193 | CCDC126 | 383.477 | 0.247 | 0.025136307 |
| ENSG00000066557 | LRRC40 | 697.698 | 0.247 | 0.009072365 |
| ENSG00000197362 | ZNF786 | 507.566 | 0.247 | 0.010172439 |
| ENSG00000196470 | SIAH1 | 728.657 | 0.246 | 0.017311666 |
| ENSG00000040275 | SPDL1 | 699.480 | 0.246 | 0.031441056 |
| ENSG00000118655 | DCLRE1B | 625.159 | 0.246 | 0.034708653 |
| ENSG00000172171 | TEFM | 376.689 | 0.246 | 0.00389176 |
| ENSG00000113360 | DROSHA | 3385.575 | 0.246 | 0.014479214 |
| ENSG00000143363 | PRUNE1 | 1806.448 | 0.245 | 0.025127338 |
| ENSG00000143924 | EML4 | 3850.163 | 0.245 | 0.027519035 |
| ENSG00000157800 | SLC37A3 | 1877.812 | 0.244 | 0.001860313 |
| ENSG00000160917 | CPSF4 | 1711.098 | 0.244 | 0.00827346 |
| ENSG00000163214 | DHX57 | 1424.374 | 0.244 | 0.002122803 |
| ENSG00000136811 | ODF2 | 2857.131 | 0.244 | 0.003115158 |
| ENSG00000173960 | UBXN2A | 1306.483 | 0.244 | 0.003861964 |
| ENSG00000183513 | COA5 | 1255.372 | 0.244 | 0.008104622 |
| ENSG00000108587 | GOSR1 | 2373.124 | 0.244 | 0.000574422 |
| ENSG00000149480 | MTA2 | 8328.434 | 0.244 | 0.000509704 |
| ENSG00000169087 | HSPBAP1 | 549.092 | 0.243 | 0.03721929 |
| ENSG00000272604 | | 64.617 | 0.243 | 0.036876387 |
| ENSG00000132763 | MMACHC | 599.691 | 0.243 | 0.010207078 |
| ENSG00000106344 | RBM28 | 1842.935 | 0.242 | 0.000950222 |

| | | | | |
|---|---|---|---|---|
| ENSG00000125633 | CCDC93 | 1371.542 | 0.242 | 0.013533816 |
| ENSG00000159352 | PSMD4 | 9832.556 | 0.242 | 0.02388534 |
| ENSG00000009954 | BAZ1B | 5581.594 | 0.241 | 0.000830816 |
| ENSG00000144736 | SHQ1 | 981.168 | 0.241 | 0.020735149 |
| ENSG00000074356 | NCBP3 | 2232.932 | 0.241 | 0.008408708 |
| ENSG00000271147 | ARMCX5-GPRAS | 154.273 | 0.241 | 0.016095408 |
| ENSG00000176973 | FAM89B | 1620.588 | 0.241 | 0.049694027 |
| ENSG00000101126 | ADNP | 6537.850 | 0.240 | 0.002698458 |
| ENSG00000131899 | LLGL1 | 2169.591 | 0.240 | 0.037864582 |
| ENSG00000006704 | GTF2IRD1 | 2596.697 | 0.240 | 0.027050005 |
| ENSG00000088038 | CNOT3 | 3537.617 | 0.240 | 0.001850381 |
| ENSG00000198920 | KIAA0753 | 885.952 | 0.239 | 0.022001016 |
| ENSG00000163156 | SCNM1 | 1852.304 | 0.239 | 0.042144777 |
| ENSG00000105778 | AVL9 | 2717.147 | 0.239 | 0.014636956 |
| ENSG00000132383 | RPA1 | 4452.109 | 0.239 | 0.018139669 |
| ENSG00000151233 | GXYLT1 | 1052.806 | 0.239 | 0.038371061 |
| ENSG00000204392 | LSM2 | 2346.899 | 0.238 | 0.028599657 |
| ENSG00000267796 | LIN37 | 229.622 | 0.238 | 0.029546173 |
| ENSG00000165609 | NUDT5 | 3642.296 | 0.238 | 0.026033498 |
| ENSG00000185730 | ZNF696 | 372.613 | 0.238 | 0.03284533 |
| ENSG00000119203 | CPSF3 | 2776.866 | 0.238 | 0.00183227 |
| ENSG00000125450 | NUP85 | 3062.376 | 0.238 | 0.001850381 |
| ENSG00000275111 | ZNF2 | 257.132 | 0.238 | 0.00299313 |
| ENSG00000084774 | CAD | 3774.246 | 0.238 | 0.033060369 |
| ENSG00000164889 | SLC4A2 | 4158.491 | 0.238 | 0.01262226 |
| ENSG00000111676 | ATN1 | 9889.142 | 0.238 | 0.004730388 |
| ENSG00000111196 | MAGOHB | 935.916 | 0.237 | 0.032146532 |
| ENSG00000143368 | SF3B4 | 6302.413 | 0.237 | 0.007931583 |
| ENSG00000160551 | TAOK1 | 4029.520 | 0.237 | 0.039089549 |
| ENSG00000116641 | DOCK7 | 1677.319 | 0.237 | 0.026882374 |
| ENSG00000006634 | DBF4 | 770.413 | 0.237 | 0.038762484 |
| ENSG00000168000 | BSCL2 | 352.233 | 0.237 | 0.02399852 |
| ENSG00000136937 | NCBP1 | 2168.214 | 0.237 | 0.009246763 |
| ENSG00000151849 | CENPJ | 740.478 | 0.237 | 0.040658976 |
| ENSG00000196505 | GDAP2 | 907.882 | 0.237 | 0.0083842 |
| ENSG00000080845 | DLGAP4 | 5448.343 | 0.236 | 0.004458488 |
| ENSG00000213516 | RBMXL1 | 1083.312 | 0.236 | 0.016069211 |
| ENSG00000162065 | TBC1D24 | 723.116 | 0.235 | 0.016653705 |
| ENSG00000137500 | CCDC90B | 1791.332 | 0.235 | 0.004933013 |
| ENSG00000158623 | COPG2 | 2837.364 | 0.234 | 0.020754117 |
| ENSG00000120685 | PROSER1 | 2399.573 | 0.234 | 0.046813755 |
| ENSG00000176390 | CRLF3 | 924.966 | 0.234 | 0.012154863 |
| ENSG00000004142 | POLDIP2 | 6625.280 | 0.234 | 0.002915901 |
| ENSG00000134077 | THUMPD3 | 1925.150 | 0.233 | 0.010818498 |
| ENSG00000163781 | TOPBP1 | 2479.389 | 0.233 | 0.031067985 |
| ENSG00000163257 | DCAF16 | 1646.912 | 0.233 | 0.023332431 |
| ENSG00000141013 | GAS8 | 908.720 | 0.233 | 0.035327125 |
| ENSG00000152219 | ARL14EP | 778.845 | 0.233 | 0.009169916 |
| ENSG00000105968 | H2AZ2 | 6459.677 | 0.233 | 0.007339643 |
| ENSG00000160688 | FLAD1 | 3114.895 | 0.232 | 0.01843053 |
| ENSG00000073584 | SMARCE1 | 3998.165 | 0.232 | 0.01426157 |
| ENSG00000178177 | LCORL | 449.703 | 0.232 | 0.037724632 |
| ENSG00000155111 | CDK19 | 1168.410 | 0.232 | 0.02542107 |
| ENSG00000108953 | YWHAE | 25177.189 | 0.232 | 0.015312948 |
| ENSG00000175931 | UBE2O | 2791.496 | 0.232 | 0.003113934 |
| ENSG00000164828 | SUN1 | 6055.013 | 0.231 | 0.028112937 |
| ENSG00000163001 | CFAP36 | 1475.955 | 0.231 | 0.016750721 |
| ENSG00000213300 | HNRNPA3P6 | 46.250 | 0.231 | 0.046282432 |
| ENSG00000144021 | CIAO1 | 3728.174 | 0.231 | 0.000234009 |

| | | | | |
|---|---|---|---|---|
| ENSG00000177084 | POLE | 3063.083 | 0.231 | 0.042384641 |
| ENSG00000106305 | AIMP2 | 1055.612 | 0.231 | 0.032987554 |
| ENSG00000090447 | TFAP4 | 983.179 | 0.230 | 0.043091157 |
| ENSG00000132768 | DPH2 | 1548.914 | 0.230 | 0.013056033 |
| ENSG00000136715 | SAP130 | 1721.488 | 0.229 | 0.005746497 |
| ENSG00000196214 | ZNF766 | 1130.313 | 0.229 | 0.012441472 |
| ENSG00000141076 | UTP4 | 2704.189 | 0.229 | 0.018599136 |
| ENSG00000064703 | DDX20 | 864.755 | 0.229 | 0.002562008 |
| ENSG00000134748 | PRPF38A | 2368.885 | 0.228 | 0.000539548 |
| ENSG00000159593 | NAE1 | 2572.973 | 0.228 | 0.007817202 |
| ENSG00000129351 | ILF3 | 17371.224 | 0.228 | 0.003212962 |
| ENSG00000141956 | PRDM15 | 630.523 | 0.228 | 0.019697265 |
| ENSG00000184634 | MED12 | 2668.013 | 0.227 | 0.016412955 |
| ENSG00000135972 | MRPS9 | 1391.112 | 0.227 | 0.005913632 |
| ENSG00000100150 | DEPDC5 | 974.901 | 0.227 | 0.023868362 |
| ENSG00000182670 | TTC3 | 7853.686 | 0.227 | 0.042626005 |
| ENSG00000089094 | KDM2B | 1984.958 | 0.226 | 0.003223787 |
| ENSG00000168522 | FNTA | 3038.020 | 0.226 | 0.034708653 |
| ENSG00000172939 | OXSR1 | 3430.977 | 0.226 | 0.027022954 |
| ENSG00000108423 | TUBD1 | 446.605 | 0.226 | 0.025636288 |
| ENSG00000142528 | ZNF473 | 658.193 | 0.225 | 0.024206955 |
| ENSG00000184677 | ZBTB40 | 1439.541 | 0.225 | 0.03333025 |
| ENSG00000187742 | SECISBP2 | 2038.164 | 0.225 | 0.013530427 |
| ENSG00000105821 | DNAJC2 | 1979.946 | 0.224 | 0.005562571 |
| ENSG00000129219 | PLD2 | 2196.131 | 0.224 | 0.042916272 |
| ENSG00000168883 | USP39 | 3047.660 | 0.224 | 0.00073983 |
| ENSG00000259623 | | 741.355 | 0.224 | 0.014308309 |
| ENSG00000153879 | CEBPG | 3375.252 | 0.223 | 0.024652009 |
| ENSG00000156170 | NDUFAF6 | 722.756 | 0.223 | 0.023992693 |
| ENSG00000028310 | BRD9 | 2619.867 | 0.223 | 0.030172251 |
| ENSG00000161956 | SENP3 | 1237.974 | 0.223 | 0.019405438 |
| ENSG00000115241 | PPM1G | 9011.602 | 0.222 | 0.010420465 |
| ENSG00000142453 | CARM1 | 2573.402 | 0.222 | 0.012452996 |
| ENSG00000189164 | ZNF527 | 229.179 | 0.222 | 0.043483169 |
| ENSG00000160803 | UBQLN4 | 3032.518 | 0.221 | 0.019786609 |
| ENSG00000204843 | DCTN1 | 6527.802 | 0.221 | 0.009078732 |
| ENSG00000070061 | ELP1 | 1767.750 | 0.221 | 0.016052872 |
| ENSG00000221838 | AP4M1 | 1264.428 | 0.221 | 0.030614432 |
| ENSG00000153561 | RMND5A | 2968.944 | 0.220 | 0.036950755 |
| ENSG00000105865 | DUS4L | 482.179 | 0.220 | 0.014268734 |
| ENSG00000158435 | CNOT11 | 3557.108 | 0.220 | 0.015966835 |
| ENSG00000157020 | SEC13 | 7095.245 | 0.220 | 0.043204227 |
| ENSG00000133316 | WDR74 | 1761.428 | 0.219 | 0.036117843 |
| ENSG00000163125 | RPRD2 | 2713.083 | 0.219 | 0.049082448 |
| ENSG00000197622 | CDC42SE1 | 8615.922 | 0.218 | 0.034413906 |
| ENSG00000143919 | CAMKMT | 329.708 | 0.218 | 0.047596425 |
| ENSG00000158411 | MITD1 | 951.591 | 0.218 | 0.017115128 |
| ENSG00000101557 | USP14 | 4175.629 | 0.217 | 0.033060369 |
| ENSG00000092847 | AGO1 | 1903.684 | 0.217 | 0.009908836 |
| ENSG00000109917 | ZPR1 | 2493.143 | 0.217 | 0.011613955 |
| ENSG00000147274 | RBMX | 7559.460 | 0.216 | 0.000497738 |
| ENSG00000213024 | NUP62 | 4928.555 | 0.216 | 0.005071056 |
| ENSG00000126261 | UBA2 | 5882.349 | 0.216 | 0.023868362 |
| ENSG00000058804 | NDC1 | 2372.426 | 0.216 | 0.042899903 |
| ENSG00000032742 | IFT88 | 480.095 | 0.216 | 0.037419573 |
| ENSG00000146576 | C7orf26 | 1612.586 | 0.215 | 0.009529074 |
| ENSG00000102900 | NUP93 | 2955.284 | 0.215 | 0.016900686 |
| ENSG00000079785 | DDX1 | 5646.127 | 0.215 | 0.003111161 |
| ENSG00000121774 | KHDRBS1 | 9371.008 | 0.215 | 4.33762E-05 |

| | | | | |
|---|---|---|---|---|
| ENSG00000040341 | STAU2 | 1676.422 | 0.214 | 0.049727325 |
| ENSG00000141127 | PRPSAP2 | 1209.430 | 0.214 | 0.036333487 |
| ENSG00000243927 | MRPS6 | 2056.104 | 0.212 | 0.042878413 |
| ENSG00000119878 | CRIPT | 1065.508 | 0.212 | 0.017160556 |
| ENSG00000115539 | PDCL3 | 1386.230 | 0.212 | 0.016812411 |
| ENSG00000105576 | TNPO2 | 4333.771 | 0.212 | 0.005293957 |
| ENSG00000163041 | H3-3A | 1760.237 | 0.212 | 0.031780858 |
| ENSG00000264364 | DYNLL2 | 7455.523 | 0.211 | 0.027656709 |
| ENSG00000132313 | MRPL35 | 1947.049 | 0.211 | 0.001060696 |
| ENSG00000099783 | HNRNPM | 12885.434 | 0.210 | 0.002393018 |
| ENSG00000163138 | PACRGL | 490.376 | 0.209 | 0.01459316 |
| ENSG00000204574 | ABCF1 | 5690.598 | 0.209 | 0.015838664 |
| ENSG00000177733 | HNRNPA0 | 9170.852 | 0.208 | 0.005370069 |
| ENSG00000165055 | METTL2B | 1197.262 | 0.208 | 0.009634152 |
| ENSG00000105323 | HNRNPUL1 | 14680.921 | 0.207 | 0.010100583 |
| ENSG00000163479 | SSR2 | 13151.373 | 0.207 | 0.033352301 |
| ENSG00000176102 | CSTF3 | 1711.353 | 0.207 | 0.009015389 |
| ENSG00000086712 | TXLNG | 1094.577 | 0.207 | 0.043873106 |
| ENSG00000145919 | BOD1 | 1922.280 | 0.207 | 0.009246763 |
| ENSG00000182952 | HMGN4 | 2609.295 | 0.205 | 0.03706252 |
| ENSG00000064419 | TNPO3 | 3589.788 | 0.205 | 0.005422213 |
| ENSG00000119812 | FAM98A | 2477.318 | 0.205 | 0.020817029 |
| ENSG00000149187 | CELF1 | 5461.629 | 0.204 | 0.00066234 |
| ENSG00000275066 | SYNRG | 2471.429 | 0.204 | 0.029132033 |
| ENSG00000169016 | E2F6 | 579.954 | 0.204 | 0.005409545 |
| ENSG00000136450 | SRSF1 | 9548.198 | 0.204 | 0.000200429 |
| ENSG00000117448 | AKR1A1 | 6898.754 | 0.203 | 0.03706252 |
| ENSG00000111642 | CHD4 | 13548.226 | 0.203 | 0.000871447 |
| ENSG00000136699 | SMPD4 | 3571.730 | 0.203 | 0.006113677 |
| ENSG00000143933 | CALM2 | 17934.611 | 0.203 | 0.036819776 |
| ENSG00000197451 | HNRNPAB | 9096.709 | 0.202 | 0.016271117 |
| ENSG00000174442 | ZWILCH | 1128.070 | 0.202 | 0.04377522 |
| ENSG00000143207 | COP1 | 3115.128 | 0.202 | 0.048231523 |
| ENSG00000180423 | HARBI1 | 174.656 | 0.202 | 0.037284715 |
| ENSG00000137947 | GTF2B | 1603.737 | 0.201 | 0.019907235 |
| ENSG00000125351 | UPF3B | 992.648 | 0.201 | 0.047530238 |
| ENSG00000204469 | PRRC2A | 14731.799 | 0.201 | 0.00900218 |
| ENSG00000151465 | CDC123 | 3987.323 | 0.201 | 0.032989699 |
| ENSG00000168256 | NKIRAS2 | 3164.681 | 0.200 | 0.002921401 |
| ENSG00000011143 | MKS1 | 905.060 | 0.200 | 0.031864392 |
| ENSG00000170296 | GABARAP | 2440.575 | 0.200 | 0.03494497 |
| ENSG00000117360 | PRPF3 | 2645.547 | 0.200 | 0.045403698 |
| ENSG00000125962 | ARMCX5 | 561.382 | 0.199 | 0.04172789 |
| ENSG00000124380 | SNRNP27 | 1213.527 | 0.199 | 0.006367142 |
| ENSG00000204560 | DHX16 | 2775.183 | 0.199 | 0.006670951 |
| ENSG00000113838 | TBCCD1 | 648.169 | 0.199 | 0.038258344 |
| ENSG00000276234 | TADA2A | 956.665 | 0.198 | 0.02657647 |
| ENSG00000198060 | MARCHF5 | 2080.356 | 0.198 | 0.013662392 |
| ENSG00000102078 | SLC25A14 | 436.928 | 0.198 | 0.023037543 |
| ENSG00000114491 | UMPS | 2160.808 | 0.198 | 0.032356225 |
| ENSG00000147130 | ZMYM3 | 2077.191 | 0.197 | 0.03428612 |
| ENSG00000132475 | H3-3B | 37885.774 | 0.197 | 0.026043181 |
| ENSG00000206562 | METTL6 | 566.808 | 0.197 | 0.01383871 |
| ENSG00000114956 | DGUOK | 2907.017 | 0.196 | 0.02781366 |
| ENSG00000204954 | C12orf73 | 426.750 | 0.196 | 0.046849922 |
| ENSG00000157954 | WIPI2 | 4762.150 | 0.196 | 0.008558202 |
| ENSG00000105221 | AKT2 | 6407.701 | 0.195 | 0.022677887 |
| ENSG00000136273 | HUS1 | 795.918 | 0.195 | 0.012934592 |
| ENSG00000126653 | NSRP1 | 1642.211 | 0.195 | 0.010945595 |

| | | | | |
|---|---|---|---|---|
| ENSG00000181090 | EHMT1 | 3122.287 | 0.193 | 0.009057359 |
| ENSG00000074054 | CLASP1 | 2418.103 | 0.193 | 0.020149201 |
| ENSG00000198000 | NOL8 | 1699.634 | 0.193 | 0.032980611 |
| ENSG00000231074 | HCG18 | 1166.266 | 0.192 | 0.032474434 |
| ENSG00000141556 | TBCD | 5341.520 | 0.192 | 0.026836222 |
| ENSG00000010244 | ZNF207 | 7768.214 | 0.191 | 0.00441843 |
| ENSG00000172466 | ZNF24 | 3070.520 | 0.190 | 0.030989011 |
| ENSG00000173933 | RBM4 | 671.244 | 0.190 | 0.018595082 |
| ENSG00000105723 | GSK3A | 4429.019 | 0.190 | 0.019542616 |
| ENSG00000172534 | HCFC1 | 5384.728 | 0.190 | 0.010338469 |
| ENSG00000203879 | GDI1 | 7770.940 | 0.190 | 0.037352304 |
| ENSG00000138231 | DBR1 | 834.917 | 0.190 | 0.016735537 |
| ENSG00000211460 | TSN | 3704.156 | 0.189 | 0.014795356 |
| ENSG00000133884 | DPF2 | 2500.646 | 0.189 | 0.007421838 |
| ENSG00000107643 | MAPK8 | 1376.858 | 0.188 | 0.04569585 |
| ENSG00000155906 | RMND1 | 815.939 | 0.188 | 0.045489837 |
| ENSG00000196235 | SUPT5H | 8583.377 | 0.187 | 0.020560234 |
| ENSG00000198900 | TOP1 | 5152.888 | 0.187 | 0.04978925 |
| ENSG00000170144 | HNRNPA3 | 13937.656 | 0.187 | 0.005821514 |
| ENSG00000188976 | NOC2L | 6833.784 | 0.187 | 0.041928868 |
| ENSG00000198522 | GPN1 | 2158.071 | 0.186 | 0.023233255 |
| ENSG00000009844 | VTA1 | 2475.545 | 0.186 | 0.017992676 |
| ENSG00000148153 | INIP | 1124.765 | 0.186 | 0.041356882 |
| ENSG00000116560 | SFPQ | 15329.116 | 0.185 | 0.001126803 |
| ENSG00000101407 | TTI1 | 1834.213 | 0.184 | 0.035438501 |
| ENSG00000160062 | ZBTB8A | 433.488 | 0.184 | 0.043148387 |
| ENSG00000162419 | GMEB1 | 1087.561 | 0.184 | 0.010413205 |
| ENSG00000163161 | ERCC3 | 2559.534 | 0.183 | 0.003880309 |
| ENSG00000124160 | NCOA5 | 2411.330 | 0.183 | 0.024671052 |
| ENSG00000135829 | DHX9 | 8946.342 | 0.182 | 0.01477739 |
| ENSG00000088247 | KHSRP | 10374.638 | 0.182 | 0.008658662 |
| ENSG00000141219 | C17orf80 | 990.915 | 0.181 | 0.026836222 |
| ENSG00000010539 | ZNF200 | 531.799 | 0.180 | 0.032992198 |
| ENSG00000168246 | UBTD2 | 1374.115 | 0.179 | 0.040756665 |
| ENSG00000106144 | CASP2 | 2795.467 | 0.178 | 0.045095666 |
| ENSG00000134697 | GNL2 | 3980.730 | 0.178 | 0.045400735 |
| ENSG00000116809 | ZBTB17 | 1731.928 | 0.178 | 0.029290812 |
| ENSG00000167986 | DDB1 | 13574.888 | 0.177 | 0.027557677 |
| ENSG00000112983 | BRD8 | 2106.548 | 0.177 | 0.043148387 |
| ENSG00000173914 | RBM4B | 941.866 | 0.176 | 0.039932053 |
| ENSG00000087157 | PGS1 | 1766.033 | 0.175 | 0.020820196 |
| ENSG00000087995 | METTL2A | 895.890 | 0.175 | 0.030000209 |
| ENSG00000172530 | BANP | 738.690 | 0.174 | 0.024893749 |
| ENSG00000155508 | CNOT8 | 2648.100 | 0.172 | 0.027929247 |
| ENSG00000114982 | KANSL3 | 2643.576 | 0.171 | 0.024882953 |
| ENSG00000162300 | ZFPL1 | 428.277 | 0.170 | 0.03706252 |
| ENSG00000182973 | CNOT10 | 1657.736 | 0.168 | 0.032056036 |
| ENSG00000087087 | SRRT | 6930.172 | 0.168 | 0.013470563 |
| ENSG00000085760 | MTIF2 | 2015.841 | 0.168 | 0.024209264 |
| ENSG00000115875 | SRSF7 | 6279.147 | 0.168 | 0.004352281 |
| ENSG00000144233 | AMMECR1L | 1577.055 | 0.167 | 0.030275633 |
| ENSG00000115282 | TTC31 | 1348.515 | 0.167 | 0.040600605 |
| ENSG00000078369 | GNB1 | 18774.633 | 0.166 | 0.01776262 |
| ENSG00000124356 | STAMBP | 2332.091 | 0.165 | 0.012577323 |
| ENSG00000122566 | HNRNPA2B1 | 41693.634 | 0.164 | 0.005421544 |
| ENSG00000009307 | CSDE1 | 25263.975 | 0.163 | 0.043770914 |
| ENSG00000011485 | PPP5C | 3356.950 | 0.163 | 0.035692545 |
| ENSG00000066044 | ELAVL1 | 4846.802 | 0.163 | 0.004214195 |
| ENSG00000105793 | GTPBP10 | 935.473 | 0.162 | 0.023920072 |

| | | | | |
|---|---|---|---|---|
| ENSG00000146872 | TLK2 | 1752.603 | 0.162 | 0.027934014 |
| ENSG00000104064 | GABPB1 | 1588.901 | 0.162 | 0.006698207 |
| ENSG00000116752 | BCAS2 | 1887.841 | 0.162 | 0.042027658 |
| ENSG00000096746 | HNRNPH3 | 6098.825 | 0.160 | 0.015530167 |
| ENSG00000066117 | SMARCD1 | 3577.671 | 0.160 | 0.044275973 |
| ENSG00000181852 | RNF41 | 1946.269 | 0.158 | 0.034537566 |
| ENSG00000130935 | NOL11 | 2963.009 | 0.158 | 0.043600672 |
| ENSG00000070770 | CSNK2A2 | 3715.371 | 0.158 | 0.043714542 |
| ENSG00000119820 | YIPF4 | 2789.368 | 0.157 | 0.021646301 |
| ENSG00000149532 | CPSF7 | 4469.036 | 0.157 | 0.01032959 |
| ENSG00000003509 | NDUFAF7 | 668.566 | 0.155 | 0.022932462 |
| ENSG00000141568 | FOXK2 | 3731.898 | 0.153 | 0.030659094 |
| ENSG00000011451 | WIZ | 3521.929 | 0.152 | 0.046390229 |
| ENSG00000163798 | SLC4A1AP | 1519.350 | 0.152 | 0.03284533 |
| ENSG00000106459 | NRF1 | 956.571 | 0.152 | 0.004688206 |
| ENSG00000109111 | SUPT6H | 7323.180 | 0.152 | 0.035324036 |
| ENSG00000163166 | IWS1 | 2769.515 | 0.151 | 0.025260562 |
| ENSG00000143889 | HNRNPLL | 1567.682 | 0.151 | 0.013523272 |
| ENSG00000119760 | SUPT7L | 2289.903 | 0.147 | 0.021819231 |
| ENSG00000011304 | PTBP1 | 16380.769 | 0.147 | 0.009417887 |
| ENSG00000182944 | EWSR1 | 14021.900 | 0.146 | 0.014464638 |
| ENSG00000153187 | HNRNPU | 19814.934 | 0.146 | 0.01132374 |
| ENSG00000104824 | HNRNPL | 11481.759 | 0.146 | 0.010522886 |
| ENSG00000135974 | C2orf49 | 880.692 | 0.142 | 0.035906201 |
| ENSG00000165119 | HNRNPK | 25998.186 | 0.120 | 0.047862073 |
| ENSG00000120948 | TARDBP | 6160.879 | 0.117 | 0.008197086 |
| ENSG00000159322 | ADPGK | 2758.140 | -0.145 | 0.048228204 |
| ENSG00000117481 | NSUN4 | 1403.708 | -0.151 | 0.044106066 |
| ENSG00000166887 | VPS39 | 3262.027 | -0.152 | 0.035459993 |
| ENSG00000100461 | RBM23 | 3223.963 | -0.153 | 0.039089549 |
| ENSG00000100395 | L3MBTL2 | 1861.436 | -0.154 | 0.041346535 |
| ENSG00000029364 | SLC39A9 | 3677.990 | -0.154 | 0.046825923 |
| ENSG00000173011 | TADA2B | 1416.630 | -0.155 | 0.035924736 |
| ENSG00000158480 | SPATA2 | 931.987 | -0.157 | 0.047620001 |
| ENSG00000111652 | COPS7A | 4597.344 | -0.158 | 0.045551524 |
| ENSG00000109332 | UBE2D3 | 10590.766 | -0.158 | 0.033539494 |
| ENSG00000107862 | GBF1 | 4004.047 | -0.159 | 0.041908666 |
| ENSG00000070831 | CDC42 | 13412.959 | -0.160 | 0.022605204 |
| ENSG00000113558 | SKP1 | 9198.682 | -0.160 | 0.039004294 |
| ENSG00000171307 | ZDHHC16 | 2012.474 | -0.164 | 0.044132357 |
| ENSG00000164219 | PGGT1B | 1100.348 | -0.166 | 0.039845806 |
| ENSG00000140153 | WDR20 | 881.501 | -0.168 | 0.017517283 |
| ENSG00000142166 | IFNAR1 | 3670.663 | -0.172 | 0.039049006 |
| ENSG00000152700 | SAR1B | 2543.546 | -0.172 | 0.041922973 |
| ENSG00000204231 | RXRB | 2306.939 | -0.172 | 0.013840961 |
| ENSG00000135655 | USP15 | 2357.607 | -0.173 | 0.043585257 |
| ENSG00000014919 | COX15 | 1736.970 | -0.174 | 0.024041583 |
| ENSG00000120029 | ARMH3 | 1396.285 | -0.174 | 0.014410561 |
| ENSG00000138802 | SEC24B | 1934.048 | -0.174 | 0.048276722 |
| ENSG00000142687 | KIAA0319L | 3496.907 | -0.175 | 0.020638593 |
| ENSG00000125952 | MAX | 2790.791 | -0.175 | 0.015838664 |
| ENSG00000136436 | CALCOCO2 | 4616.469 | -0.176 | 0.040644579 |
| ENSG00000022840 | RNF10 | 9288.273 | -0.177 | 0.011836007 |
| ENSG00000077549 | CAPZB | 14189.184 | -0.177 | 0.024424542 |
| ENSG00000035687 | ADSS2 | 3424.120 | -0.178 | 0.028734724 |
| ENSG00000119977 | TCTN3 | 1975.805 | -0.178 | 0.021841283 |
| ENSG00000100239 | PPP6R2 | 3062.871 | -0.178 | 0.037999061 |
| ENSG00000124226 | RNF114 | 3702.380 | -0.180 | 0.032146532 |
| ENSG00000163684 | RPP14 | 791.847 | -0.181 | 0.024088638 |

| | | | | |
|---|---|---|---|---|
| ENSG00000214046 | SMIM7 | 2051.309 | -0.182 | 0.018072324 |
| ENSG00000136146 | MED4 | 2007.257 | -0.182 | 0.026881859 |
| ENSG00000100138 | SNU13 | 6246.168 | -0.183 | 0.046731941 |
| ENSG00000125734 | GPR108 | 3087.346 | -0.183 | 0.028294392 |
| ENSG00000145740 | SLC30A5 | 2133.339 | -0.185 | 0.017152181 |
| ENSG00000150459 | SAP18 | 6315.663 | -0.185 | 0.026117288 |
| ENSG00000122783 | CYREN | 1433.335 | -0.187 | 0.041283268 |
| ENSG00000184007 | PTP4A2 | 11446.849 | -0.187 | 0.034574941 |
| ENSG00000137806 | NDUFAF1 | 730.233 | -0.187 | 0.040770719 |
| ENSG00000154305 | MIA3 | 3047.773 | -0.187 | 0.017699907 |
| ENSG00000100991 | TRPC4AP | 6221.829 | -0.188 | 0.036950755 |
| ENSG00000104660 | LEPROTL1 | 1788.399 | -0.188 | 0.036842327 |
| ENSG00000139990 | DCAF5 | 2962.491 | -0.189 | 0.016119944 |
| ENSG00000113621 | TXNDC15 | 1787.941 | -0.189 | 0.034413906 |
| ENSG00000162704 | ARPC5 | 10392.161 | -0.189 | 0.024187694 |
| ENSG00000164073 | MFSD8 | 683.078 | -0.189 | 0.024826701 |
| ENSG00000103978 | TMEM87A | 4495.692 | -0.189 | 0.030327887 |
| ENSG00000144848 | ATG3 | 2283.953 | -0.189 | 0.018819126 |
| ENSG00000071537 | SEL1L | 3966.223 | -0.189 | 0.040301133 |
| ENSG00000067560 | RHOA | 23184.351 | -0.190 | 0.023842686 |
| ENSG00000135775 | COG2 | 1372.039 | -0.190 | 0.009728068 |
| ENSG00000105438 | KDELR1 | 13410.679 | -0.190 | 0.023838463 |
| ENSG00000119720 | NRDE2 | 840.615 | -0.190 | 0.021956402 |
| ENSG00000079999 | KEAP1 | 4919.022 | -0.191 | 0.032992198 |
| ENSG00000188725 | SMIM15 | 2537.733 | -0.191 | 0.018771665 |
| ENSG00000035681 | NSMAF | 1784.690 | -0.191 | 0.037233703 |
| ENSG00000104946 | TBC1D17 | 2435.285 | -0.191 | 0.040840715 |
| ENSG00000075914 | EXOSC7 | 2105.710 | -0.193 | 0.043448669 |
| ENSG00000140941 | MAP1LC3B | 3417.689 | -0.195 | 0.035201594 |
| ENSG00000151148 | UBE3B | 2480.620 | -0.195 | 0.005133584 |
| ENSG00000198663 | C6orf89 | 4273.933 | -0.196 | 0.004485456 |
| ENSG00000183283 | DAZAP2 | 12056.568 | -0.196 | 0.02335417 |
| ENSG00000169967 | MAP3K2 | 2562.158 | -0.196 | 0.038518074 |
| ENSG00000166275 | BORCS7 | 723.792 | -0.197 | 0.04098079 |
| ENSG00000103264 | FBXO31 | 1423.492 | -0.197 | 0.03816747 |
| ENSG00000163312 | HELQ | 411.686 | -0.198 | 0.00722434 |
| ENSG00000166797 | CIAO2A | 1697.024 | -0.198 | 0.007257376 |
| ENSG00000105583 | WDR83OS | 5676.780 | -0.199 | 0.032353838 |
| ENSG00000141002 | TCF25 | 5644.520 | -0.199 | 0.019535191 |
| ENSG00000140374 | ETFA | 4459.918 | -0.199 | 0.03284533 |
| ENSG00000136816 | TOR1B | 1411.200 | -0.199 | 0.040845082 |
| ENSG00000065491 | TBC1D22B | 1026.970 | -0.200 | 0.009213535 |
| ENSG00000111725 | PRKAB1 | 1598.193 | -0.200 | 0.043719541 |
| ENSG00000165678 | GHITM | 7409.210 | -0.200 | 0.033116977 |
| ENSG00000100225 | FBXO7 | 4391.568 | -0.201 | 0.00702496 |
| ENSG00000100614 | PPM1A | 2387.073 | -0.202 | 0.014734785 |
| ENSG00000077147 | TM9SF3 | 10457.127 | -0.202 | 0.011658528 |
| ENSG00000164305 | CASP3 | 2330.400 | -0.202 | 0.036199219 |
| ENSG00000198898 | CAPZA2 | 4285.259 | -0.202 | 0.038465763 |
| ENSG00000145354 | CISD2 | 1545.980 | -0.202 | 0.027028011 |
| ENSG00000177981 | ASB8 | 1249.130 | -0.202 | 0.010027575 |
| ENSG00000178425 | NT5DC1 | 1380.755 | -0.203 | 0.042604019 |
| ENSG00000127720 | METTL25 | 210.776 | -0.203 | 0.033176856 |
| ENSG00000164172 | MOCS2 | 1206.630 | -0.205 | 0.020299228 |
| ENSG00000165792 | METTL17 | 1870.227 | -0.205 | 0.041283268 |
| ENSG00000185129 | PURA | 1309.765 | -0.205 | 0.035083394 |
| ENSG00000156639 | ZFAND3 | 3873.186 | -0.206 | 0.004175761 |
| ENSG00000164329 | TENT2 | 1795.860 | -0.207 | 0.008402367 |
| ENSG00000103642 | LACTB | 987.228 | -0.207 | 0.046825923 |

| | | | | |
|---|---|---|---|---|
| ENSG00000137275 | RIPK1 | 2143.919 | -0.207 | 0.014158194 |
| ENSG00000109180 | OCIAD1 | 4987.072 | -0.208 | 0.004437993 |
| ENSG00000100568 | VTI1B | 3435.736 | -0.208 | 0.004092005 |
| ENSG00000161904 | LEMD2 | 3374.548 | -0.208 | 0.006393306 |
| ENSG00000164096 | C4orf3 | 5922.991 | -0.208 | 0.039926184 |
| ENSG00000146828 | SLC12A9 | 3093.934 | -0.209 | 0.040273337 |
| ENSG00000182400 | TRAPPC6B | 1367.245 | -0.209 | 0.00428997 |
| ENSG00000171574 | ZNF584 | 441.180 | -0.210 | 0.037699628 |
| ENSG00000163933 | RFT1 | 1353.600 | -0.210 | 0.008518741 |
| ENSG00000023318 | ERP44 | 3999.249 | -0.210 | 0.022714209 |
| ENSG00000115762 | PLEKHB2 | 4372.839 | -0.212 | 0.038334312 |
| ENSG00000106049 | HIBADH | 1912.093 | -0.212 | 0.04831862 |
| ENSG00000170348 | TMED10 | 13025.077 | -0.212 | 0.009246763 |
| ENSG00000135317 | SNX14 | 2727.214 | -0.213 | 0.021514044 |
| ENSG00000223501 | VPS52 | 2471.238 | -0.213 | 0.005571674 |
| ENSG00000183826 | BTBD9 | 1010.771 | -0.213 | 0.033710304 |
| ENSG00000122912 | SLC25A16 | 867.047 | -0.214 | 0.048265816 |
| ENSG00000104904 | OAZ1 | 25341.796 | -0.214 | 0.038341627 |
| ENSG00000125741 | OPA3 | 1636.572 | -0.214 | 0.018532793 |
| ENSG00000135778 | NTPCR | 1614.804 | -0.214 | 0.018001921 |
| ENSG00000205352 | PRR13 | 4067.682 | -0.215 | 0.024115762 |
| ENSG00000134996 | OSTF1 | 2732.056 | -0.215 | 0.033688928 |
| ENSG00000089006 | SNX5 | 5391.034 | -0.215 | 0.017549337 |
| ENSG00000144659 | SLC25A38 | 2040.750 | -0.215 | 0.037332954 |
| ENSG00000170854 | RIOX2 | 1447.768 | -0.215 | 0.039923116 |
| ENSG00000166295 | ANAPC16 | 4653.307 | -0.215 | 0.015312948 |
| ENSG00000144468 | RHBDD1 | 1152.740 | -0.217 | 0.014943546 |
| ENSG00000107521 | HPS1 | 3043.830 | -0.217 | 0.021087623 |
| ENSG00000102910 | LONP2 | 3826.295 | -0.218 | 0.016788229 |
| ENSG00000131408 | NR1H2 | 4737.537 | -0.218 | 0.009523712 |
| ENSG00000173141 | MRPL57 | 2725.704 | -0.219 | 0.039602386 |
| ENSG00000162852 | CNST | 1043.959 | -0.219 | 0.016376326 |
| ENSG00000166272 | WBP1L | 2884.500 | -0.219 | 0.01664669 |
| ENSG00000090432 | MUL1 | 2019.163 | -0.219 | 0.00787248 |
| ENSG00000054611 | TBC1D22A | 1720.755 | -0.220 | 0.010117016 |
| ENSG00000158985 | CDC42SE2 | 2591.788 | -0.220 | 0.036243248 |
| ENSG00000028528 | SNX1 | 4943.425 | -0.220 | 0.001461594 |
| ENSG00000162434 | JAK1 | 6479.549 | -0.221 | 0.037999061 |
| ENSG00000066739 | ATG2B | 1245.045 | -0.221 | 0.033783748 |
| ENSG00000148719 | DNAJB12 | 2691.907 | -0.222 | 0.003418227 |
| ENSG00000143811 | PYCR2 | 4088.223 | -0.222 | 0.028407688 |
| ENSG00000103496 | STX4 | 2820.365 | -0.222 | 0.01234622 |
| ENSG00000138138 | ATAD1 | 2219.479 | -0.222 | 0.028335004 |
| ENSG00000161048 | NAPEPLD | 780.489 | -0.222 | 0.026524863 |
| ENSG00000073169 | SELENOO | 1706.557 | -0.223 | 0.04831862 |
| ENSG00000132356 | PRKAA1 | 3285.474 | -0.223 | 0.0471781 |
| ENSG00000127870 | RNF6 | 2056.698 | -0.223 | 0.028853317 |
| ENSG00000121851 | POLR3GL | 876.533 | -0.223 | 0.036392367 |
| ENSG00000111269 | CREBL2 | 2023.996 | -0.225 | 0.04123665 |
| ENSG00000105402 | NAPA | 3780.585 | -0.225 | 0.007669961 |
| ENSG00000104805 | NUCB1 | 17271.294 | -0.225 | 0.02297913 |
| ENSG00000125826 | RBCK1 | 6358.026 | -0.225 | 0.022180753 |
| ENSG00000198171 | DDRGK1 | 3512.299 | -0.225 | 0.037419573 |
| ENSG00000023191 | RNH1 | 8143.243 | -0.225 | 0.034304317 |
| ENSG00000174485 | DENND4A | 1091.238 | -0.226 | 0.018185479 |
| ENSG00000092931 | MFSD11 | 1366.285 | -0.226 | 0.020183268 |
| ENSG00000132950 | ZMYM5 | 630.757 | -0.226 | 0.009701804 |
| ENSG00000136003 | ISCU | 3467.838 | -0.226 | 0.00428997 |
| ENSG00000081791 | DELE1 | 2179.646 | -0.227 | 0.010106823 |

| | | | | |
|---|---|---|---|---|
| ENSG00000173418 | NAA20 | 3467.138 | -0.228 | 0.032356225 |
| ENSG00000120697 | ALG5 | 1657.839 | -0.228 | 0.017063177 |
| ENSG00000255062 | | 58.772 | -0.228 | 0.049907621 |
| ENSG00000100528 | CNIH1 | 3205.623 | -0.229 | 0.010078485 |
| ENSG00000242247 | ARFGAP3 | 2219.161 | -0.229 | 0.046292208 |
| ENSG00000100650 | SRSF5 | 8936.980 | -0.229 | 0.04831862 |
| ENSG00000147316 | MCPH1 | 806.471 | -0.229 | 0.020237326 |
| ENSG00000124541 | RRP36 | 2471.328 | -0.229 | 0.043672977 |
| ENSG00000112167 | SAYSD1 | 965.565 | -0.230 | 0.01617163 |
| ENSG00000134153 | EMC7 | 2477.685 | -0.230 | 0.004407545 |
| ENSG00000168904 | LRRC28 | 642.678 | -0.231 | 0.005401527 |
| ENSG00000128159 | TUBGCP6 | 2564.970 | -0.231 | 0.045066222 |
| ENSG00000173548 | SNX33 | 3054.688 | -0.231 | 0.026322096 |
| ENSG00000132963 | POMP | 5266.449 | -0.232 | 0.019222657 |
| ENSG00000143486 | EIF2D | 2655.336 | -0.232 | 0.014724678 |
| ENSG00000166938 | DIS3L | 1250.758 | -0.232 | 0.008518741 |
| ENSG00000111843 | TMEM14C | 4268.080 | -0.232 | 0.034742454 |
| ENSG00000027847 | B4GALT7 | 1780.116 | -0.232 | 0.021673776 |
| ENSG00000186834 | HEXIM1 | 4215.034 | -0.233 | 0.048635826 |
| ENSG00000100243 | CYB5R3 | 9627.935 | -0.234 | 0.016147285 |
| ENSG00000060491 | OGFR | 3641.115 | -0.234 | 0.040563974 |
| ENSG00000129128 | SPCS3 | 5826.293 | -0.234 | 0.006091081 |
| ENSG00000170634 | ACYP2 | 509.119 | -0.235 | 0.047750914 |
| ENSG00000150991 | UBC | 33038.238 | -0.235 | 0.007602949 |
| ENSG00000147457 | CHMP7 | 2035.844 | -0.235 | 0.011858125 |
| ENSG00000186283 | TOR3A | 3378.477 | -0.235 | 0.009462891 |
| ENSG00000139644 | TMBIM6 | 44677.342 | -0.236 | 0.004210209 |
| ENSG00000180628 | PCGF5 | 1708.840 | -0.236 | 0.040890277 |
| ENSG00000198843 | SELENOT | 4423.901 | -0.236 | 0.001850381 |
| ENSG00000135766 | EGLN1 | 2242.722 | -0.236 | 0.014648539 |
| ENSG00000158805 | ZNF276 | 1124.732 | -0.237 | 0.041715826 |
| ENSG00000142459 | EVI5L | 1255.163 | -0.237 | 0.037352304 |
| ENSG00000013374 | NUB1 | 3706.895 | -0.237 | 0.014693558 |
| ENSG00000033100 | CHPF2 | 4106.670 | -0.238 | 0.020931603 |
| ENSG00000254505 | CHMP4A | 214.348 | -0.238 | 0.044572228 |
| ENSG00000158156 | XKR8 | 575.484 | -0.238 | 0.009767915 |
| ENSG00000198755 | RPL10A | 28219.852 | -0.238 | 0.027169363 |
| ENSG00000134970 | TMED7 | 3429.220 | -0.240 | 0.011703716 |
| ENSG00000164077 | MON1A | 625.178 | -0.240 | 0.024742098 |
| ENSG00000100938 | GMPR2 | 2574.898 | -0.240 | 0.001857327 |
| ENSG00000213047 | DENND1B | 1004.645 | -0.241 | 0.031984112 |
| ENSG00000182405 | PGBD4 | 133.310 | -0.241 | 0.037667144 |
| ENSG00000159346 | ADIPOR1 | 6185.953 | -0.241 | 0.001591221 |
| ENSG00000126247 | CAPNS1 | 21390.809 | -0.241 | 0.015737325 |
| ENSG00000153066 | TXNDC11 | 2671.047 | -0.241 | 0.012457409 |
| ENSG00000152601 | MBNL1 | 5164.190 | -0.242 | 0.017301553 |
| ENSG00000159461 | AMFR | 4507.699 | -0.242 | 0.001682987 |
| ENSG00000189077 | TMEM120A | 2029.609 | -0.242 | 0.035748073 |
| ENSG00000160131 | VMA21 | 2451.629 | -0.242 | 0.004710052 |
| ENSG00000128928 | IVD | 3091.625 | -0.243 | 0.029210744 |
| ENSG00000099308 | MAST3 | 647.180 | -0.243 | 0.046582654 |
| ENSG00000033800 | PIAS1 | 1434.210 | -0.243 | 0.015753203 |
| ENSG00000182359 | KBTBD3 | 179.840 | -0.244 | 0.031558621 |
| ENSG00000145348 | TBCK | 1034.150 | -0.244 | 0.025526459 |
| ENSG00000213625 | LEPROT | 4466.422 | -0.244 | 0.011989169 |
| ENSG00000125868 | DSTN | 18027.823 | -0.244 | 0.036510022 |
| ENSG00000156232 | WHAMM | 916.733 | -0.245 | 0.004410216 |
| ENSG00000115685 | PPP1R7 | 3072.879 | -0.246 | 0.009850209 |
| ENSG00000134899 | ERCC5 | 1540.188 | -0.246 | 0.01617163 |

| | | | | |
|---|---|---|---|---|
| ENSG00000138767 | CNOT6L | 1894.932 | -0.246 | 0.014606939 |
| ENSG00000100865 | CINP | 1343.542 | -0.246 | 0.016175855 |
| ENSG00000049239 | H6PD | 4145.540 | -0.246 | 0.048891653 |
| ENSG00000100983 | GSS | 4457.621 | -0.246 | 0.014432798 |
| ENSG00000081870 | HSPB11 | 1266.281 | -0.247 | 0.038477469 |
| ENSG00000174444 | RPL4 | 74586.336 | -0.247 | 0.045957458 |
| ENSG00000024048 | UBR2 | 2383.344 | -0.247 | 0.001659673 |
| ENSG00000164062 | APEH | 6290.374 | -0.248 | 0.013202693 |
| ENSG00000178149 | DALRD3 | 977.128 | -0.248 | 0.013895789 |
| ENSG00000104774 | MAN2B1 | 4129.438 | -0.248 | 0.016812411 |
| ENSG00000060642 | PIGV | 877.336 | -0.249 | 0.006614887 |
| ENSG00000115520 | COQ10B | 1442.880 | -0.249 | 0.002102002 |
| ENSG00000143801 | PSEN2 | 992.879 | -0.249 | 0.034788515 |
| ENSG00000147364 | FBXO25 | 1159.123 | -0.249 | 0.022588652 |
| ENSG00000124104 | SNX21 | 1007.464 | -0.249 | 0.041569646 |
| ENSG00000109775 | UFSP2 | 785.303 | -0.249 | 0.006240535 |
| ENSG00000108219 | TSPAN14 | 10304.099 | -0.250 | 0.031643876 |
| ENSG00000164010 | ERMAP | 685.421 | -0.250 | 0.025905408 |
| ENSG00000117151 | CTBS | 918.500 | -0.250 | 0.015368163 |
| ENSG00000105135 | ILVBL | 4746.770 | -0.251 | 0.040614179 |
| ENSG00000129636 | ITFG1 | 2329.644 | -0.251 | 0.016634437 |
| ENSG00000177239 | MAN1B1 | 5058.895 | -0.251 | 0.024041583 |
| ENSG00000142541 | RPL13A | 97879.828 | -0.252 | 0.041762168 |
| ENSG00000113595 | TRIM23 | 407.661 | -0.252 | 0.006419659 |
| ENSG00000111727 | HCFC2 | 504.833 | -0.253 | 0.020541448 |
| ENSG00000149084 | HSD17B12 | 2835.347 | -0.253 | 0.016671088 |
| ENSG00000204370 | SDHD | 2737.958 | -0.253 | 0.007053147 |
| ENSG00000117758 | STX12 | 2275.158 | -0.253 | 0.000405934 |
| ENSG00000124222 | STX16 | 3714.584 | -0.253 | 0.007758818 |
| ENSG00000156017 | CARNMT1 | 873.842 | -0.254 | 0.025488061 |
| ENSG00000138777 | PPA2 | 2360.157 | -0.254 | 0.00786281 |
| ENSG00000069966 | GNB5 | 1238.337 | -0.254 | 0.033999133 |
| ENSG00000161558 | TMEM143 | 301.544 | -0.254 | 0.014458993 |
| ENSG00000247796 | | 107.121 | -0.255 | 0.044884028 |
| ENSG00000166136 | NDUFB8 | 1697.395 | -0.255 | 0.043687978 |
| ENSG00000114942 | EEF1B2 | 11914.025 | -0.256 | 0.03785059 |
| ENSG00000108639 | SYNGR2 | 13591.831 | -0.256 | 0.030711112 |
| ENSG00000132906 | CASP9 | 981.671 | -0.256 | 0.016894477 |
| ENSG00000140391 | TSPAN3 | 7380.140 | -0.256 | 0.030751196 |
| ENSG00000198931 | APRT | 6353.760 | -0.256 | 0.049632839 |
| ENSG00000122008 | POLK | 560.566 | -0.256 | 0.014287435 |
| ENSG00000136169 | SETDB2 | 526.332 | -0.257 | 0.025280396 |
| ENSG00000112062 | MAPK14 | 2965.427 | -0.257 | 0.002597817 |
| ENSG00000165476 | REEP3 | 2912.077 | -0.257 | 0.031737153 |
| ENSG00000143622 | RIT1 | 1817.260 | -0.257 | 0.019264689 |
| ENSG00000120696 | KBTBD7 | 468.301 | -0.258 | 0.043417264 |
| ENSG00000215271 | HOMEZ | 1276.577 | -0.258 | 0.045792171 |
| ENSG00000103174 | NAGPA | 799.425 | -0.258 | 0.010452229 |
| ENSG00000013563 | DNASE1L1 | 1099.261 | -0.258 | 0.017032 |
| ENSG00000100418 | DESI1 | 2337.385 | -0.258 | 0.005222747 |
| ENSG00000011198 | ABHD5 | 1125.611 | -0.258 | 0.046624908 |
| ENSG00000073969 | NSF | 2389.030 | -0.258 | 0.020299228 |
| ENSG00000161203 | AP2M1 | 22237.631 | -0.258 | 0.001234109 |
| ENSG00000109323 | MANBA | 2335.498 | -0.259 | 0.046582654 |
| ENSG00000157326 | DHRS4 | 618.069 | -0.259 | 0.031954501 |
| ENSG00000143545 | RAB13 | 5848.946 | -0.259 | 0.016900686 |
| ENSG00000176108 | CHMP6 | 1276.552 | -0.259 | 0.018442854 |
| ENSG00000112679 | DUSP22 | 1411.066 | -0.259 | 0.02669057 |
| ENSG00000198718 | TOGARAM1 | 626.718 | -0.260 | 0.011178676 |

| | | | | |
|---|---|---|---|---|
| ENSG00000131507 | NDFIP1 | 4801.406 | -0.261 | 0.000301131 |
| ENSG00000103066 | PLA2G15 | 1769.283 | -0.261 | 0.041277996 |
| ENSG00000155463 | OXA1L | 4842.300 | -0.261 | 0.001316677 |
| ENSG00000163956 | LRPAP1 | 8249.967 | -0.261 | 0.037857561 |
| ENSG00000124172 | ATP5F1E | 13711.329 | -0.261 | 0.02179224 |
| ENSG00000196072 | BLOC1S2 | 1979.838 | -0.262 | 0.004424445 |
| ENSG00000135185 | TMEM243 | 982.881 | -0.262 | 0.00681772 |
| ENSG00000112078 | KCTD20 | 2910.902 | -0.262 | 0.001795298 |
| ENSG00000145741 | BTF3 | 18527.038 | -0.264 | 0.013129813 |
| ENSG00000176715 | ACSF3 | 1276.726 | -0.264 | 0.009591662 |
| ENSG00000122026 | RPL21 | 11284.672 | -0.264 | 0.034633147 |
| ENSG00000067221 | STOML1 | 784.723 | -0.264 | 0.025199256 |
| ENSG00000175348 | TMEM9B | 2556.263 | -0.265 | 0.000688242 |
| ENSG00000112146 | FBXO9 | 1895.748 | -0.265 | 0.001959934 |
| ENSG00000049245 | VAMP3 | 5201.487 | -0.265 | 0.000513065 |
| ENSG00000124831 | LRRFIP1 | 6300.825 | -0.265 | 0.011131934 |
| ENSG00000100852 | ARHGAP5 | 3133.818 | -0.265 | 0.043142522 |
| ENSG00000165502 | RPL36AL | 7452.385 | -0.265 | 0.0178013 |
| ENSG00000197879 | MYO1C | 10678.396 | -0.266 | 0.010678859 |
| ENSG00000113552 | GNPDA1 | 3230.466 | -0.267 | 0.015845116 |
| ENSG00000075399 | VPS9D1 | 1373.909 | -0.267 | 0.039775819 |
| ENSG00000165572 | KBTBD6 | 625.959 | -0.267 | 0.024209264 |
| ENSG00000050130 | JKAMP | 1519.348 | -0.267 | 0.001321463 |
| ENSG00000110330 | BIRC2 | 2771.853 | -0.268 | 0.043031681 |
| ENSG00000135404 | CD63 | 34493.513 | -0.268 | 0.029500606 |
| ENSG00000155957 | TMBIM4 | 1883.303 | -0.269 | 0.014863408 |
| ENSG00000171503 | ETFDH | 786.968 | -0.269 | 0.005211553 |
| ENSG00000160408 | ST6GALNAC6 | 1826.906 | -0.269 | 0.046914376 |
| ENSG00000100916 | BRMS1L | 402.820 | -0.269 | 0.03284533 |
| ENSG00000126062 | TMEM115 | 2938.207 | -0.270 | 0.004459657 |
| ENSG00000078902 | TOLLIP | 4267.986 | -0.270 | 0.020911563 |
| ENSG00000013441 | CLK1 | 3432.690 | -0.270 | 0.021856822 |
| ENSG00000170791 | CHCHD7 | 1570.752 | -0.271 | 0.013986414 |
| ENSG00000138380 | CARF | 236.837 | -0.271 | 0.040060215 |
| ENSG00000096092 | TMEM14A | 1293.545 | -0.272 | 0.027519035 |
| ENSG00000156050 | FAM161B | 254.014 | -0.272 | 0.038762484 |
| ENSG00000049860 | HEXB | 4930.477 | -0.272 | 0.019895059 |
| ENSG00000213977 | TAX1BP3 | 2311.316 | -0.272 | 0.035267343 |
| ENSG00000136104 | RNASEH2B | 1231.645 | -0.273 | 0.010613122 |
| ENSG00000079805 | DNM2 | 11797.877 | -0.273 | 0.032932265 |
| ENSG00000128951 | DUT | 2746.583 | -0.273 | 0.001593255 |
| ENSG00000083845 | RPS5 | 30644.577 | -0.273 | 0.026755191 |
| ENSG00000137824 | RMDN3 | 2068.835 | -0.273 | 0.000472862 |
| ENSG00000177565 | TBL1XR1 | 8507.159 | -0.274 | 0.020045613 |
| ENSG00000231500 | RPS18 | 73930.385 | -0.274 | 0.036304758 |
| ENSG00000104671 | DCTN6 | 975.477 | -0.274 | 0.003232982 |
| ENSG00000130813 | SHFL | 1560.094 | -0.274 | 0.037277372 |
| ENSG00000051382 | PIK3CB | 1770.748 | -0.275 | 0.025972706 |
| ENSG00000089157 | RPLP0 | 84425.967 | -0.275 | 0.016735537 |
| ENSG00000229809 | ZNF688 | 688.534 | -0.275 | 0.046855907 |
| ENSG00000140988 | RPS2 | 51333.074 | -0.276 | 0.037598669 |
| ENSG00000211451 | GNRHR2 | 55.892 | -0.276 | 0.028600424 |
| ENSG00000197265 | GTF2E2 | 1720.375 | -0.276 | 0.009719309 |
| ENSG00000130313 | PGLS | 3609.123 | -0.276 | 0.039567814 |
| ENSG00000238045 | | 283.157 | -0.276 | 0.020911563 |
| ENSG00000032219 | ARID4A | 846.495 | -0.276 | 0.003966342 |
| ENSG00000196850 | PPTC7 | 1803.366 | -0.276 | 0.015876458 |
| ENSG00000114316 | USP4 | 2636.760 | -0.277 | 0.001516671 |
| ENSG00000180776 | ZDHHC20 | 6493.156 | -0.277 | 0.035748073 |

| | | | | |
|---|---|---|---|---|
| ENSG00000117305 | HMGCL | 2488.781 | -0.277 | 0.006878711 |
| ENSG00000101363 | MANBAL | 3605.668 | -0.277 | 0.000703928 |
| ENSG00000169018 | FEM1B | 3449.957 | -0.278 | 0.003824192 |
| ENSG00000173085 | COQ2 | 577.347 | -0.279 | 0.032749089 |
| ENSG00000204227 | RING1 | 2628.789 | -0.279 | 0.000804847 |
| ENSG00000187609 | EXD3 | 660.553 | -0.279 | 0.047191803 |
| ENSG00000102805 | CLN5 | 582.644 | -0.280 | 0.005611767 |
| ENSG00000281332 | LINC00997 | 180.508 | -0.281 | 0.041283268 |
| ENSG00000240344 | PPIL3 | 662.013 | -0.281 | 0.005761619 |
| ENSG00000231312 | MAP4K3-DT | 327.974 | -0.281 | 0.047895945 |
| ENSG00000236104 | ZBTB22 | 1349.633 | -0.281 | 0.000504626 |
| ENSG00000123983 | ACSL3 | 4342.856 | -0.281 | 0.024424542 |
| ENSG00000166912 | MTMR10 | 907.456 | -0.282 | 0.005772079 |
| ENSG00000075415 | SLC25A3 | 16236.680 | -0.282 | 0.00032847 |
| ENSG00000243477 | NAA80 | 468.884 | -0.282 | 0.021917288 |
| ENSG00000116903 | EXOC8 | 678.167 | -0.282 | 0.00095033 |
| ENSG00000099992 | TBC1D10A | 1303.628 | -0.282 | 0.007224904 |
| ENSG00000198130 | HIBCH | 1403.368 | -0.283 | 0.020565541 |
| ENSG00000104695 | PPP2CB | 2872.117 | -0.283 | 0.004692654 |
| ENSG00000139428 | MMAB | 1488.885 | -0.283 | 0.020931603 |
| ENSG00000270062 | | 27.083 | -0.283 | 0.044943163 |
| ENSG00000109270 | LAMTOR3 | 1783.514 | -0.283 | 0.000238975 |
| ENSG00000147419 | CCDC25 | 1549.446 | -0.284 | 0.002598393 |
| ENSG00000136636 | KCTD3 | 2528.730 | -0.284 | 0.011830543 |
| ENSG00000140280 | LYSMD2 | 452.627 | -0.284 | 0.041114588 |
| ENSG00000067182 | TNFRSF1A | 9487.057 | -0.284 | 0.023454808 |
| ENSG00000120805 | ARL1 | 3939.368 | -0.285 | 0.000489854 |
| ENSG00000142327 | RNPEPL1 | 5271.682 | -0.285 | 0.013659841 |
| ENSG00000075142 | SRI | 5422.871 | -0.287 | 0.012309184 |
| ENSG00000145214 | DGKQ | 1543.045 | -0.287 | 0.010157199 |
| ENSG00000103363 | ELOB | 8104.191 | -0.287 | 0.033665684 |
| ENSG00000054282 | SDCCAG8 | 910.685 | -0.288 | 0.000278303 |
| ENSG00000133639 | BTG1 | 6832.004 | -0.289 | 0.038465763 |
| ENSG00000145779 | TNFAIP8 | 1608.443 | -0.289 | 0.046662842 |
| ENSG00000196588 | MRTFA | 2623.127 | -0.289 | 0.000595985 |
| ENSG00000143641 | GALNT2 | 5899.675 | -0.290 | 0.023855413 |
| ENSG00000114480 | GBE1 | 1327.200 | -0.291 | 0.04135175 |
| ENSG00000125520 | SLC2A4RG | 3854.379 | -0.291 | 0.022302782 |
| ENSG00000163110 | PDLIM5 | 4155.518 | -0.291 | 0.035958027 |
| ENSG00000070778 | PTPN21 | 810.385 | -0.291 | 0.042058886 |
| ENSG00000145685 | LHFPL2 | 2155.254 | -0.292 | 0.034225839 |
| ENSG00000166548 | TK2 | 1303.758 | -0.292 | 0.003521029 |
| ENSG00000104613 | INTS10 | 2163.267 | -0.292 | 0.001895998 |
| ENSG00000102893 | PHKB | 2587.728 | -0.292 | 0.001622521 |
| ENSG00000168938 | PPIC | 2914.422 | -0.292 | 0.042878413 |
| ENSG00000276045 | ORAI1 | 1417.768 | -0.293 | 0.026033498 |
| ENSG00000272040 | | 34.255 | -0.293 | 0.041199065 |
| ENSG00000164323 | CFAP97 | 1424.185 | -0.293 | 0.006766429 |
| ENSG00000005700 | IBTK | 2176.965 | -0.293 | 0.001263527 |
| ENSG00000188846 | RPL14 | 32557.064 | -0.293 | 0.030696877 |
| ENSG00000077150 | NFKB2 | 3758.594 | -0.293 | 0.023404858 |
| ENSG00000143353 | LYPLAL1 | 720.730 | -0.293 | 0.004421652 |
| ENSG00000162542 | TMCO4 | 1923.576 | -0.294 | 0.027644324 |
| ENSG00000125817 | CENPB | 5996.473 | -0.294 | 0.004956352 |
| ENSG00000184903 | IMMP2L | 384.967 | -0.294 | 0.038465763 |
| ENSG00000183864 | TOB2 | 3088.392 | -0.294 | 0.003158032 |
| ENSG00000120725 | SIL1 | 2820.986 | -0.294 | 0.019952107 |
| ENSG00000182544 | MFSD5 | 2724.519 | -0.294 | 0.009559122 |
| ENSG00000118855 | MFSD1 | 3636.695 | -0.294 | 0.007096023 |

| | | | | |
|---|---|---|---|---|
| ENSG00000187630 | DHRS4L2 | 676.064 | -0.295 | 0.031933657 |
| ENSG00000120860 | WASHC3 | 1078.189 | -0.295 | 0.0007487 |
| ENSG00000119655 | NPC2 | 9669.751 | -0.295 | 0.048671716 |
| ENSG00000111229 | ARPC3 | 13905.891 | -0.295 | 0.001403424 |
| ENSG00000161011 | SQSTM1 | 12210.463 | -0.296 | 0.025991337 |
| ENSG00000178951 | ZBTB7A | 3169.961 | -0.296 | 0.00161032 |
| ENSG00000103544 | VPS35L | 2133.952 | -0.297 | 0.025605074 |
| ENSG00000108784 | NAGLU | 2320.326 | -0.297 | 0.007153722 |
| ENSG00000185650 | ZFP36L1 | 19177.384 | -0.297 | 0.032482938 |
| ENSG00000159720 | ATP6V0D1 | 5107.827 | -0.297 | 0.007471925 |
| ENSG00000147439 | BIN3 | 640.645 | -0.297 | 0.000975659 |
| ENSG00000099330 | OCEL1 | 1008.214 | -0.297 | 0.03696188 |
| ENSG00000128294 | TPST2 | 2104.092 | -0.297 | 0.040208825 |
| ENSG00000164331 | ANKRA2 | 637.003 | -0.298 | 0.004712992 |
| ENSG00000100600 | LGMN | 6782.401 | -0.298 | 0.045610387 |
| ENSG00000126903 | SLC10A3 | 2716.615 | -0.299 | 0.00475405 |
| ENSG00000090581 | GNPTG | 2844.386 | -0.299 | 0.001642071 |
| ENSG00000179627 | ZBTB42 | 705.105 | -0.299 | 0.048738494 |
| ENSG00000149577 | SIDT2 | 1730.573 | -0.300 | 0.005421544 |
| ENSG00000197892 | KIF13B | 1735.463 | -0.301 | 0.040614179 |
| ENSG00000197448 | GSTK1 | 7878.548 | -0.301 | 0.009806766 |
| ENSG00000156110 | ADK | 1941.073 | -0.302 | 0.036824146 |
| ENSG00000129158 | SERGEF | 606.312 | -0.302 | 0.017786834 |
| ENSG00000275964 | | 186.216 | -0.302 | 0.035478364 |
| ENSG00000103145 | HCFC1R1 | 2308.112 | -0.303 | 0.034870398 |
| ENSG00000250365 | | 14.679 | -0.303 | 0.030311865 |
| ENSG00000109390 | NDUFC1 | 2059.813 | -0.303 | 0.003270982 |
| ENSG00000092036 | HAUS4 | 893.339 | -0.303 | 0.007669961 |
| ENSG00000177410 | ZFAS1 | 5169.348 | -0.304 | 0.041920206 |
| ENSG00000137720 | C11orf1 | 664.085 | -0.304 | 0.03284533 |
| ENSG00000174917 | MICOS13 | 2561.186 | -0.304 | 0.018685763 |
| ENSG00000197746 | PSAP | 67393.620 | -0.304 | 0.029019774 |
| ENSG00000171100 | MTM1 | 487.482 | -0.304 | 0.008178519 |
| ENSG00000103152 | MPG | 1462.621 | -0.304 | 0.004933013 |
| ENSG00000153113 | CAST | 13037.138 | -0.304 | 0.03370066 |
| ENSG00000234390 | USP27X-DT | 57.950 | -0.305 | 0.023526957 |
| ENSG00000055211 | GINM1 | 2174.169 | -0.305 | 0.000219186 |
| ENSG00000091947 | TMEM101 | 2019.705 | -0.305 | 0.007978 |
| ENSG00000186615 | KTN1-AS1 | 185.334 | -0.305 | 0.049281344 |
| ENSG00000162244 | RPL29 | 31825.120 | -0.306 | 0.035342844 |
| ENSG00000139154 | AEBP2 | 1928.684 | -0.306 | 0.006138921 |
| ENSG00000168028 | RPSA | 38047.423 | -0.307 | 0.017739932 |
| ENSG00000114573 | ATP6V1A | 3633.668 | -0.307 | 0.001125325 |
| ENSG00000027697 | IFNGR1 | 4540.479 | -0.307 | 0.033733646 |
| ENSG00000117691 | NENF | 3324.572 | -0.308 | 0.020448178 |
| ENSG00000213741 | RPS29 | 14619.993 | -0.308 | 0.01451108 |
| ENSG00000109475 | RPL34 | 22653.088 | -0.308 | 0.026258548 |
| ENSG00000116209 | TMEM59 | 12898.767 | -0.308 | 0.000621915 |
| ENSG00000221821 | C6orf226 | 333.966 | -0.308 | 0.048832319 |
| ENSG00000113240 | CLK4 | 683.781 | -0.309 | 0.006564949 |
| ENSG00000155975 | VPS37A | 1303.200 | -0.309 | 0.00096976 |
| ENSG00000102699 | PARP4 | 4500.295 | -0.310 | 0.012927915 |
| ENSG00000154359 | LONRF1 | 843.091 | -0.310 | 0.035327125 |
| ENSG00000156711 | MAPK13 | 6323.400 | -0.310 | 0.021806859 |
| ENSG00000256628 | ZBTB11-AS1 | 161.292 | -0.311 | 0.002550495 |
| ENSG00000137288 | UQCC2 | 2071.543 | -0.311 | 0.009538781 |
| ENSG00000112511 | PHF1 | 2473.174 | -0.312 | 0.000751914 |
| ENSG00000177674 | AGTRAP | 3142.467 | -0.312 | 0.032670347 |
| ENSG00000130724 | CHMP2A | 6604.513 | -0.313 | 0.006505592 |

| | | | | |
|---|---|---|---|---|
| ENSG00000228172 | | 36.502 | -0.313 | 0.010228933 |
| ENSG00000107798 | LIPA | 3290.861 | -0.313 | 0.043894644 |
| ENSG00000006282 | SPATA20 | 3377.257 | -0.313 | 0.033061136 |
| ENSG00000153786 | ZDHHC7 | 3368.044 | -0.313 | 0.001594642 |
| ENSG00000163378 | EOGT | 1475.002 | -0.314 | 0.013652224 |
| ENSG00000136100 | VPS36 | 2116.114 | -0.314 | 0.006632089 |
| ENSG00000131779 | PEX11B | 1655.348 | -0.315 | 7.43203E-05 |
| ENSG00000168056 | LTBP3 | 4687.446 | -0.315 | 0.047468694 |
| ENSG00000060762 | MPC1 | 994.980 | -0.315 | 0.002772167 |
| ENSG00000106785 | TRIM14 | 2093.085 | -0.315 | 0.033060907 |
| ENSG00000114853 | ZBTB47 | 909.955 | -0.315 | 0.00824977 |
| ENSG00000113916 | BCL6 | 3935.284 | -0.315 | 0.032851221 |
| ENSG00000030582 | GRN | 31888.706 | -0.316 | 0.008518741 |
| ENSG00000151151 | IPMK | 595.154 | -0.316 | 0.017514618 |
| ENSG00000121900 | TMEM54 | 5510.398 | -0.316 | 0.040563974 |
| ENSG00000245937 | LINC01184 | 723.847 | -0.316 | 0.000278746 |
| ENSG00000120910 | PPP3CC | 614.147 | -0.317 | 0.006980466 |
| ENSG00000143337 | TOR1AIP1 | 2436.277 | -0.317 | 0.000128355 |
| ENSG00000182372 | CLN8 | 708.573 | -0.317 | 0.004075422 |
| ENSG00000166507 | NDST2 | 103.734 | -0.317 | 0.000253353 |
| ENSG00000163681 | SLMAP | 2859.712 | -0.317 | 0.016718825 |
| ENSG00000151773 | CCDC122 | 176.426 | -0.317 | 0.03741573 |
| ENSG00000179604 | CDC42EP4 | 4211.837 | -0.317 | 0.030180388 |
| ENSG00000130363 | RSPH3 | 789.367 | -0.318 | 0.000902417 |
| ENSG00000140464 | PML | 5633.928 | -0.318 | 0.032543957 |
| ENSG00000089159 | PXN | 7989.714 | -0.318 | 0.015656119 |
| ENSG00000172340 | SUCLG2 | 2556.908 | -0.318 | 0.003059517 |
| ENSG00000159840 | ZYX | 11144.989 | -0.319 | 0.024826701 |
| ENSG00000101986 | ABCD1 | 1145.051 | -0.320 | 0.027056806 |
| ENSG00000114395 | CYB561D2 | 1111.584 | -0.320 | 0.009057359 |
| ENSG00000179476 | C14orf28 | 195.708 | -0.320 | 0.000920138 |
| ENSG00000013288 | MAN2B2 | 3185.843 | -0.320 | 0.001850381 |
| ENSG00000229358 | DPY19L1P1 | 115.103 | -0.321 | 0.033217311 |
| ENSG00000268061 | NAPA-AS1 | 76.690 | -0.321 | 0.019520876 |
| ENSG00000166189 | HPS6 | 1215.583 | -0.321 | 0.000130963 |
| ENSG00000243646 | IL10RB | 2192.147 | -0.321 | 0.00291428 |
| ENSG00000124201 | ZNFX1 | 3729.628 | -0.322 | 0.008577592 |
| ENSG00000100284 | TOM1 | 2850.913 | -0.322 | 0.005637163 |
| ENSG00000163820 | FYCO1 | 2269.729 | -0.322 | 0.012653755 |
| ENSG00000150347 | ARID5B | 2875.584 | -0.323 | 0.035795883 |
| ENSG00000149925 | ALDOA | 81808.159 | -0.323 | 0.012529782 |
| ENSG00000135828 | RNASEL | 796.440 | -0.323 | 0.005528943 |
| ENSG00000134864 | GGACT | 239.294 | -0.323 | 0.028272296 |
| ENSG00000203485 | INF2 | 7436.576 | -0.323 | 0.014602315 |
| ENSG00000008282 | SYPL1 | 9367.588 | -0.324 | 0.00644444 |
| ENSG00000126822 | PLEKHG3 | 3053.993 | -0.324 | 0.04123665 |
| ENSG00000197930 | ERO1A | 7674.949 | -0.324 | 0.03991284 |
| ENSG00000125753 | VASP | 8652.615 | -0.325 | 0.004712992 |
| ENSG00000167074 | TEF | 1135.983 | -0.325 | 0.012637953 |
| ENSG00000100439 | ABHD4 | 2281.622 | -0.326 | 0.022605204 |
| ENSG00000221914 | PPP2R2A | 2504.457 | -0.326 | 0.001167447 |
| ENSG00000100429 | HDAC10 | 720.656 | -0.327 | 0.035267343 |
| ENSG00000132825 | PPP1R3D | 708.063 | -0.327 | 0.003469733 |
| ENSG00000103876 | FAH | 1466.135 | -0.327 | 0.028734724 |
| ENSG00000122873 | CISD1 | 996.737 | -0.328 | 0.006669946 |
| ENSG00000138119 | MYOF | 11180.977 | -0.328 | 0.033806005 |
| ENSG00000272760 | | 94.595 | -0.329 | 0.015836763 |
| ENSG00000120519 | SLC10A7 | 468.478 | -0.329 | 0.001146966 |
| ENSG00000181523 | SGSH | 2226.209 | -0.329 | 0.003498539 |

| | | | | |
|---|---|---|---|---|
| ENSG00000196776 | CD47 | 6107.389 | -0.329 | 0.049082303 |
| ENSG00000006025 | OSBPL7 | 953.268 | -0.329 | 0.028602406 |
| ENSG00000164031 | DNAJB14 | 1468.610 | -0.331 | 8.67395E-05 |
| ENSG00000155252 | PI4K2A | 2104.711 | -0.332 | 0.002463246 |
| ENSG00000259976 | | 377.792 | -0.332 | 0.043714542 |
| ENSG00000179981 | TSHZ1 | 945.546 | -0.332 | 0.018395633 |
| ENSG00000257433 | | 41.703 | -0.333 | 0.032529868 |
| ENSG00000183617 | MRPL54 | 1637.567 | -0.333 | 0.015443268 |
| ENSG00000171314 | PGAM1 | 2486.139 | -0.334 | 0.020299228 |
| ENSG00000186010 | NDUFA13 | 2361.703 | -0.334 | 0.034611744 |
| ENSG00000220848 | RPS18P9 | 72.277 | -0.335 | 0.003556861 |
| ENSG00000121310 | ECHDC2 | 2964.311 | -0.335 | 0.029019774 |
| ENSG00000271797 | | 10.667 | -0.335 | 0.023821602 |
| ENSG00000116717 | GADD45A | 2571.370 | -0.335 | 0.035201594 |
| ENSG00000172831 | CES2 | 2170.133 | -0.336 | 0.009482405 |
| ENSG00000164117 | FBXO8 | 821.541 | -0.336 | 0.00102787 |
| ENSG00000233554 | B4GALT1-AS1 | 32.600 | -0.336 | 0.046292208 |
| ENSG00000158092 | NCK1 | 1265.364 | -0.337 | 0.01935453 |
| ENSG00000107566 | ERLIN1 | 2682.528 | -0.338 | 0.000293774 |
| ENSG00000270959 | LPP-AS2 | 143.942 | -0.338 | 0.03064687 |
| ENSG00000139160 | ETFBKMT | 132.825 | -0.338 | 0.000797536 |
| ENSG00000146859 | TMEM140 | 910.771 | -0.338 | 0.018860521 |
| ENSG00000100147 | CCDC134 | 190.721 | -0.339 | 0.016147264 |
| ENSG00000177106 | EPS8L2 | 7545.270 | -0.339 | 0.042574046 |
| ENSG00000261737 | CLCA4-AS1 | 46.950 | -0.339 | 0.011658528 |
| ENSG00000006757 | PNPLA4 | 641.662 | -0.340 | 0.013840961 |
| ENSG00000105643 | ARRDC2 | 1686.229 | -0.340 | 0.005614053 |
| ENSG00000168297 | PXK | 608.189 | -0.340 | 0.009767915 |
| ENSG00000241852 | C8orf58 | 455.951 | -0.340 | 0.006981518 |
| ENSG00000168310 | IRF2 | 2099.657 | -0.341 | 8.38015E-05 |
| ENSG00000100994 | PYGB | 8430.093 | -0.341 | 0.03706252 |
| ENSG00000274425 | | 49.495 | -0.342 | 0.00333962 |
| ENSG00000205593 | DENND6B | 562.497 | -0.342 | 0.01706104 |
| ENSG00000126790 | L3HYPDH | 347.780 | -0.342 | 0.018258314 |
| ENSG00000253540 | FAM86HP | 75.073 | -0.342 | 0.042711013 |
| ENSG00000116663 | FBXO6 | 1144.370 | -0.342 | 0.020817366 |
| ENSG00000135535 | CD164 | 13084.300 | -0.343 | 0.000306323 |
| ENSG00000198736 | MSRB1 | 2316.970 | -0.344 | 0.018660851 |
| ENSG00000156642 | NPTN | 4601.135 | -0.344 | 0.000341451 |
| ENSG00000173193 | PARP14 | 5910.732 | -0.344 | 0.030696877 |
| ENSG00000120318 | ARAP3 | 1886.621 | -0.344 | 0.03567989 |
| ENSG00000139974 | SLC38A6 | 392.537 | -0.344 | 0.001924121 |
| ENSG00000177738 | | 217.278 | -0.344 | 0.021011475 |
| ENSG00000205730 | ITPRIPL2 | 3223.842 | -0.345 | 0.031751576 |
| ENSG00000087086 | FTL | 131030.468 | -0.345 | 0.03978039 |
| ENSG00000197217 | ENTPD4 | 2243.053 | -0.345 | 0.001001184 |
| ENSG00000129187 | DCTD | 3517.575 | -0.346 | 1.10908E-05 |
| ENSG00000105373 | NOP53 | 15877.263 | -0.346 | 0.0091663 |
| ENSG00000174132 | FAM174A | 870.504 | -0.346 | 0.00082597 |
| ENSG00000173013 | CCDC96 | 142.152 | -0.346 | 0.010753419 |
| ENSG00000117676 | RPS6KA1 | 3170.278 | -0.347 | 0.006519324 |
| ENSG00000138162 | TACC2 | 2499.681 | -0.347 | 0.042451634 |
| ENSG00000248487 | ABHD14A | 835.219 | -0.347 | 0.04527157 |
| ENSG00000108091 | CCDC6 | 3640.099 | -0.347 | 0.000168351 |
| ENSG00000265666 | RARA-AS1 | 138.881 | -0.348 | 0.018874727 |
| ENSG00000104881 | PPP1R13L | 6248.117 | -0.348 | 0.037064237 |
| ENSG00000142634 | EFHD2 | 12206.432 | -0.348 | 0.008123931 |
| ENSG00000104228 | TRIM35 | 816.713 | -0.348 | 0.001183754 |
| ENSG00000204054 | LINC00963 | 2010.464 | -0.348 | 0.003475693 |

| | | | | |
|---|---|---|---|---|
| ENSG00000227946 | | 113.671 | -0.348 | 0.004933013 |
| ENSG00000084764 | MAPRE3 | 917.784 | -0.348 | 0.036876387 |
| ENSG00000124496 | TRERF1 | 1582.840 | -0.348 | 0.035593224 |
| ENSG00000196576 | PLXNB2 | 22935.354 | -0.348 | 0.001152363 |
| ENSG00000110628 | SLC22A18 | 948.781 | -0.349 | 0.034829476 |
| ENSG00000126522 | ASL | 1510.364 | -0.349 | 0.001963148 |
| ENSG00000149932 | TMEM219 | 5995.063 | -0.349 | 0.00441843 |
| ENSG00000177200 | CHD9 | 1680.267 | -0.349 | 0.002025387 |
| ENSG00000183401 | CCDC159 | 652.094 | -0.349 | 0.028241493 |
| ENSG00000197343 | ZNF655 | 2629.321 | -0.349 | 0.006920638 |
| ENSG00000127419 | TMEM175 | 1145.385 | -0.350 | 0.002803854 |
| ENSG00000143669 | LYST | 1291.704 | -0.350 | 0.018442854 |
| ENSG00000188811 | NHLRC3 | 1342.653 | -0.350 | 0.000641236 |
| ENSG00000176473 | WDR25 | 449.660 | -0.350 | 0.000367282 |
| ENSG00000135722 | FBXL8 | 347.028 | -0.351 | 0.009957885 |
| ENSG00000205133 | TRIQK | 1101.676 | -0.351 | 0.007711048 |
| ENSG00000138814 | PPP3CA | 2715.070 | -0.351 | 0.002395898 |
| ENSG00000066027 | PPP2R5A | 2290.468 | -0.351 | 4.48796E-05 |
| ENSG00000067066 | SP100 | 3036.254 | -0.352 | 0.010499191 |
| ENSG00000183172 | SMDT1 | 1807.206 | -0.353 | 0.00345406 |
| ENSG00000178226 | PRSS36 | 118.335 | -0.353 | 0.0413791 |
| ENSG00000178035 | IMPDH2 | 9646.499 | -0.353 | 0.002412375 |
| ENSG00000232533 | | 455.010 | -0.354 | 0.016073021 |
| ENSG00000143776 | CDC42BPA | 2149.183 | -0.355 | 0.014230056 |
| ENSG00000109572 | CLCN3 | 4355.033 | -0.355 | 0.008577592 |
| ENSG00000100300 | TSPO | 10537.410 | -0.355 | 0.032039175 |
| ENSG00000144579 | CTDSP1 | 6568.234 | -0.355 | 0.000145544 |
| ENSG00000143774 | GUK1 | 9768.579 | -0.356 | 0.000830816 |
| ENSG00000267317 | | 233.205 | -0.356 | 0.012968711 |
| ENSG00000159761 | C16orf86 | 100.243 | -0.356 | 0.041346535 |
| ENSG00000169964 | TMEM42 | 655.189 | -0.356 | 0.000196039 |
| ENSG00000037042 | TUBG2 | 1187.623 | -0.356 | 0.020015263 |
| ENSG00000104765 | BNIP3L | 4079.433 | -0.357 | 0.003437794 |
| ENSG00000120889 | TNFRSF10B | 3306.058 | -0.357 | 0.014078052 |
| ENSG00000232940 | HCG25 | 25.425 | -0.358 | 0.004664971 |
| ENSG00000065621 | GSTO2 | 1247.862 | -0.358 | 0.041824082 |
| ENSG00000257704 | INAFM1 | 781.734 | -0.358 | 0.010100583 |
| ENSG00000123989 | CHPF | 8171.626 | -0.358 | 0.030614379 |
| ENSG00000178209 | PLEC | 30682.465 | -0.358 | 0.02669057 |
| ENSG00000101782 | RIOK3 | 4033.114 | -0.359 | 0.002544372 |
| ENSG00000158863 | FAM160B2 | 3041.905 | -0.359 | 0.000670535 |
| ENSG00000224536 | | 20.870 | -0.360 | 0.039654592 |
| ENSG00000168556 | ING2 | 478.042 | -0.360 | 0.000472862 |
| ENSG00000135776 | ABCB10 | 917.067 | -0.360 | 0.000234132 |
| ENSG00000171603 | CLSTN1 | 13425.145 | -0.361 | 0.006066923 |
| ENSG00000139684 | ESD | 2981.329 | -0.361 | 0.001492698 |
| ENSG00000148180 | GSN | 19337.709 | -0.362 | 0.047069292 |
| ENSG00000092010 | PSME1 | 11165.336 | -0.362 | 0.001107743 |
| ENSG00000162413 | KLHL21 | 2499.572 | -0.362 | 0.0091663 |
| ENSG00000117266 | CDK18 | 1697.468 | -0.363 | 0.049384755 |
| ENSG00000161677 | JOSD2 | 1715.014 | -0.363 | 0.010746203 |
| ENSG00000111671 | SPSB2 | 687.637 | -0.364 | 0.011795928 |
| ENSG00000255031 | | 94.894 | -0.364 | 0.046949903 |
| ENSG00000104756 | KCTD9 | 1007.880 | -0.364 | 0.006138921 |
| ENSG00000186577 | SMIM29 | 1378.088 | -0.364 | 0.001826912 |
| ENSG00000100711 | ZFYVE21 | 2419.474 | -0.365 | 8.31771E-06 |
| ENSG00000112893 | MAN2A1 | 2273.704 | -0.365 | 0.001010923 |
| ENSG00000119711 | ALDH6A1 | 979.335 | -0.366 | 0.004282946 |
| ENSG00000132294 | EFR3A | 3136.259 | -0.366 | 0.000654797 |

| | | | | |
|---|---|---|---|---|
| ENSG00000146416 | AIG1 | 2747.920 | -0.366 | 0.015386846 |
| ENSG00000166173 | LARP6 | 611.179 | -0.366 | 0.043091157 |
| ENSG00000237489 | C10orf143 | 55.970 | -0.366 | 0.035222642 |
| ENSG00000172465 | TCEAL1 | 867.793 | -0.366 | 0.005323567 |
| ENSG00000182057 | OGFRP1 | 27.293 | -0.367 | 0.033150407 |
| ENSG00000135205 | CCDC146 | 298.122 | -0.367 | 0.048240426 |
| ENSG00000168936 | TMEM129 | 3101.917 | -0.368 | 0.003148989 |
| ENSG00000131171 | SH3BGRL | 4033.418 | -0.369 | 0.014260864 |
| ENSG00000215440 | NPEPL1 | 1064.215 | -0.369 | 0.011109832 |
| ENSG00000182165 | TP53TG1 | 1533.027 | -0.370 | 0.026237352 |
| ENSG00000109320 | NFKB1 | 2916.243 | -0.370 | 0.00057 |
| ENSG00000103335 | PIEZO1 | 7655.765 | -0.371 | 0.00853114 |
| ENSG00000145349 | CAMK2D | 1768.797 | -0.371 | 0.012773793 |
| ENSG00000189067 | LITAF | 7745.706 | -0.371 | 0.009192908 |
| ENSG00000226200 | SGMS1-AS1 | 142.348 | -0.371 | 0.015364123 |
| ENSG00000102871 | TRADD | 1847.936 | -0.371 | 0.001321727 |
| ENSG00000277369 | | 32.611 | -0.371 | 0.046511376 |
| ENSG00000067225 | PKM | 83755.200 | -0.371 | 0.008119717 |
| ENSG00000100106 | TRIOBP | 2157.764 | -0.372 | 0.001535807 |
| ENSG00000114023 | FAM162A | 4674.114 | -0.372 | 0.026241837 |
| ENSG00000140264 | SERF2 | 25699.825 | -0.372 | 0.001673567 |
| ENSG00000112699 | GMDS | 1519.047 | -0.374 | 0.001593255 |
| ENSG00000270194 | GOLGA4-AS1 | 65.979 | -0.375 | 0.004091537 |
| ENSG00000261338 | | 126.512 | -0.375 | 0.036884139 |
| ENSG00000088808 | PPP1R13B | 1587.610 | -0.375 | 0.001019378 |
| ENSG00000258655 | ARHGAP5-AS1 | 178.801 | -0.376 | 0.015108502 |
| ENSG00000244538 | | 25.615 | -0.376 | 0.010156393 |
| ENSG00000261423 | TMEM202-AS1 | 31.534 | -0.377 | 0.002049444 |
| ENSG00000131981 | LGALS3 | 10889.751 | -0.377 | 0.023155725 |
| ENSG00000103227 | LMF1 | 634.691 | -0.377 | 0.027266908 |
| ENSG00000107331 | ABCA2 | 2019.119 | -0.379 | 0.036030947 |
| ENSG00000142733 | MAP3K6 | 2570.350 | -0.379 | 0.008291843 |
| ENSG00000176124 | DLEU1 | 277.521 | -0.380 | 0.024863684 |
| ENSG00000090006 | LTBP4 | 3545.095 | -0.380 | 0.047795356 |
| ENSG00000001461 | NIPAL3 | 1723.690 | -0.380 | 0.000732666 |
| ENSG00000144746 | ARL6IP5 | 5676.929 | -0.381 | 0.000566291 |
| ENSG00000110921 | MVK | 1875.760 | -0.381 | 0.006628509 |
| ENSG00000228084 | | 29.188 | -0.381 | 0.004933013 |
| ENSG00000230551 | | 935.383 | -0.381 | 0.019005041 |
| ENSG00000130304 | SLC27A1 | 1082.653 | -0.381 | 0.003031029 |
| ENSG00000178449 | COX14 | 2093.731 | -0.382 | 0.000939153 |
| ENSG00000115657 | ABCB6 | 338.742 | -0.383 | 0.037352304 |
| ENSG00000138744 | NAAA | 927.859 | -0.384 | 0.008472038 |
| ENSG00000154889 | MPPE1 | 606.757 | -0.384 | 0.000287187 |
| ENSG00000170088 | TMEM192 | 1078.740 | -0.384 | 5.04104E-05 |
| ENSG00000069943 | PIGB | 469.967 | -0.385 | 6.9165E-06 |
| ENSG00000233276 | GPX1 | 13961.151 | -0.385 | 0.005721701 |
| ENSG00000162777 | DENND2D | 3239.683 | -0.386 | 0.028680813 |
| ENSG00000268364 | SMC5-DT | 26.774 | -0.386 | 0.012977609 |
| ENSG00000123178 | SPRYD7 | 696.723 | -0.387 | 0.00118519 |
| ENSG00000216866 | RPS2P55 | 33.494 | -0.387 | 0.042286363 |
| ENSG00000151718 | WWC2 | 602.478 | -0.387 | 0.031160105 |
| ENSG00000115963 | RND3 | 3689.818 | -0.387 | 0.048228204 |
| ENSG00000213757 | RPS3AP20 | 97.330 | -0.388 | 0.030275633 |
| ENSG00000122884 | P4HA1 | 3902.327 | -0.388 | 0.011196612 |
| ENSG00000123096 | SSPN | 896.796 | -0.389 | 0.041394591 |
| ENSG00000236439 | | 157.498 | -0.389 | 0.036570074 |
| ENSG00000102796 | DHRS12 | 455.256 | -0.389 | 0.002406417 |
| ENSG00000177697 | CD151 | 15692.649 | -0.389 | 0.002153487 |

| | | | | |
|---|---|---|---|---|
| ENSG00000050405 | LIMA1 | 7477.299 | -0.389 | 0.025621239 |
| ENSG00000231747 | | 87.958 | -0.389 | 0.044106066 |
| ENSG00000102189 | EEA1 | 1750.298 | -0.389 | 0.002213298 |
| ENSG00000114529 | C3orf52 | 956.667 | -0.390 | 0.041045779 |
| ENSG00000225131 | PSME2P2 | 70.998 | -0.391 | 0.017517283 |
| ENSG00000113441 | LNPEP | 2940.319 | -0.391 | 0.000115784 |
| ENSG00000197019 | SERTAD1 | 1485.473 | -0.392 | 0.004083282 |
| ENSG00000010704 | HFE | 841.743 | -0.392 | 0.019302235 |
| ENSG00000173821 | RNF213 | 14073.178 | -0.393 | 0.018971266 |
| ENSG00000075234 | TTC38 | 1056.051 | -0.393 | 0.000789431 |
| ENSG00000106211 | HSPB1 | 70187.193 | -0.393 | 0.046220498 |
| ENSG00000171428 | NAT1 | 240.621 | -0.393 | 0.012259353 |
| ENSG00000089060 | SLC8B1 | 2518.932 | -0.393 | 0.00044881 |
| ENSG00000100288 | CHKB | 303.161 | -0.394 | 0.01113974 |
| ENSG00000150977 | RILPL2 | 1121.316 | -0.394 | 0.001062769 |
| ENSG00000151876 | FBXO4 | 558.097 | -0.395 | 0.001149257 |
| ENSG00000162616 | DNAJB4 | 765.180 | -0.395 | 0.011843009 |
| ENSG00000268996 | MAN1B1-DT | 97.356 | -0.395 | 0.016356795 |
| ENSG00000145730 | PAM | 3866.565 | -0.396 | 0.030460545 |
| ENSG00000198018 | ENTPD7 | 1063.691 | -0.396 | 0.027651902 |
| ENSG00000275601 | | 15.028 | -0.396 | 0.043848429 |
| ENSG00000280383 | | 47.428 | -0.396 | 0.02616888 |
| ENSG00000101439 | CST3 | 17709.168 | -0.397 | 0.009790221 |
| ENSG00000243989 | ACY1 | 364.249 | -0.397 | 0.014738911 |
| ENSG00000105339 | DENND3 | 870.289 | -0.397 | 0.011229221 |
| ENSG00000156381 | ANKRD9 | 1483.958 | -0.397 | 0.005323567 |
| ENSG00000183828 | NUDT14 | 1048.573 | -0.398 | 0.012623458 |
| ENSG00000163931 | TKT | 20442.340 | -0.398 | 0.007230564 |
| ENSG00000117791 | MTARC2 | 423.999 | -0.399 | 0.007659955 |
| ENSG00000077238 | IL4R | 5710.913 | -0.399 | 0.01069268 |
| ENSG00000163659 | TIPARP | 2060.938 | -0.399 | 0.023855053 |
| ENSG00000107902 | LHPP | 1135.580 | -0.399 | 0.004295218 |
| ENSG00000278396 | | 26.662 | -0.400 | 0.019222657 |
| ENSG00000043143 | JADE2 | 1892.592 | -0.401 | 0.025704916 |
| ENSG00000000938 | FGR | 507.048 | -0.401 | 0.049391081 |
| ENSG00000196923 | PDLIM7 | 4023.067 | -0.401 | 0.04723942 |
| ENSG00000168040 | FADD | 672.359 | -0.401 | 0.005222747 |
| ENSG00000059122 | FLYWCH1 | 1907.966 | -0.402 | 0.000171222 |
| ENSG00000257261 | | 70.262 | -0.402 | 0.026489141 |
| ENSG00000217027 | TPT1P4 | 13.939 | -0.403 | 0.042076302 |
| ENSG00000088970 | KIZ | 1048.164 | -0.403 | 0.010067738 |
| ENSG00000075643 | MOCOS | 1255.060 | -0.403 | 0.00859921 |
| ENSG00000120899 | PTK2B | 2059.350 | -0.404 | 0.012687026 |
| ENSG00000171604 | CXXC5 | 2505.057 | -0.404 | 0.036243248 |
| ENSG00000178977 | LINC00324 | 162.761 | -0.404 | 0.008538526 |
| ENSG00000131069 | ACSS2 | 2210.105 | -0.404 | 0.000547369 |
| ENSG00000100307 | CBX7 | 1108.459 | -0.404 | 0.011911737 |
| ENSG00000114473 | IQCG | 491.558 | -0.405 | 0.008885698 |
| ENSG00000239763 | | 23.612 | -0.405 | 0.048879164 |
| ENSG00000177432 | NAP1L5 | 231.945 | -0.405 | 0.019460743 |
| ENSG00000163517 | HDAC11 | 1281.729 | -0.405 | 0.013050179 |
| ENSG00000108771 | DHX58 | 1240.895 | -0.406 | 0.005608831 |
| ENSG00000172728 | FUT10 | 232.906 | -0.406 | 0.000542055 |
| ENSG00000196187 | TMEM63A | 3587.879 | -0.407 | 0.004621803 |
| ENSG00000090238 | YPEL3 | 4044.188 | -0.407 | 0.02295239 |
| ENSG00000182718 | ANXA2 | 50136.028 | -0.408 | 0.029685749 |
| ENSG00000159228 | CBR1 | 4072.206 | -0.408 | 0.027651902 |
| ENSG00000253982 | | 331.187 | -0.408 | 0.001497041 |
| ENSG00000158711 | ELK4 | 2916.176 | -0.408 | 2.92625E-05 |

| | | | | |
|---|---|---|---|---|
| ENSG00000123179 | EBPL | 1721.100 | -0.409 | 0.007978 |
| ENSG00000181619 | GPR135 | 60.550 | -0.409 | 0.038972438 |
| ENSG00000176978 | DPP7 | 6120.933 | -0.410 | 0.002884727 |
| ENSG00000259330 | INAFM2 | 516.252 | -0.412 | 0.001568565 |
| ENSG00000272419 | LINC01145 | 121.375 | -0.413 | 0.008518741 |
| ENSG00000205559 | CHKB-DT | 44.433 | -0.413 | 0.014352761 |
| ENSG00000241014 | GPR199P | 108.589 | -0.413 | 0.001146251 |
| ENSG00000107829 | FBXW4 | 2902.237 | -0.413 | 2.54301E-05 |
| ENSG00000107872 | FBXL15 | 795.906 | -0.413 | 0.003166817 |
| ENSG00000180758 | GPR157 | 1655.861 | -0.413 | 0.041445529 |
| ENSG00000228903 | RASA4CP | 106.847 | -0.413 | 0.024665302 |
| ENSG00000125246 | CLYBL | 240.158 | -0.414 | 0.018819144 |
| ENSG00000239704 | CDRT4 | 34.534 | -0.414 | 0.042113294 |
| ENSG00000090554 | FLT3LG | 136.240 | -0.414 | 0.007922515 |
| ENSG00000107438 | PDLIM1 | 12917.339 | -0.415 | 0.006552822 |
| ENSG00000107960 | STN1 | 1328.469 | -0.415 | 0.000576003 |
| ENSG00000156521 | TYSND1 | 2048.067 | -0.416 | 0.000335951 |
| ENSG00000203883 | SOX18 | 427.842 | -0.416 | 0.03849707 |
| ENSG00000173928 | SWSAP1 | 251.765 | -0.416 | 0.00085059 |
| ENSG00000254473 | | 59.437 | -0.417 | 0.001568565 |
| ENSG00000171298 | GAA | 8340.913 | -0.417 | 0.010946144 |
| ENSG00000100299 | ARSA | 2492.313 | -0.418 | 0.000957318 |
| ENSG00000140398 | NEIL1 | 950.002 | -0.418 | 0.037878192 |
| ENSG00000173295 | FAM86B3P | 137.226 | -0.419 | 0.030415237 |
| ENSG00000258561 | | 8.467 | -0.419 | 0.047446928 |
| ENSG00000105518 | TMEM205 | 3515.528 | -0.419 | 0.001974571 |
| ENSG00000281005 | LINC00921 | 48.373 | -0.420 | 0.002598393 |
| ENSG00000104689 | TNFRSF10A | 649.995 | -0.420 | 0.012591553 |
| ENSG00000272221 | | 75.586 | -0.420 | 0.048228204 |
| ENSG00000185641 | | 378.476 | -0.421 | 0.017757112 |
| ENSG00000121316 | PLBD1 | 4784.567 | -0.421 | 0.031237187 |
| ENSG00000090020 | SLC9A1 | 5579.662 | -0.422 | 0.00123897 |
| ENSG00000182700 | IGIP | 416.437 | -0.422 | 5.71004E-05 |
| ENSG00000276931 | | 12.526 | -0.422 | 0.038457463 |
| ENSG00000213614 | HEXA | 3647.261 | -0.422 | 1.43897E-05 |
| ENSG00000002549 | LAP3 | 4878.112 | -0.423 | 0.02037589 |
| ENSG00000173281 | PPP1R3B | 1626.836 | -0.423 | 0.016237632 |
| ENSG00000164124 | TMEM144 | 477.434 | -0.423 | 0.001078614 |
| ENSG00000149489 | ROM1 | 182.512 | -0.424 | 0.002813967 |
| ENSG00000108679 | LGALS3BP | 42095.054 | -0.424 | 0.009357058 |
| ENSG00000279982 | | 7.644 | -0.424 | 0.032670347 |
| ENSG00000151726 | ACSL1 | 5582.067 | -0.424 | 0.028115137 |
| ENSG00000186352 | ANKRD37 | 313.001 | -0.424 | 0.029290812 |
| ENSG00000073331 | ALPK1 | 1010.751 | -0.426 | 0.000212941 |
| ENSG00000126821 | SGPP1 | 973.819 | -0.427 | 0.046390229 |
| ENSG00000196605 | ZNF846 | 466.912 | -0.428 | 0.005591059 |
| ENSG00000111801 | BTN3A3 | 1191.012 | -0.428 | 0.020328597 |
| ENSG00000129625 | REEP5 | 7795.749 | -0.428 | 6.88245E-08 |
| ENSG00000109686 | SH3D19 | 2057.982 | -0.429 | 0.000308538 |
| ENSG00000180389 | ATP5F1EP2 | 25.469 | -0.429 | 0.026237352 |
| ENSG00000104691 | UBXN8 | 366.057 | -0.429 | 0.000405934 |
| ENSG00000145103 | ILDR1 | 547.778 | -0.430 | 0.047380173 |
| ENSG00000060140 | STYK1 | 595.972 | -0.430 | 0.020288796 |
| ENSG00000186470 | BTN3A2 | 2327.388 | -0.431 | 0.013445846 |
| ENSG00000220793 | RPL21P119 | 28.327 | -0.431 | 0.019434635 |
| ENSG00000167895 | TMC8 | 757.735 | -0.431 | 0.049923562 |
| ENSG00000186594 | MIR22HG | 973.598 | -0.432 | 0.008592302 |
| ENSG00000164023 | SGMS2 | 925.309 | -0.432 | 0.037381128 |
| ENSG00000272768 | | 27.373 | -0.432 | 0.014261774 |

| | | | | |
|---|---|---|---|---|
| ENSG00000161642 | ZNF385A | 6518.805 | -0.433 | 0.036617736 |
| ENSG00000159166 | LAD1 | 16236.869 | -0.433 | 0.041854681 |
| ENSG00000213700 | RPL17P50 | 56.726 | -0.433 | 0.013449747 |
| ENSG00000148120 | AOPEP | 1897.000 | -0.433 | 0.007385917 |
| ENSG00000145743 | FBXL17 | 964.999 | -0.434 | 3.02127E-06 |
| ENSG00000007384 | RHBDF1 | 1959.122 | -0.434 | 0.001005416 |
| ENSG00000155254 | MARVELD1 | 2884.708 | -0.434 | 0.027478866 |
| ENSG00000181227 | DLSTP1 | 14.510 | -0.434 | 0.041450446 |
| ENSG00000226360 | RPL10AP6 | 103.838 | -0.435 | 0.007537606 |
| ENSG00000162599 | NFIA | 1541.040 | -0.435 | 0.018066213 |
| ENSG00000168918 | INPP5D | 1857.862 | -0.435 | 0.030257495 |
| ENSG00000243749 | TMEM35B | 226.932 | -0.435 | 0.000109944 |
| ENSG00000128833 | MYO5C | 1014.515 | -0.435 | 0.04569585 |
| ENSG00000030110 | BAK1 | 2207.103 | -0.436 | 0.002513718 |
| ENSG00000170275 | CRTAP | 6820.733 | -0.437 | 2.23606E-05 |
| ENSG00000064687 | ABCA7 | 1388.404 | -0.437 | 0.010350727 |
| ENSG00000103490 | PYCARD | 2003.096 | -0.437 | 0.022605204 |
| ENSG00000198336 | MYL4 | 14.015 | -0.437 | 0.044943163 |
| ENSG00000157514 | TSC22D3 | 4826.716 | -0.437 | 0.023478343 |
| ENSG00000103852 | TTC23 | 729.224 | -0.438 | 1.56098E-05 |
| ENSG00000050327 | ARHGEF5 | 1075.199 | -0.438 | 0.00448427 |
| ENSG00000173926 | MARCHF3 | 206.777 | -0.438 | 0.026156355 |
| ENSG00000271424 | GLRX5P3 | 7.115 | -0.438 | 0.03230253 |
| ENSG00000137496 | IL18BP | 746.784 | -0.438 | 0.018185479 |
| ENSG00000260916 | CCPG1 | 567.224 | -0.439 | 0.000433337 |
| ENSG00000186951 | PPARA | 991.396 | -0.439 | 0.001234109 |
| ENSG00000266208 | | 940.887 | -0.440 | 0.008149836 |
| ENSG00000134955 | SLC37A2 | 1950.587 | -0.440 | 0.046952732 |
| ENSG00000138448 | ITGAV | 5868.361 | -0.440 | 0.003993853 |
| ENSG00000270426 | | 23.644 | -0.440 | 0.011989059 |
| ENSG00000060656 | PTPRU | 5306.184 | -0.440 | 0.024746495 |
| ENSG00000107819 | SFXN3 | 2111.470 | -0.441 | 0.001195582 |
| ENSG00000116977 | LGALS8 | 2999.493 | -0.441 | 6.12049E-06 |
| ENSG00000197324 | LRP10 | 9879.952 | -0.442 | 5.47519E-05 |
| ENSG00000234782 | TPT1P9 | 16.993 | -0.442 | 0.039461872 |
| ENSG00000261794 | GOLGA8H | 6.714 | -0.442 | 0.042771415 |
| ENSG00000267185 | PTP4A2P1 | 12.697 | -0.443 | 0.015347646 |
| ENSG00000139192 | TAPBPL | 1662.983 | -0.443 | 0.005206102 |
| ENSG00000214530 | STARD10 | 4240.643 | -0.444 | 0.016750721 |
| ENSG00000132326 | PER2 | 791.356 | -0.444 | 0.001797674 |
| ENSG00000266094 | RASSF5 | 1858.146 | -0.444 | 0.003512636 |
| ENSG00000150527 | MIA2 | 180.049 | -0.444 | 4.01372E-05 |
| ENSG00000149573 | MPZL2 | 9805.048 | -0.444 | 0.023404858 |
| ENSG00000260461 | | 145.493 | -0.444 | 0.007439823 |
| ENSG00000100258 | LMF2 | 4742.044 | -0.444 | 2.97145E-06 |
| ENSG00000271780 | | 92.464 | -0.444 | 0.002134876 |
| ENSG00000117308 | GALE | 3719.754 | -0.444 | 0.000579281 |
| ENSG00000100100 | PIK3IP1 | 3108.222 | -0.445 | 0.010158809 |
| ENSG00000137818 | RPLP1 | 77431.741 | -0.445 | 0.000658112 |
| ENSG00000202441 | RNY4P10 | 12.145 | -0.445 | 0.005673558 |
| ENSG00000110013 | SIAE | 1334.192 | -0.446 | 0.001110223 |
| ENSG00000232912 | RERE-AS1 | 7.329 | -0.447 | 0.049584149 |
| ENSG00000224424 | PRKAR2A-AS1 | 69.519 | -0.448 | 0.002802326 |
| ENSG00000087077 | TRIP6 | 4960.895 | -0.448 | 0.000326171 |
| ENSG00000070019 | GUCY2C | 74.507 | -0.449 | 0.021087623 |
| ENSG00000133112 | TPT1 | 118557.193 | -0.449 | 5.63081E-05 |
| ENSG00000161653 | NAGS | 222.042 | -0.449 | 0.022334626 |
| ENSG00000196189 | SEMA4A | 2331.777 | -0.449 | 0.004614475 |
| ENSG00000184719 | RNLS | 284.095 | -0.449 | 0.016399222 |

| | | | | |
|---|---|---|---|---|
| ENSG00000147459 | DOCK5 | 1266.064 | -0.450 | 0.005363556 |
| ENSG00000164081 | TEX264 | 3506.207 | -0.450 | 1.66309E-05 |
| ENSG00000196177 | ACADSB | 1069.167 | -0.450 | 0.016425039 |
| ENSG00000184584 | STING1 | 2234.613 | -0.451 | 0.012531311 |
| ENSG00000089057 | SLC23A2 | 2934.719 | -0.451 | 0.022922477 |
| ENSG00000219133 | | 7.033 | -0.451 | 0.048240426 |
| ENSG00000164414 | SLC35A1 | 596.913 | -0.454 | 1.76041E-05 |
| ENSG00000145217 | SLC26A1 | 151.244 | -0.454 | 0.015182011 |
| ENSG00000197536 | IRF1-AS1 | 221.975 | -0.454 | 0.013188745 |
| ENSG00000136147 | PHF11 | 1544.069 | -0.454 | 0.000180352 |
| ENSG00000005469 | CROT | 3176.176 | -0.455 | 0.0348225 |
| ENSG00000168785 | TSPAN5 | 1140.562 | -0.455 | 0.035714645 |
| ENSG00000133872 | SARAF | 8306.032 | -0.456 | 4.66861E-07 |
| ENSG00000148175 | STOM | 7265.061 | -0.457 | 0.042027658 |
| ENSG00000237669 | | 33.217 | -0.457 | 0.021818781 |
| ENSG00000254531 | FLJ20021 | 271.528 | -0.457 | 0.003439465 |
| ENSG00000275183 | LENG9 | 447.907 | -0.457 | 0.004830846 |
| ENSG00000125648 | SLC25A23 | 1642.752 | -0.457 | 0.011698523 |
| ENSG00000126005 | MMP24OS | 2347.239 | -0.457 | 0.001487006 |
| ENSG00000261438 | | 30.639 | -0.457 | 0.038559863 |
| ENSG00000271858 | CYB561D2 | 72.716 | -0.458 | 0.038659205 |
| ENSG00000105516 | DBP | 634.043 | -0.458 | 0.009818516 |
| ENSG00000145020 | AMT | 687.086 | -0.458 | 0.039348348 |
| ENSG00000165837 | ERICH6B | 12.809 | -0.458 | 0.032486473 |
| ENSG00000122378 | PRXL2A | 8390.586 | -0.458 | 0.013778794 |
| ENSG00000243176 | | 16.112 | -0.459 | 0.002333952 |
| ENSG00000233621 | LINC01137 | 323.061 | -0.459 | 0.004500082 |
| ENSG00000170430 | MGMT | 1747.772 | -0.459 | 0.006364661 |
| ENSG00000273014 | | 88.798 | -0.459 | 0.000118981 |
| ENSG00000008394 | MGST1 | 10880.417 | -0.459 | 0.019508279 |
| ENSG00000215840 | | 13.078 | -0.460 | 0.022520766 |
| ENSG00000213859 | KCTD11 | 2248.111 | -0.460 | 0.006519324 |
| ENSG00000142694 | EVA1B | 718.890 | -0.460 | 0.018075766 |
| ENSG00000151338 | MIPOL1 | 439.602 | -0.461 | 0.008970193 |
| ENSG00000132357 | CARD6 | 899.580 | -0.461 | 0.008738225 |
| ENSG00000100911 | PSME2 | 9446.428 | -0.461 | 0.001120743 |
| ENSG00000166676 | TVP23A | 67.653 | -0.462 | 0.036997611 |
| ENSG00000139508 | SLC46A3 | 501.437 | -0.462 | 0.017558277 |
| ENSG00000204267 | TAP2 | 2984.241 | -0.463 | 0.019368776 |
| ENSG00000215068 | | 106.513 | -0.463 | 0.002338645 |
| ENSG00000189171 | S100A13 | 3023.864 | -0.465 | 0.005132369 |
| ENSG00000114779 | ABHD14B | 3599.617 | -0.465 | 0.00042337 |
| ENSG00000136161 | RCBTB2 | 539.343 | -0.466 | 0.000719861 |
| ENSG00000109654 | TRIM2 | 1843.613 | -0.466 | 0.024166275 |
| ENSG00000272182 | | 9.650 | -0.467 | 0.012027351 |
| ENSG00000121797 | CCRL2 | 247.300 | -0.467 | 0.04202521 |
| ENSG00000169242 | EFNA1 | 5416.740 | -0.468 | 0.012457409 |
| ENSG00000129675 | ARHGEF6 | 879.944 | -0.469 | 0.016356795 |
| ENSG00000129422 | MTUS1 | 2784.477 | -0.470 | 0.003585949 |
| ENSG00000230325 | | 13.535 | -0.470 | 0.00561624 |
| ENSG00000116678 | LEPR | 349.159 | -0.470 | 0.025133338 |
| ENSG00000213901 | SLC23A3 | 111.017 | -0.471 | 0.045573144 |
| ENSG00000151470 | C4orf33 | 515.456 | -0.471 | 2.39731E-05 |
| ENSG00000232573 | RPL3P4 | 793.514 | -0.471 | 0.028602406 |
| ENSG00000203546 | | 18.539 | -0.472 | 0.00480316 |
| ENSG00000099840 | IZUMO4 | 160.904 | -0.472 | 0.017158954 |
| ENSG00000280734 | LINC01232 | 378.958 | -0.472 | 0.012062032 |
| ENSG00000174307 | PHLDA3 | 7360.122 | -0.473 | 0.00325102 |
| ENSG00000116704 | SLC35D1 | 1396.179 | -0.473 | 0.000242578 |

| | | | | |
|---|---|---|---|---|
| ENSG00000204261 | PSMB8-AS1 | 489.065 | -0.473 | 0.016125526 |
| ENSG00000280739 | EIF1B-AS1 | 46.530 | -0.474 | 0.00066234 |
| ENSG00000135821 | GLUL | 23386.498 | -0.475 | 0.002599017 |
| ENSG00000278224 | PRICKLE4 | 27.843 | -0.475 | 0.016591404 |
| ENSG00000272799 | | 7.716 | -0.475 | 0.040551793 |
| ENSG00000117335 | CD46 | 13150.331 | -0.475 | 5.73931E-05 |
| ENSG00000229417 | NPM1P25 | 43.026 | -0.475 | 0.048016611 |
| ENSG00000251201 | TMED7-TICAM2 | 13.215 | -0.476 | 0.011609725 |
| ENSG00000253476 | | 6.914 | -0.476 | 0.042838578 |
| ENSG00000204592 | HLA-E | 37259.436 | -0.476 | 0.000773389 |
| ENSG00000099860 | GADD45B | 2468.315 | -0.476 | 0.038005498 |
| ENSG00000130702 | LAMA5 | 14410.610 | -0.477 | 0.002138119 |
| ENSG00000110057 | UNC93B1 | 4188.368 | -0.477 | 0.001583655 |
| ENSG00000244187 | TMEM141 | 2673.038 | -0.477 | 0.00100685 |
| ENSG00000213057 | C1orf220 | 51.139 | -0.477 | 0.043789952 |
| ENSG00000237004 | ZNRF2P1 | 7.916 | -0.478 | 0.007731766 |
| ENSG00000107864 | CPEB3 | 207.069 | -0.478 | 6.01358E-05 |
| ENSG00000164294 | GPX8 | 1162.285 | -0.478 | 0.037277372 |
| ENSG00000245025 | | 20.146 | -0.479 | 0.007776644 |
| ENSG00000163803 | PLB1 | 528.229 | -0.480 | 0.048135216 |
| ENSG00000185043 | CIB1 | 7810.420 | -0.481 | 0.00011177 |
| ENSG00000176222 | ZNF404 | 148.666 | -0.481 | 0.00378618 |
| ENSG00000246582 | | 144.837 | -0.481 | 0.000820491 |
| ENSG00000244073 | RPS4XP6 | 10.317 | -0.481 | 0.020824878 |
| ENSG00000148834 | GSTO1 | 4849.305 | -0.481 | 0.00072571 |
| ENSG00000227959 | | 32.361 | -0.481 | 0.014589167 |
| ENSG00000150667 | FSIP1 | 42.699 | -0.481 | 0.040336877 |
| ENSG00000168071 | CCDC88B | 1471.169 | -0.481 | 0.016797891 |
| ENSG00000152527 | PLEKHH2 | 245.478 | -0.482 | 0.046159291 |
| ENSG00000213433 | RPLP1P6 | 25.064 | -0.482 | 0.013864887 |
| ENSG00000127528 | KLF2 | 753.170 | -0.482 | 0.035463008 |
| ENSG00000244479 | OR2A1-AS1 | 110.141 | -0.483 | 0.011926885 |
| ENSG00000090013 | BLVRB | 3342.351 | -0.483 | 0.00179468 |
| ENSG00000133943 | DGLUCY | 1888.238 | -0.483 | 0.000904643 |
| ENSG00000140450 | ARRDC4 | 889.512 | -0.484 | 0.0083842 |
| ENSG00000085276 | MECOM | 3954.963 | -0.484 | 0.03706252 |
| ENSG00000110318 | CEP126 | 197.012 | -0.485 | 0.019344277 |
| ENSG00000246898 | LINC00920 | 58.606 | -0.486 | 0.025502862 |
| ENSG00000230310 | | 4.569 | -0.486 | 0.029676183 |
| ENSG00000138641 | HERC3 | 1147.184 | -0.486 | 0.001121211 |
| ENSG00000198363 | ASPH | 11841.953 | -0.486 | 0.005571674 |
| ENSG00000259687 | LINC01220 | 12.112 | -0.487 | 0.017537568 |
| ENSG00000105538 | RASIP1 | 1061.294 | -0.488 | 0.049149337 |
| ENSG00000122042 | UBL3 | 2335.567 | -0.488 | 0.000851227 |
| ENSG00000188643 | S100A16 | 19154.645 | -0.489 | 0.026117288 |
| ENSG00000261488 | TBILA | 173.969 | -0.489 | 0.047000115 |
| ENSG00000280153 | | 168.381 | -0.489 | 0.045147592 |
| ENSG00000220842 | RPL21P16 | 247.827 | -0.489 | 0.019967718 |
| ENSG00000173221 | GLRX | 1079.237 | -0.489 | 0.01258543 |
| ENSG00000277895 | | 5.550 | -0.489 | 0.026755191 |
| ENSG00000228201 | | 8.203 | -0.490 | 0.030662342 |
| ENSG00000277662 | | 20.334 | -0.490 | 0.004686549 |
| ENSG00000143344 | RGL1 | 841.785 | -0.491 | 0.011410876 |
| ENSG00000129465 | RIPK3 | 497.020 | -0.491 | 0.004567065 |
| ENSG00000166710 | B2M | 121174.986 | -0.492 | 0.008577592 |
| ENSG00000237172 | B3GNT9 | 898.638 | -0.492 | 0.000193391 |
| ENSG00000260219 | CD2BP2-DT | 163.579 | -0.492 | 0.005046957 |
| ENSG00000003400 | CASP10 | 981.819 | -0.492 | 0.002235599 |
| ENSG00000124713 | GNMT | 24.680 | -0.493 | 0.02262168 |

| | | | | |
|---|---|---|---|---|
| ENSG00000231925 | TAPBP | 17273.028 | -0.493 | 0.000174987 |
| ENSG00000115840 | SLC25A12 | 1025.899 | -0.494 | 7.89525E-07 |
| ENSG00000112773 | TENT5A | 1293.398 | -0.494 | 0.003207325 |
| ENSG00000178537 | SLC25A20 | 584.060 | -0.495 | 0.000112495 |
| ENSG00000272444 | | 19.137 | -0.495 | 0.014907102 |
| ENSG00000275202 | | 78.291 | -0.496 | 0.004083282 |
| ENSG00000171345 | KRT19 | 138180.477 | -0.496 | 0.0348225 |
| ENSG00000255306 | | 20.199 | -0.496 | 0.013872957 |
| ENSG00000107281 | NPDC1 | 2520.519 | -0.496 | 0.019638791 |
| ENSG00000159348 | CYB5R1 | 5201.906 | -0.497 | 0.004030164 |
| ENSG00000253552 | HOXA-AS2 | 473.929 | -0.497 | 0.043421339 |
| ENSG00000026103 | FAS | 713.231 | -0.498 | 0.027652242 |
| ENSG00000250479 | CHCHD10 | 2068.395 | -0.498 | 0.014078052 |
| ENSG00000120594 | PLXDC2 | 1901.616 | -0.500 | 0.043687978 |
| ENSG00000155066 | PROM2 | 12352.677 | -0.500 | 0.049835477 |
| ENSG00000187642 | PERM1 | 550.798 | -0.501 | 0.045452001 |
| ENSG00000142961 | MOB3C | 1655.613 | -0.501 | 1.30698E-06 |
| ENSG00000135926 | TMBIM1 | 14251.795 | -0.502 | 0.000852441 |
| ENSG00000163131 | CTSS | 5443.865 | -0.502 | 0.010089818 |
| ENSG00000261889 | | 27.829 | -0.502 | 0.001684444 |
| ENSG00000115295 | CLIP4 | 1056.180 | -0.503 | 0.013220081 |
| ENSG00000159176 | CSRP1 | 12760.058 | -0.503 | 0.006643831 |
| ENSG00000172738 | TMEM217 | 48.750 | -0.504 | 0.003001006 |
| ENSG00000177989 | ODF3B | 925.225 | -0.504 | 0.016265798 |
| ENSG00000113448 | PDE4D | 981.259 | -0.505 | 0.003110617 |
| ENSG00000273361 | | 8.672 | -0.505 | 0.036304758 |
| ENSG00000173598 | NUDT4 | 2970.206 | -0.505 | 0.000756197 |
| ENSG00000106351 | AGFG2 | 1172.694 | -0.505 | 3.59215E-05 |
| ENSG00000126460 | PRRG2 | 792.939 | -0.505 | 0.001593827 |
| ENSG00000259065 | | 61.548 | -0.506 | 0.006707445 |
| ENSG00000101417 | PXMP4 | 1277.007 | -0.506 | 0.001539666 |
| ENSG00000134470 | IL15RA | 981.666 | -0.506 | 0.046493873 |
| ENSG00000253930 | TNFRSF10A-AS1 | 58.695 | -0.507 | 0.002721135 |
| ENSG00000088836 | SLC4A11 | 1304.794 | -0.507 | 0.038334312 |
| ENSG00000155287 | SLC25A28 | 2136.383 | -0.508 | 1.87659E-06 |
| ENSG00000233483 | EFCAB15P | 93.920 | -0.509 | 0.048411363 |
| ENSG00000186197 | EDARADD | 1202.979 | -0.509 | 0.042841982 |
| ENSG00000178980 | SELENOW | 10973.928 | -0.509 | 6.58425E-06 |
| ENSG00000151150 | ANK3 | 1282.131 | -0.512 | 0.004602264 |
| ENSG00000188064 | WNT7B | 3931.936 | -0.513 | 0.027843487 |
| ENSG00000160179 | ABCG1 | 1914.733 | -0.514 | 0.006762577 |
| ENSG00000246526 | LINC02481 | 81.867 | -0.514 | 0.047117957 |
| ENSG00000051523 | CYBA | 8256.165 | -0.514 | 0.006299198 |
| ENSG00000131831 | RAI2 | 272.194 | -0.515 | 0.042027658 |
| ENSG00000270344 | POC1B-AS1 | 86.452 | -0.516 | 9.89835E-05 |
| ENSG00000256576 | LINC02361 | 28.069 | -0.517 | 0.020187611 |
| ENSG00000054598 | FOXC1 | 1425.502 | -0.517 | 0.018514723 |
| ENSG00000172037 | LAMB2 | 7506.054 | -0.518 | 0.001596151 |
| ENSG00000171223 | JUNB | 17750.953 | -0.518 | 0.000433052 |
| ENSG00000178429 | RPS3AP5 | 37.119 | -0.519 | 0.022510521 |
| ENSG00000278949 | | 13.761 | -0.520 | 0.024299548 |
| ENSG00000204257 | HLA-DMA | 3571.566 | -0.520 | 0.025110828 |
| ENSG00000267519 | MIR23AHG | 1119.564 | -0.520 | 0.006902987 |
| ENSG00000170955 | CAVIN3 | 1009.554 | -0.521 | 0.031911693 |
| ENSG00000247809 | NR2F2-AS1 | 43.235 | -0.521 | 0.032355289 |
| ENSG00000023892 | DEF6 | 1804.954 | -0.522 | 4.82563E-05 |
| ENSG00000100399 | CHADL | 279.996 | -0.522 | 0.007613395 |
| ENSG00000215811 | BTNL10 | 27.123 | -0.522 | 0.033150407 |
| ENSG00000132481 | TRIM47 | 2009.335 | -0.523 | 0.012396521 |

| | | | | |
|---|---|---|---|---|
| ENSG00000157110 | RBPMS | 3531.464 | -0.523 | 0.002459266 |
| ENSG00000274092 | | 11.923 | -0.523 | 0.027647212 |
| ENSG00000109466 | KLHL2 | 851.111 | -0.524 | 2.81683E-05 |
| ENSG00000160932 | LY6E | 20432.906 | -0.524 | 0.048741893 |
| ENSG00000150687 | PRSS23 | 5784.122 | -0.524 | 0.016111641 |
| ENSG00000130755 | GMFG | 1025.048 | -0.524 | 0.035748073 |
| ENSG00000169403 | PTAFR | 1021.307 | -0.525 | 0.020735926 |
| ENSG00000234807 | LINC01135 | 5.683 | -0.526 | 0.023233255 |
| ENSG00000112874 | NUDT12 | 529.990 | -0.526 | 0.001626708 |
| ENSG00000199024 | MIR103A2 | 9.940 | -0.526 | 0.002852561 |
| ENSG00000106327 | TFR2 | 263.466 | -0.526 | 0.042775307 |
| ENSG00000259921 | | 8.448 | -0.527 | 0.012921829 |
| ENSG00000026297 | RNASET2 | 3762.794 | -0.527 | 0.005688973 |
| ENSG00000197776 | KLHDC1 | 73.620 | -0.528 | 0.000670535 |
| ENSG00000164674 | SYTL3 | 305.303 | -0.528 | 0.007192999 |
| ENSG00000139597 | N4BP2L1 | 516.736 | -0.528 | 0.00045661 |
| ENSG00000129521 | EGLN3 | 3869.570 | -0.528 | 0.044070233 |
| ENSG00000125968 | ID1 | 13368.893 | -0.528 | 0.042626005 |
| ENSG00000163959 | SLC51A | 244.883 | -0.529 | 0.007460073 |
| ENSG00000088881 | EBF4 | 728.441 | -0.529 | 0.025974311 |
| ENSG00000130005 | GAMT | 488.235 | -0.529 | 0.044070233 |
| ENSG00000275389 | | 4.942 | -0.529 | 0.011274567 |
| ENSG00000187699 | C2orf88 | 157.809 | -0.531 | 0.041283268 |
| ENSG00000204264 | PSMB8 | 5610.088 | -0.531 | 0.003737336 |
| ENSG00000275074 | NUDT18 | 447.279 | -0.531 | 0.000301973 |
| ENSG00000263293 | EFCAB13-DT | 11.007 | -0.531 | 0.04270061 |
| ENSG00000233690 | EBAG9P1 | 6.881 | -0.532 | 0.029474329 |
| ENSG00000150540 | HNMT | 1753.836 | -0.532 | 0.004214195 |
| ENSG00000143797 | MBOAT2 | 2380.384 | -0.533 | 0.012843068 |
| ENSG00000137642 | SORL1 | 6374.718 | -0.533 | 0.049088973 |
| ENSG00000117394 | SLC2A1 | 26949.613 | -0.534 | 0.027249848 |
| ENSG00000068137 | PLEKHH3 | 2840.317 | -0.537 | 2.41693E-05 |
| ENSG00000219755 | | 18.396 | -0.537 | 0.005192755 |
| ENSG00000256043 | CTSO | 1174.230 | -0.537 | 0.000315093 |
| ENSG00000154102 | C16orf74 | 1419.281 | -0.538 | 0.025978147 |
| ENSG00000235863 | B3GALT4 | 936.725 | -0.538 | 0.000195316 |
| ENSG00000241429 | EEF1A1P25 | 9.930 | -0.540 | 0.017826353 |
| ENSG00000124593 | | 720.603 | -0.540 | 0.001420708 |
| ENSG00000152213 | ARL11 | 182.088 | -0.542 | 0.015365088 |
| ENSG00000102763 | VWA8 | 1347.771 | -0.542 | 5.007E-05 |
| ENSG00000122359 | ANXA11 | 14713.077 | -0.542 | 3.03026E-06 |
| ENSG00000140876 | NUDT7 | 63.421 | -0.542 | 0.016376326 |
| ENSG00000163932 | PRKCD | 4794.740 | -0.543 | 2.18241E-05 |
| ENSG00000272979 | | 14.160 | -0.543 | 0.010268932 |
| ENSG00000118515 | SGK1 | 4840.410 | -0.543 | 0.036754044 |
| ENSG00000275022 | MIR6753 | 4.679 | -0.543 | 0.015216473 |
| ENSG00000265778 | ZNF516-AS1 | 45.078 | -0.544 | 0.032245677 |
| ENSG00000179715 | PCED1B | 824.183 | -0.544 | 0.002272737 |
| ENSG00000231006 | RPL7P32 | 13.224 | -0.544 | 0.012580148 |
| ENSG00000275484 | | 24.126 | -0.544 | 0.022405324 |
| ENSG00000273628 | | 5.676 | -0.545 | 0.038659205 |
| ENSG00000156966 | B3GNT7 | 526.719 | -0.546 | 0.047191803 |
| ENSG00000259583 | | 130.981 | -0.547 | 0.002180507 |
| ENSG00000182704 | TSKU | 4822.295 | -0.547 | 0.001599924 |
| ENSG00000203724 | C1orf53 | 125.856 | -0.547 | 0.000283322 |
| ENSG00000149131 | SERPING1 | 11604.821 | -0.547 | 0.041978869 |
| ENSG00000153714 | LURAP1L | 568.331 | -0.547 | 0.012689672 |
| ENSG00000035664 | DAPK2 | 295.120 | -0.548 | 0.024559806 |
| ENSG00000236305 | SLC12A9-AS1 | 29.452 | -0.548 | 0.006213269 |

| | | | | |
|---|---|---|---|---|
| ENSG00000196502 | SULT1A1 | 835.995 | -0.549 | 0.028550225 |
| ENSG00000261655 | | 28.276 | -0.549 | 0.033688928 |
| ENSG00000103260 | METRN | 772.213 | -0.549 | 0.015732998 |
| ENSG00000139899 | CBLN3 | 143.770 | -0.549 | 0.000254628 |
| ENSG00000196274 | Metazoa_SRP | 7.941 | -0.550 | 0.008319533 |
| ENSG00000270504 | | 249.965 | -0.550 | 0.00475405 |
| ENSG00000238005 | | 27.653 | -0.552 | 0.008088143 |
| ENSG00000254718 | | 15.276 | -0.552 | 0.019073443 |
| ENSG00000013364 | MVP | 12115.341 | -0.552 | 1.86037E-05 |
| ENSG00000125122 | LRRC29 | 148.774 | -0.552 | 9.73145E-05 |
| ENSG00000260325 | HSPB9 | 26.074 | -0.553 | 0.011766405 |
| ENSG00000276945 | | 6.120 | -0.553 | 0.00853114 |
| ENSG00000179397 | CATSPERE | 39.675 | -0.553 | 0.002925035 |
| ENSG00000121858 | TNFSF10 | 7989.353 | -0.554 | 0.041871477 |
| ENSG00000268204 | | 31.500 | -0.554 | 0.041534504 |
| ENSG00000205683 | DPF3 | 30.676 | -0.554 | 0.020299228 |
| ENSG00000279422 | | 5.220 | -0.555 | 0.047585883 |
| ENSG00000016402 | IL20RA | 1378.244 | -0.555 | 0.022154568 |
| ENSG00000184524 | CEND1 | 16.455 | -0.555 | 0.031863209 |
| ENSG00000267370 | | 53.580 | -0.555 | 0.006619348 |
| ENSG00000249846 | LINC02021 | 23.599 | -0.555 | 0.011479598 |
| ENSG00000183044 | ABAT | 1423.754 | -0.555 | 0.037419573 |
| ENSG00000125170 | DOK4 | 1304.649 | -0.555 | 9.27964E-06 |
| ENSG00000174500 | GCSAM | 210.600 | -0.557 | 0.007537383 |
| ENSG00000258940 | | 26.469 | -0.558 | 4.78272E-05 |
| ENSG00000112619 | PRPH2 | 51.461 | -0.560 | 0.0363332 |
| ENSG00000275367 | | 21.634 | -0.560 | 0.04569585 |
| ENSG00000065357 | DGKA | 4342.024 | -0.561 | 0.000177478 |
| ENSG00000248677 | LINC02102 | 5.468 | -0.561 | 0.041726916 |
| ENSG00000172794 | RAB37 | 110.106 | -0.561 | 0.028540815 |
| ENSG00000149716 | LTO1 | 1330.494 | -0.562 | 0.002852561 |
| ENSG00000278058 | | 17.780 | -0.562 | 0.026294048 |
| ENSG00000118507 | AKAP7 | 318.357 | -0.562 | 0.003971369 |
| ENSG00000205220 | PSMB10 | 2163.765 | -0.563 | 0.000335951 |
| ENSG00000206503 | HLA-A | 70513.089 | -0.563 | 0.010604769 |
| ENSG00000082196 | C1QTNF3 | 337.016 | -0.563 | 0.03230253 |
| ENSG00000111796 | KLRB1 | 84.383 | -0.563 | 0.03284533 |
| ENSG00000276600 | RAB7B | 562.688 | -0.563 | 0.041346535 |
| ENSG00000014257 | ACP3 | 263.688 | -0.564 | 0.037999061 |
| ENSG00000188396 | DYNLT4 | 5.613 | -0.564 | 0.027275355 |
| ENSG00000136156 | ITM2B | 27553.837 | -0.564 | 1.36188E-05 |
| ENSG00000160211 | G6PD | 6365.894 | -0.565 | 0.011683863 |
| ENSG00000164185 | ZNF474 | 14.143 | -0.565 | 0.019895059 |
| ENSG00000085733 | CTTN | 17863.438 | -0.565 | 6.38337E-05 |
| ENSG00000160712 | IL6R | 572.454 | -0.566 | 0.016111641 |
| ENSG00000105991 | HOXA1 | 359.529 | -0.566 | 0.005851257 |
| ENSG00000236383 | CCDC200 | 21.756 | -0.566 | 0.006628749 |
| ENSG00000179256 | SMCO3 | 13.582 | -0.566 | 0.02475294 |
| ENSG00000260038 | | 5.262 | -0.566 | 0.01843053 |
| ENSG00000213145 | CRIP1 | 194.285 | -0.567 | 0.023236946 |
| ENSG00000223509 | | 72.872 | -0.567 | 0.000554884 |
| ENSG00000254122 | PCDHGB7 | 67.594 | -0.567 | 0.026184813 |
| ENSG00000105281 | SLC1A5 | 13853.052 | -0.567 | 3.69099E-05 |
| ENSG00000232415 | ELN-AS1 | 7.095 | -0.567 | 0.038379453 |
| ENSG00000244021 | | 11.237 | -0.567 | 0.022025812 |
| ENSG00000073060 | SCARB1 | 2856.058 | -0.568 | 0.000788298 |
| ENSG00000271303 | SRXN1 | 121.669 | -0.568 | 0.011634804 |
| ENSG00000072310 | SREBF1 | 9787.042 | -0.568 | 0.000848976 |
| ENSG00000244198 | ARHGEF35-AS1 | 17.697 | -0.569 | 0.008046758 |

| | | | | |
|---|---|---|---|---|
| ENSG00000259577 | CERNA1 | 26.341 | -0.569 | 0.000851709 |
| ENSG00000119630 | PGF | 1267.677 | -0.569 | 0.02653942 |
| ENSG00000214274 | ANG | 226.984 | -0.570 | 0.004412076 |
| ENSG00000242193 | CRYZL2P | 465.347 | -0.571 | 0.009246763 |
| ENSG00000214650 | | 5.988 | -0.571 | 0.043148387 |
| ENSG00000239282 | CASTOR1 | 127.979 | -0.572 | 0.000308013 |
| ENSG00000164877 | MICALL2 | 2924.113 | -0.572 | 1.76505E-05 |
| ENSG00000167701 | GPT | 138.265 | -0.572 | 0.03431377 |
| ENSG00000016391 | CHDH | 312.797 | -0.573 | 0.043692688 |
| ENSG00000261187 | | 14.290 | -0.573 | 0.001345225 |
| ENSG00000229619 | MBNL1-AS1 | 195.726 | -0.574 | 0.019952107 |
| ENSG00000274515 | | 5.991 | -0.575 | 0.006365077 |
| ENSG00000132692 | BCAN | 60.525 | -0.575 | 0.026247926 |
| ENSG00000121577 | POPDC2 | 236.530 | -0.575 | 0.047829342 |
| ENSG00000159733 | ZFYVE28 | 375.638 | -0.575 | 0.000937292 |
| ENSG00000272449 | | 98.976 | -0.576 | 0.016608218 |
| ENSG00000101825 | MXRA5 | 5364.975 | -0.576 | 0.049880037 |
| ENSG00000118308 | IRAG2 | 1283.579 | -0.577 | 0.048331708 |
| ENSG00000140199 | SLC12A6 | 1693.412 | -0.577 | 0.000207206 |
| ENSG00000213071 | LPAL2 | 26.272 | -0.577 | 0.013645879 |
| ENSG00000228839 | PIK3IP1-DT | 16.213 | -0.578 | 0.002194184 |
| ENSG00000234785 | EEF1GP5 | 11.832 | -0.578 | 0.034611744 |
| ENSG00000235241 | | 59.232 | -0.579 | 0.032631055 |
| ENSG00000109452 | INPP4B | 2058.572 | -0.580 | 0.004593125 |
| ENSG00000174791 | RIN1 | 1198.541 | -0.580 | 0.003870087 |
| ENSG00000270265 | | 6.634 | -0.581 | 0.01359924 |
| ENSG00000243896 | OR2A7 | 21.335 | -0.581 | 0.016685372 |
| ENSG00000259275 | | 40.601 | -0.581 | 0.02773769 |
| ENSG00000103811 | CTSH | 20517.652 | -0.581 | 0.004782983 |
| ENSG00000213654 | GPSM3 | 1108.792 | -0.582 | 0.005323567 |
| ENSG00000198467 | TPM2 | 11878.925 | -0.582 | 0.043091157 |
| ENSG00000145244 | CORIN | 83.810 | -0.582 | 0.027537391 |
| ENSG00000175806 | MSRA | 508.813 | -0.583 | 7.53917E-05 |
| ENSG00000019582 | CD74 | 81174.175 | -0.584 | 0.030696119 |
| ENSG00000258818 | RNASE4 | 44.265 | -0.584 | 0.004710052 |
| ENSG00000128284 | APOL3 | 1708.361 | -0.584 | 0.009701237 |
| ENSG00000255046 | | 7.961 | -0.584 | 0.013729905 |
| ENSG00000272030 | | 37.822 | -0.586 | 0.000848976 |
| ENSG00000147027 | TMEM47 | 2277.548 | -0.586 | 0.02467023 |
| ENSG00000215915 | ATAD3C | 376.075 | -0.586 | 0.017155124 |
| ENSG00000131401 | NAPSB | 276.734 | -0.586 | 0.044883217 |
| ENSG00000176919 | C8G | 44.226 | -0.587 | 0.003861165 |
| ENSG00000104611 | SH2D4A | 1881.347 | -0.587 | 0.001179636 |
| ENSG00000130653 | PNPLA7 | 206.336 | -0.588 | 0.002317742 |
| ENSG00000108352 | RAPGEFL1 | 6528.550 | -0.588 | 0.02074914 |
| ENSG00000107742 | SPOCK2 | 1535.037 | -0.589 | 0.033155872 |
| ENSG00000110031 | LPXN | 752.117 | -0.589 | 0.007224904 |
| ENSG00000115041 | KCNIP3 | 147.855 | -0.590 | 0.034714817 |
| ENSG00000129538 | RNASE1 | 5317.773 | -0.591 | 0.016111641 |
| ENSG00000166949 | SMAD3 | 6541.808 | -0.591 | 0.000226797 |
| ENSG00000277678 | RNU1-153P | 4.783 | -0.592 | 0.033954156 |
| ENSG00000132386 | SERPINF1 | 4917.756 | -0.592 | 0.039691411 |
| ENSG00000181585 | TMIE | 62.535 | -0.592 | 0.020149201 |
| ENSG00000112053 | SLC26A8 | 15.215 | -0.592 | 0.015732998 |
| ENSG00000235314 | LINC00957 | 110.136 | -0.594 | 0.000536985 |
| ENSG00000231346 | LINC01160 | 14.016 | -0.594 | 0.02835292 |
| ENSG00000259712 | | 10.557 | -0.595 | 0.029224247 |
| ENSG00000104361 | NIPAL2 | 2019.698 | -0.595 | 8.6836E-05 |
| ENSG00000129038 | LOXL1 | 2116.987 | -0.595 | 0.012520844 |

| | | | | |
|---|---|---|---|---|
| ENSG00000265972 | TXNIP | 22701.095 | -0.596 | 0.000848976 |
| ENSG00000125746 | EML2 | 3221.047 | -0.596 | 5.30666E-07 |
| ENSG00000228748 | | 23.835 | -0.596 | 0.008970193 |
| ENSG00000235529 | AGAP1-IT1 | 11.173 | -0.597 | 0.031938882 |
| ENSG00000201499 | RNU6-312P | 3.970 | -0.598 | 0.017589427 |
| ENSG00000142552 | RCN3 | 2423.536 | -0.598 | 0.032301925 |
| ENSG00000102890 | ELMO3 | 2577.739 | -0.598 | 0.001576413 |
| ENSG00000007312 | CD79B | 200.565 | -0.598 | 0.049993567 |
| ENSG00000106560 | GIMAP2 | 291.967 | -0.599 | 0.001162657 |
| ENSG00000175879 | HOXD8 | 323.154 | -0.600 | 0.000799739 |
| ENSG00000070404 | FSTL3 | 2636.300 | -0.600 | 0.043091157 |
| ENSG00000229953 | | 63.947 | -0.602 | 0.046093593 |
| ENSG00000053918 | KCNQ1 | 1271.906 | -0.602 | 0.021598643 |
| ENSG00000102780 | DGKH | 1858.156 | -0.603 | 0.000258562 |
| ENSG00000177363 | LRRN4CL | 122.805 | -0.603 | 0.024090844 |
| ENSG00000226891 | LINC01359 | 8.676 | -0.603 | 0.043585257 |
| ENSG00000185033 | SEMA4B | 13355.469 | -0.604 | 0.002802326 |
| ENSG00000125966 | MMP24 | 66.415 | -0.604 | 0.006614887 |
| ENSG00000238120 | LINC01589 | 20.543 | -0.604 | 0.026591913 |
| ENSG00000171444 | MCC | 1908.123 | -0.604 | 0.002207666 |
| ENSG00000214279 | SCART1 | 97.817 | -0.605 | 0.010865047 |
| ENSG00000116690 | PRG4 | 48.415 | -0.605 | 0.029101626 |
| ENSG00000162882 | HAAO | 228.288 | -0.606 | 0.00408364 |
| ENSG00000231563 | | 42.394 | -0.606 | 0.025891017 |
| ENSG00000278514 | | 31.020 | -0.606 | 0.027307551 |
| ENSG00000251287 | ALG1L2 | 26.760 | -0.607 | 0.002274093 |
| ENSG00000261504 | LINC01686 | 24.372 | -0.607 | 0.022520766 |
| ENSG00000166979 | EVA1C | 620.137 | -0.607 | 0.016237632 |
| ENSG00000258745 | | 6.158 | -0.608 | 0.034495023 |
| ENSG00000248124 | RRN3P1 | 123.425 | -0.608 | 0.005400645 |
| ENSG00000099194 | SCD | 22524.099 | -0.609 | 0.014705216 |
| ENSG00000117318 | ID3 | 4368.796 | -0.609 | 0.004202146 |
| ENSG00000104219 | ZDHHC2 | 1205.522 | -0.610 | 0.010705857 |
| ENSG00000112561 | TFEB | 1014.605 | -0.611 | 0.00096772 |
| ENSG00000105329 | TGFB1 | 4878.139 | -0.612 | 0.000670535 |
| ENSG00000142910 | TINAGL1 | 21345.019 | -0.612 | 0.008006687 |
| ENSG00000250722 | SELENOP | 1573.987 | -0.612 | 0.040751806 |
| ENSG00000142235 | LMTK3 | 461.018 | -0.612 | 0.016513399 |
| ENSG00000159871 | LYPD5 | 529.556 | -0.613 | 0.012410153 |
| ENSG00000108370 | RGS9 | 64.968 | -0.613 | 0.037667144 |
| ENSG00000257298 | | 18.911 | -0.613 | 0.011589004 |
| ENSG00000261390 | MAFTRR | 10.508 | -0.613 | 0.030702039 |
| ENSG00000000971 | CFH | 3563.137 | -0.614 | 0.013703986 |
| ENSG00000232926 | | 5.574 | -0.615 | 0.029348998 |
| ENSG00000126458 | RRAS | 2218.491 | -0.615 | 0.000568505 |
| ENSG00000146192 | FGD2 | 396.039 | -0.617 | 0.007688988 |
| ENSG00000157613 | CREB3L1 | 1594.186 | -0.617 | 0.031097344 |
| ENSG00000130707 | ASS1 | 10841.811 | -0.617 | 0.047069292 |
| ENSG00000174514 | MFSD4A | 210.728 | -0.617 | 0.020885359 |
| ENSG00000267750 | RUNDC3A-AS1 | 51.542 | -0.619 | 0.016376326 |
| ENSG00000166501 | PRKCB | 406.931 | -0.619 | 0.041279168 |
| ENSG00000109743 | BST1 | 499.784 | -0.620 | 0.030965698 |
| ENSG00000180739 | S1PR5 | 808.759 | -0.620 | 0.021207591 |
| ENSG00000248429 | FAM198B-AS1 | 141.803 | -0.620 | 0.01137452 |
| ENSG00000254503 | | 17.789 | -0.622 | 0.011902334 |
| ENSG00000082438 | COBLL1 | 2485.823 | -0.622 | 0.000848976 |
| ENSG00000166619 | BLCAP | 10476.422 | -0.622 | 3.03026E-06 |
| ENSG00000052802 | MSMO1 | 3440.296 | -0.622 | 2.24352E-05 |
| ENSG00000258711 | | 83.928 | -0.622 | 0.033867471 |

| | | | | |
|---|---|---|---|---|
| ENSG00000236060 | HSPB1P1 | 176.521 | -0.623 | 0.005140266 |
| ENSG00000279894 | | 16.834 | -0.623 | 0.005400645 |
| ENSG00000104687 | GSR | 4258.325 | -0.623 | 0.000172412 |
| ENSG00000151458 | ANKRD50 | 2727.622 | -0.623 | 0.000804847 |
| ENSG00000233818 | | 54.127 | -0.623 | 0.016729428 |
| ENSG00000254827 | SLC22A18AS | 76.220 | -0.624 | 0.006262082 |
| ENSG00000235169 | SMIM1 | 63.843 | -0.625 | 0.017296021 |
| ENSG00000272369 | | 28.812 | -0.626 | 0.004233606 |
| ENSG00000185201 | IFITM2 | 3055.594 | -0.626 | 0.004846628 |
| ENSG00000226332 | | 301.182 | -0.626 | 0.002107113 |
| ENSG00000184988 | TMEM106A | 203.090 | -0.627 | 0.007654967 |
| ENSG00000100767 | PAPLN | 731.760 | -0.627 | 0.00690707 |
| ENSG00000251131 | | 5.130 | -0.628 | 0.015099696 |
| ENSG00000170684 | ZNF296 | 440.575 | -0.630 | 0.001019191 |
| ENSG00000099139 | PCSK5 | 407.885 | -0.630 | 0.029274724 |
| ENSG00000170345 | FOS | 18411.520 | -0.630 | 0.010975812 |
| ENSG00000168010 | ATG16L2 | 1463.374 | -0.631 | 0.000504626 |
| ENSG00000091262 | ABCC6 | 67.358 | -0.631 | 0.010338469 |
| ENSG00000237523 | LINC00857 | 173.591 | -0.632 | 0.006607144 |
| ENSG00000177191 | B3GNT8 | 401.348 | -0.632 | 0.000711832 |
| ENSG00000275765 | | 64.527 | -0.633 | 2.34574E-05 |
| ENSG00000025708 | TYMP | 8419.071 | -0.633 | 0.026524863 |
| ENSG00000140945 | CDH13 | 867.309 | -0.633 | 0.022525075 |
| ENSG00000235770 | LINC00607 | 19.677 | -0.634 | 0.035730444 |
| ENSG00000257252 | | 10.133 | -0.635 | 0.014207891 |
| ENSG00000074370 | ATP2A3 | 2067.094 | -0.636 | 0.00516298 |
| ENSG00000228037 | | 13.611 | -0.636 | 0.019675742 |
| ENSG00000167261 | DPEP2 | 296.716 | -0.637 | 0.03883497 |
| ENSG00000133106 | EPSTI1 | 1859.687 | -0.637 | 0.017680396 |
| ENSG00000260657 | | 6.004 | -0.638 | 0.045087961 |
| ENSG00000187098 | MITF | 332.074 | -0.639 | 0.003352863 |
| ENSG00000227258 | SMIM2-AS1 | 34.604 | -0.640 | 0.010093137 |
| ENSG00000253123 | | 26.527 | -0.640 | 0.018373308 |
| ENSG00000278954 | | 25.929 | -0.640 | 0.026997296 |
| ENSG00000148357 | HMCN2 | 183.997 | -0.640 | 0.041888899 |
| ENSG00000112782 | CLIC5 | 246.168 | -0.641 | 0.044995736 |
| ENSG00000110719 | TCIRG1 | 6693.614 | -0.641 | 9.1291E-06 |
| ENSG00000251000 | GGCTP1 | 19.090 | -0.642 | 0.000554176 |
| ENSG00000105472 | CLEC11A | 733.001 | -0.642 | 0.009940436 |
| ENSG00000104432 | IL7 | 119.041 | -0.643 | 0.001532449 |
| ENSG00000279296 | PRAL | 12.015 | -0.644 | 0.037352304 |
| ENSG00000019549 | SNAI2 | 2033.225 | -0.644 | 0.018009033 |
| ENSG00000075073 | TACR2 | 91.466 | -0.645 | 0.03284533 |
| ENSG00000253981 | ALG1L13P | 19.935 | -0.646 | 0.009861593 |
| ENSG00000225931 | | 4.432 | -0.646 | 0.029306186 |
| ENSG00000158525 | CPA5 | 19.081 | -0.646 | 0.031339424 |
| ENSG00000268536 | | 4.728 | -0.646 | 0.007009481 |
| ENSG00000214922 | HLA-F-AS1 | 82.044 | -0.647 | 0.004472977 |
| ENSG00000117009 | KMO | 90.360 | -0.647 | 0.01673482 |
| ENSG00000233901 | LINC01503 | 857.151 | -0.649 | 0.002831856 |
| ENSG00000101335 | MYL9 | 15223.370 | -0.650 | 0.032397324 |
| ENSG00000166428 | PLD4 | 93.813 | -0.652 | 0.025199256 |
| ENSG00000120129 | DUSP1 | 10029.098 | -0.652 | 0.007669961 |
| ENSG00000198483 | ANKRD35 | 345.574 | -0.652 | 0.012138044 |
| ENSG00000127824 | TUBA4A | 4342.043 | -0.652 | 0.006633364 |
| ENSG00000279805 | | 14.307 | -0.654 | 0.007784641 |
| ENSG00000255471 | | 22.411 | -0.654 | 0.008338887 |
| ENSG00000176454 | LPCAT4 | 3999.843 | -0.655 | 9.27042E-06 |
| ENSG00000112303 | VNN2 | 94.790 | -0.656 | 0.046738983 |

| | | | | |
|---|---|---|---|---|
| ENSG00000103472 | RRN3P2 | 36.994 | -0.657 | 0.003177429 |
| ENSG00000140287 | HDC | 64.354 | -0.657 | 0.036243248 |
| ENSG00000261357 | | 29.747 | -0.657 | 0.039968275 |
| ENSG00000228078 | HLA-U | 14.743 | -0.658 | 0.035967031 |
| ENSG00000164237 | CMBL | 1088.503 | -0.659 | 0.003993853 |
| ENSG00000179163 | FUCA1 | 3872.562 | -0.660 | 8.64706E-06 |
| ENSG00000168899 | VAMP5 | 1685.040 | -0.660 | 0.002172977 |
| ENSG00000225864 | | 12.953 | -0.661 | 0.031154353 |
| ENSG00000130943 | PKDREJ | 17.272 | -0.661 | 0.00433105 |
| ENSG00000131435 | PDLIM4 | 1070.984 | -0.662 | 0.022611353 |
| ENSG00000137501 | SYTL2 | 1830.860 | -0.662 | 0.00351588 |
| ENSG00000227262 | HCG4B | 48.169 | -0.663 | 0.010149152 |
| ENSG00000197182 | MIRLET7BHG | 165.606 | -0.663 | 0.00254431 |
| ENSG00000204653 | ASPDH | 12.205 | -0.664 | 0.017101352 |
| ENSG00000106123 | EPHB6 | 4995.539 | -0.664 | 0.033642086 |
| ENSG00000204618 | RNF39 | 381.581 | -0.665 | 0.00397074 |
| ENSG00000113578 | FGF1 | 110.771 | -0.666 | 0.016042318 |
| ENSG00000172830 | SSH3 | 8771.381 | -0.667 | 0.000359281 |
| ENSG00000008853 | RHOBTB2 | 2320.705 | -0.668 | 0.002852561 |
| ENSG00000184730 | APOBR | 1052.383 | -0.668 | 0.006196007 |
| ENSG00000122224 | LY9 | 90.696 | -0.669 | 0.047524536 |
| ENSG00000198121 | LPAR1 | 744.383 | -0.669 | 0.000550196 |
| ENSG00000168675 | LDLRAD4 | 548.167 | -0.669 | 0.002410753 |
| ENSG00000260001 | TGFBR3L | 28.393 | -0.669 | 0.033025722 |
| ENSG00000272767 | JMJD1C-AS1 | 49.090 | -0.669 | 0.001010031 |
| ENSG00000175416 | CLTB | 6338.509 | -0.670 | 4.93862E-05 |
| ENSG00000253641 | LINCR-0001 | 19.256 | -0.670 | 0.047446972 |
| ENSG00000103024 | NME3 | 2123.637 | -0.670 | 6.42117E-05 |
| ENSG00000272647 | | 3.595 | -0.671 | 0.046273511 |
| ENSG00000246223 | LINC01550 | 23.970 | -0.671 | 0.042574046 |
| ENSG00000187492 | CDHR4 | 6.465 | -0.671 | 0.027314425 |
| ENSG00000120093 | HOXB3 | 1637.885 | -0.672 | 0.008363753 |
| ENSG00000258733 | LINC02328 | 23.170 | -0.673 | 0.027808201 |
| ENSG00000166780 | BMERB1 | 600.761 | -0.674 | 0.001458435 |
| ENSG00000278831 | | 13.027 | -0.674 | 0.005556638 |
| ENSG00000269889 | | 10.741 | -0.674 | 0.009468191 |
| ENSG00000187554 | TLR5 | 296.777 | -0.674 | 0.000772196 |
| ENSG00000006555 | TTC22 | 1422.567 | -0.675 | 0.001111184 |
| ENSG00000169884 | WNT10B | 46.543 | -0.675 | 0.033096074 |
| ENSG00000197580 | BCO2 | 80.960 | -0.675 | 0.005098621 |
| ENSG00000165457 | FOLR2 | 584.502 | -0.676 | 0.034807558 |
| ENSG00000180353 | HCLS1 | 3150.099 | -0.677 | 0.006916472 |
| ENSG00000121064 | SCPEP1 | 8664.456 | -0.679 | 0.000422989 |
| ENSG00000089356 | FXYD3 | 31289.082 | -0.680 | 0.004990284 |
| ENSG00000249352 | LINC02198 | 12.228 | -0.680 | 0.044287801 |
| ENSG00000181690 | PLAG1 | 327.316 | -0.681 | 0.004948103 |
| ENSG00000267795 | SMIM22 | 2828.955 | -0.681 | 0.024189589 |
| ENSG00000225217 | HSPA7 | 286.411 | -0.682 | 0.016369664 |
| ENSG00000276850 | | 513.866 | -0.683 | 0.007211234 |
| ENSG00000166126 | AMN | 375.766 | -0.683 | 0.012284224 |
| ENSG00000105499 | PLA2G4C | 226.494 | -0.684 | 0.005046957 |
| ENSG00000255650 | FAM222A-AS1 | 7.319 | -0.684 | 0.0378215 |
| ENSG00000101605 | MYOM1 | 148.728 | -0.685 | 0.010775685 |
| ENSG00000280062 | | 4.236 | -0.685 | 0.039775819 |
| ENSG00000198133 | TMEM229B | 593.297 | -0.686 | 0.000120676 |
| ENSG00000186891 | TNFRSF18 | 348.701 | -0.687 | 0.020652649 |
| ENSG00000257594 | GALNT4 | 21.051 | -0.687 | 0.015124107 |
| ENSG00000173210 | ABLIM3 | 1711.282 | -0.688 | 0.001999168 |
| ENSG00000019991 | HGF | 196.443 | -0.688 | 0.044275973 |

| | | | | |
|---|---|---|---|---|
| ENSG00000230943 | LINC02541 | 113.286 | -0.688 | 0.016031832 |
| ENSG00000065989 | PDE4A | 851.609 | -0.688 | 3.9287E-06 |
| ENSG00000079263 | SP140 | 237.353 | -0.689 | 0.029593069 |
| ENSG00000118596 | SLC16A7 | 514.836 | -0.690 | 0.005984602 |
| ENSG00000214944 | ARHGEF28 | 876.149 | -0.691 | 0.000151488 |
| ENSG00000157368 | IL34 | 223.786 | -0.692 | 0.007214249 |
| ENSG00000237429 | | 4.467 | -0.692 | 0.021611668 |
| ENSG00000143153 | ATP1B1 | 7073.042 | -0.693 | 0.001921758 |
| ENSG00000235052 | | 12.084 | -0.693 | 0.014261774 |
| ENSG00000185669 | SNAI3 | 65.341 | -0.693 | 0.003165587 |
| ENSG00000253258 | | 7.615 | -0.694 | 0.048228204 |
| ENSG00000231672 | DIRC3 | 27.353 | -0.694 | 0.045315023 |
| ENSG00000267801 | | 38.921 | -0.694 | 0.000915471 |
| ENSG00000089012 | SIRPG | 138.406 | -0.695 | 0.027266908 |
| ENSG00000082074 | FYB1 | 1197.605 | -0.695 | 0.033369386 |
| ENSG00000232855 | | 48.306 | -0.695 | 0.038848882 |
| ENSG00000170961 | HAS2 | 749.493 | -0.695 | 0.040225312 |
| ENSG00000163364 | LINC01116 | 188.629 | -0.695 | 0.019618527 |
| ENSG00000272505 | | 31.493 | -0.697 | 0.006219051 |
| ENSG00000125848 | FLRT3 | 1473.075 | -0.697 | 0.026033498 |
| ENSG00000119632 | IFI27L2 | 1274.854 | -0.697 | 0.001292662 |
| ENSG00000150281 | CTF1 | 522.633 | -0.697 | 0.000108614 |
| ENSG00000092607 | TBX15 | 75.551 | -0.697 | 0.02524066 |
| ENSG00000260641 | | 18.434 | -0.698 | 0.021828169 |
| ENSG00000168062 | BATF2 | 622.307 | -0.698 | 0.03064687 |
| ENSG00000253187 | HOXA10-AS | 47.111 | -0.699 | 0.016718825 |
| ENSG00000146006 | LRRTM2 | 11.672 | -0.699 | 0.006476328 |
| ENSG00000175294 | CATSPER1 | 186.742 | -0.699 | 0.04946102 |
| ENSG00000120915 | EPHX2 | 505.864 | -0.700 | 0.003980974 |
| ENSG00000267279 | | 42.268 | -0.706 | 0.034751076 |
| ENSG00000224769 | MUC20P1 | 292.463 | -0.706 | 0.040565147 |
| ENSG00000139344 | AMDHD1 | 39.664 | -0.706 | 0.035593224 |
| ENSG00000080031 | PTPRH | 378.595 | -0.706 | 0.038319368 |
| ENSG00000261616 | | 29.070 | -0.707 | 0.013061879 |
| ENSG00000103888 | CEMIP | 1003.332 | -0.708 | 0.015005678 |
| ENSG00000172236 | TPSAB1 | 384.894 | -0.708 | 0.025280396 |
| ENSG00000240065 | PSMB9 | 3604.417 | -0.710 | 0.011658528 |
| ENSG00000274414 | | 32.447 | -0.710 | 0.009567065 |
| ENSG00000004468 | CD38 | 364.553 | -0.710 | 0.043123269 |
| ENSG00000276317 | | 12.675 | -0.711 | 0.011902334 |
| ENSG00000231530 | | 9.906 | -0.711 | 0.000330291 |
| ENSG00000146021 | KLHL3 | 342.508 | -0.711 | 0.000111328 |
| ENSG00000237238 | BMS1P10 | 32.245 | -0.712 | 0.001708002 |
| ENSG00000240990 | HOXA11-AS | 313.144 | -0.713 | 0.006551585 |
| ENSG00000270055 | | 168.140 | -0.714 | 0.000117489 |
| ENSG00000198113 | TOR4A | 1299.986 | -0.715 | 0.000407273 |
| ENSG00000007516 | BAIAP3 | 790.966 | -0.715 | 0.009583872 |
| ENSG00000261487 | | 13.959 | -0.715 | 0.000166399 |
| ENSG00000240038 | AMY2B | 205.972 | -0.715 | 0.009208457 |
| ENSG00000146285 | SCML4 | 43.113 | -0.715 | 0.036349236 |
| ENSG00000277901 | | 10.924 | -0.716 | 0.029209712 |
| ENSG00000116774 | OLFML3 | 1379.110 | -0.717 | 0.022389278 |
| ENSG00000262503 | | 11.045 | -0.717 | 0.041394591 |
| ENSG00000090382 | LYZ | 4613.961 | -0.717 | 0.04569585 |
| ENSG00000166825 | ANPEP | 1365.991 | -0.718 | 0.03706252 |
| ENSG00000261578 | | 164.718 | -0.719 | 0.001429015 |
| ENSG00000117154 | IGSF21 | 90.300 | -0.719 | 0.029474329 |
| ENSG00000175482 | POLD4 | 1157.897 | -0.721 | 2.91079E-11 |
| ENSG00000116176 | TPSG1 | 9.192 | -0.722 | 0.021419975 |

| | | | | |
|---|---|---|---|---|
| ENSG00000224577 | LINC01117 | 11.206 | -0.722 | 0.016788229 |
| ENSG00000278621 | THBS1-AS1 | 9.318 | -0.723 | 0.037667144 |
| ENSG00000261707 | | 4.616 | -0.723 | 0.009759467 |
| ENSG00000165152 | PGAP4 | 517.046 | -0.725 | 0.015368163 |
| ENSG00000224389 | C4B | 333.388 | -0.725 | 0.046620045 |
| ENSG00000204338 | CYP21A1P | 12.291 | -0.725 | 0.037697163 |
| ENSG00000068079 | IFI35 | 2613.267 | -0.725 | 0.000149759 |
| ENSG00000186564 | FOXD2 | 119.082 | -0.725 | 0.002272737 |
| ENSG00000256139 | | 7.951 | -0.726 | 0.004232104 |
| ENSG00000237512 | UNC5B-AS1 | 50.462 | -0.727 | 0.009236506 |
| ENSG00000070526 | ST6GALNAC1 | 468.503 | -0.727 | 0.038756123 |
| ENSG00000107738 | VSIR | 2734.496 | -0.728 | 1.14229E-05 |
| ENSG00000204228 | HSD17B8 | 697.527 | -0.728 | 1.9279E-07 |
| ENSG00000260618 | | 21.541 | -0.728 | 0.012252645 |
| ENSG00000144045 | DQX1 | 399.992 | -0.729 | 0.008402367 |
| ENSG00000233093 | LINC00892 | 10.448 | -0.729 | 0.028474434 |
| ENSG00000222017 | | 15.113 | -0.729 | 0.010570949 |
| ENSG00000275719 | | 25.931 | -0.730 | 5.57872E-06 |
| ENSG00000184313 | MROH7 | 27.959 | -0.730 | 0.008423771 |
| ENSG00000063660 | GPC1 | 11370.786 | -0.731 | 9.97149E-05 |
| ENSG00000215302 | | 20.151 | -0.731 | 0.000159469 |
| ENSG00000128594 | LRRC4 | 98.107 | -0.731 | 0.012039406 |
| ENSG00000257038 | | 20.637 | -0.733 | 0.000188327 |
| ENSG00000244734 | HBB | 816.597 | -0.734 | 0.028674287 |
| ENSG00000244731 | C4A | 266.538 | -0.734 | 0.030519763 |
| ENSG00000197766 | CFD | 1076.408 | -0.735 | 0.011131934 |
| ENSG00000119943 | PYROXD2 | 563.392 | -0.735 | 0.00041157 |
| ENSG00000157873 | TNFRSF14 | 3229.278 | -0.735 | 7.4144E-08 |
| ENSG00000175318 | GRAMD2A | 263.319 | -0.736 | 0.048239436 |
| ENSG00000223704 | LINC01422 | 10.765 | -0.736 | 0.027904144 |
| ENSG00000110852 | CLEC2B | 1218.111 | -0.736 | 0.012373103 |
| ENSG00000130720 | FIBCD1 | 79.569 | -0.737 | 0.039513731 |
| ENSG00000267530 | LINC01836 | 21.577 | -0.737 | 0.047596425 |
| ENSG00000214145 | LINC00887 | 28.519 | -0.741 | 0.018259119 |
| ENSG00000177409 | SAMD9L | 1569.670 | -0.742 | 0.004998619 |
| ENSG00000273289 | | 4.522 | -0.742 | 0.012160919 |
| ENSG00000164850 | GPER1 | 83.802 | -0.742 | 0.001045205 |
| ENSG00000175147 | TMEM51-AS1 | 446.736 | -0.743 | 0.004593125 |
| ENSG00000224478 | | 4.866 | -0.745 | 0.025223667 |
| ENSG00000161714 | PLCD3 | 5367.016 | -0.745 | 0.000902417 |
| ENSG00000222898 | RN7SKP97 | 4.615 | -0.746 | 0.007211234 |
| ENSG00000142765 | SYTL1 | 5891.600 | -0.747 | 0.000103734 |
| ENSG00000167964 | RAB26 | 117.242 | -0.747 | 0.005372573 |
| ENSG00000178015 | GPR150 | 22.097 | -0.748 | 0.002549562 |
| ENSG00000226051 | ZNF503-AS1 | 150.895 | -0.750 | 0.019079182 |
| ENSG00000144488 | ESPNL | 40.377 | -0.750 | 0.028478059 |
| ENSG00000213214 | ARHGEF35 | 280.666 | -0.750 | 0.000258562 |
| ENSG00000019102 | VSIG2 | 5914.634 | -0.751 | 0.049764081 |
| ENSG00000184731 | FAM110C | 2125.559 | -0.752 | 0.001662917 |
| ENSG00000167711 | SERPINF2 | 286.194 | -0.753 | 0.004614475 |
| ENSG00000258689 | LINC01269 | 17.186 | -0.753 | 0.037737205 |
| ENSG00000211689 | TRGC1 | 17.011 | -0.753 | 0.029176113 |
| ENSG00000007933 | FMO3 | 106.242 | -0.754 | 0.030536542 |
| ENSG00000251429 | AIDAP2 | 9.584 | -0.754 | 0.010038854 |
| ENSG00000127954 | STEAP4 | 1179.260 | -0.755 | 0.021482558 |
| ENSG00000176678 | FOXL1 | 348.729 | -0.756 | 0.027213636 |
| ENSG00000130303 | BST2 | 10514.486 | -0.756 | 0.004223038 |
| ENSG00000224666 | ETV7-AS1 | 9.079 | -0.756 | 0.001453916 |
| ENSG00000274322 | | 15.882 | -0.757 | 0.002287093 |

| | | | | |
|---|---|---|---|---|
| ENSG00000180316 | PNPLA1 | 45.694 | -0.757 | 0.028391147 |
| ENSG00000122223 | CD244 | 85.174 | -0.758 | 0.016294478 |
| ENSG00000163406 | SLC15A2 | 621.351 | -0.760 | 0.000756991 |
| ENSG00000143365 | RORC | 439.953 | -0.761 | 0.044943163 |
| ENSG00000088386 | SLC15A1 | 749.113 | -0.762 | 0.02964135 |
| ENSG00000182107 | TMEM30B | 3002.237 | -0.765 | 3.67319E-06 |
| ENSG00000277496 | | 89.942 | -0.765 | 0.001551815 |
| ENSG00000170577 | SIX2 | 629.311 | -0.765 | 0.038457463 |
| ENSG00000230795 | HLA-K | 201.687 | -0.766 | 0.00254431 |
| ENSG00000171236 | LRG1 | 1766.636 | -0.766 | 0.001444764 |
| ENSG00000276409 | CCL14 | 38.222 | -0.768 | 0.048879164 |
| ENSG00000075275 | CELSR1 | 3137.979 | -0.769 | 0.00120993 |
| ENSG00000240859 | | 78.783 | -0.770 | 0.009279963 |
| ENSG00000023909 | GCLM | 2671.726 | -0.771 | 0.000442528 |
| ENSG00000215386 | MIR99AHG | 47.577 | -0.771 | 0.032932265 |
| ENSG00000143416 | SELENBP1 | 1960.148 | -0.773 | 0.004999377 |
| ENSG00000249092 | PPIAP77 | 6.385 | -0.774 | 0.012921032 |
| ENSG00000272729 | | 6.789 | -0.774 | 0.029066676 |
| ENSG00000169435 | RASSF6 | 1058.513 | -0.774 | 0.004933013 |
| ENSG00000237380 | HOXD-AS2 | 84.946 | -0.774 | 0.002044279 |
| ENSG00000130592 | LSP1 | 4447.357 | -0.774 | 0.009132927 |
| ENSG00000225969 | ABHD11-AS1 | 158.280 | -0.774 | 0.024019511 |
| ENSG00000116194 | ANGPTL1 | 112.441 | -0.775 | 0.040220795 |
| ENSG00000269646 | | 56.700 | -0.775 | 0.011193221 |
| ENSG00000168743 | NPNT | 1057.103 | -0.775 | 0.008830455 |
| ENSG00000188171 | ZNF626 | 971.354 | -0.776 | 0.025342539 |
| ENSG00000185442 | FAM174B | 4253.308 | -0.778 | 0.004423074 |
| ENSG00000159231 | CBR3 | 573.033 | -0.779 | 0.00023331 |
| ENSG00000178038 | ALS2CL | 4744.651 | -0.779 | 0.000113335 |
| ENSG00000161055 | SCGB3A1 | 47.036 | -0.780 | 0.045493104 |
| ENSG00000160808 | MYL3 | 8.731 | -0.780 | 0.028459443 |
| ENSG00000184860 | SDR42E1 | 673.768 | -0.781 | 0.002991571 |
| ENSG00000130988 | RGN | 61.397 | -0.781 | 0.016078075 |
| ENSG00000133710 | SPINK5 | 815.950 | -0.782 | 0.031456188 |
| ENSG00000148926 | ADM | 3304.998 | -0.782 | 0.000780626 |
| ENSG00000152137 | HSPB8 | 2936.692 | -0.782 | 0.012678002 |
| ENSG00000232682 | | 5.584 | -0.784 | 0.005912796 |
| ENSG00000272144 | | 35.374 | -0.784 | 4.49701E-05 |
| ENSG00000235821 | IFITM4P | 5.942 | -0.784 | 0.035342844 |
| ENSG00000272551 | | 5.335 | -0.784 | 0.045490103 |
| ENSG00000099953 | MMP11 | 9054.746 | -0.787 | 0.03398285 |
| ENSG00000211445 | GPX3 | 4790.058 | -0.787 | 0.012819448 |
| ENSG00000111846 | GCNT2 | 374.852 | -0.787 | 0.003450813 |
| ENSG00000042980 | ADAM28 | 548.589 | -0.787 | 0.006091081 |
| ENSG00000184374 | COLEC10 | 16.137 | -0.788 | 0.025199256 |
| ENSG00000005243 | COPZ2 | 648.474 | -0.791 | 0.001051365 |
| ENSG00000238142 | | 135.452 | -0.791 | 0.002203441 |
| ENSG00000102174 | PHEX | 111.783 | -0.793 | 0.006313103 |
| ENSG00000244459 | | 55.366 | -0.793 | 0.000848976 |
| ENSG00000151689 | INPP1 | 1608.717 | -0.796 | 7.07128E-07 |
| ENSG00000197721 | CR1L | 10.746 | -0.797 | 0.024213378 |
| ENSG00000223855 | HRAT92 | 34.871 | -0.797 | 0.002512083 |
| ENSG00000253649 | PRSS51 | 16.800 | -0.798 | 0.029329848 |
| ENSG00000160221 | GATD3A | 78.041 | -0.798 | 0.03764381 |
| ENSG00000197191 | CYSRT1 | 425.865 | -0.799 | 0.005511409 |
| ENSG00000015520 | NPC1L1 | 18.784 | -0.799 | 0.011609725 |
| ENSG00000054219 | LY75 | 970.834 | -0.799 | 3.52704E-05 |
| ENSG00000240891 | PLCXD2 | 199.929 | -0.800 | 0.007351665 |
| ENSG00000107821 | KAZALD1 | 290.693 | -0.801 | 0.00018269 |

| | | | | |
|---|---|---|---|---|
| ENSG00000079112 | CDH17 | 20.123 | -0.801 | 0.021141309 |
| ENSG00000224272 | | 11.532 | -0.802 | 0.024826701 |
| ENSG00000172322 | CLEC12A | 187.890 | -0.802 | 0.032135196 |
| ENSG00000182179 | UBA7 | 3340.225 | -0.802 | 1.19589E-07 |
| ENSG00000108846 | ABCC3 | 4632.590 | -0.804 | 0.002721203 |
| ENSG00000139269 | INHBE | 22.308 | -0.804 | 0.000968295 |
| ENSG00000196337 | CGB7 | 64.796 | -0.804 | 0.001696992 |
| ENSG00000012124 | CD22 | 138.326 | -0.806 | 0.017281936 |
| ENSG00000165548 | TMEM63C | 80.299 | -0.807 | 0.047946797 |
| ENSG00000128218 | VPREB3 | 32.616 | -0.807 | 0.014158194 |
| ENSG00000277287 | | 109.326 | -0.809 | 0.000442528 |
| ENSG00000150750 | C11orf53 | 177.517 | -0.809 | 0.039454308 |
| ENSG00000189377 | CXCL17 | 3940.700 | -0.809 | 0.044275973 |
| ENSG00000163554 | SPTA1 | 7.635 | -0.809 | 0.049289464 |
| ENSG00000183092 | BEGAIN | 48.395 | -0.809 | 0.00310391 |
| ENSG00000243836 | WDR86-AS1 | 73.775 | -0.809 | 0.033774796 |
| ENSG00000143819 | EPHX1 | 5784.293 | -0.810 | 1.05271E-05 |
| ENSG00000224331 | | 7.553 | -0.810 | 0.000975029 |
| ENSG00000099812 | MISP | 2426.138 | -0.810 | 0.01415082 |
| ENSG00000225756 | DBH-AS1 | 53.587 | -0.812 | 0.006479847 |
| ENSG00000006534 | ALDH3B1 | 1318.914 | -0.813 | 0.000349308 |
| ENSG00000231437 | LINC01750 | 9.339 | -0.813 | 0.030567724 |
| ENSG00000164749 | HNF4G | 103.588 | -0.813 | 0.026294048 |
| ENSG00000074047 | GLI2 | 159.742 | -0.814 | 0.010027575 |
| ENSG00000153029 | MR1 | 2131.956 | -0.814 | 1.30224E-11 |
| ENSG00000271447 | MMP28 | 1956.001 | -0.816 | 0.006473298 |
| ENSG00000272382 | | 15.856 | -0.817 | 4.63725E-05 |
| ENSG00000126337 | KRT36 | 7.456 | -0.818 | 0.02642931 |
| ENSG00000144476 | ACKR3 | 3161.392 | -0.818 | 0.000998816 |
| ENSG00000105523 | FAM83E | 337.499 | -0.819 | 0.030567724 |
| ENSG00000260228 | | 12.981 | -0.819 | 0.005615065 |
| ENSG00000226944 | RNF207-AS1 | 22.339 | -0.819 | 0.000212056 |
| ENSG00000109667 | SLC2A9 | 679.665 | -0.819 | 0.000539683 |
| ENSG00000278592 | | 13.483 | -0.820 | 0.007136134 |
| ENSG00000276842 | | 21.651 | -0.820 | 0.017797464 |
| ENSG00000254109 | RBPMS-AS1 | 48.372 | -0.820 | 2.70013E-05 |
| ENSG00000112139 | MDGA1 | 466.986 | -0.821 | 0.00746112 |
| ENSG00000267612 | | 23.050 | -0.822 | 0.021602711 |
| ENSG00000281831 | HCP5B | 7.470 | -0.825 | 0.020735926 |
| ENSG00000119986 | AVPI1 | 1437.872 | -0.825 | 5.9059E-06 |
| ENSG00000123496 | IL13RA2 | 128.631 | -0.825 | 0.027848503 |
| ENSG00000257906 | LINC02156 | 4.347 | -0.825 | 0.004956352 |
| ENSG00000225978 | HAR1A | 13.339 | -0.826 | 0.002441506 |
| ENSG00000169026 | SLC49A3 | 329.445 | -0.826 | 1.40754E-05 |
| ENSG00000244219 | TMEM225B | 26.970 | -0.827 | 0.001195178 |
| ENSG00000280424 | | 13.669 | -0.827 | 0.004156027 |
| ENSG00000229393 | | 10.337 | -0.831 | 0.029205558 |
| ENSG00000261786 | | 114.696 | -0.831 | 0.049281344 |
| ENSG00000041880 | PARP3 | 1167.555 | -0.832 | 4.84745E-11 |
| ENSG00000132744 | ACY3 | 59.493 | -0.833 | 0.007043197 |
| ENSG00000140105 | WARS1 | 15762.295 | -0.834 | 0.003117218 |
| ENSG00000225193 | RPS12P26 | 11.760 | -0.834 | 0.001107675 |
| ENSG00000091513 | TF | 124.148 | -0.834 | 0.03320956 |
| ENSG00000277639 | | 214.328 | -0.834 | 0.012704779 |
| ENSG00000227619 | | 45.816 | -0.835 | 0.007174418 |
| ENSG00000225982 | | 10.393 | -0.836 | 0.005266938 |
| ENSG00000258534 | | 51.730 | -0.837 | 0.00045032 |
| ENSG00000227802 | DNAJB3 | 10.971 | -0.837 | 0.032542348 |
| ENSG00000185614 | INKA1 | 438.714 | -0.839 | 9.09285E-05 |

| | | | | |
|---|---|---|---|---|
| ENSG00000120075 | HOXB5 | 446.752 | -0.842 | 0.004324649 |
| ENSG00000163568 | AIM2 | 1107.030 | -0.842 | 0.033475489 |
| ENSG00000253508 | | 39.082 | -0.844 | 0.008203298 |
| ENSG00000220517 | ASS1P1 | 17.131 | -0.844 | 0.00737961 |
| ENSG00000267908 | ZSCAN5DP | 7.820 | -0.846 | 0.020299228 |
| ENSG00000224093 | BCAR3-AS1 | 52.076 | -0.846 | 0.008161117 |
| ENSG00000064787 | BCAS1 | 3897.274 | -0.847 | 0.028485341 |
| ENSG00000204389 | HSPA1A | 4785.410 | -0.847 | 0.001919951 |
| ENSG00000265121 | | 10.109 | -0.848 | 0.020326089 |
| ENSG00000273132 | | 60.947 | -0.849 | 0.02295239 |
| ENSG00000262884 | | 13.605 | -0.849 | 0.005500592 |
| ENSG00000103534 | TMC5 | 559.225 | -0.849 | 0.013029701 |
| ENSG00000269720 | CCDC194 | 7.071 | -0.849 | 0.013358022 |
| ENSG00000243491 | | 48.112 | -0.850 | 0.014863408 |
| ENSG00000182103 | FAM181B | 286.479 | -0.850 | 0.013523272 |
| ENSG00000251576 | LINC01267 | 4.226 | -0.851 | 0.02536967 |
| ENSG00000204882 | GPR20 | 9.379 | -0.851 | 0.004458488 |
| ENSG00000267737 | | 10.433 | -0.851 | 0.001525722 |
| ENSG00000018236 | CNTN1 | 553.528 | -0.851 | 0.047702629 |
| ENSG00000196196 | HRCT1 | 73.814 | -0.851 | 0.004741265 |
| ENSG00000147168 | IL2RG | 1526.822 | -0.852 | 0.005046957 |
| ENSG00000123454 | DBH | 14.384 | -0.852 | 0.00611499 |
| ENSG00000257335 | MGAM | 65.971 | -0.852 | 0.019252746 |
| ENSG00000112299 | VNN1 | 79.973 | -0.853 | 0.019776792 |
| ENSG00000071909 | MYO3B | 225.882 | -0.854 | 0.030662342 |
| ENSG00000259030 | FPGT-TNNI3K | 18.173 | -0.854 | 0.001275465 |
| ENSG00000255122 | | 6.910 | -0.856 | 0.012110095 |
| ENSG00000155269 | GPR78 | 511.514 | -0.856 | 0.031780858 |
| ENSG00000233695 | GAS6-AS1 | 441.820 | -0.858 | 0.000923498 |
| ENSG00000198774 | RASSF9 | 121.823 | -0.858 | 0.023815806 |
| ENSG00000142494 | SLC47A1 | 76.350 | -0.858 | 0.005204925 |
| ENSG00000169245 | CXCL10 | 3201.562 | -0.859 | 0.041130299 |
| ENSG00000237254 | TRBV30 | 27.962 | -0.859 | 0.041635702 |
| ENSG00000172116 | CD8B | 157.476 | -0.859 | 0.011425263 |
| ENSG00000230148 | HOXB-AS1 | 68.786 | -0.863 | 4.63725E-05 |
| ENSG00000166278 | C2 | 1717.735 | -0.864 | 0.001136712 |
| ENSG00000260868 | LINC01960 | 5.238 | -0.865 | 0.011609725 |
| ENSG00000174788 | PCP2 | 74.572 | -0.865 | 0.002122803 |
| ENSG00000184106 | TREML3P | 10.080 | -0.865 | 0.03956445 |
| ENSG00000183971 | NPW | 24.724 | -0.865 | 0.012213657 |
| ENSG00000100031 | GGT1 | 301.931 | -0.866 | 0.000282078 |
| ENSG00000137270 | GCM1 | 7.225 | -0.867 | 0.022219648 |
| ENSG00000139910 | NOVA1 | 72.633 | -0.867 | 0.009276268 |
| ENSG00000079393 | DUSP13 | 14.314 | -0.870 | 0.02602357 |
| ENSG00000106819 | ASPN | 995.316 | -0.871 | 0.011658528 |
| ENSG00000263961 | RHEX | 548.257 | -0.871 | 0.042293717 |
| ENSG00000102359 | SRPX2 | 1773.057 | -0.875 | 2.21678E-05 |
| ENSG00000160870 | CYP3A7 | 65.111 | -0.875 | 0.007577008 |
| ENSG00000163993 | S100P | 34164.681 | -0.876 | 0.007534797 |
| ENSG00000196754 | S100A2 | 41510.838 | -0.877 | 0.033694492 |
| ENSG00000228236 | TXNP5 | 5.296 | -0.878 | 0.015466871 |
| ENSG00000230490 | | 6.883 | -0.879 | 0.005180831 |
| ENSG00000231106 | LINC01436 | 79.456 | -0.879 | 0.037352304 |
| ENSG00000251359 | WWC2-AS2 | 17.115 | -0.880 | 7.66976E-06 |
| ENSG00000205755 | CRLF2 | 7.413 | -0.880 | 0.00561624 |
| ENSG00000235649 | MXRA5Y | 41.641 | -0.881 | 0.035540943 |
| ENSG00000122367 | LDB3 | 81.405 | -0.883 | 0.000210527 |
| ENSG00000148346 | LCN2 | 7978.608 | -0.884 | 0.04377522 |
| ENSG00000219607 | PPP1R3G | 324.411 | -0.884 | 3.02029E-05 |

| | | | | |
|---|---|---|---|---|
| ENSG00000233110 | | 7.595 | -0.884 | 0.022946159 |
| ENSG00000175287 | PHYHD1 | 313.518 | -0.885 | 0.002784979 |
| ENSG00000181634 | TNFSF15 | 700.527 | -0.885 | 0.000708661 |
| ENSG00000169583 | CLIC3 | 2935.629 | -0.885 | 0.016506388 |
| ENSG00000272463 | | 37.441 | -0.886 | 4.98848E-05 |
| ENSG00000214318 | ATP5MC1P6 | 5.738 | -0.886 | 0.040396255 |
| ENSG00000126233 | SLURP1 | 26.130 | -0.887 | 0.030180388 |
| ENSG00000274447 | | 7.725 | -0.887 | 0.02748607 |
| ENSG00000123999 | INHA | 58.631 | -0.888 | 0.002068673 |
| ENSG00000132321 | IQCA1 | 227.888 | -0.889 | 0.0117034 |
| ENSG00000212658 | KRTAP29-1 | 3.700 | -0.890 | 0.038371061 |
| ENSG00000135709 | KIAA0513 | 834.170 | -0.892 | 2.78823E-08 |
| ENSG00000064270 | ATP2C2 | 1022.266 | -0.892 | 0.001977462 |
| ENSG00000233217 | MROH3P | 85.951 | -0.893 | 0.009078732 |
| ENSG00000161640 | SIGLEC11 | 17.228 | -0.894 | 0.007275124 |
| ENSG00000240505 | TNFRSF13B | 25.478 | -0.894 | 0.039558502 |
| ENSG00000143867 | OSR1 | 1241.490 | -0.895 | 0.004024291 |
| ENSG00000087085 | ACHE | 197.967 | -0.896 | 0.002616109 |
| ENSG00000189325 | BNIP5 | 32.954 | -0.898 | 0.043014945 |
| ENSG00000224016 | | 3.206 | -0.899 | 0.013530427 |
| ENSG00000183671 | GPR1 | 92.535 | -0.900 | 0.021656392 |
| ENSG00000154342 | WNT3A | 27.825 | -0.900 | 0.030998535 |
| ENSG00000109819 | PPARGC1A | 65.155 | -0.900 | 0.035893849 |
| ENSG00000211745 | TRBV4-2 | 9.252 | -0.901 | 0.048331708 |
| ENSG00000275385 | CCL18 | 1249.983 | -0.901 | 0.031840366 |
| ENSG00000147576 | ADHFE1 | 175.391 | -0.902 | 0.000453787 |
| ENSG00000157168 | NRG1 | 434.205 | -0.902 | 0.040193089 |
| ENSG00000250413 | SLC2A9-AS1 | 15.547 | -0.904 | 0.021207591 |
| ENSG00000173705 | SUSD5 | 100.830 | -0.905 | 0.003439465 |
| ENSG00000257831 | | 5.187 | -0.905 | 0.001281355 |
| ENSG00000244476 | ERVFRD-1 | 9.617 | -0.906 | 0.023868362 |
| ENSG00000173662 | TAS1R1 | 8.372 | -0.906 | 0.005046957 |
| ENSG00000152785 | BMP3 | 1972.556 | -0.906 | 0.044248175 |
| ENSG00000188015 | S100A3 | 255.820 | -0.909 | 0.001226486 |
| ENSG00000136872 | ALDOB | 19.198 | -0.909 | 0.04836031 |
| ENSG00000260604 | | 34.642 | -0.911 | 0.020652649 |
| ENSG00000277893 | SRD5A2 | 33.165 | -0.914 | 0.03741573 |
| ENSG00000124875 | CXCL6 | 370.792 | -0.915 | 0.036278706 |
| ENSG00000168350 | DEGS2 | 1109.988 | -0.916 | 0.006067411 |
| ENSG00000189334 | S100A14 | 13103.965 | -0.916 | 0.005073197 |
| ENSG00000175938 | ORAI3 | 898.760 | -0.916 | 7.1424E-11 |
| ENSG00000060718 | COL11A1 | 1915.116 | -0.916 | 0.035950757 |
| ENSG00000260908 | | 4.819 | -0.916 | 0.025488061 |
| ENSG00000092068 | SLC7A8 | 3179.083 | -0.917 | 1.46264E-05 |
| ENSG00000170356 | OR2A20P | 11.653 | -0.920 | 0.022605204 |
| ENSG00000102970 | CCL17 | 91.964 | -0.920 | 0.001473449 |
| ENSG00000225434 | LINC01504 | 34.302 | -0.921 | 0.000212941 |
| ENSG00000255364 | SMILR | 26.860 | -0.921 | 0.014573596 |
| ENSG00000114251 | WNT5A | 4475.541 | -0.921 | 0.003198421 |
| ENSG00000007171 | NOS2 | 127.652 | -0.922 | 0.009781321 |
| ENSG00000216921 | FAM240C | 4.228 | -0.922 | 0.023492343 |
| ENSG00000010030 | ETV7 | 840.670 | -0.923 | 5.31544E-06 |
| ENSG00000162779 | AXDND1 | 83.418 | -0.925 | 0.036974612 |
| ENSG00000236915 | CLCA4-AS1 | 7.225 | -0.925 | 0.015576053 |
| ENSG00000066405 | CLDN18 | 33.662 | -0.925 | 0.011109832 |
| ENSG00000149527 | PLCH2 | 2134.289 | -0.925 | 0.001365203 |
| ENSG00000107984 | DKK1 | 1643.810 | -0.926 | 0.018714745 |
| ENSG00000116661 | FBXO2 | 645.713 | -0.926 | 0.002000985 |
| ENSG00000263718 | SEPTIN9-DT | 124.508 | -0.927 | 0.001107038 |

| | | | | |
|---|---|---|---|---|
| ENSG00000197921 | HES5 | 20.412 | -0.927 | 0.004295218 |
| ENSG00000238042 | LINC02257 | 19.266 | -0.929 | 0.038545981 |
| ENSG00000233916 | ZDHHC20P1 | 4.387 | -0.929 | 0.019794813 |
| ENSG00000149043 | SYT8 | 7645.500 | -0.930 | 0.015190061 |
| ENSG00000171227 | TMEM37 | 527.828 | -0.933 | 5.66402E-05 |
| ENSG00000101425 | BPI | 8.009 | -0.935 | 0.013050179 |
| ENSG00000166816 | LDHD | 449.264 | -0.936 | 0.000378686 |
| ENSG00000268108 | | 4.464 | -0.936 | 0.004539344 |
| ENSG00000204642 | HLA-F | 5181.631 | -0.939 | 0.000175162 |
| ENSG00000162931 | TRIM17 | 454.599 | -0.939 | 0.002121097 |
| ENSG00000160973 | FOXH1 | 58.899 | -0.940 | 0.004898627 |
| ENSG00000185499 | MUC1 | 13998.448 | -0.941 | 0.00254431 |
| ENSG00000101463 | SYNDIG1 | 135.715 | -0.943 | 0.015474172 |
| ENSG00000168505 | GBX2 | 6.731 | -0.943 | 0.021611668 |
| ENSG00000204930 | FAM221B | 7.877 | -0.944 | 0.001897864 |
| ENSG00000197181 | PIWIL2 | 64.519 | -0.945 | 0.021331949 |
| ENSG00000260337 | | 239.640 | -0.946 | 0.037600321 |
| ENSG00000102962 | CCL22 | 368.004 | -0.946 | 0.000865638 |
| ENSG00000145945 | FAM50B | 244.000 | -0.946 | 0.000328711 |
| ENSG00000271763 | | 6.364 | -0.947 | 0.014070757 |
| ENSG00000258414 | | 10.700 | -0.948 | 0.010700931 |
| ENSG00000253417 | LINC02159 | 22.836 | -0.948 | 0.035571246 |
| ENSG00000258661 | | 13.985 | -0.948 | 0.048144226 |
| ENSG00000260528 | FAM157C | 23.098 | -0.949 | 0.000146642 |
| ENSG00000182950 | ODF3L1 | 21.229 | -0.949 | 0.000317181 |
| ENSG00000176153 | GPX2 | 13868.997 | -0.949 | 0.017607909 |
| ENSG00000230882 | | 53.948 | -0.950 | 0.000893288 |
| ENSG00000249641 | HOXC13-AS | 90.178 | -0.953 | 0.013208878 |
| ENSG00000221340 | RNU6ATAC18P | 6.756 | -0.954 | 0.001952537 |
| ENSG00000085741 | WNT11 | 283.270 | -0.956 | 0.017406829 |
| ENSG00000158458 | NRG2 | 84.908 | -0.959 | 0.004293855 |
| ENSG00000111275 | ALDH2 | 2727.624 | -0.959 | 0.000144486 |
| ENSG00000225972 | MTND1P23 | 140.425 | -0.960 | 0.016968349 |
| ENSG00000187510 | PLEKHG7 | 44.844 | -0.960 | 0.006231359 |
| ENSG00000164342 | TLR3 | 489.122 | -0.961 | 5.07673E-06 |
| ENSG00000230387 | | 192.602 | -0.963 | 0.021419975 |
| ENSG00000276603 | | 21.279 | -0.964 | 0.000332808 |
| ENSG00000224034 | LINC02561 | 4.747 | -0.965 | 0.017565475 |
| ENSG00000204572 | KRTAP5-10 | 60.246 | -0.965 | 0.026393081 |
| ENSG00000255760 | LINC02422 | 5.857 | -0.966 | 0.028187556 |
| ENSG00000278389 | | 37.246 | -0.969 | 6.97005E-05 |
| ENSG00000186453 | FAM228A | 11.115 | -0.970 | 0.004954393 |
| ENSG00000130487 | KLHDC7B | 5602.884 | -0.970 | 0.018395633 |
| ENSG00000125998 | FAM83C | 789.076 | -0.971 | 0.015349349 |
| ENSG00000228561 | | 6.947 | -0.971 | 0.018603957 |
| ENSG00000131095 | GFAP | 22.579 | -0.971 | 0.00291579 |
| ENSG00000186399 | GOLGA8R | 6.741 | -0.972 | 9.4241E-05 |
| ENSG00000253414 | LINC01605 | 49.359 | -0.972 | 0.004412094 |
| ENSG00000162496 | DHRS3 | 8033.081 | -0.972 | 9.41597E-07 |
| ENSG00000180801 | ARSJ | 242.369 | -0.973 | 0.001444764 |
| ENSG00000164220 | F2RL2 | 121.794 | -0.975 | 0.003474573 |
| ENSG00000278385 | | 29.935 | -0.975 | 1.01387E-05 |
| ENSG00000158050 | DUSP2 | 1797.020 | -0.975 | 0.000464296 |
| ENSG00000224063 | | 25.002 | -0.975 | 0.015723247 |
| ENSG00000261039 | LINC02544 | 20.499 | -0.976 | 0.017751933 |
| ENSG00000234465 | PINLYP | 118.551 | -0.976 | 0.000132685 |
| ENSG00000134121 | CHL1 | 393.657 | -0.976 | 0.02483967 |
| ENSG00000189316 | | 6.784 | -0.977 | 0.011890145 |
| ENSG00000269526 | ERVV-1 | 63.932 | -0.978 | 0.039654592 |

| | | | | |
|---|---|---|---|---|
| ENSG00000144908 | ALDH1L1 | 2515.205 | -0.979 | 0.006003907 |
| ENSG00000136514 | RTP4 | 676.689 | -0.979 | 9.05106E-06 |
| ENSG00000175920 | DOK7 | 310.072 | -0.979 | 0.000366838 |
| ENSG00000256069 | A2MP1 | 12.441 | -0.980 | 0.002620061 |
| ENSG00000167483 | NIBAN3 | 39.918 | -0.982 | 0.002044279 |
| ENSG00000174123 | TLR10 | 40.365 | -0.982 | 0.002506598 |
| ENSG00000128645 | HOXD1 | 48.056 | -0.983 | 0.002251896 |
| ENSG00000164761 | TNFRSF11B | 280.394 | -0.983 | 0.004423477 |
| ENSG00000074410 | CA12 | 5970.996 | -0.983 | 0.000751914 |
| ENSG00000160868 | CYP3A4 | 6.473 | -0.986 | 0.013089396 |
| ENSG00000255303 | OR5BA1P | 5.179 | -0.987 | 0.007092112 |
| ENSG00000211652 | IGLV7-43 | 88.051 | -0.992 | 0.039908366 |
| ENSG00000236671 | PRKG1-AS1 | 7.774 | -0.993 | 0.003149589 |
| ENSG00000168490 | PHYHIP | 206.981 | -0.994 | 0.002218196 |
| ENSG00000273340 | MICE | 24.639 | -0.997 | 0.000451931 |
| ENSG00000181240 | SLC25A41 | 18.855 | -0.997 | 0.001822994 |
| ENSG00000250596 | | 21.732 | -0.998 | 0.014308309 |
| ENSG00000110848 | CD69 | 354.506 | -0.998 | 0.001830652 |
| ENSG00000128438 | TBC1D27P | 15.872 | -0.999 | 0.017869892 |
| ENSG00000180176 | TH | 553.512 | -0.999 | 0.041530668 |
| ENSG00000225950 | NTF4 | 222.298 | -1.000 | 0.000162566 |
| ENSG00000140279 | DUOX2 | 3617.783 | -1.000 | 0.008159685 |
| ENSG00000196878 | LAMB3 | 17654.257 | -1.001 | 0.000404678 |
| ENSG00000138311 | ZNF365 | 304.595 | -1.002 | 0.002499926 |
| ENSG00000184564 | SLITRK6 | 3719.612 | -1.003 | 0.005177388 |
| ENSG00000224649 | | 3.789 | -1.005 | 0.010358329 |
| ENSG00000188039 | NWD1 | 41.890 | -1.007 | 0.005753499 |
| ENSG00000186466 | AQP7P1 | 3.889 | -1.009 | 0.015084808 |
| ENSG00000279205 | | 3.923 | -1.010 | 0.001021226 |
| ENSG00000204889 | KRT40 | 11.291 | -1.011 | 0.019776792 |
| ENSG00000234869 | | 50.707 | -1.011 | 6.33832E-05 |
| ENSG00000204622 | HLA-J | 217.885 | -1.012 | 5.54096E-05 |
| ENSG00000188037 | CLCN1 | 33.943 | -1.013 | 0.000483876 |
| ENSG00000260710 | | 73.461 | -1.013 | 0.012241925 |
| ENSG00000152580 | IGSF10 | 85.850 | -1.013 | 0.017002862 |
| ENSG00000138449 | SLC40A1 | 2839.790 | -1.015 | 9.54017E-06 |
| ENSG00000171385 | KCND3 | 278.626 | -1.016 | 0.000927117 |
| ENSG00000271392 | | 13.215 | -1.017 | 0.010253508 |
| ENSG00000198807 | PAX9 | 281.095 | -1.018 | 0.000210464 |
| ENSG00000180438 | TPRXL | 338.392 | -1.018 | 0.011013365 |
| ENSG00000151632 | AKR1C2 | 11346.945 | -1.020 | 0.009855743 |
| ENSG00000196979 | GPRACR | 5.521 | -1.023 | 0.024706446 |
| ENSG00000239605 | STPG4 | 44.183 | -1.023 | 0.010798607 |
| ENSG00000230107 | | 7.752 | -1.025 | 0.003292175 |
| ENSG00000232328 | | 7.743 | -1.027 | 0.049770721 |
| ENSG00000166819 | PLIN1 | 115.662 | -1.027 | 0.005337512 |
| ENSG00000270765 | GAS2L2 | 36.961 | -1.027 | 0.008960967 |
| ENSG00000102854 | MSLN | 1078.895 | -1.029 | 0.043679975 |
| ENSG00000227028 | SLC8A1-AS1 | 22.014 | -1.030 | 0.005932267 |
| ENSG00000164303 | ENPP6 | 23.465 | -1.031 | 0.001319023 |
| ENSG00000003436 | TFPI | 1682.110 | -1.031 | 5.84617E-06 |
| ENSG00000121380 | BCL2L14 | 202.722 | -1.031 | 6.11148E-05 |
| ENSG00000238755 | LINC02006 | 8.799 | -1.032 | 0.004214702 |
| ENSG00000124678 | TCP11 | 14.174 | -1.033 | 0.015803674 |
| ENSG00000106031 | HOXA13 | 770.347 | -1.034 | 6.91993E-05 |
| ENSG00000232679 | LINC01705 | 18.024 | -1.035 | 0.026755191 |
| ENSG00000156968 | MPV17L | 218.947 | -1.036 | 0.000848976 |
| ENSG00000262920 | | 5.895 | -1.037 | 0.01706861 |
| ENSG00000214711 | CAPN14 | 115.199 | -1.037 | 0.000859454 |

| | | | | |
|---|---|---|---|---|
| ENSG00000244345 | | 5.702 | -1.038 | 0.008865358 |
| ENSG00000259663 | | 7.494 | -1.039 | 0.026836222 |
| ENSG00000127083 | OMD | 63.423 | -1.039 | 0.029329848 |
| ENSG00000259881 | | 15.759 | -1.040 | 0.006926761 |
| ENSG00000127129 | EDN2 | 305.185 | -1.040 | 0.00618478 |
| ENSG00000231226 | TRIM31-AS1 | 23.836 | -1.043 | 0.001576413 |
| ENSG00000260581 | | 48.680 | -1.044 | 0.012198694 |
| ENSG00000211892 | IGHG4 | 7430.001 | -1.045 | 0.020565541 |
| ENSG00000123843 | C4BPB | 38.032 | -1.046 | 0.017716997 |
| ENSG00000099834 | CDHR5 | 58.071 | -1.046 | 0.007251511 |
| ENSG00000261064 | LINC02256 | 6.868 | -1.048 | 1.34219E-05 |
| ENSG00000254154 | CRYZL2P-SEC16E | 19.759 | -1.049 | 8.81513E-06 |
| ENSG00000168334 | XIRP1 | 68.687 | -1.049 | 0.008320509 |
| ENSG00000215267 | AKR1C7P | 7.035 | -1.049 | 0.019903172 |
| ENSG00000232653 | GOLGA8N | 37.613 | -1.050 | 3.97716E-08 |
| ENSG00000114854 | TNNC1 | 504.886 | -1.050 | 0.004654603 |
| ENSG00000207980 | MIR23A | 5.026 | -1.050 | 0.001826135 |
| ENSG00000171060 | C8orf74 | 17.214 | -1.051 | 0.023350042 |
| ENSG00000255571 | MIR9-3HG | 183.182 | -1.052 | 0.002236562 |
| ENSG00000198125 | MB | 211.299 | -1.053 | 0.010067738 |
| ENSG00000017427 | IGF1 | 79.084 | -1.054 | 0.005562571 |
| ENSG00000010310 | GIPR | 155.501 | -1.054 | 2.0367E-05 |
| ENSG00000125775 | SDCBP2 | 1818.052 | -1.055 | 9.22375E-06 |
| ENSG00000138755 | CXCL9 | 3035.863 | -1.057 | 0.009585741 |
| ENSG00000261664 | TTC39A-AS1 | 8.128 | -1.058 | 0.000120964 |
| ENSG00000259420 | | 4.857 | -1.061 | 0.005585444 |
| ENSG00000099957 | P2RX6 | 79.475 | -1.062 | 0.000668966 |
| ENSG00000228509 | | 5.757 | -1.064 | 0.006329131 |
| ENSG00000147206 | NXF3 | 18.774 | -1.064 | 0.001142038 |
| ENSG00000268941 | LINC01711 | 26.160 | -1.065 | 0.010876174 |
| ENSG00000257556 | LINC02298 | 88.042 | -1.066 | 1.02112E-06 |
| ENSG00000177455 | CD19 | 101.976 | -1.067 | 0.005516149 |
| ENSG00000134443 | GRP | 30.591 | -1.068 | 0.049880037 |
| ENSG00000203709 | MIR29B2CHG | 498.232 | -1.072 | 0.000659171 |
| ENSG00000187912 | CLEC17A | 12.601 | -1.074 | 0.015170702 |
| ENSG00000236256 | DIAPH2-AS1 | 17.585 | -1.075 | 0.000151771 |
| ENSG00000163501 | IHH | 56.408 | -1.075 | 0.040193089 |
| ENSG00000263316 | | 11.857 | -1.075 | 0.002513718 |
| ENSG00000253348 | | 6.348 | -1.076 | 0.013129813 |
| ENSG00000120341 | SEC16B | 15.918 | -1.079 | 1.3072E-06 |
| ENSG00000117322 | CR2 | 146.404 | -1.083 | 0.041356882 |
| ENSG00000177508 | IRX3 | 1041.963 | -1.083 | 0.00163546 |
| ENSG00000225187 | | 10.239 | -1.083 | 0.001447121 |
| ENSG00000171916 | LGALS9C | 15.882 | -1.084 | 0.005700424 |
| ENSG00000183775 | KCTD16 | 35.730 | -1.085 | 0.002258268 |
| ENSG00000171798 | KNDC1 | 17.655 | -1.086 | 0.00193626 |
| ENSG00000185633 | NDUFA4L2 | 4652.506 | -1.086 | 0.00087655 |
| ENSG00000258955 | LINC00519 | 46.092 | -1.087 | 0.000620758 |
| ENSG00000186377 | CYP4X1 | 333.598 | -1.087 | 0.000511786 |
| ENSG00000167779 | IGFBP6 | 1795.843 | -1.090 | 0.000330291 |
| ENSG00000122035 | RASL11A | 317.840 | -1.092 | 1.26284E-06 |
| ENSG00000226816 | | 55.000 | -1.092 | 0.003959897 |
| ENSG00000137648 | TMPRSS4 | 9605.861 | -1.092 | 0.001796423 |
| ENSG00000223756 | TSSC2 | 32.164 | -1.093 | 9.89835E-05 |
| ENSG00000169752 | NRG4 | 172.668 | -1.094 | 4.07221E-06 |
| ENSG00000235741 | LINC02784 | 10.578 | -1.096 | 0.034014281 |
| ENSG00000151655 | ITIH2 | 31.207 | -1.097 | 0.002511113 |
| ENSG00000100253 | MIOX | 28.737 | -1.098 | 0.001712056 |
| ENSG00000101213 | PTK6 | 4065.019 | -1.102 | 8.2978E-07 |

| | | | | |
|---|---|---|---|---|
| ENSG00000258647 | LINC00930 | 267.845 | -1.105 | 0.035403977 |
| ENSG00000154263 | ABCA10 | 325.358 | -1.106 | 0.000339329 |
| ENSG00000089558 | KCNH4 | 70.352 | -1.107 | 0.000245274 |
| ENSG00000243709 | LEFTY1 | 7.410 | -1.108 | 0.0014635 |
| ENSG00000248243 | LINC02014 | 17.869 | -1.108 | 2.15639E-06 |
| ENSG00000099937 | SERPIND1 | 48.033 | -1.110 | 0.004083282 |
| ENSG00000234362 | LINC01914 | 8.502 | -1.111 | 0.006650041 |
| ENSG00000233539 | | 24.441 | -1.111 | 0.004677912 |
| ENSG00000237737 | DCTN1-AS1 | 11.728 | -1.115 | 0.001705013 |
| ENSG00000167588 | GPD1 | 111.598 | -1.116 | 0.000937292 |
| ENSG00000268628 | | 39.662 | -1.117 | 3.67744E-05 |
| ENSG00000183273 | CCDC60 | 85.796 | -1.121 | 0.018192934 |
| ENSG00000168032 | ENTPD3 | 1504.767 | -1.122 | 8.36628E-05 |
| ENSG00000256262 | USP30-AS1 | 56.417 | -1.122 | 0.000165047 |
| ENSG00000105131 | EPHX3 | 1187.142 | -1.124 | 0.002438976 |
| ENSG00000102287 | GABRE | 2762.956 | -1.125 | 0.000852505 |
| ENSG00000087495 | PHACTR3 | 95.819 | -1.126 | 0.001192227 |
| ENSG00000244137 | | 5.372 | -1.126 | 0.01028931 |
| ENSG00000101842 | VSIG1 | 120.296 | -1.129 | 0.000360948 |
| ENSG00000280604 | | 20.408 | -1.129 | 0.008547362 |
| ENSG00000114771 | AADAC | 406.769 | -1.132 | 0.025420278 |
| ENSG00000271573 | | 4.017 | -1.133 | 0.031590226 |
| ENSG00000055957 | ITIH1 | 9.638 | -1.133 | 0.016827827 |
| ENSG00000268964 | ERVV-2 | 21.599 | -1.134 | 0.035342844 |
| ENSG00000221986 | MYBPHL | 12.818 | -1.136 | 0.005400645 |
| ENSG00000205054 | LINC01121 | 13.494 | -1.136 | 0.00033029 |
| ENSG00000023839 | ABCC2 | 55.470 | -1.136 | 3.98555E-05 |
| ENSG00000261172 | | 23.604 | -1.137 | 0.00072571 |
| ENSG00000138131 | LOXL4 | 595.448 | -1.138 | 0.000378686 |
| ENSG00000250033 | SLC7A11-AS1 | 10.930 | -1.138 | 0.021207591 |
| ENSG00000156738 | MS4A1 | 165.110 | -1.139 | 0.01703249 |
| ENSG00000101017 | CD40 | 1550.483 | -1.140 | 1.22869E-07 |
| ENSG00000149922 | TBX6 | 561.287 | -1.141 | 3.44188E-07 |
| ENSG00000016082 | ISL1 | 16.288 | -1.141 | 0.015678207 |
| ENSG00000205363 | INSYN1 | 124.778 | -1.141 | 0.000781484 |
| ENSG00000180638 | SLC47A2 | 59.565 | -1.141 | 0.007293248 |
| ENSG00000232316 | LINC02518 | 26.194 | -1.141 | 0.002005333 |
| ENSG00000229754 | CXCR2P1 | 77.995 | -1.143 | 0.005400645 |
| ENSG00000175513 | TSGA10IP | 15.188 | -1.148 | 0.00545018 |
| ENSG00000120885 | CLU | 21154.076 | -1.149 | 9.93897E-05 |
| ENSG00000257042 | | 11.962 | -1.151 | 0.015512023 |
| ENSG00000240747 | KRBOX1 | 62.754 | -1.153 | 0.001397867 |
| ENSG00000254226 | LINC01933 | 21.524 | -1.154 | 0.007534797 |
| ENSG00000198099 | ADH4 | 27.975 | -1.155 | 0.009169916 |
| ENSG00000174837 | ADGRE1 | 115.751 | -1.157 | 0.002068673 |
| ENSG00000211710 | TRBV4-1 | 6.036 | -1.159 | 0.008101939 |
| ENSG00000186204 | CYP4F12 | 1767.169 | -1.159 | 0.001974591 |
| ENSG00000089250 | NOS1 | 13.008 | -1.161 | 0.005400645 |
| ENSG00000164929 | BAALC | 309.179 | -1.161 | 0.00182259 |
| ENSG00000144820 | ADGRG7 | 19.498 | -1.161 | 0.041569149 |
| ENSG00000114248 | LRRC31 | 14.465 | -1.163 | 0.003120264 |
| ENSG00000187997 | C17orf99 | 17.835 | -1.166 | 2.27867E-05 |
| ENSG00000112818 | MEP1A | 12.924 | -1.170 | 0.003661044 |
| ENSG00000130234 | ACE2 | 184.789 | -1.171 | 0.000496995 |
| ENSG00000198670 | LPA | 28.431 | -1.174 | 0.011288733 |
| ENSG00000114378 | HYAL1 | 172.186 | -1.177 | 5.9059E-06 |
| ENSG00000166143 | PPP1R14D | 143.836 | -1.180 | 0.006469576 |
| ENSG00000115718 | PROC | 119.768 | -1.180 | 1.42258E-05 |
| ENSG00000276972 | | 6.185 | -1.185 | 0.011843009 |

| | | | | |
|---|---|---|---|---|
| ENSG00000165682 | CLEC1B | 8.270 | -1.186 | 0.002502121 |
| ENSG00000217455 | | 16.049 | -1.186 | 0.008658662 |
| ENSG00000207146 | Y_RNA | 9.663 | -1.187 | 0.000554345 |
| ENSG00000144035 | NAT8 | 11.131 | -1.189 | 0.00153021 |
| ENSG00000171819 | ANGPTL7 | 14.816 | -1.191 | 0.011868879 |
| ENSG00000234184 | LINC01781 | 7.172 | -1.192 | 0.017602691 |
| ENSG00000108511 | HOXB6 | 721.906 | -1.193 | 1.07894E-05 |
| ENSG00000198842 | STYXL2 | 48.358 | -1.194 | 0.036922265 |
| ENSG00000173212 | MAB21L3 | 128.439 | -1.199 | 0.000453289 |
| ENSG00000135226 | UGT2B28 | 29.446 | -1.200 | 0.037054017 |
| ENSG00000198610 | AKR1C4 | 10.536 | -1.201 | 0.011658528 |
| ENSG00000167617 | CDC42EP5 | 1197.848 | -1.201 | 4.21488E-07 |
| ENSG00000241119 | UGT1A9 | 12.593 | -1.207 | 0.014955536 |
| ENSG00000175874 | CREG2 | 171.231 | -1.207 | 0.000103459 |
| ENSG00000256948 | | 9.956 | -1.208 | 0.00853114 |
| ENSG00000248810 | LINC02432 | 71.343 | -1.208 | 0.021532224 |
| ENSG00000169174 | PCSK9 | 374.050 | -1.212 | 0.004730388 |
| ENSG00000160180 | TFF3 | 365.698 | -1.212 | 0.006213269 |
| ENSG00000150045 | KLRF1 | 32.546 | -1.213 | 5.95024E-05 |
| ENSG00000226025 | LGALS17A | 35.015 | -1.214 | 0.01706861 |
| ENSG00000145287 | PLAC8 | 509.359 | -1.217 | 0.000200429 |
| ENSG00000168874 | ATOH8 | 669.691 | -1.221 | 0.000447013 |
| ENSG00000101144 | BMP7 | 1409.192 | -1.224 | 0.001732628 |
| ENSG00000180525 | DIP2C-AS1 | 34.591 | -1.227 | 2.05692E-05 |
| ENSG00000019186 | CYP24A1 | 3688.476 | -1.228 | 0.002166891 |
| ENSG00000136573 | BLK | 53.437 | -1.231 | 0.003578722 |
| ENSG00000260284 | TPSP2 | 118.070 | -1.231 | 0.007053175 |
| ENSG00000269404 | SPIB | 92.533 | -1.235 | 0.000600292 |
| ENSG00000135773 | CAPN9 | 188.771 | -1.236 | 0.005108165 |
| ENSG00000164530 | PI16 | 101.175 | -1.238 | 0.003540275 |
| ENSG00000068078 | FGFR3 | 13010.781 | -1.240 | 0.000114555 |
| ENSG00000261857 | MIA | 23.911 | -1.241 | 0.003840169 |
| ENSG00000198758 | EPS8L3 | 308.408 | -1.241 | 0.011860167 |
| ENSG00000162006 | MSLNL | 4.931 | -1.243 | 0.038518074 |
| ENSG00000254851 | | 50.907 | -1.243 | 0.007669961 |
| ENSG00000140961 | OSGIN1 | 584.954 | -1.243 | 8.74093E-07 |
| ENSG00000271503 | CCL5 | 3099.347 | -1.243 | 0.000141269 |
| ENSG00000258837 | | 65.787 | -1.244 | 0.013105582 |
| ENSG00000130055 | GDPD2 | 444.764 | -1.246 | 0.002062233 |
| ENSG00000196260 | SFTA2 | 67.742 | -1.248 | 0.015312948 |
| ENSG00000118785 | SPP1 | 9966.449 | -1.248 | 0.000229625 |
| ENSG00000231971 | CT69 | 15.143 | -1.249 | 0.004919394 |
| ENSG00000176083 | ZNF683 | 129.182 | -1.255 | 0.000591561 |
| ENSG00000073734 | ABCB11 | 13.266 | -1.256 | 0.003457091 |
| ENSG00000119919 | NKX2-3 | 4.404 | -1.257 | 0.035438501 |
| ENSG00000269842 | | 12.914 | -1.258 | 0.001421671 |
| ENSG00000156959 | LHFPL4 | 10.976 | -1.259 | 0.032992198 |
| ENSG00000259023 | LINC00524 | 4.709 | -1.261 | 0.035967031 |
| ENSG00000229196 | | 5.351 | -1.264 | 0.000622653 |
| ENSG00000256612 | CYP2B7P | 27.899 | -1.269 | 0.001021226 |
| ENSG00000274214 | | 6.477 | -1.270 | 0.001746383 |
| ENSG00000235947 | EGOT | 10.639 | -1.271 | 0.001607946 |
| ENSG00000176194 | CIDEA | 49.525 | -1.271 | 0.002863167 |
| ENSG00000123838 | C4BPA | 12.348 | -1.272 | 0.014230056 |
| ENSG00000244586 | WNT5A-AS1 | 51.808 | -1.273 | 4.14537E-05 |
| ENSG00000253616 | | 17.900 | -1.274 | 9.96071E-07 |
| ENSG00000242795 | | 10.230 | -1.276 | 0.002246309 |
| ENSG00000239552 | HOXB-AS2 | 16.050 | -1.277 | 0.000555618 |
| ENSG00000270937 | | 3.711 | -1.279 | 0.028115137 |

| | | | | |
|---|---|---|---|---|
| ENSG00000106541 | AGR2 | 9353.912 | -1.280 | 0.000157837 |
| ENSG00000254303 | | 3.381 | -1.281 | 0.007540833 |
| ENSG00000268287 | | 16.680 | -1.283 | 1.19112E-06 |
| ENSG00000106258 | CYP3A5 | 1362.729 | -1.288 | 0.000147164 |
| ENSG00000203797 | DDO | 100.781 | -1.288 | 2.26429E-07 |
| ENSG00000272068 | BCAN-AS1 | 181.833 | -1.289 | 0.000252001 |
| ENSG00000235491 | LINC01889 | 18.491 | -1.290 | 0.027048733 |
| ENSG00000173917 | HOXB2 | 713.390 | -1.291 | 2.70403E-07 |
| ENSG00000255325 | | 42.581 | -1.292 | 0.000341451 |
| ENSG00000215182 | MUC5AC | 110.416 | -1.295 | 0.020935259 |
| ENSG00000174343 | CHRNA9 | 8.743 | -1.297 | 0.008320509 |
| ENSG00000175164 | ABO | 1189.673 | -1.303 | 1.63767E-05 |
| ENSG00000223722 | IFITM3P2 | 18.410 | -1.303 | 0.000999633 |
| ENSG00000110092 | CCND1 | 15421.541 | -1.307 | 1.25392E-07 |
| ENSG00000096006 | CRISP3 | 217.730 | -1.314 | 0.008115606 |
| ENSG00000042832 | TG | 116.339 | -1.317 | 1.53734E-05 |
| ENSG00000204539 | CDSN | 8.971 | -1.318 | 0.039042931 |
| ENSG00000012223 | LTF | 584.267 | -1.320 | 0.001684444 |
| ENSG00000118094 | TREH | 14.434 | -1.320 | 0.000156355 |
| ENSG00000162878 | PKDCC | 438.868 | -1.321 | 0.000147164 |
| ENSG00000240602 | AADACP1 | 61.786 | -1.321 | 0.008863039 |
| ENSG00000171346 | KRT15 | 11450.532 | -1.325 | 0.000620758 |
| ENSG00000153923 | CLCA3P | 19.138 | -1.328 | 0.000323694 |
| ENSG00000094796 | KRT31 | 64.247 | -1.328 | 0.018592083 |
| ENSG00000279756 | | 6.066 | -1.329 | 0.01556842 |
| ENSG00000135744 | AGT | 303.819 | -1.330 | 2.81683E-05 |
| ENSG00000179593 | ALOX15B | 236.109 | -1.332 | 2.94515E-05 |
| ENSG00000078725 | BRINP1 | 40.817 | -1.333 | 0.00421043 |
| ENSG00000182983 | ZNF662 | 501.450 | -1.333 | 2.18586E-05 |
| ENSG00000142619 | PADI3 | 7291.323 | -1.336 | 0.001116897 |
| ENSG00000278966 | | 9.852 | -1.336 | 0.000820491 |
| ENSG00000233101 | HOXB-AS3 | 103.342 | -1.336 | 9.38206E-05 |
| ENSG00000253593 | | 15.047 | -1.352 | 0.004461576 |
| ENSG00000114948 | ADAM23 | 366.228 | -1.353 | 1.96494E-05 |
| ENSG00000138152 | BTBD16 | 2137.179 | -1.354 | 0.002263349 |
| ENSG00000137033 | IL33 | 943.821 | -1.355 | 0.000115002 |
| ENSG00000185303 | SFTPA2 | 62.116 | -1.360 | 0.008115606 |
| ENSG00000133742 | CA1 | 5.662 | -1.362 | 0.01452037 |
| ENSG00000233536 | | 4.117 | -1.363 | 0.00871537 |
| ENSG00000228100 | LINC01820 | 6.671 | -1.365 | 0.007534797 |
| ENSG00000143512 | HHIPL2 | 24.392 | -1.365 | 4.84364E-05 |
| ENSG00000164078 | MST1R | 2056.050 | -1.366 | 5.54423E-08 |
| ENSG00000205436 | EXOC3L4 | 159.728 | -1.368 | 6.14151E-06 |
| ENSG00000238164 | TNFRSF14-AS1 | 337.238 | -1.370 | 1.69149E-09 |
| ENSG00000242948 | EPS15P1 | 4.223 | -1.373 | 0.012636002 |
| ENSG00000236740 | | 44.843 | -1.375 | 0.00507392 |
| ENSG00000261083 | LINC02516 | 9.968 | -1.377 | 0.000132685 |
| ENSG00000254290 | | 26.784 | -1.385 | 0.005053462 |
| ENSG00000163534 | FCRL1 | 17.840 | -1.387 | 0.004156027 |
| ENSG00000277249 | MIR6784 | 4.788 | -1.388 | 0.000162566 |
| ENSG00000139304 | PTPRQ | 214.833 | -1.388 | 0.002921401 |
| ENSG00000227502 | MROCKI | 34.445 | -1.390 | 5.04578E-06 |
| ENSG00000206199 | ANKUB1 | 8.063 | -1.391 | 0.000764086 |
| ENSG00000161849 | KRT84 | 7.363 | -1.397 | 0.039553634 |
| ENSG00000280109 | PLAC4 | 45.319 | -1.398 | 0.000171222 |
| ENSG00000243766 | HOTTIP | 198.649 | -1.399 | 3.09872E-05 |
| ENSG00000086548 | CEACAM6 | 8098.647 | -1.403 | 0.004598212 |
| ENSG00000213886 | UBD | 496.220 | -1.406 | 0.000659171 |
| ENSG00000188822 | CNR2 | 17.080 | -1.406 | 0.00014846 |

| | | | | |
|---|---|---|---|---|
| ENSG00000091831 | ESR1 | 251.671 | -1.406 | 5.456E-07 |
| ENSG00000254233 | LINC02365 | 5.724 | -1.407 | 0.000968295 |
| ENSG00000267308 | LINC01764 | 46.193 | -1.407 | 0.001421671 |
| ENSG00000279141 | LINC01451 | 280.508 | -1.408 | 3.25797E-05 |
| ENSG00000159374 | M1AP | 60.245 | -1.413 | 0.000204132 |
| ENSG00000113924 | HGD | 20.986 | -1.413 | 3.1986E-05 |
| ENSG00000105852 | PON3 | 72.394 | -1.414 | 0.00045032 |
| ENSG00000215853 | RPTN | 18.240 | -1.418 | 0.046099607 |
| ENSG00000136059 | VILL | 1627.156 | -1.420 | 3.23582E-08 |
| ENSG00000237988 | OR2I1P | 332.565 | -1.423 | 0.000723444 |
| ENSG00000175967 | LINC02880 | 4.760 | -1.432 | 0.00050345 |
| ENSG00000198732 | SMOC1 | 246.134 | -1.434 | 0.000660435 |
| ENSG00000224961 | LINC01752 | 24.132 | -1.435 | 5.54247E-06 |
| ENSG00000256039 | LINC02446 | 55.779 | -1.436 | 0.001523372 |
| ENSG00000278484 | | 32.307 | -1.436 | 0.00853114 |
| ENSG00000100652 | SLC10A1 | 16.934 | -1.437 | 0.000803916 |
| ENSG00000260115 | | 8.234 | -1.437 | 0.001461594 |
| ENSG00000116745 | RPE65 | 12.172 | -1.439 | 0.02335417 |
| ENSG00000119457 | SLC46A2 | 49.582 | -1.440 | 4.93862E-05 |
| ENSG00000211655 | IGLV1-36 | 82.410 | -1.442 | 0.005591059 |
| ENSG00000181126 | HLA-V | 91.252 | -1.445 | 1.58725E-05 |
| ENSG00000233441 | CYP2AB1P | 5.439 | -1.455 | 0.010228544 |
| ENSG00000231574 | LINC02015 | 36.903 | -1.458 | 0.000408727 |
| ENSG00000255666 | LINC02700 | 7.095 | -1.460 | 0.009691827 |
| ENSG00000129988 | LBP | 57.028 | -1.461 | 0.000309366 |
| ENSG00000260685 | | 7.662 | -1.467 | 1.0539E-06 |
| ENSG00000103355 | PRSS33 | 7.464 | -1.467 | 0.005326481 |
| ENSG00000158816 | VWA5B1 | 121.245 | -1.468 | 0.003226091 |
| ENSG00000143839 | REN | 148.380 | -1.468 | 8.86146E-05 |
| ENSG00000255191 | | 6.965 | -1.471 | 0.002089549 |
| ENSG00000143125 | PROK1 | 11.958 | -1.473 | 0.000689544 |
| ENSG00000249307 | LINC01088 | 61.335 | -1.475 | 3.41716E-05 |
| ENSG00000253838 | | 5.987 | -1.475 | 0.012217989 |
| ENSG00000165325 | DEUP1 | 6.436 | -1.476 | 0.000998409 |
| ENSG00000231453 | LINC01305 | 10.726 | -1.478 | 0.003177429 |
| ENSG00000266634 | MIR3972 | 13.275 | -1.479 | 0.00248395 |
| ENSG00000244921 | MTCYBP18 | 43.681 | -1.481 | 0.000116204 |
| ENSG00000188263 | IL17REL | 9.160 | -1.481 | 0.000659171 |
| ENSG00000224322 | | 16.715 | -1.486 | 0.011422785 |
| ENSG00000213889 | PPM1N | 412.585 | -1.497 | 2.1248E-07 |
| ENSG00000105650 | PDE4C | 170.964 | -1.498 | 4.72394E-08 |
| ENSG00000104921 | FCER2 | 32.592 | -1.498 | 0.000852441 |
| ENSG00000205076 | LGALS7 | 72.879 | -1.503 | 0.038005498 |
| ENSG00000196660 | SLC30A10 | 13.115 | -1.508 | 0.000799739 |
| ENSG00000146453 | PNLDC1 | 18.337 | -1.511 | 5.83188E-05 |
| ENSG00000198203 | SULT1C2 | 136.219 | -1.514 | 8.43223E-06 |
| ENSG00000243955 | GSTA1 | 788.557 | -1.515 | 0.007297377 |
| ENSG00000186481 | ANKRD20A5P | 88.918 | -1.524 | 7.62521E-06 |
| ENSG00000117472 | TSPAN1 | 5966.075 | -1.527 | 4.21406E-07 |
| ENSG00000164509 | IL31RA | 34.914 | -1.527 | 0.000845957 |
| ENSG00000181143 | MUC16 | 981.277 | -1.531 | 0.003885001 |
| ENSG00000101076 | HNF4A | 36.669 | -1.531 | 0.000175104 |
| ENSG00000188385 | JAKMIP3 | 40.499 | -1.534 | 1.97857E-06 |
| ENSG00000204876 | | 19.055 | -1.536 | 0.000400159 |
| ENSG00000261175 | LINC02188 | 34.136 | -1.537 | 0.005502933 |
| ENSG00000240902 | APOOP2 | 7.108 | -1.541 | 0.010356393 |
| ENSG00000105641 | SLC5A5 | 57.446 | -1.543 | 6.4961E-05 |
| ENSG00000081051 | AFP | 6.521 | -1.548 | 0.011561027 |
| ENSG00000271579 | | 53.367 | -1.557 | 0.001516579 |

| | | | | |
|---|---|---|---|---|
| ENSG00000241935 | HOGA1 | 57.841 | -1.559 | 9.2767E-09 |
| ENSG00000271856 | LINC01215 | 37.288 | -1.561 | 1.18041E-05 |
| ENSG00000232079 | LINC01697 | 17.892 | -1.563 | 0.001409317 |
| ENSG00000169562 | GJB1 | 76.453 | -1.565 | 0.001944217 |
| ENSG00000259905 | PWRN1 | 54.776 | -1.565 | 0.013576667 |
| ENSG00000251584 | | 18.826 | -1.568 | 8.8821E-05 |
| ENSG00000266145 | RHOT1P1 | 6.578 | -1.569 | 0.000709062 |
| ENSG00000223956 | LINC01767 | 22.942 | -1.579 | 9.83658E-06 |
| ENSG00000244414 | CFHR1 | 5.279 | -1.582 | 0.002608207 |
| ENSG00000185823 | NPAP1 | 19.822 | -1.586 | 0.011762645 |
| ENSG00000236028 | | 36.868 | -1.588 | 0.008497703 |
| ENSG00000197353 | LYPD2 | 44.408 | -1.596 | 0.00254431 |
| ENSG00000253154 | | 13.042 | -1.600 | 0.023866739 |
| ENSG00000162494 | LRRC38 | 6.901 | -1.602 | 0.000536985 |
| ENSG00000213030 | CGB8 | 9.782 | -1.604 | 0.004800411 |
| ENSG00000233845 | | 10.300 | -1.605 | 0.000112495 |
| ENSG00000172927 | MYEOV | 1517.535 | -1.605 | 5.86082E-05 |
| ENSG00000215559 | ANKRD20A11P | 37.621 | -1.610 | 1.01733E-05 |
| ENSG00000257345 | LINC02413 | 5.377 | -1.610 | 0.000357313 |
| ENSG00000170477 | KRT4 | 3820.135 | -1.612 | 0.003046953 |
| ENSG00000268104 | SLC6A14 | 429.119 | -1.613 | 0.00034053 |
| ENSG00000172551 | MUCL1 | 48.447 | -1.618 | 0.000196039 |
| ENSG00000204872 | NAT8B | 35.920 | -1.618 | 2.18822E-07 |
| ENSG00000186038 | HTR3E | 4.680 | -1.620 | 0.000152513 |
| ENSG00000169894 | MUC3A | 417.953 | -1.628 | 1.47963E-05 |
| ENSG00000257322 | | 9.590 | -1.630 | 3.66538E-07 |
| ENSG00000151012 | SLC7A11 | 950.170 | -1.632 | 2.68695E-07 |
| ENSG00000181333 | HEPHL1 | 324.411 | -1.633 | 0.000364748 |
| ENSG00000100721 | TCL1A | 40.386 | -1.637 | 0.000278746 |
| ENSG00000188833 | ENTPD8 | 37.251 | -1.637 | 6.00926E-07 |
| ENSG00000164825 | DEFB1 | 740.422 | -1.638 | 2.63096E-05 |
| ENSG00000165794 | SLC39A2 | 490.876 | -1.644 | 8.26905E-05 |
| ENSG00000156510 | HKDC1 | 134.799 | -1.648 | 3.19613E-05 |
| ENSG00000260877 | | 65.815 | -1.653 | 0.000133363 |
| ENSG00000176998 | HCG4 | 30.036 | -1.655 | 3.83828E-06 |
| ENSG00000250654 | | 14.738 | -1.658 | 2.29751E-06 |
| ENSG00000279429 | | 5.469 | -1.669 | 0.005738827 |
| ENSG00000171903 | CYP4F11 | 1119.945 | -1.690 | 4.21488E-07 |
| ENSG00000224511 | LINC00365 | 22.645 | -1.694 | 1.49726E-06 |
| ENSG00000162078 | ZG16B | 518.903 | -1.698 | 1.62938E-09 |
| ENSG00000137745 | MMP13 | 1401.316 | -1.698 | 3.38048E-05 |
| ENSG00000148735 | PLEKHS1 | 350.166 | -1.698 | 1.79458E-05 |
| ENSG00000164690 | SHH | 1123.325 | -1.701 | 0.000539 |
| ENSG00000104827 | CGB3 | 9.713 | -1.702 | 0.001325558 |
| ENSG00000257017 | HP | 48.758 | -1.702 | 0.000449152 |
| ENSG00000234476 | LINC02765 | 8.407 | -1.709 | 6.61419E-05 |
| ENSG00000073067 | CYP2W1 | 249.181 | -1.711 | 5.54096E-05 |
| ENSG00000187689 | AMTN | 30.943 | -1.712 | 0.003418227 |
| ENSG00000145113 | MUC4 | 2418.465 | -1.713 | 5.82036E-05 |
| ENSG00000138615 | CILP | 840.512 | -1.713 | 3.52704E-05 |
| ENSG00000143536 | CRNN | 14.719 | -1.719 | 0.037598669 |
| ENSG00000186529 | CYP4F3 | 1033.496 | -1.720 | 3.95155E-06 |
| ENSG00000267235 | ZNF861P | 4.058 | -1.739 | 0.004095921 |
| ENSG00000125207 | PIWIL1 | 10.725 | -1.739 | 0.000235509 |
| ENSG00000160181 | TFF2 | 170.938 | -1.740 | 0.000861645 |
| ENSG00000126838 | PZP | 23.268 | -1.744 | 1.28173E-06 |
| ENSG00000253525 | CPP | 5.043 | -1.753 | 0.000797536 |
| ENSG00000166869 | CHP2 | 636.444 | -1.755 | 0.000122401 |
| ENSG00000165078 | CPA6 | 28.406 | -1.760 | 7.80894E-06 |

| | | | | |
|---|---|---|---|---|
| ENSG00000138109 | CYP2C9 | 60.209 | -1.764 | 0.0005294 |
| ENSG00000255462 | | 5.944 | -1.771 | 1.17568E-06 |
| ENSG00000134668 | SPOCD1 | 1937.263 | -1.772 | 1.07427E-09 |
| ENSG00000162897 | FCAMR | 14.367 | -1.773 | 0.000214247 |
| ENSG00000166391 | MOGAT2 | 249.049 | -1.773 | 0.000179459 |
| ENSG00000248485 | PCP4L1 | 1372.627 | -1.777 | 2.67661E-06 |
| ENSG00000165887 | ANKRD2 | 38.713 | -1.784 | 8.13135E-10 |
| ENSG00000251577 | | 8.511 | -1.788 | 0.000597615 |
| ENSG00000189052 | CGB5 | 24.538 | -1.801 | 0.000707039 |
| ENSG00000136327 | NKX2-8 | 45.172 | -1.806 | 7.92839E-05 |
| ENSG00000254545 | | 55.733 | -1.816 | 4.00536E-08 |
| ENSG00000253308 | | 93.065 | -1.818 | 0.000761426 |
| ENSG00000181577 | C6orf223 | 164.929 | -1.821 | 1.09787E-06 |
| ENSG00000135346 | CGA | 16.990 | -1.844 | 0.00150138 |
| ENSG00000211654 | IGLV5-37 | 14.483 | -1.851 | 0.0033821 |
| ENSG00000240241 | | 10.529 | -1.858 | 0.015141643 |
| ENSG00000226308 | | 10.784 | -1.862 | 0.000119631 |
| ENSG00000254287 | | 5.651 | -1.865 | 0.000785455 |
| ENSG00000275896 | PRSS2 | 284.544 | -1.870 | 0.001049597 |
| ENSG00000101441 | CST4 | 52.978 | -1.874 | 5.63081E-05 |
| ENSG00000100181 | TPTEP1 | 214.916 | -1.891 | 2.15877E-07 |
| ENSG00000134193 | REG4 | 64.713 | -1.894 | 3.22041E-06 |
| ENSG00000007306 | CEACAM7 | 350.342 | -1.899 | 0.000433052 |
| ENSG00000230257 | NFE4 | 7.517 | -1.949 | 2.67899E-05 |
| ENSG00000259087 | | 44.423 | -1.954 | 4.82336E-07 |
| ENSG00000214797 | | 4.399 | -1.956 | 7.2691E-05 |
| ENSG00000073737 | DHRS9 | 566.823 | -1.958 | 8.87615E-07 |
| ENSG00000069764 | PLA2G10 | 50.393 | -1.970 | 2.96591E-11 |
| ENSG00000178934 | LGALS7B | 209.076 | -1.971 | 0.000212406 |
| ENSG00000249628 | LINC00942 | 41.516 | -1.973 | 2.52015E-08 |
| ENSG00000153802 | TMPRSS11D | 80.980 | -1.981 | 0.000513065 |
| ENSG00000095713 | CRTAC1 | 1934.115 | -1.988 | 7.40187E-05 |
| ENSG00000140274 | DUOXA2 | 661.187 | -2.016 | 3.16138E-07 |
| ENSG00000188761 | BCL2L15 | 125.114 | -2.017 | 1.72786E-08 |
| ENSG00000271134 | IFITM3P9 | 7.272 | -2.032 | 0.001282802 |
| ENSG00000157765 | SLC34A2 | 92.228 | -2.038 | 0.000141506 |
| ENSG00000136352 | NKX2-1 | 25.313 | -2.040 | 0.003563648 |
| ENSG00000203697 | CAPN8 | 1027.767 | -2.052 | 3.50121E-07 |
| ENSG00000170323 | FABP4 | 4567.265 | -2.054 | 9.51026E-06 |
| ENSG00000228314 | CYP4F29P | 219.283 | -2.063 | 4.10851E-07 |
| ENSG00000169271 | HSPB3 | 12.924 | -2.066 | 3.12304E-05 |
| ENSG00000242908 | AADACL2-AS1 | 3.968 | -2.069 | 0.000432213 |
| ENSG00000163331 | DAPL1 | 183.825 | -2.079 | 1.54381E-05 |
| ENSG00000109511 | ANXA10 | 1699.584 | -2.083 | 0.000115487 |
| ENSG00000156298 | TSPAN7 | 753.869 | -2.087 | 3.3761E-11 |
| ENSG00000240216 | CPHL1P | 27.736 | -2.091 | 3.28335E-09 |
| ENSG00000203985 | LDLRAD1 | 7.454 | -2.102 | 8.92509E-06 |
| ENSG00000053438 | NNAT | 299.949 | -2.111 | 7.29615E-10 |
| ENSG00000187134 | AKR1C1 | 5100.197 | -2.115 | 2.19947E-09 |
| ENSG00000167080 | B4GALNT2 | 24.085 | -2.116 | 9.97149E-05 |
| ENSG00000145879 | SPINK7 | 23.035 | -2.120 | 1.46448E-05 |
| ENSG00000213606 | AKR1B10P1 | 7.413 | -2.128 | 0.000583584 |
| ENSG00000167941 | SOST | 15.569 | -2.132 | 0.000148529 |
| ENSG00000187908 | DMBT1 | 146.618 | -2.137 | 3.35783E-06 |
| ENSG00000204544 | MUC21 | 50.852 | -2.145 | 2.00152E-05 |
| ENSG00000255774 | LINC02747 | 6.225 | -2.155 | 5.77518E-05 |
| ENSG00000120094 | HOXB1 | 18.803 | -2.170 | 6.42098E-09 |
| ENSG00000131203 | IDO1 | 2393.304 | -2.171 | 6.13662E-08 |
| ENSG00000278912 | | 11.630 | -2.183 | 0.000104745 |

| | | | | |
|---|---|---|---|---|
| ENSG00000159182 | PRAC1 | 8.284 | -2.183 | 0.002856785 |
| ENSG00000171658 | NMRAL2P | 73.979 | -2.189 | 1.30698E-06 |
| ENSG00000267056 | | 107.927 | -2.190 | 1.58635E-08 |
| ENSG00000260676 | LINC01541 | 27.826 | -2.202 | 0.027275355 |
| ENSG00000198691 | ABCA4 | 295.574 | -2.203 | 1.05151E-10 |
| ENSG00000272620 | AFAP1-AS1 | 524.670 | -2.216 | 8.74939E-07 |
| ENSG00000206072 | SERPINB11 | 32.126 | -2.229 | 0.000159362 |
| ENSG00000228295 | LINC00392 | 6.474 | -2.241 | 0.002005333 |
| ENSG00000250271 | | 16.357 | -2.262 | 0.00010383 |
| ENSG00000232756 | DDX3ILA1 | 12.687 | -2.272 | 4.93422E-06 |
| ENSG00000173467 | AGR3 | 59.112 | -2.272 | 1.02112E-06 |
| ENSG00000229807 | XIST | 848.021 | -2.274 | 0.000576003 |
| ENSG00000108602 | ALDH3A1 | 1648.312 | -2.292 | 1.72786E-08 |
| ENSG00000134398 | ERN2 | 877.383 | -2.308 | 7.02725E-08 |
| ENSG00000185860 | CCDC190 | 86.120 | -2.366 | 2.01971E-05 |
| ENSG00000181617 | FDCSP | 167.757 | -2.367 | 1.07049E-05 |
| ENSG00000127324 | TSPAN8 | 347.337 | -2.370 | 2.96981E-08 |
| ENSG00000169605 | GKN1 | 24.117 | -2.397 | 6.65519E-05 |
| ENSG00000178172 | SPINK6 | 15.674 | -2.398 | 2.16204E-05 |
| ENSG00000134827 | TCN1 | 1258.637 | -2.406 | 2.59032E-07 |
| ENSG00000163586 | FABP1 | 13.136 | -2.418 | 0.017936776 |
| ENSG00000173080 | RXFP4 | 5.509 | -2.428 | 8.10025E-10 |
| ENSG00000228695 | CES1P1 | 8.077 | -2.429 | 7.67014E-06 |
| ENSG00000207296 | RNU6-140P | 9.333 | -2.442 | 0.001196683 |
| ENSG00000231131 | LNCAROD | 17.041 | -2.474 | 3.71539E-05 |
| ENSG00000173702 | MUC13 | 53.961 | -2.474 | 3.71878E-08 |
| ENSG00000127831 | VIL1 | 17.765 | -2.482 | 1.77936E-08 |
| ENSG00000188393 | CLEC2A | 9.839 | -2.531 | 0.003074537 |
| ENSG00000145321 | GC | 19.532 | -2.564 | 0.013496039 |
| ENSG00000265787 | CYP4F35P | 137.195 | -2.590 | 2.4403E-10 |
| ENSG00000140297 | GCNT3 | 171.380 | -2.595 | 2.47905E-14 |
| ENSG00000124664 | SPDEF | 347.415 | -2.613 | 8.33491E-15 |
| ENSG00000204983 | PRSS1 | 97.669 | -2.644 | 5.16991E-06 |
| ENSG00000248474 | LINC02122 | 5.595 | -2.705 | 8.42697E-05 |
| ENSG00000105388 | CEACAM5 | 3464.076 | -2.710 | 2.12941E-08 |
| ENSG00000198848 | CES1 | 980.141 | -2.752 | 2.73836E-13 |
| ENSG00000204616 | TRIM31 | 2052.050 | -2.791 | 7.27391E-14 |
| ENSG00000162009 | SSTR5 | 12.872 | -2.844 | 1.94274E-06 |
| ENSG00000233041 | PHGR1 | 180.490 | -2.871 | 1.37635E-05 |
| ENSG00000242317 | | 13.561 | -2.899 | 1.21271E-05 |
| ENSG00000214814 | FER1L6 | 36.151 | -2.919 | 3.83276E-11 |
| ENSG00000250920 | | 77.206 | -3.007 | 0.000183882 |
| ENSG00000160182 | TFF1 | 608.562 | -3.025 | 5.48666E-10 |
| ENSG00000198183 | BPIFA1 | 27.839 | -3.062 | 2.40346E-05 |
| ENSG00000251676 | SNHG27 | 4.054 | -3.062 | 0.00044881 |
| ENSG00000196188 | CTSE | 2771.533 | -3.156 | 2.58949E-12 |
| ENSG00000047457 | CP | 2381.020 | -3.230 | 8.88843E-17 |
| ENSG00000253474 | | 12.124 | -3.297 | 6.37518E-05 |
| ENSG00000242366 | UGT1A8 | 260.465 | -3.368 | 5.5464E-10 |
| ENSG00000131050 | BPIFA2 | 9.001 | -3.402 | 1.05115E-09 |
| ENSG00000196091 | MYBPC1 | 392.306 | -3.412 | 8.31784E-12 |
| ENSG00000196344 | ADH7 | 307.261 | -3.423 | 7.18042E-09 |
| ENSG00000223829 | | 10.002 | -3.466 | 3.58239E-06 |
| ENSG00000198074 | AKR1B10 | 3503.547 | -3.635 | 4.61943E-17 |
| ENSG00000249837 | | 6.043 | -3.701 | 1.81265E-08 |
| ENSG00000171747 | LGALS4 | 1029.019 | -3.704 | 1.16397E-20 |
| ENSG00000244468 | | 40.912 | -3.830 | 1.5209E-15 |
| ENSG00000125999 | BPIFB1 | 483.721 | -3.840 | 1.15843E-16 |
| ENSG00000102837 | OLFM4 | 1691.973 | -3.913 | 1.72283E-14 |

| | | | | |
|---|---|---|---|---|
| ENSG00000162896 | PIGR | 2812.475 | -4.048 | 3.15637E-23 |
| ENSG00000261713 | SSTR5-AS1 | 78.512 | -4.657 | 1.00263E-14 |
| ENSG00000149021 | SCGB1A1 | 175.163 | -4.797 | 7.65574E-13 |
