## Supplementary material for "*E2F3* amplification primes bladder cancer cells for premature mitosis": Table S2

**Table S2** DE gene set enrichment analysis TCGA bladder cancer

**MSigDB pathway enrichment - UP in E2F3-amplified bladder cancers TCGA**

| Term | Overlap | Adjusted P-val | Odds Ratio | Combined |
|---|---|---|---|---|
| G2-M Checkpoint | 72/200 | 8.64184E-19 | 4.586598474 | 208.6183 |
| E2F Targets | 70/200 | 8.85387E-18 | 4.386036596 | 186.2502 |
| Mitotic Spindle | 56/199 | 5.63883E-10 | 3.167013803 | 76.29141 |
| Spermatogenesis | 29/135 | 0.004790963 | 2.190043449 | 17.18428 |
| Myc Targets V1 | 38/200 | 0.00718194 | 1.879425929 | 13.56676 |

**MSigDB pathway enrichment - DOWN in E2F3-amplified bladder cancers TCGA**

| Term | Overlap | Adjusted P-val | Odds Ratio | Combined |
|---|---|---|---|---|
| Interferon Gamma Response | 58/200 | 4.42373E-12 | 3.667821445 | 110.2403 |
| Interferon Alpha Response | 33/97 | 4.8378E-09 | 4.59263004 | 102.7173 |
| Complement | 41/200 | 0.000176159 | 2.293692776 | 26.28006 |
| Hypoxia | 37/200 | 0.002589705 | 2.014641311 | 16.63851 |
| Xenobiotic Metabolism | 37/200 | 0.002589705 | 2.014641311 | 16.63851 |
| Androgen Response | 21/100 | 0.006651161 | 2.351630663 | 16.19447 |
| Myogenesis | 35/200 | 0.006651161 | 1.880554423 | 12.87342 |
| p53 Pathway | 35/200 | 0.006651161 | 1.880554423 | 12.87342 |
| Reactive Oxygen Species Pathway | 18233 | 0.016096065 | 2.863160698 | 16.48771 |
| Coagulation | 25/138 | 0.016096065 | 1.957366807 | 11.23258 |
| heme Metabolism | 33/200 | 0.017472794 | 1.749918779 | 9.731715 |
| IL-6/JAK/STAT3 Signaling | 17/87 | 0.026603743 | 2.14529806 | 10.84195 |
| Allograft Rejection | 32/200 | 0.026741952 | 1.685856432 | 8.376338 |
| Estrogen Response Late | 31/200 | 0.041461216 | 1.622611142 | 7.15501 |
| Apoptosis | 26/161 | 0.041461216 | 1.702667549 | 7.469551 |

**CHEA Transcription factor binding site enrichment - UP in E2F3-amplified bladder cancers TCGA**

| Term | Overlap | Adjusted P-val | Odds Ratio | Combined |
|---|---|---|---|---|
| FOXM1 25889361 ChIP-Seq OE33 AND U2OS Human | 183/784 | 3.13E-20 | 2.546934446 | 131.2557 |
| FOXM1 23109430 ChIP-Seq U2OS Human | 74/226 | 1.01E-15 | 3.967954651 | 160.5322 |
| KDM5B 21448134 ChIP-Seq MESCs Mouse | 466/3030 | 2.63E-12 | 1.561752544 | 50.27174 |
| E2F4 17652178 ChIP-ChIP JURKAT Human | 131/608 | 1.12E-11 | 2.256586237 | 68.72876 |
| E2F1 26619117 ChIP-Seq Hepatocytes Mouse Liver | 396/2717 | 4.25E-07 | 1.43284136 | 28.20905 |
| MYC 30127528 ChIP-Seq KELLY Human Brain Neuroblastoma | 570/4188 | 2.60E-06 | 1.338617224 | 23.68526 |
| FOXM1 26456572 ChIP-Seq MCF-7 Human BreastCancer | 93/478 | 6.24E-06 | 1.959983994 | 32.65991 |
| CREM 20920259 ChIP-Seq GC1-SPG Mouse | 616/4643 | 2.06E-05 | 1.298003148 | 19.90852 |
| MYCN 21190229 ChIP-Seq SHEP-21N Human | 57/271 | 1.45E-04 | 2.14637239 | 28.48217 |
| MYCN 28898695 ChIP-Seq NB1643 Human Nerve Neuroblastoma | 307/2184 | 4.73E-04 | 1.34783368 | 16.14647 |
| NR4A2 19515692 ChIP-ChIP MN9D Mouse | 32/128 | 5.71E-04 | 2.673475564 | 31.19944 |
| E2F1 21310950 ChIP-Seq MCF-7 Human | 137/858 | 5.71E-04 | 1.54365891 | 17.91867 |
| MYC 22102868 ChIP-Seq CA46 Human Blood BurkittsLymphoma | 319/2314 | 0.001205793 | 1.316085098 | 14.18879 |
| CCND1 20090754 ChIP-ChIP RETINA Mouse | 213/1472 | 0.001767522 | 1.381066411 | 14.25884 |
| MYBL2 22936984 ChIP-ChIP MESCs Mouse | 206/1419 | 0.001833174 | 1.385398937 | 14.15746 |
| CRX 20693478 ChIP-Seq ADULT RETINA Mouse | 90/544 | 0.004080339 | 1.599858291 | 14.96568 |
| POU2F1 27270436 ChIP-Seq VCaP Human Prostate Carcinoma | 140/933 | 0.006835959 | 1.430223284 | 12.55412 |
| MYCN 18555785 ChIP-Seq MESCs Mouse | 232/1668 | 0.007671592 | 1.31708009 | 11.33381 |
| MYC 19030024 ChIP-ChIP MESCs Mouse | 373/2842 | 0.010466941 | 1.241040459 | 10.22678 |
| XRN2 22483619 ChIP-Seq HELA Human | 181/1296 | 0.029352441 | 1.315883696 | 9.419148 |
| MYBL1 21750041 ChIP-ChIP SPERMATOCYTES Mouse | 23/103 | 0.029540026 | 2.298570298 | 16.23295 |
| SOX2 18555785 ChIP-Seq MESCs Mouse | 61/365 | 0.029540026 | 1.611631785 | 11.37225 |
| GTF3C2 35216376 ChIP-Seq H9 Human BoneMarrow Lymphoma | 178/1277 | 0.030834428 | 1.312258503 | 9.145158 |
| MBD3 35695185 ChIP-Seq nicBasalRootGanglia Mouse Embryo | 463/3656 | 0.031130361 | 1.192442695 | 8.248019 |
| SOX9 26525672 Chip-Seq Limbbuds Mouse | 142/993 | 0.033508831 | 1.348453212 | 9.172803 |

| | | | | |
|---|---|---|---|---|
| RUNX2 24655370 ChIP-Seq MC3T3E1 Mouse Bone | 545/4374 | 0.039902441 | 1.173714768 | 7.733153 |
| MYC 18358816 ChIP-ChIP MESCs Mouse | 308/2369 | 0.048816783 | 1.217871021 | 7.732551 |

**CHEA Transcription factor binding site enrichment - DOWN in E2F3-amplified bladder cancers TCGA**

| Term | Overlap | Adjusted P-val | Odds Ratio | Combined |
|---|---|---|---|---|
| RXR 22158963 ChIP-Seq LIVER Mouse | 251/1546 | 1.16466E-11 | 1.803273338 | 57.3514 |
| PPARA 22158963 ChIP-Seq LIVER Mouse | 246/1552 | 1.65967E-10 | 1.746433285 | 49.69323 |
| SOX2 20726797 ChIP-Seq SW620 Human | 291/1931 | 3.09679E-10 | 1.653597944 | 45.34978 |
| P63 26484246 Chip-Seq KERATINOCYTES Human | 242/1627 | 8.90489E-08 | 1.608770959 | 34.54969 |
| LXR 22158963 ChIP-Seq LIVER Mouse | 227/1578 | 4.79976E-06 | 1.537393585 | 26.54394 |
| RACK7 27058665 Chip-Seq MCF-7 Human | 237/1679 | 9.64608E-06 | 1.503871585 | 24.6413 |
| UBF1/2 26484160 Chip-Seq HMEC-DERIVED Human | 231/1650 | 2.3929E-05 | 1.486689867 | 22.77988 |
| SOX2 27498859 Chip-Seq STOMACH Mouse | 219/1573 | 6.61916E-05 | 1.473165388 | 20.84071 |
| Nerf2 26677805 Chip-Seq MACROPHAGESS Mouse | 149/993 | 6.61916E-05 | 1.596043543 | 22.43043 |
| FOSL1 28411283 ChIP-Seq MDA231-LM2-4175 Human BreastCancer | 178/1238 | 7.86362E-05 | 1.522311523 | 20.83225 |
| KLF4 26769127 Chip-Seq PDAC-Cell Line Human | 230/1678 | 7.86362E-05 | 1.447265315 | 19.71679 |
| IRF8 27001747 Chip-Seq BMDM Mouse | 209/1500 | 7.86362E-05 | 1.472024418 | 20.01043 |
| FOXA1 26769127 Chip-Seq PDAC-Cell Line Human | 227/1658 | 8.94259E-05 | 1.444453685 | 19.31421 |
| TP63 30713093 ChIP-Seq Epithelial Human Tongue SCC | 278/2100 | 8.94259E-05 | 1.3957053 | 18.57835 |
| FOXA1 33576154 ChIP-Seq Human HilarLN ProstateCancer | 121/781 | 9.04552E-05 | 1.650858928 | 21.84193 |
| KLF6 26769127 Chip-Seq PDAC-Cell Line Human | 227/1666 | 0.000104905 | 1.435728056 | 18.69018 |
| EGR1 23403033 ChIP-Seq LIVER Mouse | 86/516 | 0.00014695 | 1.792229039 | 22.61837 |
| RELA 24523406 ChIP-Seq FIBROSARCOMA Human | 158/1104 | 0.000256189 | 1.50813572 | 18.10859 |
| SMAD2 18955504 ChIP-ChIP HaCaT Human | 205/1513 | 0.000392081 | 1.42053258 | 16.30253 |
| SMAD3 18955504 ChIP-ChIP HaCaT Human | 205/1513 | 0.000392081 | 1.42053258 | 16.30253 |
| IRF8 22096565 ChIP-ChIP GC-B Mouse | 76/463 | 0.00074907 | 1.755155472 | 18.92092 |
| ESR1 17901129 ChIP-ChIP LIVER Mouse | 61/354 | 0.001045648 | 1.856484728 | 19.25954 |
| RARG 19884340 ChIP-ChIP MEFs Mouse | 56/317 | 0.001045648 | 1.91191212 | 19.79912 |
| IRF8 22096565 ChIP-ChIP GC-B Human | 32/150 | 0.00132184 | 2.406948309 | 24.25895 |
| STAT5 23275557 ChIP-Seq MAMMARY-EPITHELIUM Mouse | 126/897 | 0.003576924 | 1.465955338 | 13.20218 |
| GATA4 25053715 ChIP-Seq YYC3 Human | 208/1602 | 0.003576924 | 1.347616909 | 12.13286 |
| SMRT 22465074 ChIP-Seq MACROPHAGES Mouse | 205/1580 | 0.004019437 | 1.345876182 | 11.90941 |
| NCOR 22465074 ChIP-Seq MACROPHAGES Mouse | 207/1604 | 0.004871258 | 1.3372864 | 11.52773 |
| ELF3 26769127 Chip-Seq PDAC-Cell Line Human | 215/1680 | 0.005688645 | 1.324835469 | 11.1684 |
| STAT5A 24692510 ChIP-Seq Epithelium Mouse Mammary | 248/1977 | 0.006244108 | 1.297555698 | 10.77355 |
| IRF1 21803131 ChIP-Seq MONOCYTES Human | 47/276 | 0.007546956 | 1.823911307 | 14.69145 |
| E2A 27217539 Chip-Seq RAMOS-Cell Line Human | 205/1603 | 0.007546956 | 1.321897757 | 10.63984 |
| TAF7L 23326641 ChIP-Seq C3H10T1-2 Mouse | 82/554 | 0.008444305 | 1.549909201 | 12.25327 |
| TP63 23658742 ChIP-Seq EP156T Human | 351/2937 | 0.011435622 | 1.233858519 | 9.343644 |
| ESR1 21235772 ChIP-Seq MCF-7 Human | 36/200 | 0.011520914 | 1.947158524 | 14.67434 |
| ESR2 21235772 ChIP-Seq MCF-7 Human | 58/370 | 0.013199207 | 1.653400279 | 12.18907 |
| TBP 23326641 ChIP-Seq C3H10T1-2 Mouse | 87/607 | 0.014418822 | 1.492333268 | 10.82888 |
| CEBPB 24764292 ChIP-Seq MC3T3 Mouse | 186/1461 | 0.015031434 | 1.311290568 | 9.425644 |
| FOXA1 27197147 Chip-Seq ENDOMETRIOID-ADENOCARCINOMA Human | 69/462 | 0.015713179 | 1.563067054 | 11.1255 |
| ESR1 26153859 ChIP-Seq MCF-7 Human BreastCancer | 420/3597 | 0.01634489 | 1.204717011 | 8.496871 |
| JUND 26020271 ChIP-Seq SMOOTH MUSCLE Human | 198/1573 | 0.016681614 | 1.294984816 | 9.075147 |
| FOXO1 23066095 ChIP-Seq LIVER Mouse | 45/276 | 0.019042564 | 1.729249719 | 11.81506 |
| MYB 21317192 ChIP-Seq ERMYB Mouse | 105/767 | 0.019042564 | 1.416274271 | 9.670207 |
| CREB1 26743006 Chip-Seq LNCaP Human | 205/1645 | 0.020844848 | 1.280087552 | 8.595148 |
| NR1H3 23393188 ChIP-Seq ATHEROSCLEROTIC-FOAM Human | 77/537 | 0.023022412 | 1.49059188 | 9.826974 |
| BRD4 27068464 Chip-Seq AML-cells Mouse | 199/1597 | 0.023587771 | 1.279030504 | 8.37308 |
| MECOM 23826213 ChIP-Seq KASUMI Mouse | 191/1527 | 0.02406337 | 1.28381891 | 8.34971 |
| FOXO1 32281255 ChIP-Seq Chondrocytes Human Osteoarthritis | 76/531 | 0.02406337 | 1.487069187 | 9.627796 |
| GATA2 20887958 ChIP-Seq HPC-7 Mouse | 168/1324 | 0.02406337 | 1.302995277 | 8.421631 |

| | | | | |
|---|---|---|---|---|
| GATA1 22383799 ChIP-Seq G1ME Mouse | 199/1606 | 0.027871402 | 1.270158393 | 7.978222 |
| FOXA1 27270436 ChIP-Seq LNCaP Human Prostate Carcinoma | 223/1822 | 0.027871402 | 1.254239438 | 7.872071 |
| TCF3 18692474 ChIP-Seq MEFs Mouse | 85/611 | 0.028485634 | 1.439426763 | 8.975046 |
| IRF1 21803131 ChIP-Seq PrimaryMonocytes Human Connective | 17/79 | 0.032135553 | 2.423193947 | 14.77068 |
| GATA3 24758297 ChIP-Seq MCF-7 Human | 200/1629 | 0.038908595 | 1.255899017 | 7.388575 |
| TCFCP2L1 18555785 ChIP-Seq MESCs Mouse | 184/1486 | 0.038908595 | 1.266869385 | 7.428856 |
| VDR 24763502 ChIP-Seq THP-1 Human | 133/1034 | 0.038908595 | 1.31893554 | 7.714757 |
| JUN 26020271 ChIP-Seq SMOOTH MUSCLE Human | 195/1587 | 0.039879794 | 1.256452682 | 7.287345 |
| ESR1 20079471 ChIP-ChIP T-47D Human | 29/167 | 0.039879794 | 1.860299359 | 10.77017 |
| ATF3 27146783 Chip-Seq COLON Human | 199/1628 | 0.044477801 | 1.248941487 | 7.073092 |
| TP63 22573176 ChIP-Seq HFKS Human | 382/3315 | 0.047793013 | 1.178847678 | 6.571572 |
