## Supplementary material for "*E2F3* amplification primes bladder cancer cells for premature mitosis": Table S3

**Supplemental table S3** *TP53* mutational status of cell lines used in this study.

| Cell line | TP53 mutation status | Anticipated effect | Transcriptionally active p53 expression? |
| --- | --- | --- | --- |
| 5637 | R280T | DNA-binding domain mutation | No |
| TCCSUP | E349* | Truncating mutation of tetramerization motif | No |
| HT1376 | P250L | DNA-binding domain mutation | No |
| T24 | Y126* truncation | Truncating mutation in DNA binding domain | No |
| UMUC3 | F113C | DNA-binding domain mutation | No |
| RPE1-hTERT | Intact | n/a | Yes |
| RPE1-hTERT-*TP53KO* | X111 in/del | Protein deletion. | No |
