## Supplementary material for "*E2F3* amplification primes bladder cancer cells for premature mitosis": Table S5

**Table S5** Gene set enrichment analysis DE genes in RP-6306-resistant E2F3-amplified bladder cancer cell lines

**MSigDB pathway enrichment - UP in all PKMYT1i-resistant bladder cancer cell lines**

| Term | Overlap | Adjusted P-v | Odds Ratio | Combined Score |
|---|---|---|---|---|
| Epithelial Mesenchymal Transition | 21/200 | 1.76E-05 | 4.06809603 | 60.450777 |
| Mitotic Spindle | 18/199 | 4.82E-04 | 3.42951346 | 37.2316381 |
| Myogenesis | 17/200 | 0.00119146 | 3.1975229 | 30.5235108 |
| UV Response Dn | 13/144 | 0.0034428 | 3.40066167 | 27.8758986 |
| Apical Junction | 15/200 | 0.00711744 | 2.78060413 | 20.1532426 |
| Hedgehog Signaling | 5/36 | 0.02923411 | 5.4779391 | 30.965051 |

**MSigDB pathway enrichment - DOWN in all PKMYT1i-resistant bladder cancer cell lines**

| Term | Overlap | Adjusted P-v | Odds Ratio | Combined Score |
|---|---|---|---|---|
| Estrogen Response Late | 28/200 | 2.1186E-09 | 5.15611329 | 123.047076 |
| Estrogen Response Early | 22/200 | 1.0318E-05 | 3.87508262 | 56.8873347 |
| TNF-alpha Signaling via NF-kB | 20/200 | 5.996E-05 | 3.47195078 | 41.6780366 |
| mTORC1 Signaling | 20/200 | 5.996E-05 | 3.47195078 | 41.6780366 |
| p53 Pathway | 20/200 | 5.996E-05 | 3.47195078 | 41.6780366 |
| Androgen Response | 13/100 | 0.00013348 | 4.63892128 | 51.1284057 |
| Myc Targets V1 | 19/200 | 0.00015071 | 3.27462606 | 35.1892578 |
| Oxidative Phosphorylation | 18/200 | 0.0004388 | 3.08006279 | 29.3956198 |
| Cholesterol Homeostasis | 27273 | 0.0029275 | 4.27591385 | 32.1899386 |
| Xenobiotic Metabolism | 16/200 | 0.00325561 | 2.69902913 | 19.7477437 |
| Interferon Gamma Response | 15/200 | 0.00752299 | 2.51246562 | 15.8202547 |
| KRAS Signaling Up | 15/200 | 0.00752299 | 2.51246562 | 15.8202547 |
| Adipogenesis | 14/200 | 0.01673453 | 2.32847728 | 12.4411572 |
| Epithelial Mesenchymal Transition | 14/200 | 0.01673453 | 2.32847728 | 12.4411572 |
| IL-6/JAK/STAT3 Signaling | 31990 | 0.02069888 | 3.11999029 | 15.7916613 |
| Wnt-beta Catenin Signaling | 15462 | 0.03116542 | 4.15266618 | 19.0510939 |
| Inflammatory Response | 13/200 | 0.03151515 | 2.14702007 | 9.57299471 |
| heme Metabolism | 13/200 | 0.03151515 | 2.14702007 | 9.57299471 |

**CHEA TF binding site enrichment - UP in all PKMYT1i-resistant bladder cancer cell lines**

| Term | Overlap | Adjusted P-v | Odds Ratio | Combined Score |
|---|---|---|---|---|
| MBD3 35695185 ChIP-Seq nicBasalRootG | 194/3656 | 1.692E-16 | 2.34152398 | 100.518872 |
| CHD4 35695185 ChIP-Seq nicBasalRootGa | 124/2119 | 2.3538E-12 | 2.39669528 | 78.3604966 |
| WT1 25993318 ChIP-Seq PODOCYTE Hum | 153/2909 | 4.2762E-12 | 2.18753667 | 69.3289944 |
| EBF1 22473956 ChIP-Seq BONE MARROW | 97/1502 | 5.8789E-12 | 2.59711142 | 80.7357394 |
| EGR1 20690147 ChIP-Seq ERYTHROLEUKE | 222/4931 | 7.5722E-12 | 1.95966253 | 59.9862295 |
| EBF1 22473956 ChIP-Seq LYMPHODE Mo | 95/1505 | 2.9671E-11 | 2.52330399 | 73.333469 |
| PU1 27457419 Chip-Seq LIVER Mouse | 94/1510 | 7.8215E-11 | 2.4801791 | 69.2938066 |
| SUZ12 20075857 ChIP-Seq MESCs Mouse | 173/3611 | 7.8781E-11 | 1.99606644 | 55.4872446 |
| WT1 20215353 ChIP-ChIP NEPHRON PRO | 82/1307 | 1.5272E-09 | 2.46602991 | 60.9503006 |
| P53 22387025 ChIP-Seq ESCs Mouse | 90/1511 | 2.0289E-09 | 2.34615685 | 57.0739196 |
| SUZ12 18555785 Chip-Seq ESCs Mouse | 89/1491 | 2.1655E-09 | 2.3491785 | 56.7705297 |
| RUNX1 27457419 Chip-Seq LIVER Mouse | 88/1475 | 2.6829E-09 | 2.34504237 | 55.9640611 |
| SOX2 18555785 Chip-Seq ESCs Mouse | 89/1508 | 3.3241E-09 | 2.3188453 | 54.6563427 |
| RUNX2 24655370 ChIP-Seq MC3T3E1 Mo | 192/4374 | 4.4963E-09 | 1.82233381 | 42.2678273 |

| | | | | |
|---|---|---|---|---|
| NR3C1 27076634 ChIP-Seq BEAS2B Huma | 147/3084 | 6.8526E-09 | 1.92352455 | 43.6716437 |
| CTCF 18555785 Chip-Seq ESCs Mouse | 86/1492 | 1.9555E-08 | 2.249177 | 48.5615897 |
| NANOG 18555785 Chip-Seq ESCs Mouse | 87/1522 | 2.074E-08 | 2.23031145 | 47.7888089 |
| CMYC 18555785 Chip-Seq ESCs Mouse | 86/1499 | 2.074E-08 | 2.2371651 | 47.7862276 |
| P300 18555785 Chip-Seq ESCs Mouse | 86/1499 | 2.074E-08 | 2.2371651 | 47.7862276 |
| NMYC 18555785 Chip-Seq ESCs Mouse | 86/1510 | 2.8193E-08 | 2.21852786 | 46.5931427 |
| ZFP281 18757296 ChIP-ChIP E14 Mouse | 84/1466 | 3.276E-08 | 2.2289012 | 46.3676599 |
| ZFP281 27345836 Chip-Seq ESCs Mouse | 91/1646 | 3.434E-08 | 2.15610303 | 44.6513545 |
| FOXA2 19822575 ChIP-Seq HepG2 Humar | 124/2553 | 6.5462E-08 | 1.91945045 | 38.4267829 |
| OCT4 18555785 Chip-Seq ESCs Mouse | 84/1492 | 6.6098E-08 | 2.1845898 | 43.6206937 |
| KLF4 18555785 Chip-Seq ESCs Mouse | 83/1486 | 1.1615E-07 | 2.16248144 | 41.8718397 |
| E2F1 18555785 Chip-Seq ESCs Mouse | 84/1528 | 1.8536E-07 | 2.12586987 | 40.085913 |
| ZFX 18555785 Chip-Seq ESCs Mouse | 82/1485 | 2.2772E-07 | 2.13210269 | 39.6841886 |
| SOX2 20726797 ChIP-Seq SW620 Human | 99/1931 | 2.2841E-07 | 1.99299662 | 36.9707537 |
| TCFCP2L1 18555785 Chip-Seq ESCs Mous | 83/1513 | 2.2841E-07 | 2.1184726 | 39.2726883 |
| EOMES 21245162 ChIP-Seq HESCs Humar | 54/820 | 3.465E-07 | 2.51976231 | 45.5763957 |
| NFE2 27457419 Chip-Seq LIVER Mouse | 50/735 | 4.248E-07 | 2.60039966 | 46.4198539 |
| TP53 23651856 ChIP-Seq MEFs Mouse | 120/2541 | 4.7457E-07 | 1.84819235 | 32.7287306 |
| BRD4 25478319 ChIP-Seq HGPS Human | 91/1770 | 7.7956E-07 | 1.98301026 | 34.0631607 |
| SMAD1 18555785 Chip-Seq ESCs Mouse | 80/1483 | 7.7956E-07 | 2.07171269 | 35.5331094 |
| MTF2 20144788 ChIP-Seq MESCs Mouse | 117/2488 | 8.6947E-07 | 1.83333747 | 31.1913361 |
| STAT3 18555785 Chip-Seq ESCs Mouse | 80/1495 | 1.0392E-06 | 2.05277556 | 34.5009032 |
| KDM2B 26808549 Chip-Seq SUP-B15 Hum | 86/1666 | 1.4847E-06 | 1.98215448 | 32.5524602 |
| PCGF2 27294783 Chip-Seq ESCs Mouse | 32/385 | 1.7554E-06 | 3.17797034 | 51.5740405 |
| MITF 21258399 ChIP-Seq MELANOMA Hu | 187/4614 | 2.0688E-06 | 1.62849667 | 26.1184464 |
| GATA4 21415370 ChIP-Seq HL-1 Mouse | 74/1374 | 2.4651E-06 | 2.05512718 | 32.5486739 |
| LEF1 29337183 ChIP-Seq mESC Mouse Ste | 109/2335 | 3.5553E-06 | 1.80327683 | 27.8550065 |
| AR 19668381 ChIP-Seq PC3 Human | 124/2803 | 7.7719E-06 | 1.71472977 | 25.1048573 |
| SUZ12 27294783 Chip-Seq NPCs Mouse | 43/658 | 9.4696E-06 | 2.46735612 | 35.5783018 |
| CDX2 21402776 ChIP-Seq INTESTINAL-VIL | 77/1514 | 1.1885E-05 | 1.93148859 | 27.3681321 |
| RING1B 27294783 Chip-Seq NPCs Mouse | 81/1632 | 1.5291E-05 | 1.88566907 | 26.2013722 |
| P300 27058665 Chip-Seq ZR-75-30cells Hu | 83/1689 | 1.6067E-05 | 1.86786114 | 25.8203882 |
| TCFAP2C 20176728 ChIP-ChIP TROPHOBL | 93/1980 | 2.2544E-05 | 1.78944856 | 24.0749494 |
| DMRT1 23473982 ChIP-Seq TESTES Mous | 82/1678 | 2.2544E-05 | 1.85420028 | 24.9245716 |
| SRY 25088423 ChIP-ChIP EMBRYONIC GO | 115/2614 | 2.5983E-05 | 1.68950444 | 22.4359945 |
| TCF4 23295773 ChIP-Seq U87 Human | 134/3177 | 2.775E-05 | 1.63200313 | 21.5320752 |
| MYC 19915707 ChIP-ChIP AK7 Human | 83/1732 | 3.9186E-05 | 1.81476388 | 23.2811574 |
| HAND2 30127528 ChIP-Seq KELLY Human | 140/3383 | 4.073E-05 | 1.60213813 | 20.4336576 |
| TBX2 33478486 ChIP-Seq IMR90 Human L | 105/2354 | 4.073E-05 | 1.70245516 | 21.6962177 |
| TP63 23658742 ChIP-Seq EP156T Human | 125/2937 | 4.073E-05 | 1.63734454 | 20.8482333 |
| PCGF2 27294783 Chip-Seq NPCs Mouse | 27/346 | 4.1266E-05 | 2.94542371 | 37.4113902 |
| ESRRB 18555785 Chip-Seq ESCs Mouse | 74/1496 | 4.3611E-05 | 1.86616123 | 23.5663065 |
| MYC 30127528 ChIP-Seq KELLY Human Br | 166/4188 | 4.4868E-05 | 1.55045542 | 19.5080021 |
| SOX6 21985497 ChIP-Seq MYOTUBES Mo | 74/1504 | 5.0965E-05 | 1.8548965 | 23.0699095 |
| EZH2 27294783 Chip-Seq NPCs Mouse | 56/1032 | 5.1135E-05 | 2.03556116 | 25.2753282 |
| ISL1 27105846 Chip-Seq CPCs Mouse | 66/1293 | 5.2484E-05 | 1.9192387 | 23.7487513 |
| STAT3 23295773 ChIP-Seq U87 Human | 114/2637 | 5.4096E-05 | 1.65294357 | 20.3762548 |

| | | | | |
|---|---|---|---|---|
| OCT4 21477851 ChIP-Seq ESCs Mouse | 74/1517 | 6.471E-05 | 1.8368577 | 22.2844832 |
| SMARCD1 25818293 ChIP-Seq ESCs Mous | 80/1683 | 6.6765E-05 | 1.79310969 | 21.6689915 |
| ESR1 22446102 ChIP-Seq UTERUS Mouse | 61/1183 | 8.5759E-05 | 1.93209421 | 22.8344046 |
| STAT3 24763339 ChIP-Seq IMN-ESCs Mou | 69/1398 | 8.9942E-05 | 1.85299709 | 21.7826064 |
| TP53 22127205 ChIP-Seq IMR90 Human | 44/766 | 0.00012944 | 2.14235728 | 24.355174 |
| ELK3 25401928 ChIP-Seq HUVEC Human | 79/1689 | 0.00012944 | 1.75875876 | 19.9765405 |
| SOX9 26525672 Chip-Seq Limbbuds Mous | 53/993 | 0.00012944 | 1.99270168 | 22.609605 |
| POU3F1 26484290 ChIP-Seq ESCss Mouse | 74/1557 | 0.00014353 | 1.78333732 | 20.0237769 |
| FLI1 27457419 Chip-Seq LIVER Mouse | 23/294 | 0.0001648 | 2.93947806 | 32.5260441 |
| SMAD3 30307970 ChIP-Seq HCASMC Hun | 81/1759 | 0.0001648 | 1.73056669 | 19.1426544 |
| PCGF4 22325352 ChIP-Seq 293T-Rex Hum | 75/1594 | 0.00016882 | 1.76457487 | 19.4515567 |
| JUN 27471255 ChIP-Seq Chondrocytes Mo | 161/4145 | 0.00017198 | 1.50350801 | 16.5251493 |
| TFAP2C 20629094 ChIP-Seq MCF-7 Huma | 54/1034 | 0.00018011 | 1.94693878 | 21.282538 |
| PPARD 21283829 ChIP-Seq MYOFIBROBLA | 115/2749 | 0.00018011 | 1.5901271 | 17.3607574 |
| PU 27001747 Chip-Seq BMDM Mouse | 74/1573 | 0.00018428 | 1.76272894 | 19.1814726 |
| TEAD4 22529382 ChIP-Seq TROPHECTODI | 80/1748 | 0.00021453 | 1.71694902 | 18.3999009 |
| CRX 20693478 ChIP-Seq RETINA Mouse | 70/1492 | 0.00033627 | 1.75133281 | 17.9583041 |
| TCF3 18467660 ChIP-ChIP MESCs Mouse | 51/982 | 0.00033627 | 1.92960565 | 19.7620997 |
| CTCF 33576154 ChIP-Seq Human Liver Pro | 144/3679 | 0.00036287 | 1.49825554 | 15.2115155 |
| TP53 18474530 ChIP-ChIP U2OS Human | 39/687 | 0.00043398 | 2.10435202 | 20.9624035 |
| CTCF 31629814 ChIP-Seq Hepatocytes Mo | 124/3086 | 0.00046314 | 1.52468734 | 15.0702451 |
| CBX2 22325352 ChIP-Seq 293T-Rex Huma | 73/1595 | 0.00047323 | 1.70703244 | 16.8150821 |
| MYB 21317192 ChIP-Seq ERMYB Mouse | 42/767 | 0.0004889 | 2.02856257 | 19.8919112 |
| SOX2 21211035 ChIP-Seq LN229 Gbm | 110/2673 | 0.0005053 | 1.55289285 | 15.1579214 |
| SOX9 24532713 ChIP-Seq HFSC Mouse | 58/1203 | 0.00072202 | 1.78749642 | 16.7890491 |
| SUZ12 18974828 ChIP-Seq MESCs Mouse | 71/1564 | 0.00074203 | 1.68854125 | 15.7939381 |
| LMO2 26923725 Chip-Seq HEMANGIOBLA | 65/1404 | 0.0008445 | 1.71802861 | 15.8095372 |
| RUNX2 24764292 ChIP-Seq MC3T3 Mouse | 72/1600 | 0.0008445 | 1.67314884 | 15.395497 |
| PRDM14 21183938 ChIP-Seq MESCs Mou | 69/1520 | 0.00091027 | 1.68575437 | 15.3662292 |
| SOX2 18358816 ChIP-ChIP MESCs Mouse | 34/591 | 0.00091263 | 2.12484514 | 19.3397182 |
| NR3C2 34362910 ChIP-Seq WistarRat Hip | 68/1496 | 0.00096424 | 1.68691414 | 15.2425624 |
| GATA1 21571218 ChIP-Seq MEGAKARYOC | 90/2132 | 0.00102349 | 1.57634128 | 14.1191675 |
| TCF3 18347094 ChIP-ChIP MESCs Mouse | 72/1613 | 0.00102349 | 1.65782887 | 14.8452335 |
| SMAD4 21799915 ChIP-Seq A2780 Human | 91/2167 | 0.00110783 | 1.56791212 | 13.8993151 |
| SMAD 19615063 ChIP-ChIP OVARY Huma | 11/100 | 0.00111287 | 4.22959133 | 37.4312375 |
| AR 27270436 ChIP-Seq VCaP Human Pros | 137/3565 | 0.00116772 | 1.45621879 | 12.8021676 |
| SMAD4 21741376 ChIP-Seq HESCs Human | 80/1862 | 0.00139843 | 1.5967923 | 13.7337262 |
| KDM2B 26808549 Chip-Seq HPB-ALL Hum | 72/1636 | 0.00148585 | 1.63134819 | 13.9154522 |
| NR3C2 34362910 ChIP-Seq WistarRat Hip | 48/974 | 0.00156881 | 1.81602199 | 15.3738133 |
| POU5F1 18347094 ChIP-ChIP MESCs Mou | 69/1555 | 0.00157889 | 1.64284406 | 13.88088 |
| MNX1 26342078 ChIP-Seq MIN6-4N Mou | 68/1533 | 0.00175952 | 1.64092876 | 13.6707837 |
| NFKB1 27076634 ChIP-Seq BEAS2B Huma | 144/3823 | 0.00177632 | 1.42656519 | 11.84351 |
| KLF4 26769127 Chip-Seq PDAC-Cell Line H | 73/1678 | 0.00177632 | 1.61125087 | 13.3592711 |
| CJUN 26792858 Chip-Seq BT549 Human | 69/1564 | 0.00177632 | 1.63213473 | 13.5208096 |
| PPARG 20887899 ChIP-Seq 3T3-L1 Mouse | 110/2774 | 0.00177632 | 1.48506876 | 12.3010548 |
| AF4 26711339 ChIP-Seq SEM Human Bloc | 116/2966 | 0.0019805 | 1.46650191 | 11.9670793 |
| SUZ12 27294783 Chip-Seq ESCs Mouse | 72/1657 | 0.0019805 | 1.60784137 | 13.1130084 |

| | | | | |
|---|---|---|---|---|
| KLF4 18358816 ChIP-ChIP MESCs Mouse | 57/1238 | 0.00215698 | 1.69649998 | 13.6756233 |
| FOXM1 26100407 CHIP-SEQ Hek293 Flp-I | 69/1577 | 0.00216155 | 1.61689137 | 13.015709 |
| FOXO3 23340844 ChIP-Seq DLD1 Human | 32/576 | 0.00219675 | 2.04152249 | 16.3824596 |
| CCND1 20090754 ChIP-ChIP RETINA Mous | 65/1472 | 0.00249996 | 1.62884905 | 12.8457016 |
| ETV1 20927104 ChIP-Seq GIST48 Human | 69/1588 | 0.00252036 | 1.60419694 | 12.6017589 |
| CDX2 19796622 ChIP-Seq MESCs Mouse | 18/252 | 0.00252036 | 2.645437 | 20.7398721 |
| TBX20 22080862 ChIP-Seq HEART Mouse | 69/1589 | 0.00252036 | 1.60305201 | 12.5647246 |
| TBX20 22328084 ChIP-Seq HEART Mouse | 69/1589 | 0.00252036 | 1.60305201 | 12.5647246 |
| TET1 21490601 ChIP-Seq MESCs Mouse | 70/1618 | 0.00252036 | 1.59752224 | 12.5141266 |
| TBX5 21415370 ChIP-Seq HL-1 Mouse | 68/1561 | 0.00252036 | 1.60764407 | 12.5815065 |
| EZH2 27294783 Chip-Seq ESCs Mouse | 71/1649 | 0.00257607 | 1.59001368 | 12.3953502 |
| RING1B 27294783 Chip-Seq ESCs Mouse | 71/1651 | 0.00264334 | 1.58782304 | 12.3240554 |
| TFAP2A 35110662 ChIP-Seq HEPM Huma | 53/1144 | 0.00273555 | 1.70287578 | 13.1445269 |
| SOX2 18692474 ChIP-Seq MESCs Mouse | 93/2302 | 0.00274121 | 1.50053845 | 11.5672304 |
| STAT5A 24692510 ChIP-Seq Epithelium M | 82/1977 | 0.00279757 | 1.535447 | 11.7925451 |
| RUNX1 21571218 ChIP-Seq MEGAKARYO( | 153/4172 | 0.00312659 | 1.38641982 | 10.482601 |
| NR3C1 34362910 ChIP-Seq WistarRat Hip | 34/647 | 0.00348878 | 1.92500135 | 14.3283095 |
| SUZ12 18692474 ChIP-Seq MESCs Mouse | 66/1531 | 0.00380187 | 1.5864204 | 11.6591816 |
| POU5F1 16518401 ChIP-PET MESCs Mous | 41/836 | 0.00391183 | 1.79577735 | 13.1324231 |
| GTF3C2 35216376 ChIP-Seq H9 Human Bo | 57/1277 | 0.00391948 | 1.63875675 | 11.9680851 |
| OLIG2 23332759 ChIP-Seq OLIGODENDRO | 71/1679 | 0.00392052 | 1.55772622 | 11.363773 |
| TAL1 26923725 Chip-Seq HPCs Mouse | 62/1420 | 0.00393159 | 1.60468874 | 11.6894544 |
| MAF 26560356 Chip-Seq TH2 Human | 69/1627 | 0.00426917 | 1.5606338 | 11.2280158 |
| POU5F1 18692474 ChIP-Seq MESCs Mous | 115/3012 | 0.0042928 | 1.42312303 | 10.2200124 |
| KDM2B 26808549 Chip-Seq JURKAT Huma | 70/1660 | 0.00448093 | 1.55166928 | 11.0648912 |
| NKX2-5 21415370 ChIP-Seq HL-1 Mouse | 46/980 | 0.00448483 | 1.7181932 | 12.2380027 |
| CEBPD 21427703 ChIP-Seq 3T3-L1 Mouse | 59/1349 | 0.00490771 | 1.60422533 | 11.2697738 |
| MEF2A 21415370 ChIP-Seq HL-1 Mouse | 35/691 | 0.00512752 | 1.85090844 | 12.9079822 |
| PPARD 23176727 ChIP-Seq KERATINOCYT | 13/164 | 0.00517731 | 2.94718455 | 20.5031867 |
| SMARCA4 23332759 ChIP-Seq OLIGODEN | 84/2089 | 0.00543389 | 1.48327961 | 10.2364543 |
| TP63 19390658 ChIP-ChIP HaCaT Human | 13/167 | 0.00602899 | 2.88932205 | 19.618743 |
| RBPJ 21746931 ChIP-Seq IB4 Human | 67/1598 | 0.00638789 | 1.53838484 | 10.3418434 |
| AR 22383394 ChIP-Seq PROSTATE CANCE | 65/1540 | 0.00638789 | 1.54789214 | 10.3987023 |
| NCOR1 26117541 ChIP-Seq K562 Human | 68/1631 | 0.00669908 | 1.52964972 | 10.1925794 |
| TCFCP2L1 18555785 ChIP-Seq MESCs Mo | 63/1486 | 0.00673187 | 1.55351309 | 10.3331018 |
| ZNF217 24962896 ChIP-Seq MCF-7 Huma | 56/1289 | 0.00723041 | 1.58883274 | 10.4434451 |
| SALL4 28974232 ChIP-Seq Mouse BoneM | 21/352 | 0.00727896 | 2.18259166 | 14.3165295 |
| P300 19829295 ChIP-Seq ESCs Human | 66/1594 | 0.00911058 | 1.51567601 | 9.59134357 |
| SETDB1 19884255 ChIP-Seq MESCs Mous | 62/1478 | 0.00919874 | 1.53401921 | 9.682178 |
| SALL4 18804426 ChIP-ChIP XEN Mouse | 34/693 | 0.00940118 | 1.78625223 | 11.2231868 |
| AR 25329375 ChIP-Seq VCAP Human | 62/1481 | 0.00951691 | 1.53052103 | 9.58737134 |
| EWS-ERG 20517297 ChIP-Seq CADO-ES1 I | 39/830 | 0.00960748 | 1.71078968 | 10.6784492 |
| PIAS1 25552417 ChIP-Seq VCAP Human | 30/589 | 0.00960748 | 1.85427274 | 11.5731059 |
| EP300 20729851 ChIP-Seq FORBRAIN MIC | 56/1313 | 0.01030791 | 1.55644085 | 9.59444167 |
| TFAP2A 17053090 ChIP-ChIP MCF-7 Huma | 64/1549 | 0.01072708 | 1.51001684 | 9.23817741 |
| TRIM28 19339689 ChIP-ChIP MESCs Mou | 87/2241 | 0.01108456 | 1.42645157 | 8.67087632 |
| RNF2 27304074 Chip-Seq ESCs Mouse | 52/1205 | 0.01138905 | 1.57255219 | 9.50617489 |

| | | | | |
|---|---|---|---|---|
| CTCF 33576154 ChIP-Seq Human HilarLN | 132/3649 | 0.01145932 | 1.34464262 | 8.10851784 |
| MEIS1 20887958 ChIP-Seq HPC-7 Mouse | 48/1092 | 0.01145932 | 1.60048763 | 9.64468289 |
| CREB1 26743006 Chip-Seq LNCaP Human | 67/1645 | 0.01161417 | 1.4886442 | 8.94127189 |
| FOXP2 23625967 ChIP-Seq PFSK-1 AND SI | 35/733 | 0.01179039 | 1.7356429 | 10.3877367 |
| SMAD3 22036565 ChIP-Seq ESCs Mouse | 39/844 | 0.01212832 | 1.67977376 | 9.99536367 |
| TAL1 26923725 Chip-Seq HEMANGIOBLAS | 47/1070 | 0.01240899 | 1.59811481 | 9.44727718 |
| ARNT 22903824 ChIP-Seq MCF-7 Human | 39/846 | 0.01240899 | 1.67543076 | 9.90131006 |
| KDM2B 26808549 Chip-Seq SIL-ALL Huma | 67/1652 | 0.01240899 | 1.48148841 | 8.74769945 |
| GFI1B 20887958 ChIP-Seq HPC-7 Mouse | 61/1475 | 0.01240899 | 1.50864448 | 8.90530489 |
| TP63 22573176 ChIP-Seq HFKS Human | 121/3315 | 0.01260577 | 1.35131806 | 7.94366764 |
| TCF21 26020271 ChIP-Seq HCASMC Huma | 49/1129 | 0.01260577 | 1.57926769 | 9.27819262 |
| TBX3 20139965 ChIP-Seq ESCs Mouse | 36/767 | 0.01293588 | 1.70478796 | 9.95286389 |
| P63 26484246 Chip-Seq KERATINOCYTES | 66/1627 | 0.01293588 | 1.48089837 | 8.6442563 |
| RELA 24523406 ChIP-Seq FIBROSARCOMA | 48/1104 | 0.01317256 | 1.58126722 | 9.1920728 |
| TBX3 20139965 ChIP-Seq MESCs Mouse | 36/770 | 0.01360594 | 1.69754768 | 9.80305661 |
| PPARG 20176806 ChIP-Seq 3T3-L1 Mouse | 57/1369 | 0.01405772 | 1.51614268 | 8.68897906 |
| NFIB 24661679 ChIP-Seq LUNG Mouse | 24/453 | 0.01405772 | 1.92510388 | 11.0317563 |
| FOXO1 32281255 ChIP-Seq Chondrocytes | 27/531 | 0.01411977 | 1.8462139 | 10.5608459 |
| FOXA1 27270436 ChIP-Seq VCaP Human I | 121/3331 | 0.01413413 | 1.34325699 | 7.67468765 |
| SA1 22415368 ChIP-Seq MEFs Mouse | 63/1549 | 0.01454116 | 1.48244433 | 8.41862509 |
| NFYA 21822215 ChIP-Seq K562 Human | 64/1579 | 0.01454116 | 1.47764026 | 8.38364685 |
| TAL1 20566737 ChIP-Seq PRIMARY FETAL | 61/1492 | 0.0148404 | 1.48931496 | 8.41110941 |
| CLOCK 20551151 ChIP-Seq 293T Human | 21/382 | 0.01567791 | 1.99806843 | 11.163414 |
| GATA3 30127528 ChIP-Seq BE2C Human I | 36/782 | 0.01655542 | 1.6691689 | 9.22556351 |
| FOXA1 25552417 ChIP-Seq VCAP Human | 66/1650 | 0.01671323 | 1.45751634 | 8.0338035 |
| ATF3 23680149 ChIP-Seq GBM1-GSC Hum | 66/1652 | 0.01708662 | 1.45551517 | 7.98254974 |
| OCT4 18692474 ChIP-Seq MEFs Mouse | 66/1655 | 0.01770857 | 1.45252286 | 7.90620429 |
| KDM2B 26808549 Chip-Seq REH Human | 57/1389 | 0.01786906 | 1.49172872 | 8.09436836 |
| EOMES 20176728 ChIP-ChIP TSCs Mouse | 52/1243 | 0.01786906 | 1.51921216 | 8.23388041 |
| NANOG 16518401 ChIP-PET MESCs Mous | 66/1657 | 0.01786906 | 1.45053426 | 7.85566191 |
| YAP1 20516196 ChIP-Seq MESCs Mouse | 73/1868 | 0.01786906 | 1.42533878 | 7.71439271 |
| NANOG 18700969 ChIP-ChIP MESCs Mou | 13/196 | 0.01823889 | 2.42779218 | 13.0768786 |
| AF4 28076791 ChIP-Seq SEM Human Bloc | 31/652 | 0.01823889 | 1.72226802 | 9.26776816 |
| EBNA2 21746931 ChIP-Seq IB4-LCL Huma | 48/1137 | 0.02050308 | 1.53059521 | 8.04688 |
| CTNNB1 20460455 ChIP-Seq HCT116 Hum | 37/824 | 0.02050308 | 1.62560203 | 8.53072345 |
| AHR 22903824 ChIP-Seq MCF-7 Human | 28/577 | 0.02050308 | 1.7566777 | 9.21812636 |
| SMAD1 26771354 ChIP-Seq mESC Mouse | 43/994 | 0.02050308 | 1.56710523 | 8.21644117 |
| REST 18959480 ChIP-ChIP MESCs Mouse | 67/1701 | 0.02070702 | 1.43311459 | 7.49228934 |
| GATA2 21571218 ChIP-Seq MEGAKARYO( | 40/911 | 0.02096782 | 1.58961221 | 8.26806942 |
| KDM2B 26808549 Chip-Seq K562 Human | 66/1673 | 0.02096782 | 1.43480362 | 7.46134782 |
| SMC1 22415368 ChIP-Seq MEFs Mouse | 63/1583 | 0.02096782 | 1.4465374 | 7.52204664 |
| ISL1 30127528 ChIP-Seq BE2C Human Bra | 35/776 | 0.02315316 | 1.63116985 | 8.31212263 |
| CBP 21632823 ChIP-Seq H3396 Human | 63/1593 | 0.02374259 | 1.43628024 | 7.2748178 |
| TCF3 18692474 ChIP-Seq MESCs Mouse | 42/976 | 0.02374259 | 1.5570339 | 7.87946995 |
| SMAD4 21741376 ChIP-Seq ESCs Human | 63/1595 | 0.02415445 | 1.43424488 | 7.22623267 |
| KDM2B 26808549 Chip-Seq DND41 Huma | 65/1657 | 0.02450522 | 1.42478538 | 7.14474282 |
| SOX9 26525672 Chip-Seq HEART Mouse | 53/1299 | 0.02450522 | 1.47843501 | 7.41190537 |

| | | | | |
|---|---|---|---|---|
| NANOG 18358816 ChIP-ChIP MESCs Mou | 38/865 | 0.02450522 | 1.58832255 | 7.955983 |
| JARID2 20064375 ChIP-Seq MESCs Mouse | 41/951 | 0.02464603 | 1.55915374 | 7.79327951 |
| BRD4 28847988 ChIP-Seq BCBL1 Human E | 13/206 | 0.02505914 | 2.30080343 | 11.4508473 |
| ESR1 30970003 ChIP-Seq MCF-7 Human E | 38/868 | 0.02548835 | 1.58232633 | 7.84049331 |
| SMAD2 18955504 ChIP-ChIP HaCaT Huma | 60/1513 | 0.02585579 | 1.43816322 | 7.09167913 |
| SMAD3 18955504 ChIP-ChIP HaCaT Huma | 60/1513 | 0.02585579 | 1.43816322 | 7.09167913 |
| DNAJC2 21179169 ChIP-ChIP NT2 Human | 34/758 | 0.02642621 | 1.62025239 | 7.94644519 |
| POU5F1 18700969 ChIP-ChIP MESCs Mou | 18/330 | 0.02671715 | 1.97601323 | 9.66018797 |
| LMO2 26923725 Chip-Seq HEMOGENIC-E | 60/1517 | 0.02689513 | 1.43389571 | 6.99359141 |
| SUZ12 18692474 ChIP-Seq MEFs Mouse | 43/1019 | 0.02823771 | 1.52489773 | 7.35594656 |
| SMAD2/3 21741376 ChIP-Seq ESCs Huma | 63/1612 | 0.02833769 | 1.41715651 | 6.82453654 |
| TBX2 30127528 ChIP-Seq BE2C Human Br | 38/878 | 0.02927687 | 1.56264826 | 7.4669049 |
| CREB1 20920259 ChIP-Seq GC1-SPG Mous | 90/2451 | 0.03071423 | 1.33833749 | 6.32468229 |
| IRF8 27001747 Chip-Seq BMDM Mouse | 59/1500 | 0.03123401 | 1.42416278 | 6.69976529 |
| ESR1 20056654 ChIP-Seq MCF-7 Human E | 36/827 | 0.03206749 | 1.57041719 | 7.33918653 |
| CREM 20920259 ChIP-Seq GC1-SPG Mous | 158/4643 | 0.03245521 | 1.25904104 | 5.86308142 |
| SOX2 18692474 ChIP-Seq MEFs Mouse | 61/1566 | 0.03316173 | 1.41026223 | 6.53046093 |
| NR3C1 21868756 ChIP-Seq MCF10A Hum | 38/887 | 0.0332428 | 1.54533433 | 7.14512185 |
| SOX2 18555785 ChIP-Seq MESCs Mouse | 19/365 | 0.03326947 | 1.88085429 | 8.68641835 |
| NR0B1 18358816 ChIP-ChIP MESCs Mous | 49/1210 | 0.03410757 | 1.46259931 | 6.71178431 |
| ZFX 18555785 ChIP-Seq MESCs Mouse | 87/2372 | 0.03438658 | 1.33447262 | 6.09995856 |
| TAL1 30185409 ChIP-Seq HPC Mouse Bon | 80/2154 | 0.03438658 | 1.34927054 | 6.16458192 |
| CTCF 18555785 ChIP-Seq MESCs Mouse | 46/1123 | 0.03438658 | 1.47853401 | 6.75301572 |
| PHOX2B 30127528 ChIP-Seq KELLY Huma | 71/1877 | 0.03460113 | 1.37153164 | 6.24968309 |
| RNF2 18974828 ChIP-Seq MESCs Mouse | 44/1066 | 0.0346534 | 1.48920736 | 6.76429564 |
| EZH2 18974828 ChIP-Seq MESCs Mouse | 44/1066 | 0.0346534 | 1.48920736 | 6.76429564 |
| SOX2 27498859 Chip-Seq STOMACH Mou | 61/1573 | 0.0346534 | 1.40318488 | 6.37329629 |
| SUZ12 18555785 ChIP-Seq MESCs Mouse | 37/866 | 0.03622343 | 1.53976559 | 6.91871176 |
| HNF1A 27111144 ChIP-Seq CD8+TCells M | 110/3109 | 0.0365652 | 1.29281291 | 5.79131672 |
| KLF1 20508144 ChIP-Seq FETAL-LIVER-ER | 37/868 | 0.03713949 | 1.53589457 | 6.84966166 |
| ELK1 22589737 ChIP-Seq MCF10A Human | 33/755 | 0.0378024 | 1.57421833 | 6.98595351 |
| MEIS1 26253404 ChIP-Seq OPTIC CUPS M | 57/1461 | 0.03800602 | 1.40959768 | 6.24180242 |
| NFE2L2 22581777 ChIP-Seq LYMPHOBLAS | 21/425 | 0.0380362 | 1.78137543 | 7.87905153 |
| UBF1/2 26484160 Chip-Seq FIBROBLAST | 61/1583 | 0.03808273 | 1.3931873 | 6.15446991 |
| NANOG 18692474 ChIP-Seq MESCs Mous | 80/2172 | 0.03951701 | 1.33627336 | 5.84799736 |
| RCOR3 21632747 ChIP-Seq MESCs Mouse | 79/2142 | 0.03983804 | 1.33766145 | 5.83761674 |
| MECOM 23826213 ChIP-Seq KASUMI Mo | 59/1527 | 0.03987442 | 1.39587012 | 6.08451532 |
| TBX2 30127528 ChIP-Seq KELLY Human B | 88/2425 | 0.0400291 | 1.3184691 | 5.73651916 |
| SALL4 18804426 ChIP-ChIP MESCs Mouse | 33/761 | 0.04064886 | 1.56074312 | 6.75955216 |
| KDM5B 21448134 ChIP-Seq MESCs Mous | 107/3030 | 0.04064886 | 1.28793037 | 5.5731721 |
| CEBPB 21427703 ChIP-Seq 3T3-L1 Mouse | 59/1531 | 0.04120966 | 1.39176688 | 5.99768752 |
| MYCN 28898695 ChIP-Seq NB1643 Huma | 80/2184 | 0.04301136 | 1.32773212 | 5.64452581 |
| NR3C1 34362910 ChIP-Seq WistarRat Hip | 31/709 | 0.04301136 | 1.57269357 | 6.67557613 |
| MYC 20876797 ChIP-ChIP MEDULLOBLAS | 44/1086 | 0.04301136 | 1.45903627 | 6.19311623 |
| IRF4 20064451 ChIP-Seq CD4+T Mouse | 34/795 | 0.04301136 | 1.53842536 | 6.51895289 |
| CBP 20019798 ChIP-Seq JUKART Human | 34/795 | 0.04301136 | 1.53842536 | 6.51895289 |
| NANOG 16153702 ChIP-ChIP HESCs Huma | 43/1057 | 0.04301136 | 1.46472833 | 6.2062842 |

| | | | | |
|---|---|---|---|---|
| P63 20808887 ChIP-Seq KERATINOCYTES | 61/1598 | 0.04301136 | 1.37843485 | 5.83841967 |
| POU5F1 16153702 ChIP-ChIP HESCs Huma | 21/433 | 0.04301136 | 1.74605091 | 7.38964698 |
| RUNX2 22187159 ChIP-Seq PCA Human | 97/2722 | 0.04301136 | 1.29595626 | 5.48221388 |
| NUCKS1 24931609 ChIP-Seq HEPATOCYTE | 26/570 | 0.04324574 | 1.64064171 | 6.92490457 |
| GATA3 26560356 Chip-Seq TH1 Human | 59/1539 | 0.04347683 | 1.38362695 | 5.82725142 |
| FOSL1 28411283 ChIP-Seq MDA231-LM2- | 49/1238 | 0.04387203 | 1.42596668 | 5.98706229 |
| CRX 20693478 ChIP-Seq ADULT RETINA M | 25/544 | 0.04421163 | 1.65271411 | 6.91987131 |
| TCF7 22412390 ChIP-Seq EML Mouse | 62/1635 | 0.0451385 | 1.36887061 | 5.69417347 |
| JUND 26020271 ChIP-Seq HCASMC Huma | 28/629 | 0.0451385 | 1.60026477 | 6.65461197 |
| FOXA1 33576154 ChIP-Seq Human Lung F | 131/3825 | 0.04530483 | 1.2535542 | 5.20337539 |
| GATA1 22383799 ChIP-Seq G1ME Mouse | 61/1606 | 0.04559095 | 1.37068401 | 5.67565786 |
| FOXA1 27270436 ChIP-Seq LNCaP Human | 68/1822 | 0.04575357 | 1.34850376 | 5.57359864 |
| ZFP57 27257070 Chip-Seq ESCs Mouse | 32/744 | 0.04575357 | 1.54593523 | 6.3839783 |
| CDH4 35650610 ChIP-Seq mESC Mouse St | 33/774 | 0.04662708 | 1.53229546 | 6.29283697 |
| KLF4 19030024 ChIP-ChIP MESCs Mouse | 46/1157 | 0.04765773 | 1.43063023 | 5.83861017 |
| CEBPD 23245923 ChIP-Seq MEFs Mouse | 20/413 | 0.04790067 | 1.74190419 | 7.09352193 |
| GATA3 26560356 Chip-Seq TH2 Human | 60/1583 | 0.04872278 | 1.36671807 | 5.53725348 |
| GATA1 26923725 Chip-Seq HPCs Mouse | 10/160 | 0.04934482 | 2.27020024 | 9.16038995 |

**CHEA TF binding site enrichment - DOWN in all PKMYT1i-resistant bladder cancer cell lines**

| Term | Overlap | Adjusted P-v | Odds Ratio | Combined Score |
|---|---|---|---|---|
| STAT1 19122651 ChIP-Seq HeLaS3 Humar | 245/3434 | 2.9179E-36 | 3.19491972 | 282.557436 |
| ESR1 26153859 ChIP-Seq MCF-7 Human E | 216/3597 | 1.7235E-20 | 2.44312079 | 125.653666 |
| E2F1 18555785 ChIP-Seq MESCs Mouse | 185/3015 | 6.7318E-18 | 2.40751729 | 108.479062 |
| CREM 20920259 ChIP-Seq GC1-SPG Mous | 248/4643 | 1.3098E-17 | 2.18854916 | 96.526285 |
| MYC 19030024 ChIP-ChIP MESCs Mouse | 174/2842 | 1.1258E-16 | 2.3673913 | 98.7932429 |
| AF4 26711339 ChIP-Seq SEM Human Bloc | 175/2966 | 3.0999E-15 | 2.2642223 | 86.5681402 |
| LEF1 29337183 ChIP-Seq mESC Mouse Ste | 145/2335 | 5.5724E-14 | 2.32561093 | 81.8379123 |
| KDM5B 21448134 ChIP-Seq MESCs Mouse | 172/3030 | 2.0476E-13 | 2.15039731 | 72.5863937 |
| MYC 18358816 ChIP-ChIP MESCs Mouse | 143/2369 | 7.6726E-13 | 2.24254002 | 72.4702092 |
| YY1 33199912 ChIP-Seq 293T Human Kidr | 192/3612 | 1.4576E-12 | 2.02537112 | 63.9390222 |
| MYC 30127528 ChIP-Seq KELLY Human Br | 213/4188 | 2.5081E-12 | 1.95896742 | 60.5928194 |
| EKLF 21900194 ChIP-Seq ERYTHROCYTE M | 82/1055 | 2.5504E-12 | 2.80811252 | 86.5664513 |
| FOXA1 33576154 ChIP-Seq Human Lung F | 198/3825 | 5.1227E-12 | 1.97064448 | 59.2174549 |
| MYC 18555785 ChIP-Seq MESCs Mouse | 70/852 | 1.4084E-11 | 2.94952023 | 85.4309061 |
| CTCF 33576154 ChIP-Seq Human Liver Pro | 188/3679 | 8.8779E-11 | 1.91684447 | 51.8587372 |
| NFKB1 27076634 ChIP-Seq BEAS2B Huma | 193/3823 | 1.12E-10 | 1.89717084 | 50.763176 |
| MYC 22102868 ChIP-Seq CA46 Human Blo | 133/2314 | 1.7049E-10 | 2.09174041 | 54.9636376 |
| MYC 19079543 ChIP-ChIP MESCs Mouse | 75/1013 | 2.5576E-10 | 2.63587506 | 68.0419417 |
| ASH2L 23239880 ChIP-Seq MESCs Mouse | 144/2603 | 2.6854E-10 | 2.02057249 | 51.950888 |
| JARID1A 20064375 ChIP-Seq MESCs Mous | 106/1695 | 2.7026E-10 | 2.24598328 | 57.6169234 |
| MYC 28411283 ChIP-Seq MDA231-LM2-4 | 137/2441 | 3.3007E-10 | 2.04132259 | 51.8318477 |
| NELFA 20434984 ChIP-Seq ESCs Mouse | 95/1454 | 3.3007E-10 | 2.33537565 | 59.220633 |
| FOXO3 22982991 ChIP-Seq MACROPHAGI | 103/1639 | 3.7092E-10 | 2.25166009 | 56.7349492 |
| HNF1A 27111144 ChIP-Seq CD8+TCells M | 163/3109 | 4.2872E-10 | 1.92888843 | 48.2406512 |
| ZNF217 24962896 ChIP-Seq MCF-7 Huma | 87/1289 | 4.607E-10 | 2.40347136 | 59.8387697 |
| TCFCP2L1 18555785 ChIP-Seq MESCs Moi | 95/1486 | 9.3744E-10 | 2.27759557 | 54.9975092 |

| | | | | |
|---|---|---|---|---|
| VDR 23849224 ChIP-Seq CD4+ Human | 109/1811 | 9.9457E-10 | 2.15474903 | 51.8223045 |
| SMAD2 18955504 ChIP-ChIP HaCaT Huma | 95/1513 | 2.2484E-09 | 2.23087209 | 51.6740364 |
| SMAD3 18955504 ChIP-ChIP HaCaT Huma | 95/1513 | 2.2484E-09 | 2.23087209 | 51.6740364 |
| ZFX 18555785 ChIP-Seq MESCs Mouse | 131/2372 | 2.2927E-09 | 1.99017852 | 45.9924844 |
| YAP1 20516196 ChIP-Seq MESCs Mouse | 110/1868 | 2.5742E-09 | 2.10258014 | 48.2776788 |
| FOXA1 33576154 ChIP-Seq Human Liver F | 113/1944 | 2.8788E-09 | 2.07710354 | 47.394462 |
| TRIM28 19339689 ChIP-ChIP MESCs Mou: | 125/2241 | 3.3659E-09 | 2.00200735 | 45.3063423 |
| ESR1 26153859 ChIP-Seq T47D Human Br | 91/1452 | 5.6244E-09 | 2.21705549 | 48.9685346 |
| JUN 27471255 ChIP-Seq Chondrocytes Mo | 197/4145 | 6.7828E-09 | 1.76049861 | 38.5037409 |
| NANOG 18692474 ChIP-Seq MEFs Mouse | 94/1533 | 8.1383E-09 | 2.16860731 | 46.9732671 |
| ATF3 23680149 ChIP-Seq GBM1-GSC Hum | 99/1652 | 8.7026E-09 | 2.12249671 | 45.7740497 |
| KLF4 18555785 ChIP-Seq MESCs Mouse | 106/1830 | 1.2593E-08 | 2.05438814 | 43.4912948 |
| NR3C1 27076634 ChIP-Seq BEAS2B Huma | 156/3084 | 1.2752E-08 | 1.83220728 | 38.7170761 |
| STAT5A 24692510 ChIP-Seq Epithelium M | 112/1977 | 1.297E-08 | 2.01340688 | 42.4610171 |
| ELK1 19687146 ChIP-ChIP HELA Human | 54/691 | 1.891E-08 | 2.73741677 | 56.6300976 |
| FLI1 21867929 ChIP-Seq TH2 Mouse | 91/1496 | 2.0853E-08 | 2.14237628 | 44.0590006 |
| MYCN 18555785 ChIP-Seq MESCs Mouse | 98/1668 | 2.6349E-08 | 2.07244985 | 42.0643394 |
| FOXA1 27270436 ChIP-Seq VCaP Human I | 164/3331 | 2.6349E-08 | 1.78478592 | 36.2044713 |
| E2F1 26619117 ChIP-Seq Hepatocytes Mo | 140/2717 | 3.4131E-08 | 1.84633923 | 36.9337972 |
| ESR2 21235772 ChIP-Seq MCF-7 Human | 36/370 | 4.3367E-08 | 3.43035668 | 67.7232092 |
| TCFAP2C 20176728 ChIP-ChIP TROPHOBL | 110/1980 | 4.8965E-08 | 1.96407723 | 38.4947587 |
| CREB1 23762244 ChIP-Seq HIPPOCAMPUS | 111/2006 | 5.0051E-08 | 1.95672521 | 38.2665653 |
| ELF5 23300383 ChIP-Seq T47D Human | 61/860 | 5.3089E-08 | 2.47383182 | 48.1405141 |
| PPAR 26484153 Chip-Seq NCI-H1993 Hum | 65/945 | 5.3089E-08 | 2.3997244 | 46.6906518 |
| MYCN 28898695 ChIP-Seq NB1643 Huma | 118/2184 | 5.4311E-08 | 1.91490879 | 37.1762543 |
| SMARCA4 23332759 ChIP-Seq OLIGODEN | 114/2089 | 5.9546E-08 | 1.9304518 | 37.2628783 |
| CRX 20693478 ChIP-Seq ADULT RETINA M | 45/544 | 6.851E-08 | 2.88868059 | 55.299168 |
| CREB1 20920259 ChIP-Seq GC1-SPG Mou: | 128/2451 | 7.6888E-08 | 1.85590733 | 35.2795646 |
| ENL 26711339 ChIP-Seq SEM Human Bloc | 90/1523 | 8.22E-08 | 2.07038145 | 39.180278 |
| ELK1 22589737 ChIP-Seq MCF10A Human | 55/755 | 1.1667E-07 | 2.53301258 | 47.0024992 |
| GATA4 21415370 ChIP-Seq HL-1 Mouse | 83/1374 | 1.2575E-07 | 2.1090096 | 38.9393555 |
| FLI1 21571218 ChIP-Seq MEGAKARYOCYT | 213/4775 | 1.5467E-07 | 1.64180398 | 29.9447914 |
| NANOG 18692474 ChIP-Seq MESCs Mous | 115/2172 | 2.4208E-07 | 1.86452482 | 33.1397893 |
| ESRRB 18555785 ChIP-Seq MESCs Mouse | 69/1081 | 2.8697E-07 | 2.21488335 | 38.9186246 |
| SOX2 20726797 ChIP-Seq SW620 Human | 105/1931 | 2.8697E-07 | 1.90661253 | 33.4999654 |
| TAF7L 23326641 ChIP-Seq C3H10T1-2 Mo | 44/554 | 2.9216E-07 | 2.75727484 | 48.3521975 |
| ETS1 20019798 ChIP-Seq JURKAT Human | 79/1332 | 5.916E-07 | 2.0576591 | 34.5989378 |
| TAL1 21186366 ChIP-Seq BM-HSCs Mouse | 85/1477 | 6.5087E-07 | 1.99918085 | 33.3932813 |
| BRD4 25478319 ChIP-Seq HGPS Human | 97/1770 | 7.0358E-07 | 1.9103062 | 31.7303845 |
| TTF2 22483619 ChIP-Seq HELA Human | 76/1272 | 7.6721E-07 | 2.0692032 | 34.1417537 |
| CTCF 33576154 ChIP-Seq Human HilarLN | 169/3649 | 7.6721E-07 | 1.65908664 | 27.3636761 |
| SOX2 18692474 ChIP-Seq MEFs Mouse | 88/1566 | 1.0112E-06 | 1.95064115 | 31.6047207 |
| HOXB13 26457646 ChIP-Seq LNCaP Huma | 82/1424 | 1.0421E-06 | 1.99514028 | 32.2366052 |
| FOXM1 26456572 ChIP-Seq MCF-7 Human | 89/1595 | 1.1268E-06 | 1.93664486 | 31.1123542 |
| EOMES 20176728 ChIP-ChIP TSCs Mouse | 74/1243 | 1.258E-06 | 2.05697483 | 32.7896393 |
| FOXA1 26457646 ChIP-Seq LHSAR Human | 106/2020 | 1.2673E-06 | 1.8305231 | 29.1407122 |
| NCOR 22465074 ChIP-Seq MACROPHAGE | 89/1604 | 1.3886E-06 | 1.92416992 | 30.4290754 |

| | | | | |
|---|---|---|---|---|
| YY1 23942234 ChIP-Seq MYOBLASTS AND | 70/1173 | 2.5058E-06 | 2.05501759 | 31.2572834 |
| RUNX1 27514584 ChIP-Seq MCF-7 Humar | 80/1413 | 2.8107E-06 | 1.95352087 | 29.4629914 |
| ERG 20887958 ChIP-Seq HPC-7 Mouse | 83/1486 | 2.8452E-06 | 1.92862456 | 29.0384141 |
| AR 27270436 ChIP-Seq VCaP Human Pros | 163/3565 | 2.965E-06 | 1.62395544 | 24.3629582 |
| NANOG 18347094 ChIP-ChIP MESCs Mou: | 79/1399 | 3.5955E-06 | 1.94599236 | 28.793923 |
| ESR1 30970003 ChIP-Seq MCF-7 Human E | 56/868 | 3.6143E-06 | 2.21381697 | 32.7170714 |
| TP63 23658742 ChIP-Seq EP156T Human | 139/2937 | 4.2625E-06 | 1.66276922 | 24.2781408 |
| ATF3 27146783 Chip-Seq COLON Human | 88/1628 | 4.6086E-06 | 1.86562009 | 27.0711591 |
| OCT4 18692474 ChIP-Seq MEFs Mouse | 89/1655 | 4.8423E-06 | 1.8561871 | 26.8196686 |
| TBX5 21415370 ChIP-Seq HL-1 Mouse | 85/1561 | 5.349E-06 | 1.87659504 | 26.9050603 |
| TBX2 30127528 ChIP-Seq KELLY Human B | 119/2425 | 5.6427E-06 | 1.70946202 | 24.396995 |
| POU5F1 18692474 ChIP-Seq MESCs Mous | 141/3012 | 6.0701E-06 | 1.64320338 | 23.3119328 |
| SOX2 18692474 ChIP-Seq MESCs Mouse | 114/2302 | 6.4044E-06 | 1.72118197 | 24.3058 |
| STAT3 20064451 ChIP-Seq CD4+T Mouse | 54/843 | 6.6693E-06 | 2.1921201 | 30.8420267 |
| SOX2 21211035 ChIP-Seq LN229 Gbm | 128/2673 | 6.8247E-06 | 1.67195074 | 23.4658674 |
| CEBPB 21427703 ChIP-Seq 3T3-L1 Mouse | 83/1531 | 8.2283E-06 | 1.86400668 | 25.7917047 |
| E2F4 21247883 ChIP-Seq LYMPHOBLASTC | 118/2422 | 8.5589E-06 | 1.69347882 | 23.3465234 |
| TAL1 20887958 ChIP-Seq HPC-7 Mouse | 85/1584 | 8.8894E-06 | 1.84542579 | 25.350967 |
| CNOT3 19339689 ChIP-ChIP MESCs Mous | 64/1083 | 9.5885E-06 | 2.02160357 | 27.5960141 |
| HOXC9 25013753 ChIP-Seq NEUROBLAST( | 87/1639 | 1.0009E-05 | 1.82558331 | 24.8221613 |
| CUX1 19635798 ChIP-ChIP MULTIPLE CAN | 88/1669 | 1.1254E-05 | 1.81305938 | 24.4199091 |
| MBD3 35695185 ChIP-Seq nicBasalRootG | 163/3656 | 1.19E-05 | 1.57263207 | 21.0772344 |
| FOXP3 21729870 ChIP-Seq TREG Human | 69/1210 | 1.22E-05 | 1.9506643 | 26.0748054 |
| TOP2B 26459242 ChIP-Seq MCF-7 Human | 84/1575 | 1.2437E-05 | 1.83098592 | 24.4208566 |
| KAP1 27257070 Chip-Seq ESCs Mouse | 79/1461 | 1.554E-05 | 1.8523044 | 24.2735546 |
| SFPI1 20887958 ChIP-Seq HPC-7 Mouse | 95/1861 | 1.564E-05 | 1.756535 | 22.9894168 |
| XRN2 22483619 ChIP-Seq HELA Human | 72/1296 | 1.7496E-05 | 1.89889052 | 24.6206192 |
| SOX2 27498859 Chip-Seq STOMACH Mou | 83/1573 | 2.11E-05 | 1.80721811 | 23.0754916 |
| NR3C1 23031785 ChIP-Seq PC12 Mouse | 47/730 | 2.6432E-05 | 2.19020954 | 27.4507184 |
| RELA 24523406 ChIP-Seq FIBROSARCOM/ | 63/1104 | 3.2694E-05 | 1.94221675 | 23.9106291 |
| GABP 19822575 ChIP-Seq HepG2 Human | 105/2155 | 3.3117E-05 | 1.67659182 | 20.6027768 |
| STAT5 23275557 ChIP-Seq MAMMARY-EF | 54/897 | 3.526E-05 | 2.04573567 | 24.991123 |
| GATA2 22383799 ChIP-Seq G1ME Mouse | 80/1522 | 3.5859E-05 | 1.79493984 | 21.8801172 |
| KLF4 19030024 ChIP-ChIP MESCs Mouse | 65/1157 | 3.6702E-05 | 1.91166625 | 23.2405875 |
| PPARG 20887899 ChIP-Seq 3T3-L1 Mouse | 128/2774 | 3.8094E-05 | 1.59847514 | 19.3586967 |
| TCF3 18347094 ChIP-ChIP MESCs Mouse | 83/1613 | 5.094E-05 | 1.75603241 | 20.7403449 |
| RUNX1 27514584 Chip-Seq MCF-7 Humar | 81/1568 | 5.7096E-05 | 1.76113295 | 20.5835832 |
| AR 21915096 ChIP-Seq LNCaP-1F5 Humar | 83/1619 | 5.7471E-05 | 1.74858449 | 20.4096711 |
| TBX2 35687133 ChIP-Seq MCF-7 Human E | 47/756 | 6.0625E-05 | 2.10695535 | 24.4611415 |
| KLF4 18358816 ChIP-ChIP MESCs Mouse | 67/1238 | 8.3196E-05 | 1.83606017 | 20.7025188 |
| FOSL1 28411283 ChIP-Seq MDA231-LM2- | 67/1238 | 8.3196E-05 | 1.83606017 | 20.7025188 |
| RUNX1 21571218 ChIP-Seq MEGAKARYO( | 176/4172 | 0.00010343 | 1.47807196 | 16.3312903 |
| SPI1 22096565 ChIP-ChIP GC-B Mouse | 58/1032 | 0.0001105 | 1.90140885 | 20.8637143 |
| CTCF 33576154 ChIP-Seq Human Perigast | 109/2328 | 0.0001105 | 1.60434602 | 17.5927828 |
| OLIG2 23332759 ChIP-Seq OLIGODENDRC | 84/1679 | 0.00011723 | 1.70164035 | 18.5445718 |
| GATA3 24758297 ChIP-Seq MCF-7 Humar | 82/1629 | 0.0001188 | 1.71107005 | 18.6102608 |
| SMRT 22465074 ChIP-Seq MACROPHAGE: | 80/1580 | 0.00012262 | 1.71995187 | 18.638035 |

| | | | | |
|---|---|---|---|---|
| AR 22383394 ChIP-Seq PROSTATE CANCE | 78/1540 | 0.00015436 | 1.71799742 | 18.2070359 |
| TAL1 20566737 ChIP-Seq PRIMARY FETAL | 76/1492 | 0.00016225 | 1.7265557 | 18.1974672 |
| TBX3 20139965 ChIP-Seq ESCs Mouse | 46/767 | 0.00016905 | 2.02305471 | 21.2229436 |
| JUND 26020271 ChIP-Seq SMOOTH MUS( | 79/1573 | 0.00017631 | 1.70277265 | 17.7598707 |
| NKX2-5 21415370 ChIP-Seq HL-1 Mouse | 55/980 | 0.00017631 | 1.89376838 | 19.7424944 |
| TCF3 18692474 ChIP-Seq MEFs Mouse | 39/611 | 0.00017631 | 2.15362872 | 22.4502764 |
| PGR 26153859 ChIP-Seq MCF-7 Human Bı | 48/816 | 0.00017875 | 1.98357509 | 20.6325682 |
| TBX3 20139965 ChIP-Seq MESCs Mouse | 46/770 | 0.00017875 | 2.01434773 | 20.9389045 |
| GATA4 25053715 ChIP-Seq YYC3 Human | 80/1602 | 0.000184 | 1.69300322 | 17.5291976 |
| FOXM1 32153563 ChIP-Seq SKOV3 Huma | 16/154 | 0.000184 | 3.60733549 | 37.3374696 |
| PPARA 22158963 ChIP-Seq LIVER Mouse | 78/1552 | 0.00018482 | 1.70286891 | 17.5953172 |
| MECOM 23826213 ChIP-Seq KASUMI Mo | 77/1527 | 0.00018482 | 1.70808147 | 17.6336927 |
| TCF21 26020271 ChIP-Seq SMOOTH MUS | 81/1629 | 0.00018482 | 1.6859624 | 17.4044966 |
| CDX2 20551321 ChIP-Seq CACO-2 Human | 27/360 | 0.00019412 | 2.54236609 | 26.101388 |
| EVI1 22308434 ChIP-Seq SKOV3 Human C | 56/1010 | 0.00019442 | 1.86987588 | 19.1805103 |
| MITF 21258399 ChIP-Seq MELANOMA Hu | 189/4614 | 0.0002253 | 1.43406285 | 14.4880675 |
| KLF5 25053715 ChIP-Seq YYC3 Human | 79/1591 | 0.00023764 | 1.680807 | 16.8789337 |
| MYBL2 22936984 ChIP-ChIP MESCs Mous | 72/1419 | 0.00026926 | 1.71379657 | 16.9836711 |
| SPI1 22790984 ChIP-Seq ERYTHROLEUKEI | 78/1572 | 0.00026944 | 1.67819479 | 16.6176235 |
| HSF1 23293686 ChIP-Seq STHDH STRIATA | 40/649 | 0.000274 | 2.07405196 | 20.4877727 |
| SPI1 23547873 ChIP-Seq NB4 Human | 121/2723 | 0.00031309 | 1.51963177 | 14.7976504 |
| POU5F1 18347094 ChIP-ChIP MESCs Mou | 77/1555 | 0.0003164 | 1.67310379 | 16.2626995 |
| SOX9 24532713 ChIP-Seq HFSC Mouse | 63/1203 | 0.00032262 | 1.76396903 | 17.0991951 |
| VDR 33458620 ChIP-Seq PrimaryEpithelia | 61/1154 | 0.00032396 | 1.77977419 | 17.2326052 |
| HNF4A 19761587 ChIP-ChIP CACO-2 Hum | 45/771 | 0.00033168 | 1.96158217 | 18.9206004 |
| KLF4 25985364 ChIP-Seq ATHEROSCLERO | 49/865 | 0.00033168 | 1.90411849 | 18.3654816 |
| TRP63 18441228 ChIP-ChIP KERATINOCYT | 36495 | 0.00035245 | 4.27519681 | 40.9460498 |
| DCP1A 22483619 ChIP-Seq HELA Human | 39/636 | 0.00035572 | 2.06069845 | 19.6902241 |
| SOX2 18358816 ChIP-ChIP MESCs Mouse | 37/591 | 0.00035572 | 2.10451778 | 20.1082897 |
| ARNT 22903824 ChIP-Seq MCF-7 Human | 48/846 | 0.00036896 | 1.90592522 | 18.1283941 |
| STAT5B 24692510 ChIP-Seq Epithelium M | 32/484 | 0.00039424 | 2.22432599 | 20.994698 |
| PRDM5 23873026 ChIP-Seq MEFs Mouse | 49/873 | 0.00039923 | 1.88481869 | 17.7540321 |
| HOXB7 26014856 ChIP-Seq BT474 Humar | 77/1573 | 0.00042674 | 1.65130954 | 15.4336183 |
| TP63 30713093 ChIP-Seq Epithelial Huma | 97/2100 | 0.00044125 | 1.56581794 | 14.572036 |
| AHR 22903824 ChIP-Seq MCF-7 Human | 36/577 | 0.00045756 | 2.09478298 | 19.4051692 |
| SALL4 18804426 ChIP-ChIP MESCs Mouse | 44/761 | 0.00046315 | 1.93971113 | 17.9326257 |
| GATA3 30127528 ChIP-Seq KELLY Human | 65/1277 | 0.0005132 | 1.71108192 | 15.6324591 |
| HOXB4 20404135 ChIP-ChIP EML Mouse | 61/1180 | 0.00054941 | 1.73594755 | 15.7302387 |
| CHD1 26751641 Chip-Seq LNCaP Human | 79/1647 | 0.00063893 | 1.61569452 | 14.3864939 |
| GATA2 21186366 ChIP-Seq BM-HSCs Mou | 73/1495 | 0.00068052 | 1.64202461 | 14.5070886 |
| Nerf2 26677805 Chip-Seq MACROPHAGE! | 53/993 | 0.0006992 | 1.7881459 | 15.7384841 |
| CCND1 20090754 ChIP-ChIP RETINA Mous | 72/1472 | 0.00071094 | 1.64406711 | 14.4328221 |
| ADNP 35650610 ChIP-Seq mESC Mouse S | 30/458 | 0.00071291 | 2.19773163 | 19.2736855 |
| GATA1 21571218 ChIP-Seq MEGAKARYO( | 97/2132 | 0.00072023 | 1.53835532 | 13.4643082 |
| FOXM1 26100407 CHIP-SEQ Hek293 Flp-I | 76/1577 | 0.00072023 | 1.62106982 | 14.1801502 |
| GABP 17652178 ChIP-ChIP JURKAT Huma | 37/617 | 0.00076122 | 2.00739906 | 17.4362818 |
| CEBPD 21427703 ChIP-Seq 3T3-L1 Mouse | 67/1349 | 0.0007675 | 1.66685652 | 14.4546204 |

| | | | | |
|---|---|---|---|---|
| TP53 18474530 ChIP-ChIP U2OS Human | 40/687 | 0.00076861 | 1.94828241 | 16.8806303 |
| SMARCA4 20176728 ChIP-ChIP TSCs Mou | 46/830 | 0.00079074 | 1.85420137 | 16.001833 |
| FOXA1 27197147 Chip-Seq ENDOMETRIO | 30/462 | 0.0007937 | 2.17692237 | 18.7628631 |
| TCF3 18692474 ChIP-Seq MESCs Mouse | 52/976 | 0.0007937 | 1.78326713 | 15.3620319 |
| FOXO1 32281255 ChIP-Seq Chondrocytes | 33/531 | 0.00085895 | 2.08034801 | 17.7447731 |
| STAT3 23295773 ChIP-Seq U87 Human | 115/2637 | 0.00086863 | 1.47989408 | 12.5978983 |
| PPARG 23326641 ChIP-Seq C3H10T1-2 M | 33/534 | 0.00094085 | 2.06756205 | 17.4235038 |
| CEBPB 26923725 Chip-Seq MESODERM M | 45/814 | 0.00094775 | 1.84762475 | 15.5459814 |
| NR0B1 18358816 ChIP-ChIP MESCs Mous | 61/1210 | 0.0009711 | 1.68784298 | 14.1508763 |
| CLOCK 20551151 ChIP-Seq 293T Human | 26/382 | 0.00098088 | 2.28350458 | 19.1090858 |
| GATA1 19941827 ChIP-Seq MEL86 Mouse | 73/1519 | 0.00098576 | 1.61261135 | 13.4777529 |
| RXR 22158963 ChIP-Seq LIVER Mouse | 74/1546 | 0.00099917 | 1.60636161 | 13.3948191 |
| TP53 32428506 ChIP-Seq Human Liver He | 35/584 | 0.00105801 | 2.00271552 | 16.5741027 |
| FOXA1 26769127 Chip-Seq PDAC-Cell Line | 78/1658 | 0.0011288 | 1.5792141 | 12.9508633 |
| ERA 27197147 Chip-Seq ENDOMETRIOID- | 45/822 | 0.0011288 | 1.82781496 | 14.9880684 |
| SOX2 30713093 ChIP-Seq Epithelial Huma | 50/945 | 0.00115947 | 1.7669511 | 14.4319462 |
| CTNNB1 20460455 ChIP-Seq HCT116 Hum | 45/824 | 0.00117708 | 1.82292609 | 14.8517197 |
| NFIB 24661679 ChIP-Seq LUNG Mouse | 29/453 | 0.0011793 | 2.14142367 | 17.4309287 |
| NANOG 18555785 ChIP-Seq MESCs Mous | 26/388 | 0.00118854 | 2.24494766 | 18.243977 |
| STAT1 20625510 ChIP-Seq HELA Human | 32/523 | 0.00129308 | 2.04342619 | 16.4230683 |
| STAT3 1855785 ChIP-Seq MESCs Mouse | 28/435 | 0.00134489 | 2.15231793 | 17.2022026 |
| TCF4 23295773 ChIP-Seq U87 Human | 133/3177 | 0.00136191 | 1.42345139 | 11.3494289 |
| GATA2 19941826 ChIP-Seq K562 Human | 76/1618 | 0.00136191 | 1.57434603 | 12.546352 |
| TCF21 26020271 ChIP-Seq HCASMC Huma | 57/1129 | 0.00138812 | 1.68582672 | 13.3937858 |
| FOXM1 25889361 ChIP-Seq OE33 AND U2 | 43/784 | 0.00140164 | 1.82876983 | 14.5021843 |
| POU5F1 18700969 ChIP-ChIP MESCs Mou | 23/330 | 0.001453 | 2.33694429 | 18.4357777 |
| EST1 17652178 ChIP-ChIP JURKAT Human | 35/597 | 0.00147525 | 1.95503776 | 15.3831668 |
| PPARG 19300518 ChIP-PET 3T3-L1 Mouse | 16/191 | 0.00153759 | 2.83916782 | 22.2077998 |
| FLI1 26923725 Chip-Seq MACROPHAGESS | 70/1473 | 0.00165805 | 1.58905941 | 12.3015483 |
| ZFP42 18358816 ChIP-ChIP MESCs Mouse | 52/1013 | 0.00166664 | 1.71116856 | 13.2292947 |
| RING1B 27294783 Chip-Seq NPCs Mouse | 76/1632 | 0.00168414 | 1.55895551 | 12.0283359 |
| CHD1 19587682 ChIP-ChIP MESCs Mouse | 36/626 | 0.00170539 | 1.91580976 | 14.7465887 |
| PPARD 21283829 ChIP-Seq MYOFIBROBLA | 117/2749 | 0.00170539 | 1.43882905 | 11.0689764 |
| KLF1 20508144 ChIP-Seq FETAL-LIVER-ERY | 46/868 | 0.00177795 | 1.7648675 | 13.4948713 |
| STAT3 29892750 ChIP-Seq EML Mouse Bo | 24/358 | 0.00183353 | 2.24191617 | 17.0624375 |
| NANOG 21062744 ChIP-ChIP HESCs Huma | 37/654 | 0.00190398 | 1.88330361 | 14.2528559 |
| SIN3B 21632747 ChIP-Seq MESCs Mouse | 137/3329 | 0.00208559 | 1.39675018 | 10.4364976 |
| JUN 21703547 ChIP-Seq K562 Human | 59/1206 | 0.00224555 | 1.62984117 | 12.0497401 |
| YY1 26981420 ChIP-Seq C2C12 Mouse Mu | 82/1811 | 0.00225444 | 1.51532049 | 11.1897011 |
| DMRT1 23473982 ChIP-Seq TESTES Mous | 77/1678 | 0.002267 | 1.53394366 | 11.3112741 |
| FOXP2 23625967 ChIP-Seq PFSK-1 AND SK | 40/733 | 0.00227202 | 1.81448929 | 13.367254 |
| SOX17 20123909 ChIP-Seq XEN Mouse | 65/1363 | 0.002287 | 1.59014409 | 11.6964376 |
| DACH1 20351289 ChIP-Seq MDA-MB-231 | 64/1338 | 0.00232038 | 1.59449172 | 11.6977029 |
| ESR1 21235772 ChIP-Seq MCF-7 Human | 16/200 | 0.00234753 | 2.69902913 | 19.7477437 |
| PADI4 21655091 ChIP-ChIP MCF-7 Human | 45/857 | 0.00234753 | 1.7457367 | 12.7704261 |
| NUCKS1 24931609 ChIP-Seq HEPATOCYTE | 33/570 | 0.00242155 | 1.92527352 | 14.0116033 |
| LXR 22158963 ChIP-Seq LIVER Mouse | 73/1578 | 0.00242155 | 1.54429146 | 11.2211822 |

| | | | | |
|---|---|---|---|---|
| TFAP2C 20629094 ChIP-Seq MCF-7 Huma | 52/1034 | 0.00242155 | 1.67266467 | 12.1480025 |
| VDR 24763502 ChIP-Seq THP-1 Human | 52/1034 | 0.00242155 | 1.67266467 | 12.1480025 |
| BCL6 25482012 ChIP-Seq CML-JURL-MK1 | 74/1605 | 0.00242155 | 1.53936503 | 11.1721965 |
| SPI1 26923725 Chip-Seq HPCs Mouse | 69/1473 | 0.00242155 | 1.5623856 | 11.33707 |
| GATA1 22383799 ChIP-Seq G1ME Mouse | 74/1606 | 0.00244929 | 1.53827396 | 11.1375449 |
| MEF2A 21415370 ChIP-Seq HL-1 Mouse | 38/691 | 0.00252775 | 1.82712211 | 13.1629582 |
| SMAD4 19686287 ChIP-ChIP HaCaT Huma | 22/326 | 0.00259202 | 2.25406347 | 16.1708131 |
| POU3F1 26484290 ChIP-Seq ESCss Mouse | 72/1557 | 0.00259202 | 1.54262914 | 11.0607201 |
| TP53 23651856 ChIP-Seq MEFs Mouse | 108/2541 | 0.00276003 | 1.42899204 | 10.1497675 |
| TEAD4 22529382 ChIP-Seq TROPHECTODI | 79/1748 | 0.00276694 | 1.50930643 | 10.7096919 |
| TET1 21490601 ChIP-Seq MESCs Mouse | 74/1618 | 0.00292982 | 1.52529145 | 10.7290787 |
| SOX9 25088423 ChIP-ChIP EMBRYONIC G | 74/1619 | 0.00296682 | 1.52421868 | 10.695645 |
| KDM2B 26808549 Chip-Seq K562 Human | 76/1673 | 0.00298403 | 1.51543553 | 10.6185585 |
| CEBPB 26923725 Chip-Seq HEMOGENIC-E | 68/1461 | 0.00299645 | 1.55010793 | 10.8482533 |
| REST 18959480 ChIP-ChIP MESCs Mouse | 77/1701 | 0.00301896 | 1.51026125 | 10.5514761 |
| ILF3 29590119 ChIP-Seq K562 Human Bor | 38/700 | 0.00306688 | 1.80141528 | 12.5494134 |
| TP63 22573176 ChIP-Seq HFKS Human | 135/3315 | 0.00312653 | 1.37703709 | 9.56051615 |
| PRDM14 21183938 ChIP-Seq MESCs Mou | 70/1520 | 0.00327816 | 1.53352898 | 10.5677618 |
| SPI1 23127762 ChIP-Seq K562 Human | 56/1157 | 0.00347402 | 1.60728372 | 10.9758304 |
| EGR1 20690147 ChIP-Seq ERYTHROLEUKE | 190/4931 | 0.00365431 | 1.32006875 | 8.94205379 |
| SETDB1 19884255 ChIP-Seq MESCs Mous | 68/1478 | 0.00391761 | 1.52997018 | 10.2509375 |
| WT1 25993318 ChIP-Seq PODOCYTE Hum | 120/2909 | 0.00392752 | 1.38763597 | 9.28788939 |
| GATA6 25053715 ChIP-Seq YYC3 Human | 74/1639 | 0.00394871 | 1.50305112 | 10.0459552 |
| POU5F1 16518401 ChIP-PET MESCs Mous | 43/836 | 0.00409783 | 1.70407948 | 11.3192303 |
| CJUN 26792858 Chip-Seq BT549 Human | 71/1564 | 0.00427351 | 1.50968939 | 9.95830421 |
| MEIS1 20887958 ChIP-Seq HPC-7 Mouse | 53/1092 | 0.00430131 | 1.60907234 | 10.5967061 |
| STAT3 18555785 ChIP-Seq MESCs Mouse | 68/1485 | 0.00430733 | 1.5218186 | 10.0136312 |
| CDX2 22108803 ChIP-Seq LS180 Human | 73/1619 | 0.00432289 | 1.49988585 | 9.857694 |
| SALL4 18804426 ChIP-ChIP XEN Mouse | 37/693 | 0.00448776 | 1.76765433 | 11.5324645 |
| GTF3C2 35216376 ChIP-Seq H9 Human Bo | 60/1277 | 0.00448776 | 1.55883978 | 10.1699084 |
| SOX2 18555785 ChIP-Seq MESCs Mouse | 23/365 | 0.00448776 | 2.09393095 | 13.6577632 |
| ZIC3 20872845 ChIP-ChIP MESCs Mouse | 18/259 | 0.00488995 | 2.31886889 | 14.9080116 |
| FOXO3 23340844 ChIP-Seq DLD1 Human | 32/576 | 0.00488995 | 1.83916357 | 11.8232264 |
| BACH1 22875853 ChIP-PCR HELA AND SCI | 54/1127 | 0.00498333 | 1.58727062 | 10.1674711 |
| SRF 21415370 ChIP-Seq HL-1 Mouse | 49/1001 | 0.00524011 | 1.62013575 | 10.2900715 |
| ESR1 20056654 ChIP-Seq MCF-7 Human E | 42/827 | 0.00560572 | 1.67929506 | 10.5458207 |
| CEBPB 26923725 Chip-Seq MACROPHAGE | 66/1450 | 0.0056279 | 1.50972381 | 9.4689412 |
| EWS 26573619 Chip-Seq HEK293 Human | 35/656 | 0.00585596 | 1.76374204 | 10.9850612 |
| JUN 26020271 ChIP-Seq SMOOTH MUSCL | 71/1587 | 0.00587002 | 1.48487185 | 9.23873955 |
| LUZP1 20508642 ChIP-Seq ESCs Mouse | 67/1480 | 0.00587539 | 1.50136613 | 9.33407132 |
| NEUROD2 26341353 ChIP-Seq CORTEX M | 67/1481 | 0.00594951 | 1.50022077 | 9.30224703 |
| GF1 26923725 Chip-Seq HPCs Mouse | 52/1089 | 0.00629513 | 1.57921178 | 9.69668411 |
| RCOR1 19997604 ChIP-ChIP NEURONS Mo | 52/1095 | 0.00702635 | 1.56961316 | 9.4591393 |
| WT1 20215353 ChIP-ChIP NEPHRON PRO | 60/1307 | 0.00723932 | 1.51882288 | 9.10180617 |
| SA1 22415368 ChIP-Seq MEFs Mouse | 69/1549 | 0.00737164 | 1.47588376 | 8.81204479 |
| CTCF 31629814 ChIP-Seq Hepatocytes Mo | 124/3086 | 0.00755921 | 1.34653056 | 8.00069372 |
| TRIM28 17542650 ChIP-ChIP NTERA2 Hur | 35/668 | 0.00759845 | 1.72919848 | 10.2588048 |

| | | | | |
|---|---|---|---|---|
| SETDB1 19884257 ChIP-Seq MESCs Mous | 72/1634 | 0.00773707 | 1.46026857 | 8.63134113 |
| IRF8 27001747 Chip-Seq BMDM Mouse | 67/1500 | 0.00783983 | 1.47876275 | 8.71551258 |
| FOXA2 19822575 ChIP-Seq HepG2 Human | 105/2553 | 0.00812774 | 1.3717382 | 8.0300538 |
| RUNX1 30185409 ChIP-Seq HPC Mouse Bo | 111/2724 | 0.00820885 | 1.36073786 | 7.94700271 |
| LMO2 20887958 ChIP-Seq HPC-7 Mouse | 60/1317 | 0.00832002 | 1.50590838 | 8.7689007 |
| SOX11 23321250 ChIP-ChIP Z138-A519-JV | 39/773 | 0.00842192 | 1.66383807 | 9.6620267 |
| SOX2 19030024 ChIP-ChIP MESCs Mouse | 33/625 | 0.00859426 | 1.74130391 | 10.0700942 |
| MEIS1 26253404 ChIP-Seq OPTIC CUPS M | 65/1461 | 0.00968758 | 1.47049567 | 8.32242068 |
| LMO2 26923725 Chip-Seq HEMOGENIC-E | 67/1517 | 0.0099528 | 1.46004014 | 8.21839374 |
| P63 26484246 Chip-Seq KERATINOCYTES | 71/1627 | 0.01012368 | 1.44345843 | 8.09514887 |
| E2F4 17652178 ChIP-ChIP JURKAT Human | 32/608 | 0.010187 | 1.7340347 | 9.70754945 |
| FOXA1 27270436 ChIP-Seq LNCaP Human | 78/1822 | 0.01052363 | 1.41751782 | 7.88432373 |
| REST 19997604 ChIP-ChIP NEURONS Mou | 51/1095 | 0.01064749 | 1.53523196 | 8.5154807 |
| HAND2 30127528 ChIP-Seq KELLY Human | 133/3383 | 0.01076764 | 1.31639982 | 7.28211546 |
| OLIG2 29049317 ChIP-Seq Mouse SpinalC | 65/1472 | 0.0111954 | 1.45810616 | 8.0039151 |
| SMC1 22415368 ChIP-Seq MEFs Mouse | 69/1583 | 0.01156178 | 1.43999719 | 7.85293253 |
| PPARG 20176806 ChIP-Seq 3T3-L1 Mouse | 61/1369 | 0.01199911 | 1.46972851 | 7.95520682 |
| STAT3 24763339 ChIP-Seq IMN-ESCs Mou | 62/1398 | 0.01234631 | 1.46279992 | 7.87072507 |
| ESR1 17901129 ChIP-ChIP LIVER Mouse | 21/354 | 0.01244302 | 1.95804124 | 10.5087681 |
| GATA3 21867929 ChIP-Seq TH1 Mouse | 61/1372 | 0.01244302 | 1.46612167 | 7.86668681 |
| PHOX2B 30127528 ChIP-Seq BE2C Human | 39/794 | 0.01248416 | 1.61573599 | 8.65839124 |
| JUN 26020271 ChIP-Seq HCASMC Human | 27/498 | 0.01317975 | 1.78443635 | 9.45635162 |
| GATA1 30185409 ChIP-Seq HPC Mouse Bo | 42/874 | 0.01317975 | 1.58042327 | 8.37229647 |
| GATA2 20887958 ChIP-Seq HPC-7 Mouse | 59/1324 | 0.01344541 | 1.46823715 | 7.74352872 |
| PDX1 19855005 ChIP-ChIP MIN6 Mouse | 26/476 | 0.01381288 | 1.7975731 | 9.42569115 |
| RUNX2 24764292 ChIP-Seq MC3T3 Mouse | 69/1600 | 0.01423565 | 1.42265163 | 7.4083856 |
| KAP1 22055183 ChIP-Seq ESCs Mouse | 47/1009 | 0.01423565 | 1.53178181 | 7.97514277 |
| CEBPB 20176806 ChIP-Seq MACROPHAGE | 65/1492 | 0.01443279 | 1.43606913 | 7.45209149 |
| NANOG 16518401 ChIP-PET MESCs Mous | 71/1657 | 0.01455038 | 1.41376924 | 7.32001672 |
| FOXP1 21924763 ChIP-Seq HESCs Human | 128/3273 | 0.0145896 | 1.30471732 | 6.74737986 |
| NCOR1 26117541 ChIP-Seq K562 Human | 70/1631 | 0.01475206 | 1.4156569 | 7.30057362 |
| HIF1A 21447827 ChIP-Seq MCF-7 Human | 16/248 | 0.01495744 | 2.13525276 | 10.9747229 |
| FOXA1 21572438 ChIP-Seq LNCaP Human | 68/1578 | 0.01504141 | 1.42069127 | 7.28924261 |
| FOXA1 33576154 ChIP-Seq Human HilarLI | 38/781 | 0.01542417 | 1.59807871 | 8.15379266 |
| CTCF 18555785 ChIP-Seq MESCs Mouse | 51/1123 | 0.01598811 | 1.49284773 | 7.55821894 |
| KLF6 26769127 Chip-Seq PDAC-Cell Line H | 71/1666 | 0.01608854 | 1.40508026 | 7.10031868 |
| TAL1 26923725 Chip-Seq HPCs Mouse | 62/1420 | 0.01613403 | 1.43734616 | 7.25447838 |
| SMAD1 26771354 ChIP-Seq mESC Mouse | 46/994 | 0.01675685 | 1.51989868 | 7.60847215 |
| FOXO1 23066095 ChIP-Seq LIVER Mouse | 17/276 | 0.01798059 | 2.03262142 | 10.0250575 |
| CREB1 15753290 ChIP-ChIP HEK293T Hun | 24/440 | 0.01815481 | 1.79224464 | 8.81625313 |
| CEBPB 20513432 ChIP-Seq MACROPHAGE | 64/1484 | 0.0182133 | 1.41900667 | 6.96739235 |
| MYC 19915707 ChIP-ChIP AK7 Human | 73/1732 | 0.0182133 | 1.38886042 | 6.81829511 |
| PIAS1 25552417 ChIP-Seq VCAP Human | 30/589 | 0.01874459 | 1.67105996 | 8.15013603 |
| MYB 21317192 ChIP-Seq ERMYB Mouse | 37/767 | 0.01882868 | 1.5821849 | 7.7033202 |
| BRD4 27068464 Chip-Seq AML-cells Mous | 68/1597 | 0.01882868 | 1.40154423 | 6.82018327 |
| RXRA 24833708 ChIP-Seq LIVER Mouse | 62/1434 | 0.01911819 | 1.42157332 | 6.88473033 |
| FOXO1 25302145 ChIP-Seq T-LYMPHOCYT | 68/1599 | 0.01911819 | 1.3995564 | 6.77235637 |

| | | | | |
|---|---|---|---|---|
| VDR 21846776 ChIP-Seq THP-1 Human | 32/641 | 0.01911819 | 1.63719185 | 7.92080222 |
| SMC4 20622854 ChIP-Seq HELA Human | 66/1544 | 0.01911819 | 1.40631611 | 6.80368084 |
| PHOX2B 30127528 ChIP-Seq KELLY Huma | 78/1877 | 0.01913125 | 1.36989175 | 6.62211481 |
| RAD21 21589869 ChIP-Seq MESCs Mouse | 66/1546 | 0.01947627 | 1.40425866 | 6.75494951 |
| WDR5 24793694 ChIP-Seq LNCAP Human | 30/592 | 0.01947627 | 1.66187457 | 7.99244449 |
| MYB 26560356 Chip-Seq TH2 Human | 69/1629 | 0.01947627 | 1.39393465 | 6.70004249 |
| TCF4 22108803 ChIP-Seq LS180 Human | 67/1579 | 0.02083233 | 1.39532535 | 6.60837197 |
| ELF1 17652178 ChIP-ChIP JURKAT Human | 33817 | 0.02084154 | 2.9335159 | 13.8827702 |
| AR 21909140 ChIP-Seq LNCAP Human | 15/238 | 0.02166358 | 2.08020313 | 9.75745084 |
| GFI1B 20887958 ChIP-Seq HPC-7 Mouse | 63/1475 | 0.02245442 | 1.40291373 | 6.52582042 |
| TRIM28 21343339 ChIP-Seq HEK293 Hum | 27576 | 0.02250746 | 3.16835538 | 14.7205605 |
| ESR1 22217937 ChIP-Seq MCF-7 Human E | 10/133 | 0.02304179 | 2.50716594 | 11.5781709 |
| NANOG 18700969 ChIP-ChIP MESCs Mou | 13/196 | 0.02304179 | 2.19440705 | 10.130277 |
| TBX20 22080862 ChIP-Seq HEART Mouse | 67/1589 | 0.02322189 | 1.38538125 | 6.37609329 |
| TBX20 22328084 ChIP-Seq HEART Mouse | 67/1589 | 0.02322189 | 1.38538125 | 6.37609329 |
| RARB 24833708 ChIP-Seq LIVER Mouse | 61/1424 | 0.02327489 | 1.406126 | 6.4640172 |
| OCT1 27270436 Chip-Seq PROSTATE Hum | 67/1591 | 0.02369642 | 1.38340809 | 6.33048812 |
| ETS1 21867929 ChIP-Seq TH2 Mouse | 63/1483 | 0.02443859 | 1.3943884 | 6.33344801 |
| RUNX1 22897851 ChIP-Seq JUKARTE6-1 H | 67/1594 | 0.02449783 | 1.38045805 | 6.26260449 |
| NR1I2 20693526 ChIP-Seq LIVER Mouse | 31/631 | 0.02505002 | 1.6079215 | 7.25376781 |
| NANOG 18358816 ChIP-ChIP MESCs Mou | 40/865 | 0.02536078 | 1.51339659 | 6.80407484 |
| KDM6A 18722178 ChIP-ChIP U937 AND S | 21/384 | 0.02559821 | 1.79338843 | 8.04073463 |
| RUNX1 20887958 ChIP-Seq HPC-7 Mouse | 40/870 | 0.02744077 | 1.50387408 | 6.62927691 |
| CEBPA 20513432 ChIP-Seq MACROPHAGE | 63/1493 | 0.02744077 | 1.38386587 | 6.09989185 |
| TP53 16413492 ChIP-PET HCT116 Human | 16/269 | 0.02744077 | 1.95586937 | 8.61120175 |
| SMAD3 22036565 ChIP-Seq ESCs Mouse | 39/844 | 0.02744077 | 1.51130852 | 6.65276123 |
| RCOR3 21632747 ChIP-Seq MESCs Mouse | 86/2142 | 0.02830418 | 1.32127084 | 5.7713351 |
| CEBPD 23245923 ChIP-Seq MEFs Mouse | 22/413 | 0.02852039 | 1.74452134 | 7.60162577 |
| ZBTB16 27035670 ChIP-Seq hESC Human | 28/563 | 0.02898514 | 1.62631628 | 7.05543514 |
| TCF3 18467660 ChIP-ChIP MESCs Mouse | 44/982 | 0.02965501 | 1.46512956 | 6.31834393 |
| TFAP2A 35110662 ChIP-Seq HEPM Humai | 50/1144 | 0.0297342 | 1.42996669 | 6.1586667 |
| GF1B 26923725 Chip-Seq HPCs Mouse | 55/1284 | 0.03125626 | 1.40183757 | 5.9634071 |
| SALL4 22934838 ChIP-ChIP CD34+ Human | 47/1067 | 0.03148537 | 1.4401276 | 6.11154569 |
| TAL1 30185409 ChIP-Seq HPC Mouse Bon | 86/2154 | 0.03163254 | 1.31269325 | 5.56077997 |
| NRF2 31884422 ChIP-Seq A549 Human Lu | 65/1563 | 0.0320445 | 1.36259007 | 5.75054206 |
| MYC 22102868 ChIP-Seq BL Human | 28/569 | 0.03227895 | 1.60776706 | 6.76886233 |
| CEBPB 22108803 ChIP-Seq LS180 Human | 65/1565 | 0.03267235 | 1.36062097 | 5.70792026 |
| BCL3 23251550 ChIP-Seq MUSCLE Mouse | 34/725 | 0.03293228 | 1.53147612 | 6.40810359 |
| RACK7 27058665 Chip-Seq MCF-7 Human | 69/1679 | 0.03335844 | 1.34685209 | 5.61438349 |
| ZNF263 19887448 ChIP-Seq K562 Human | 23529 | 0.03372551 | 3.1805403 | 13.2141949 |
| ESR1 22446102 ChIP-Seq UTERUS Mouse | 51/1183 | 0.03431429 | 1.40908485 | 5.82589574 |
| EP300 21415370 ChIP-Seq HL-1 Mouse | 29/600 | 0.03549399 | 1.57778871 | 6.46555976 |
| AR 19668381 ChIP-Seq PC3 Human | 108/2803 | 0.03584999 | 1.27010871 | 5.18746758 |
| AR 35650195 ChIP-Seq DSRCT Human Me | 41/916 | 0.03584999 | 1.46110528 | 5.96450017 |
| RARA 24833708 ChIP-Seq LIVER Mouse | 60/1437 | 0.03679797 | 1.3655652 | 5.53497389 |
| NFI 21473784 ChIP-Seq ESCs Mouse | 64/1549 | 0.03694219 | 1.35198062 | 5.47079904 |
| JUND 26020271 ChIP-Seq HCASMC Huma | 30/629 | 0.03730698 | 1.55615319 | 6.27730427 |

| | | | | |
|---|---|---|---|---|
| SPI1 26923725 Chip-Seq MACROPHAGESS | 61/1469 | 0.03847775 | 1.3577835 | 5.43133399 |
| RARB 27405468 Chip-Seq BRAIN Mouse | 66/1611 | 0.03940011 | 1.34029126 | 5.32585343 |
| TBP 23326641 ChIP-Seq C3H10T1-2 Mous | 29/607 | 0.03988175 | 1.55810003 | 6.16806049 |
| FOXA1 21915096 ChIP-Seq LNCaP-1F5 Hu | 67/1641 | 0.04008007 | 1.3357088 | 5.27732756 |
| EOMES 21245162 ChIP-Seq HESCs Humar | 37/820 | 0.04182194 | 1.47089428 | 5.74477351 |
| GATA3 27048872 Chip-Seq THYMUS Hum | 66/1617 | 0.04185105 | 1.33465688 | 5.20804932 |
| FOXI1 33610681 ChIP-Seq GES1 Human G | 13/216 | 0.04267342 | 1.97614685 | 7.66732704 |
| UTX 26944678 Chip-Seq JUKART Human | 66/1621 | 0.04362944 | 1.33092478 | 5.13074594 |
| PU.1 20513432 ChIP-Seq MACROPHAGES | 62/1509 | 0.04368516 | 1.34227314 | 5.16908854 |
| TET1 21451524 ChIP-Seq MESCs Mouse | 58/1400 | 0.04506368 | 1.35239692 | 5.1615129 |
| SOX11 22085726 ChIP-Seq ESNs Mouse | 61/1484 | 0.04506368 | 1.34234877 | 5.12031989 |
| ETS2 20176728 ChIP-ChIP TROPHOBLAST | 10/151 | 0.04536112 | 2.18505637 | 8.31444767 |
| AR 27623747 ChIP-Seq 22Rv1 Mouse Pros | 107/2805 | 0.04560734 | 1.25433837 | 4.7627312 |
| IRF8 21731497 ChIP-ChIP J774 Mouse | 13/219 | 0.0462112 | 1.94706315 | 7.35980601 |
| PKCTHETA 26484144 Chip-Seq BREAST Hu | 21/412 | 0.0462112 | 1.6625084 | 6.28167652 |
| E2A 27217539 Chip-Seq RAMOS-Cell Line | 65/1603 | 0.04806406 | 1.32418109 | 4.94769855 |
| CTBP1 25329375 ChIP-Seq LNCAP Human | 39/885 | 0.04860203 | 1.43488885 | 5.34151607 |

s.
