## Supplementary material for "*E2F3* amplification primes bladder cancer cells for premature mitosis": Table S6

**Supplemental table S6** Overview of antibodies used in various applications.

| **Technique** | **Name** | **Company** | **Catalog** | **RRID:** | **Dilution** |
| --- | --- | --- | --- | --- | --- |
| Immuno-blots | γ-tubulin | Sigma Aldrich | T6557 | AB_477584 | 1:5000 |
|  | GAPDH | Cell Signaling | 2118 | AB_561053 | 1:5000 |
|  | γH2Ax (s139) | Cell Signaling | 2577 | AB_2118010 | 1:1000 |
|  | CHK1 phospho s296 | Cell Signaling | 2349 | AB_2080323 | 1:1000 |
|  | CHK1 phospho s345 | Cell Signaling | 2348 | AB_331212 | 1:1000 |
|  | CHK1 | Cell Signaling | 2360 | AB_2080320 | 1:1000 |
|  | pKAP1 | Bethyl | A300-767A | AB_669740 | 1:1000 |
|  | CDK1 phospho T14 | Abcam | ab58509 | AB_2074777 | 1:1000 |
|  | CDK1 phospho Y15 | Cell Signaling | 9111 | AB_331460 | 1:1000 |
|  | CDK1 phospho T161 | Cell Signaling | 9114 | AB_2074652 | 1:1000 |
|  | CDK1 | Cell Signaling | 9116 | AB_2074795 | 1:1000 |
|  | E2F3 | Santa Cruz | sc-56665 | AB_1122397 | 1:1000 |
|  | P53 | Santa Cruz | sc-126 | AB_628082 | 1:5000 |
|  | P21 | Cell Signaling | 2947 | AB_823586 | 1:1000 |
|  | RPA | Cell Signaling | 52448 | AB_2750889 | 1:1000 |
|  | RPA-phospho S4/8 | Sanbio | A300-245A | AB_210547 | 1:1000 |
|  | RB phospho S807/811 | Cell Signaling | 8516 | AB_11178658 | 1:1000 |
|  | RB | Santa Cruz | sc-102 | AB_628209 | 1:1000 |
|  | Cyclin A2 | Santa Cruz | sc-751 | AB_631329 | 1:1000 |
|  | Cyclin B1 | Santa Cruz | sc-245 | AB_627338 | 1:500 |
|  | WEE1 phospho S642 | Cell Signaling | 4910 | AB_2215870 | 1:1000 |
|  | WEE1 | Cell Signaling | 13084 | AB_2713924 | 1:1000 |
|  | Anti-mouse HRP | Cell Signaling | 7076 | AB_330924 | 1:3000 |
|  | Anti-rabbit HRP | Cell Signaling | 7074 | AB_2099233 | 1:1000 |
| Immuno-fluorescence | γH2Ax (s139) | Cell signaling | 9718 | AB_2118009 | 1:200 |
|  | GFP (against mTurq) | Abcam | ab6673 | AB_305643 | 1:3000 |
|  | H3 | Active Motif | 39763 | AB_2650522 | 1:200 |
|  | Goat anti Mouse IgG AlexaFluor 488 | Invitrogen | A11029 | AB_2534088 | 1:250 |
|  | Goat anti Rabbit IgG AlexaFluor 647 | Invitrogen | A21244 | AB_2535812 | 1:250 |
| Flow Cytometry | Histone H3 phospho S10 | Millipore | 06-570 | AB_310177 | 1:400 |
|  | Goat anti Mouse IgG AlexaFluor 488 | Invitrogen | A11029 | AB_2534088 | 1:200 |
|  | Goat anti Rabbit IgG AlexaFluor 488 | Invitrogen | A11034 | AB_2576217 | 1:200 |
|  | Goat anti Mouse IgG AlexaFluor 647 | Invitrogen | A21235 | AB_2535804 | 1:200 |
|  | Goat anti Rabbit IgG AlexaFluor 647 | Invitrogen | A21244 | AB_2535812 | 1:200 |
| DNA Fiber Assays | Mouse Anti-BrdU | BD Biosciences | 347580 | AB_10015219 | 1:100 |
|  | Rat Anti-BrdU | Abcam | ab6326 | AB_305426 | 1:100 |
|  | Goat anti Rat IgG AlexaFluor 594 | Invitrogen | A11007 | AB_10561522 | 1:200 (UMUC3) 1:300 (RPE & T24) |
|  | Goat anti Mouse IgG AlexaFluor 488 | Invitrogen | A11029 | AB_2534088 | 1:200 (UMUC3) 1:300 (RPE & T24) |
