## Supplementary material for "*E2F3* amplification primes bladder cancer cells for premature mitosis": Table S7

**Supplemental table S7** Primers used for quantitative PCR.

| Gene | Forward primer (5’-3’) | Reverse primer (5’-3’) |  |
| --- | --- | --- | --- |
| *CCNE1* | GACACCATGAAGGAGGACGG | ATTGTCCCAAGGCTGGCTC | E2F3 targets |
| *CDC45* | CTTGAAGTTCCCGCCTATGAAG | GCATGGTTTGCTCCACTATCTC |  |
| *DBF4* | GGGCAAAAGAGTTGGTAGTGG | ACTTATCGCCATCTGTTTGGATT |  |
| *CCNE2* | TTGGCTATGCGTGAGGAAGT | TGCTCTTCGGTGGTGTCATA |  |
| *C8ORF33* | CAGAAGAAGTTCCCCTAAGCG | TGCTCCAATAGCCTGCTCTTT | RS signature |
| *DDX27* | AGGAGGCTGCGAAAAGTTAAG | GGTTCCGATTAAGCCGAGGT |  |
| *MOCS3* | TGCCCGGATCAAACACCAG | GGGACCGAGGAATCTTTGGG |  |
| *MPP6* | AGACTGGGACAATTCAGGACC | GCATTTTGGCTACCTCCTCAT |  |
| *NAT10* | GCCTCTTGTAAGAAGTGTCTCG | TCTTTTCAGAGATGCCCTCGAT |  |
| *ZNF48* | GATTGGACAAGAGGCCGACT | TCACTCCCTAGACCTGTGCG |  |
| *FOXM1* | AGACCTGTGCAGATGGTGAG | CTGATGGTCTCGAAGGCTCC | Mitotic genes |
| *CCNB1* | CTGGAAACATGAGAGCCATCC | CAGCATCTTCTTGGGCACAC |  |
| *PLK1* | GCTCATCTTGTGCCCACTG | CTTGTCCACCATAGTGCGG |  |
| *BUB1* | GAAAAGAACCCAAGAGAGGCAC | CTTTCAAAGGAACAGGAGGAGC |  |
| *AURKA* | CCAAAAGAGCAAGCAGCC | CAAAGTCTTCCAAAGCCCAC |  |
| *AURKB* | CCTCATCTCCAAACTGCTCAGGC | ATCAGGCGACAGATTGAAGGGC |  |
| *CENPE* | TATATCAAAGCCAGTTGGAGGC | TTTGCCATCTATAAGGGAGGTG |  |
| *GAPDH* | CTCTGCTCCTCCTGTTCG | GCCCAATACGACCAAATCC | Housekeeping genes |
| *ACTIN* | GATCGGCGGCTCCATCCTG | GACTCGTCATACTCCTGCTTGC |  |
| *RPS18* | AGTTCCAGCATATTTTGCGAG | CTCTTGGTGAGGTCAATGTC |  |
