## Supplementary figures and images for "*E2F3* amplification primes bladder cancer cells for premature mitosis"

### Figure S1

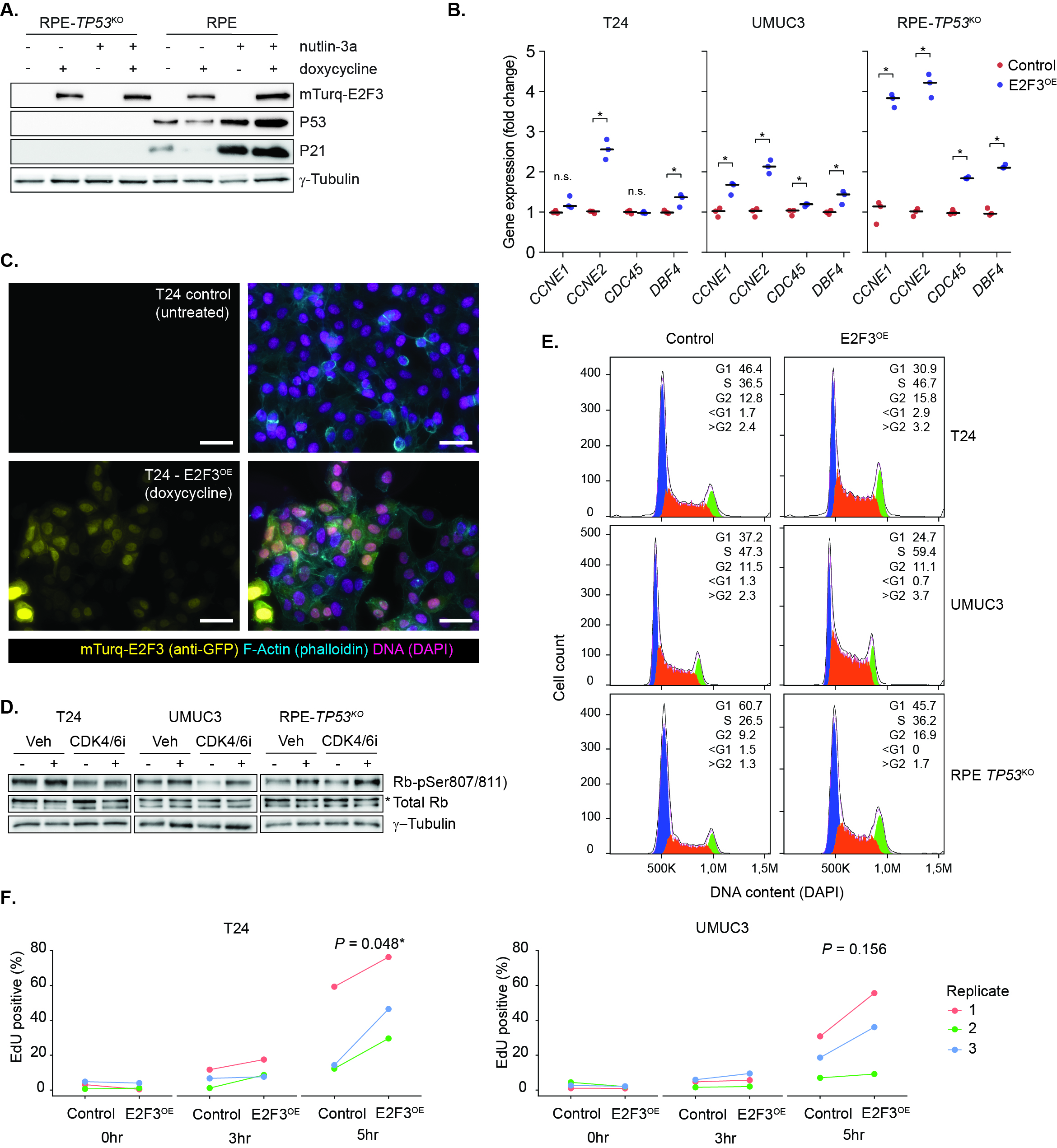

### Figure S2

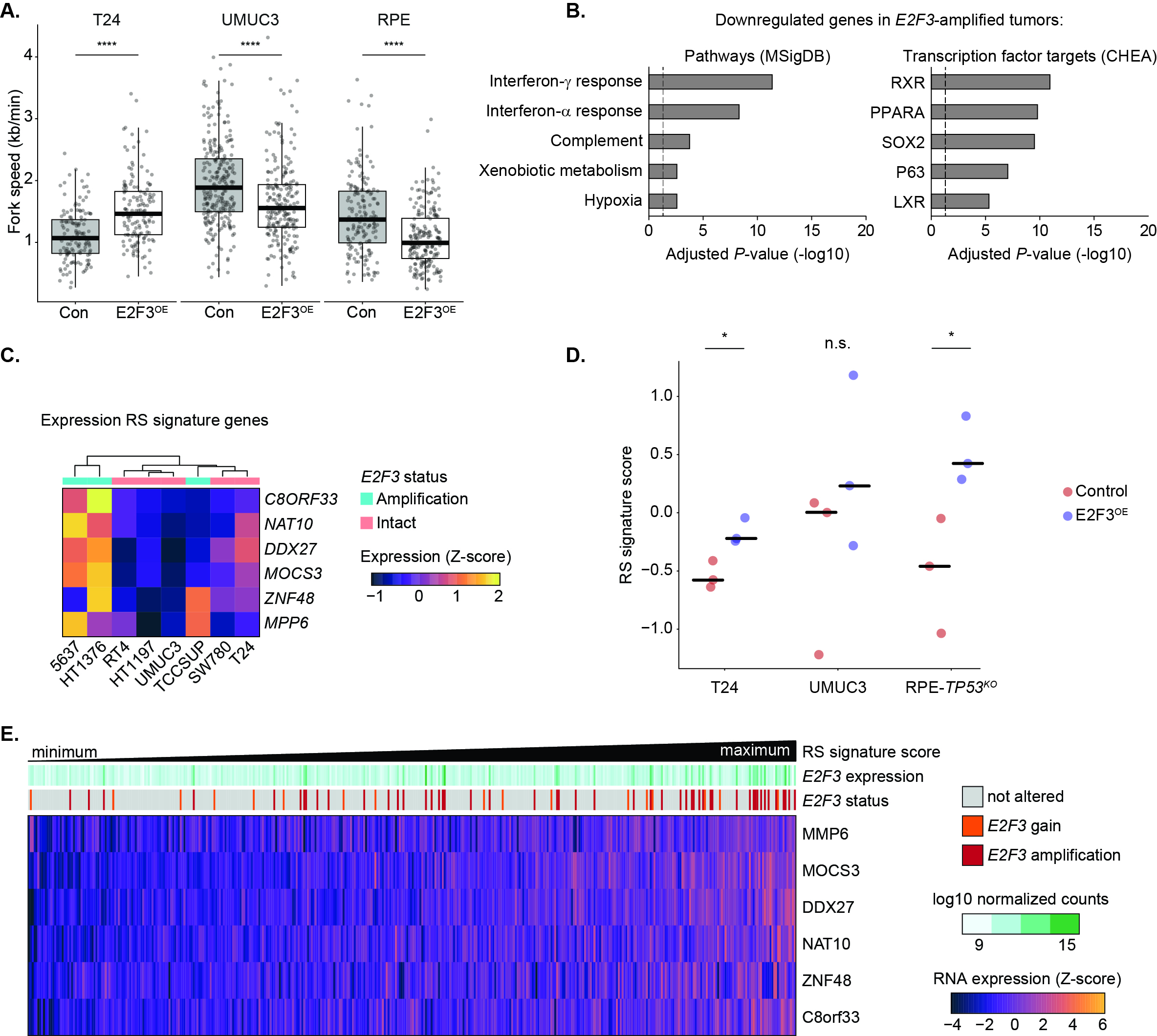

### Figure S3

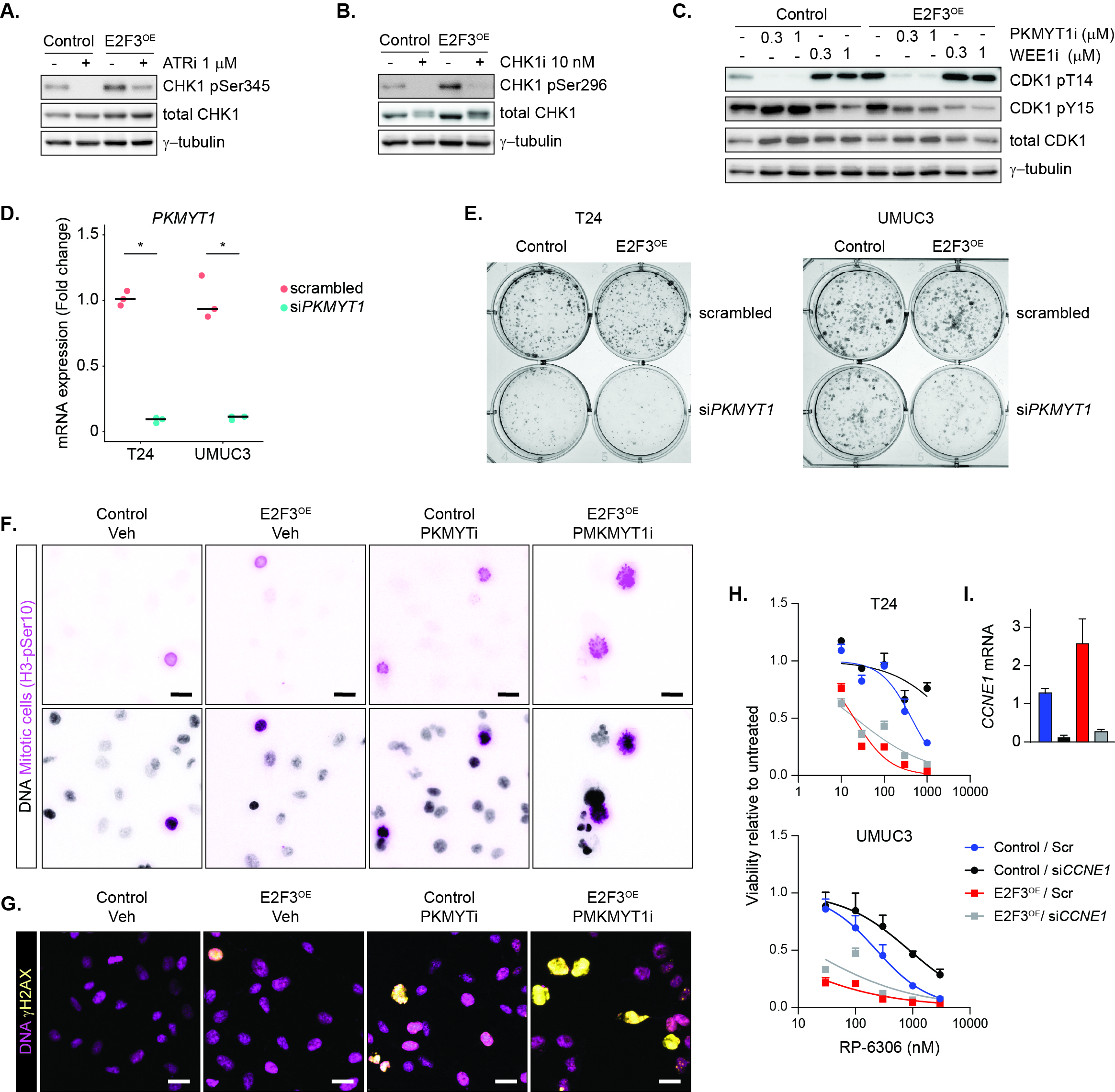

### Figure S4

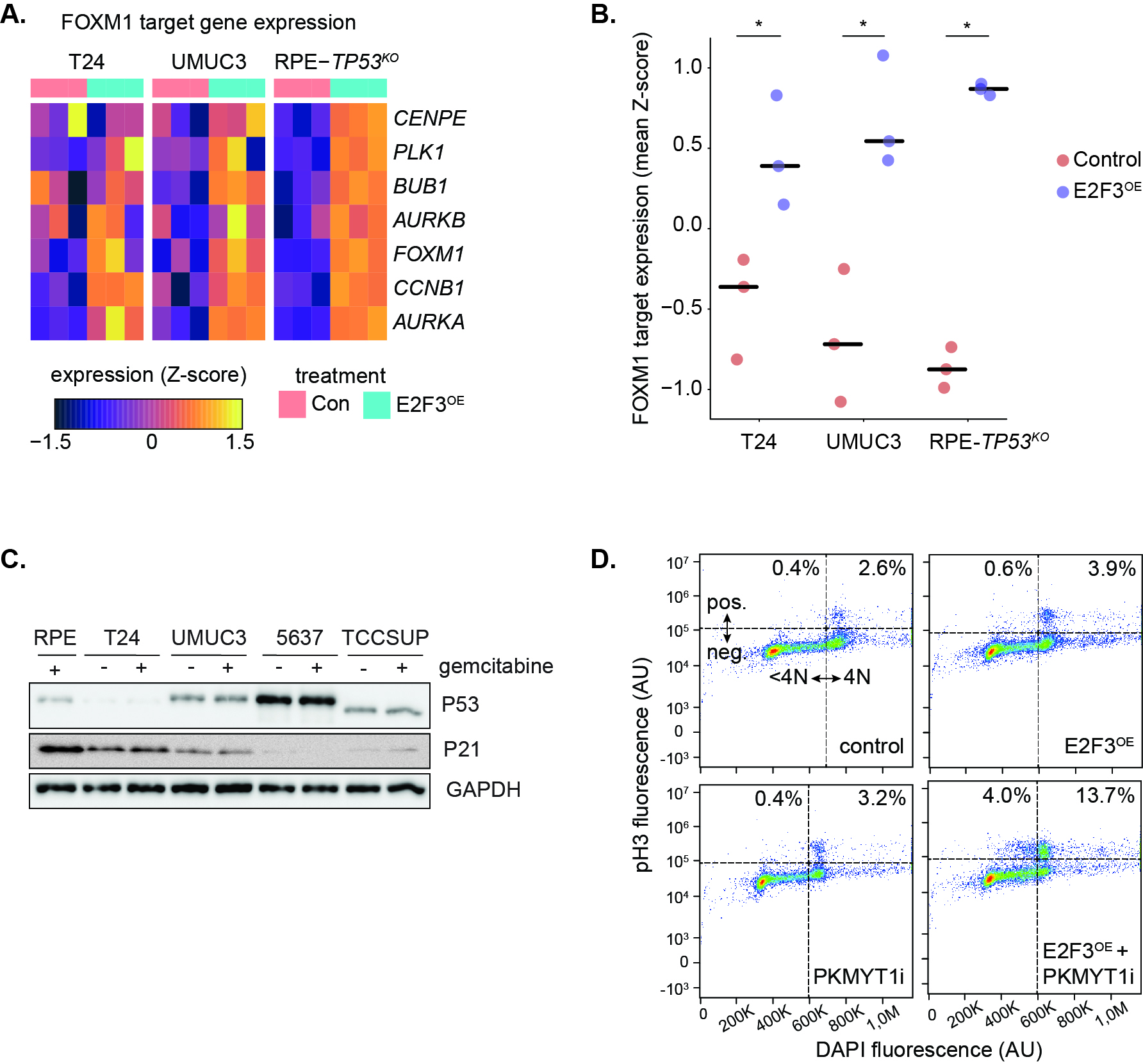

### Figure S5

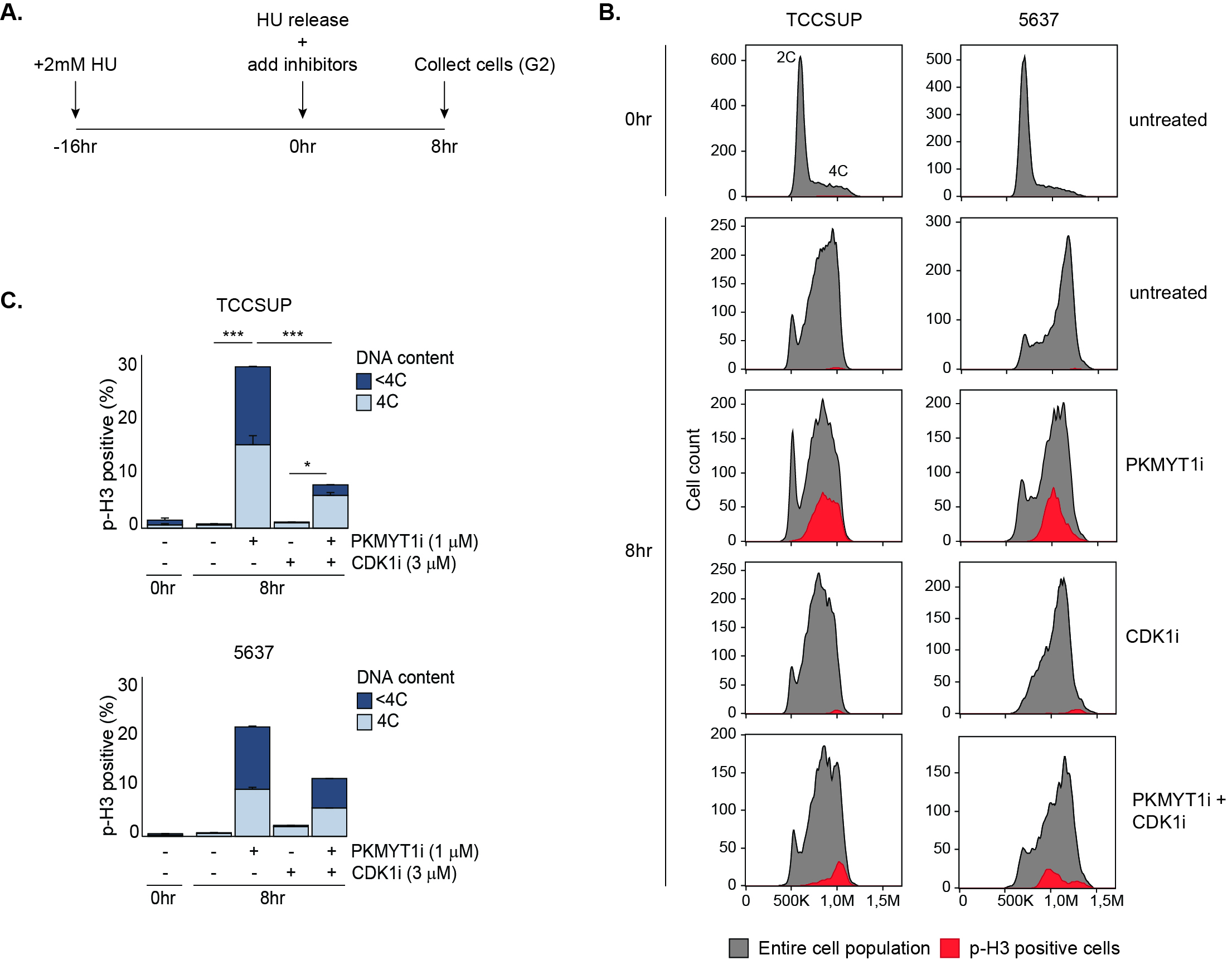

### Figure S6

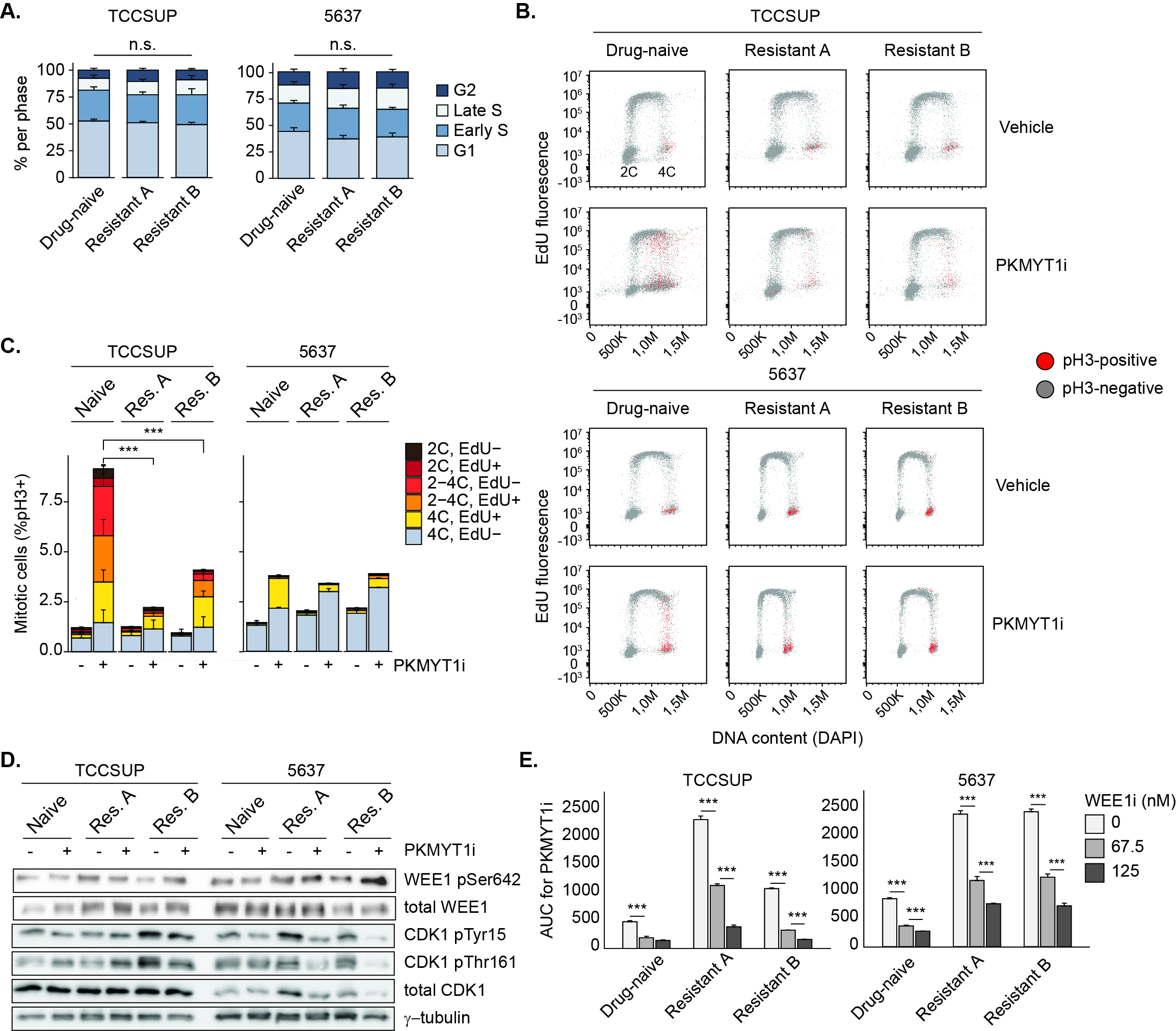
